# Cartilaginous fish inform the lineage-specific evolution and MHC association of the TLR family

**DOI:** 10.64898/2026.03.23.713620

**Authors:** Fabiana Neves, Antonio Muñoz, Ana Matos, Joana Abrantes, Raquel Xavier, Yuko Otha, Martin Flajnik, Ana Veríssimo

## Abstract

Chondrichthyans (sharks, rays and chimaeras) diverged from other vertebrate lineages over 400 million years ago and possess fully functional innate and adaptive immune systems. They comprise over 1,200 species occupying diverse aquatic habitats, from fresh to marine waters and from shallow to deep seas. Exposure to diverse pathogens through time, together with contemporary ocean warming and habitat degradation driven by climate change, has likely shaped unique immune mechanisms in these species. Toll-like receptors (TLRs) are central components of innate immunity, recognizing conserved pathogen-associated molecular patterns and initiating immune responses, yet their diversity remains poorly characterized in Chondrichthyans. Here, we characterize the TLR gene repertoire in elasmobranchs (sharks and rays) using genomic and transcriptomic data. We identify orthologs spanning all known vertebrate TLR families, including genes shared with agnathans as well as gnathostome-specific genes, and detect lineage-specific patterns of TLR duplication and loss, particularly in squaliform sharks and rays. Molecular evolution analyses indicate that most elasmobranch TLRs are under strong purifying selection, with positively selected sites in all genes, consistent with ongoing adaptation to pathogen-and environment-driven pressures. Several TLRs co-localize with genomic regions enriched in immune genes, including MHC paralogous regions, suggesting an ancestral genomic hotspot for vertebrate immunity.

## Introduction

Innate immunity, the primordial immune defense system, provides rapid and non-specific protection against invading pathogens. At its core are the pathogen recognition receptors (PRRs), which detect pathogen-associated molecular patterns (PAMPs) or released damage-associated molecular patterns (DAMPs) ^1,2^. Among these, Toll-like receptors (TLRs) are some of the most conserved and widely studied receptors found across the metazoan Tree of Life ^3,4^. These type-I transmembrane glycoproteins comprise an ectodomain (ECD) containing leucine-rich repeats (LRRs) that recognize PAMPs or DAMPs, a transmembrane (TM) domain, and an intracellular toll-interleukin 1 receptor (TIR) domain responsible for the downstream signal transduction ^1^.

TLRs are grouped in six multi- or single-gene subfamilies based on phylogenetic relationships: TLR1, TLR3, TLR4, TLR5, TLR7, and TLR11, ^2,5,6^. The different subfamilies recognize distinct antigens from viral to bacterial products, from proteins to lipopolysaccharides (LPS) to nucleic acids to lipoteichoic acid ^1,4,5,7,8^. The number of functional TLRs varies across vertebrate lineages, with several gene expansions and losses being reported. Up to date, 13 functional TLRs have been detected in mammals (1-13) ^4^, ten in birds (1-5, 7, 15, 21) ^9^, sixteen in amphibians (1-5, 7-9, 12-14, 19, 21-22) ^10^, twenty-one in fishes (1-5, 7-9, 13, 18-23, 25, 27) ^11^, and 13 in chondrichthyans (1-3, 5, 7-9, 13, 15, 21-22, 25, 27) ^4,6,12^. However, there is no consensus among studies regarding the nomenclature of TLR genes, as well as the number and composition of the different TLR subfamilies, leading to inconsistent TLR classification. Variations in gene duplication and loss events among taxonomic lineages contribute to this complexity, necessitating further comparative genomic and functional studies to unify TLR classification frameworks.

Chondrichthyans are the oldest extant jawed vertebrates to possess fully functional innate and adaptive immune systems like those in mammals ^13,14^. The group diverged from other jawed vertebrates over 400 million years ago (mya), and its living representatives are classified into the subclasses, Elasmobranchii (sharks and rays) and Holocephali (chimaeras) ^15,16^, comprising over 1,200 species with diverse biology and ecology, and inhabiting a wide range of aquatic environments ^16^. Due to their phylogenetic position, chondrichthyans provide critical insights into the early evolution of vertebrate immunity, bridging the gap between jawless vertebrates (e.g., lampreys and hagfish) and higher jawed vertebrates (mammals, reptiles, birds). Chondrichthyan immunology has largely focused in their adaptive immune system ^17–21^ with their innate immune system remaining largely unexplored, which presents a significant opportunity for further research to better understand the evolution and function of vertebrate immunity.

Previous studies on chondrichthyan TLRs report numbers ranging from nine to thirteen genes ^4,6,12,13^, though this variation likely stems from differences in the number and the species analyzed (Table 1). Indeed, comparative studies reveal species-specific and lineage-specific differences in TLR repertoires. For instance, while the elephant shark possesses orthologs for TLR14/18 and TLR15, it appears to lack orthologs for TLR21 or TLR29. In contrast, the whale shark lacks TLR15 but has TLR27 and TLR29 ^12,22^. Regardless, none of the previous studies have provided a robust assessment on TLR diversity across or within chondrichthyan lineages, as they have focused on only a limited number of species (Table 1).

**Table 1.** Overview of the studied TLRs in chondrichthyan species.

| Species | TLRs | References |
| --- | --- | --- |
| *R. typus* (whale shark) | TLR1; TLR2 (x2); TLR3; TLR7, TLR8, TLR9 (x2), TLR21 (x2), TLR22/23, TLR27, TLR28, TLR29 (x2) | <sup>4,12</sup> |
| *C. milii* (elephant shark) | TLR1, TLR2 (x2), TLR3, TLR7, TLR8, TLR9, TLR14, TLR15 TLR22/23, TLR25, TLR27 | <sup>6,12,22,23</sup> |
| *C. griseus* (bamboo shark) | TLR1, TLR2, TLR3, TLR9 | <sup>12,24</sup> |
| *A. radiata* (thorny skate) | TLR1 (x2), TLR2, TLR8, TLR9, TLR21, TLR27 | <sup>6</sup> |
| *C. carcharias* (white shark) | TLR1, TLR2 (x2), TLR3, TLR5, TLR7, TLR8, TLR9, TLR13, TLR21 (x2), TLR22 | <sup>6</sup> |

In recent years, advances in next-generation sequencing (NGS) technologies have facilitated the production of high-quality whole-genome assemblies (WGAs) for a growing number of non-model species. Currently, 37 whole genome assemblies (WGA) of elasmobranchs are available on NCBI (www.ncbi.nih.gov), providing unprecedented resources for studying their evolutionary history. This work aims to advance our understanding of key vertebrate innate immune genes by providing the first comprehensive baseline of TLR genes in elasmobranchs as ancient jawed vertebrates. Additionally, we compare chondrichthyan TLR diversity with that of other jawed and jawless vertebrates to gain deeper insights into TLR evolution. Specifically, we examine lineage-specific differences in elasmobranch TLR diversity, copy number and selective pressures, as well as conduct detailed synteny analysis and genome mapping. Furthermore, this study seeks to resolve existing inconsistencies in TLR gene nomenclature, working toward a standardized framework that facilitates comparative analyses across species.

## Material and Methods

### Bioinformatic searches

TLR sequences from reference bony vertebrates were retrieved from several lobe-finned fish (Sarcopterygii) such as: two mammals (*Homo sapiens* and TLR13 from *Mus musculus*), one bird (*Gallus gallus*), one lizard (*Pedoliscus sinensis*), one amphibian (*Xenopus tropicalis*), one lungfish (*Protopterus annectens*), and one coelacanth (*Latimeria chalumnae*); as well as ray-finned fish (Actinopterygii): two teleost fish (*Danio rerio* and *Takifugu rubripes*), the spotted gar (*Lepisostelus oculatus*), the sterlet (*Acipenser ruthenus*), and the reedfish (*Erpetoichthys calabaricus*). These were used as queries to perform exhaustive BLAST searches (until April 2025) against the genomes of elasmobranchs (sharks and rays) available on NCBI Genome Workbench (https://www.ncbi.nlm.nih.gov/tools/gbench/) (Supplementary Table 1). In turn, a total of 32 elasmobranch species from six of the nine orders of shark (Selachii) as well as all four orders of rays (Batoidea) were screened for TLR genes. Although genomes are available for other elasmobranch species, their suboptimal quality restricts its usefulness to perform detailed genomic analysis.

### Identification and classification of TLR sequences

All the retrieved putative TLR sequences (Supplementary File 1) were translated to their corresponding protein sequences in Geneious Prime (version 2024.0.7; https://www.geneious.com/). These protein sequences were evaluated for TLR-specific protein domains using HMMER (https://www.ebi.ac.uk/Tools/hmmer/search/hmmscan) and SMART (https://smart.embl-heidelberg.de/). The highly conserved TIR domain was specifically used to differentiate TLRs from other proteins containing leucine-rich repeats (LRRs). Only the full-length TLRs comprising an extracellular domain (ECD), transmembrane domain (TM), and intracellular domain (IC) were considered for further analyses.

To define TLR subfamilies and investigate the evolution of these genes in elasmobranchs, we analyzed sequences from five key species: two sharks (*Carcharodon carcharias* and *Squalus acanthias*), two rays (*Leucoraja erinaceus* and *Pristis pectinata*) and one chimaera (*Callorhinchus milii*). For broader comparative context, the TLR sequences retrieved from the other vertebrates were included (Supplementary Table 2) and translated. The corresponding protein sequences were aligned using the ClustalW alignment tool implemented in Geneious Prime (version 2024.0.7; https://www.geneious.com/). Alignments were manually corrected.

Subfamily classification and gene identity were evaluated for each validated elasmobranch TLR sequence based on synteny and phylogenetic analyses. Synteny was assessed using the NCBI Genome Data Viewer by checking at least three annotated open reading frames (ORFs) flanking the TLR genes in all studied species. In cases where the target genes or their syntenic counterparts appeared absent or were unannotated, supplementary searches were performed using BLASTn and BLASTp searches with default parameters to identify potential orthologs. For species where no TLR orthologs were identified, we examined the genomic region between the two closest syntenic genes employing Blastx searches against the NCBI non-redundant protein (nr) database.

The genomic location of TLR genes was assessed in terms of its physical proximity to key genes of the adaptive immune system (namely, Major Histocompatibility Complex, MHC; immunoglobulins, Igs; and T-cell receptor, TCR, genes). Specifically, each TLR gene location was inspected for its position relative to MHC paralogous regions, previously annotated by Veríssimo et al. (2023), in three chondrichthyan species representative of the main extant lineages: *Carcharodon carcharias* (sharks)*, Pristis pectinata* (rays) and *Callorhinchus milii* (chimaera).

Due to the high variability observed in the ectodomains across different TLRs, we focused on the conserved TIR domain to investigate the phylogenetic relationships of jawed vertebrate TLRs. TLR protein sequences from all reference bony vertebrates plus five chondrichthyans (two sharks: *C. carcharias*, *Squalus acanthias*; two rays: *Leucoraja erinacea*, *P. pectinata*; and one chimaera: *C. milii*) were aligned using ClustalW, as implemented in Geneious software. A maximum likelihood (ML) tree was reconstructed using the IQ-TREE Web Server (http://iqtree.cibiv.univie.ac.at/) based on amino acid sequence alignment (171 amino acid sites in length), using JTT+G4 as the best-fit substitution model as determined by IQ-TREE model selection ^25^. The retrieved phylogenetic tree was visualized and annotated using the interactive tree of life iTOL server version 7.1 (https://itol.embl.de/) ^26^, and rooted to the midpoint. Furthermore, to explore the evolutionary history of elasmobranch TLRs in greater depth, the same methodological approach was applied, incorporating all validated elasmobranch TLR protein sequences alongside the broader bony vertebrate dataset (Supplementary File 2), totaling 266 sequences with 173 amino acid sites, and JTT+G4 as the best-fit substitution model.

### Selective pressure analysis

We analyzed selective pressures on eleven TLRs (TLR1, TLR2, TLR3, TLR5, TLR7-9, TLR21-22, TLR27 and TLR29) retrieved for elasmobranchs using the ratio (ω) of non-synonymous substitutions per non-synonymous site (dN) to synonymous substitutions per synonymous sites (dS) (dN/dS). Due to the limited number of available sequences for some TLRs (less than 10 sequencies; e.g., TLR13 and 18), these genes were excluded from the analysis. Selective pressure analysis were conducted using the CODEML method available in the Phylogenetic Analysis by Maximum Likelihood (PAML) 4.9 package ^27,28^ and the HyPhy package implemented in the Datamonkey webserver (https://www.datamonkey.org) ^29,30^. In CODEML, site models M7 (ω ≤ 1) and M8 (ω > 1) were compared using Likelihood Ratio Tests (LRT) by calculating twice the difference in the natural logs of the likelihoods (2ΔlnL), with two degrees of freedom to determinate statistical significance (p<0.05). Sites under positive selection were identified using the Bayes Empirical Bayes (BEB) approach, with posterior probabilities > 95%. A neighbour-joining tree was constructed in MEGA X ^31^ using p-distance as the substitution model and complete deletion of gaps/missing data. Additionally, we used the site model M0 (“one-ratio model” assumes a single ω for the entire gene) to understand the overall selective pressures acting on the different TLR genes.

In the Datamonkey Web Server, all the methods can take recombination into account. Thus, prior to the selection analysis we used the GARD module available in the webserver ^32^ and the software RDP4 (version Beta 101) ^33^ to screen our sequences for recombination. Since recombination was detected for all the TLR genes, partitioned datasets from GARD were used as input for the selection models. The TLR nucleotide sequences were analyzed using the Single Likelihood Ancestor Counting (SLAC), Fixed-Effect Likelihood (FEL) ^34^, Mixed Effects Model of Evolution (MEME) ^35^ and Fast Unconstrained Bayesian AppRoximation (FUBAR) ^36^. The significance thresholds were p ≤ 0.05 for SLAC, FEL and MEME, and p ≥ 0.95 for FUBAR. To ensure robustness, only sites identified as under selection in more than one ML method were considered.

## Results and Discussion

### TLR diversity in Elasmobranchs is fully shared with other vertebrates

We screened TLR genes in the genomes of elasmobranch species spanning all major lineages of sharks and rays, making this study the most comprehensive investigation of TLRs in this ancient jawed vertebrate group to date. We identified genes encoding orthologous belonging to all six TLR subfamilies reported for vertebrates (Figure 1; more details below), including TLR1, TLR2, TLR3, TLR5, TLR7, TLR8, TLR9, TLR13, TLR18, TLR21, TLR22, TLR27, and TLR29. Additionally, we detected a new TLR, which we named/classified as TLR30. Each TLR member formed strongly supported monophyletic clades (>85% BS) with few exceptions (Fig. 1; see below). The large majority of jawed vertebrate TLRs exhibited a highly conserved synteny, except among some of the TLR11 subfamily members (Supplementary Figure 1). The repertoire of TLR genes found here in elasmobranchs is fully shared by other jawed vertebrates and partly shared by jawless vertebrates (Figure 2). Indeed, no TLR gene was found to be exclusive of elasmobranchs, unlike the pattern observed in both bony fish and jawless fish lineages. In the following sections, a detailed analysis of the elasmobranch TLR gene diversity is presented for each TLR subfamily. The results highlight a complex pattern of TLR gene gain and loss in this group, similar to observations in other taxonomically diverse groups of vertebrates ^9,37,38^.

**Figure 1.**
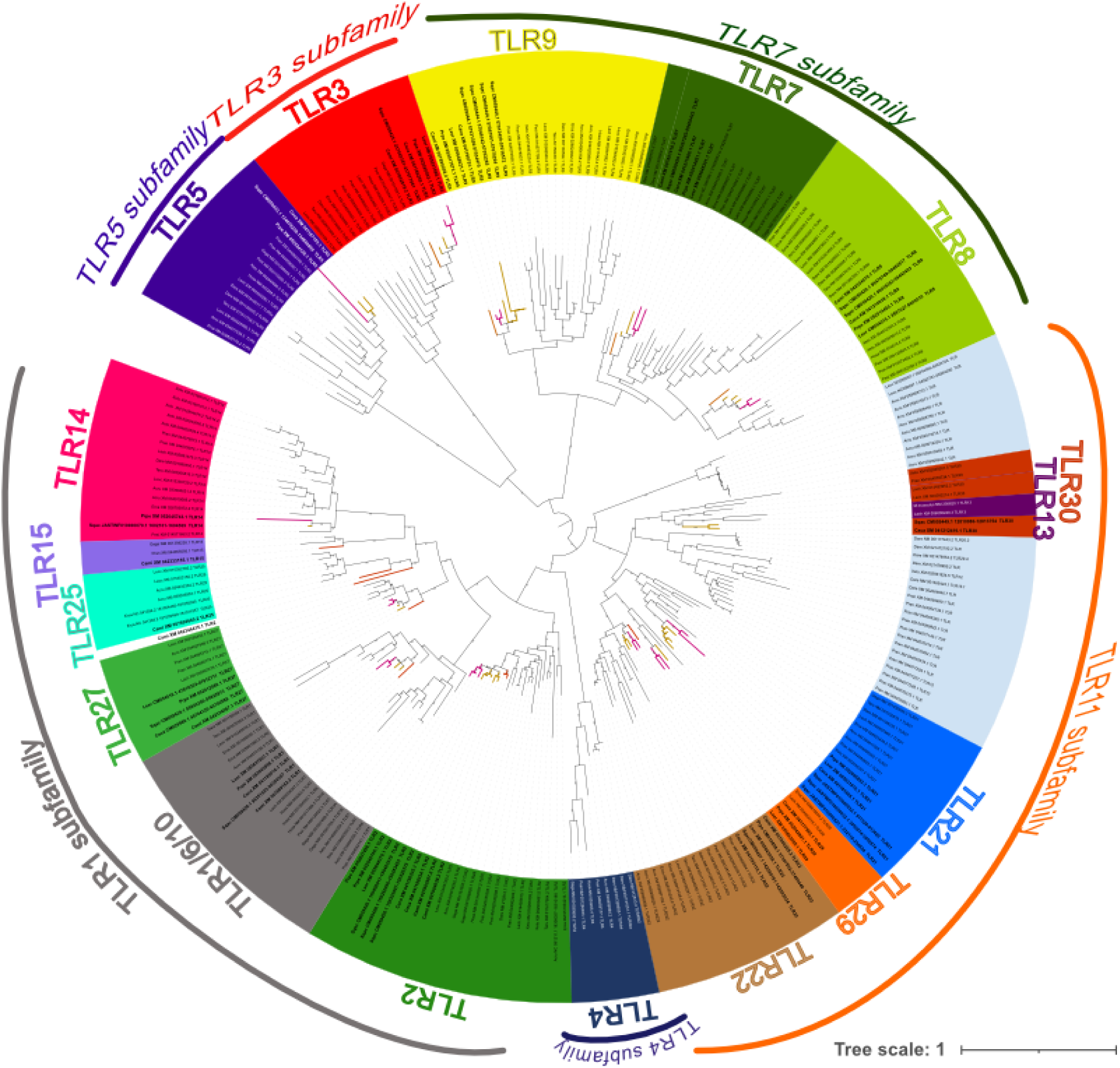
Phylogenetic tree of Toll-like receptors (TLR) using representative elasmobranch species, showing the division of TLRs into six subfamilies (TLR1, 3, 4, 5, 7 and 11). The tree was constructed using the maximum likelihood (ML) method based on 266 TIR domain sequences, aligned across 209 amino acid sites, with the JTT+G4 substitution model. The tree is midpoint-rooted. Bootstrap support values are indicated at the internal nodes. Branch colors denote major chondrichthyan lineages: orange for chimaeras, pink for batoids, and brown for sharks.

**Figure 2.**
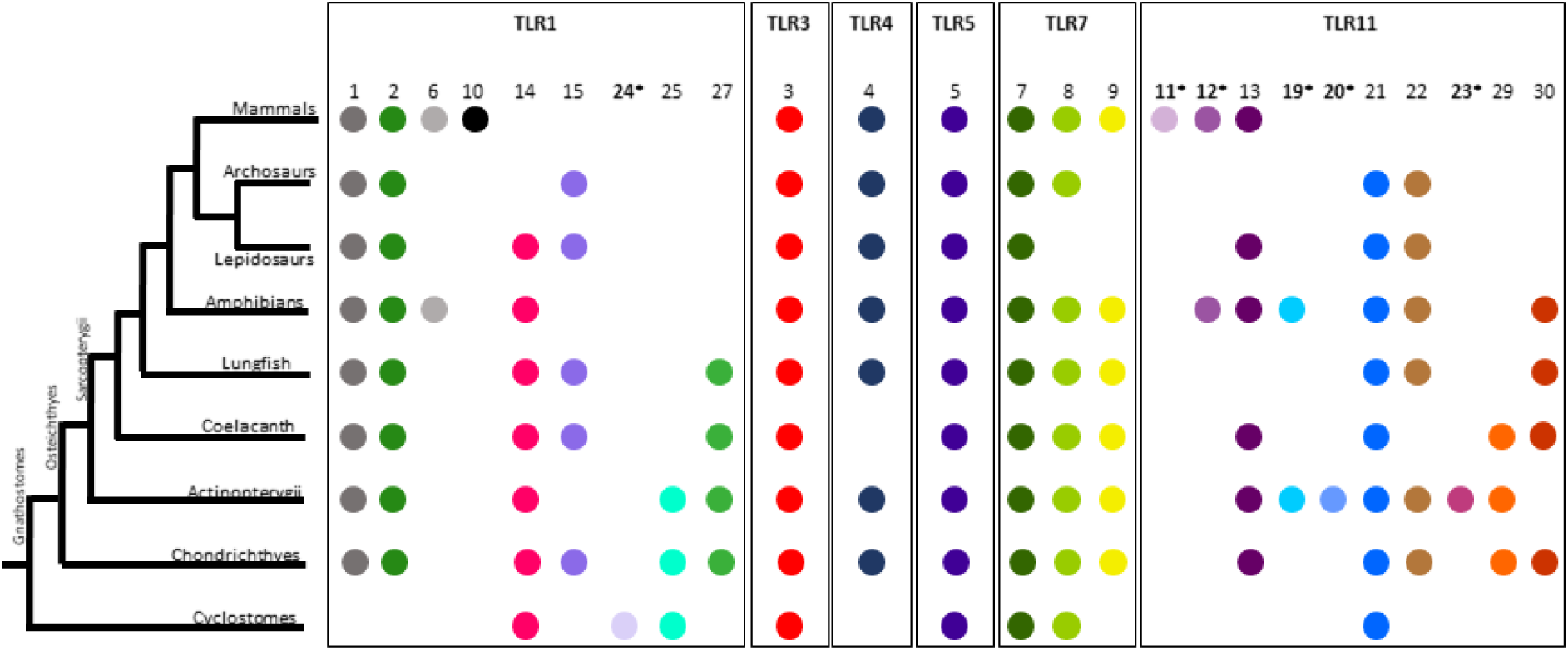
Presence/absence plot of Toll-like receptors (TLRs) across major vertebrate groups. Colored circles indicate the presence of each TLR gene in a given lineage, while the absence of a circle denotes that the gene was not detected in that group. TLRs are grouped by subfamily (e.g., TLR1, TLR3, TLR4, TLR5, TLR7, TLR11), with individual members numbered above each column. TLRs marked with an asterisk (*) were not analyzed in the present study.

Inconsistencies in the TLR gene nomenclature are common across different bibliographic sources and online databases. As previously suggested by other authors, our findings confirm that TLR14 and TLR18 are indeed orthologous genes (Figure 1, Figure 2), with variations in nomenclature across ray-finned fish (Actinopterygii), bony fish (Sarcopterygii) and jawless vertebrates (cyclostomes) ^6,7,38–40^. Therefore, we will refer to this gene as TLR14 as it was the first to be described. Furthermore, a previous study ^12^ identified a TLR30 in the whale shark (*Rhinchodon typus*) as a sister lineage of TLR9. However, we were unable to confirm the presence of such a lineage in our dataset despite extensive search efforts. Therefore, we use the designation TLR30 to refer to a newly identified putative TLR13 homolog.

#### TLR1 subfamily

The TLR1 subfamily is the largest and most evolutionary dynamic group of TLRs ^3,4^, being critical in detecting different PAMPs, such as lipopeptides, and responding to microbial infections ^1^. This subfamily has undergone significant gene expansion and loss events across various vertebrate lineages, probably facilitating class-specific adaptations that allow species to tailor their immune responses to the pathogens they encounter in their distinct ecological niches ^4^. In elasmobranchs, four TLRs belonging to the TLR1 subfamily have been identified: TLR1, TLR2, TLR14, and TLR27. TLR25 has been detected in a chimaera (*C. milii*), but it is absent from its elasmobranch relatives surveyed in this study and was not included. All members of the TLR1 subfamily present in elasmobranchs exhibit a highly conserved synteny (Supplementary Figure 1), except for TLR14 (Figure 3), supporting their evolutionary conserved nature.

**Figure 3.**
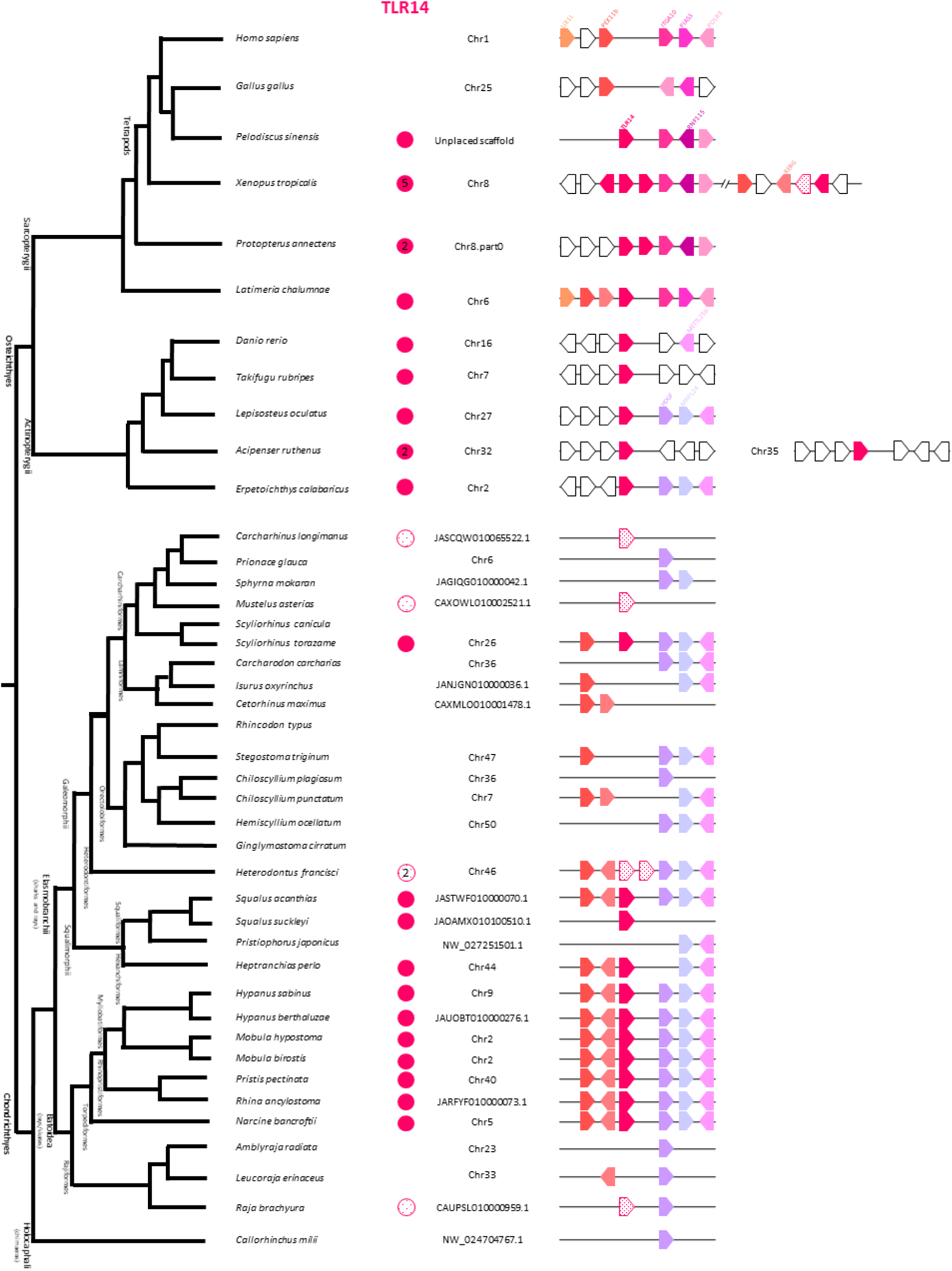
Detailed dendrogram of the *TLR14* gene across different vertebrates. The presence of TLR14 gene is marked with colored circles. When multiple gene copies were found, the gene count is indicated inside the corresponding-colored circle. Species in which these genes were not identified do not display a colored circle. These results were obtained through exhaustive BLAST searches against available vertebrate genomes. The figure is organized based on the evolutionary relationships among the analyzed vertebrates. Additionally, it incorporates a synteny analysis of the *TLR14* gene, mapping the genomic locations across the same set of vertebrates. The analyzed genomic regions are represented by horizontal lines (not to scale), with predicted genes shown as arrows, where the arrowhead indicates gene orientation. Conserved genes shared among vertebrates are consistently color-coded across species, pseudogenes are indicated by a striped background, incomplete genes indicated by a dotted background and species-specific genes represented in white. Gene names are labeled in the first vertebrate species from top to bottom.

TLR1 is present across jawed vertebrates and is generally conserved as a single-copy gene in elasmobranchs (Figure 1; Supplementary Figure 1), with the exception of some Rajiformes (or skates). The number of TLR1 gene copies varied among skate species from 1 to 3, and sequence identity among copies ranged from 63% to 98%, suggesting either different selective pressures or varying time since duplication events among copies. Notably, TLR1 copies in *A. radiata* display 63-97% amino acid (aa) identity and differences in protein length (514 aa vs. 792-793 aa), as well as variations in the number of LRRs, ranging from five to three. The expansion of TLR1 genes in Rajiformes may be a compensatory strategy for the loss of TLR5 in this batoid lineage (see below), given that both TLRs target bacterial antigens (Supplementary Figure 1).

TLR2 is also widely present across jawed vertebrates, varying from one to three copies ^6^. While the syntenic genes surrounding TLR2 are highly conserved across gnathostomes, that is not the case in teleost fishes (Supplementary Figure 1). Functionally, TLR2 exhibits a unique ability to form heterodimers with other members of the TLR1 subfamily (i.e. TLR1, TLR6 and TLR10) and, surprisingly, with TLR4, significantly expanding its antigen recognition repertoire ^41,42^. In elasmobranchs, TLR2 is generally present as a duplicated gene across most sharks, while it is mostly a single gene in rays (Supplementary Figure 1). Interestingly, the two Squaliform sharks analyzed here exhibited three tandem copies of TLR2 (Figure 1, Supplementary Figure 1) that share between 99 to 100% identity, suggesting lineage-specific differences in TLR2 copy number.

TLR14 is an ancient TLR1 subfamily member present in both jawless and jawed vertebrates but secondarily lost in birds and mammals (Figure 2). TLR14 synteny is relatively poorly conserved across vertebrates although several conserved genes are consistently present in its vicinity, including *ras-related and estrogen-regulated growth inhibitor* (RERG), *peroxisomal membrane protein 11B* (*PEX11b*) or *mitochondrial ribosomal protein L24* (*MRPL24*), among others (Figure 3). Contrary to previous work by Carlson et al. (2023), TLR14 was detected in the majority of elasmobranch orders, except in Lamniformes and Orectolobiformes. However, the retrieved TLR14 gene appears to be incomplete in some species, likely due to the low quality or fragmentation of the genome assemblies. Thus, some of these absences may need to be revisited in future higher-quality genome assemblies.

TLR27 was first described in coelacanth (*L. chalumnae*), a lobe-finned fish, and later reported in a basal Actinopterygii spotted gar (*Lepisosteus oculatus*) and in the chondrichthyan elephant shark (*Callorhinchus milii*) ^23,43^. Here, we detected a single-copy TLR27 gene in all examined elasmobranchs species, except in the blue shark (*Prionace glauca*). This finding is strongly supported by high bootstrap values (99%) in our phylogenetic analysis and is further corroborated by its location within a highly conserved syntenic cluster (Supplementary Figure 1). The TLR27 ortholog of basal ray-finned fish is located upstream of the syntenic cluster, although the gene has been lost in teleosts (Supplementary Figure 1). These findings collectively highlight the evolutionary conservation of TLR27 in cartilaginous and basal bony fishes, suggesting that it was lost independently in teleosts and tetrapods.

#### TLR3 subfamily

TLR3, the sole member of its subfamily, is a highly conserved single-copy gene in vertebrates, with no duplications or pseudogenizations, being important in the recognition of dsRNA viruses within endosomes ^4,44^. Concordantly, TLR3 is present in all elasmobranchs surveyed as a single-copy gene (Supplementary Figure 1). Its conserved synteny across jawed vertebrates supports TLR3 evolutionary stability. Additionally, phylogenetic analysis revealed that TLR3, along with TLR5, occupies a basal position relative to all other TLRs ^6^. Altogether, this emphasis the importance of TLR3 as the sole endosomal TLR that recognizes dsRNA, a key viral component.

#### TLR4 subfamily

TLR4 is another single-member TLR subfamily playing a critical role in detecting lipopolysaccharides (LPS) from Gram-negative bacteria ^45^. It has been documented across bony vertebrates ^3,4,6^, although a fragment of the TLR4 gene was also previously identified in the elephant shark genome within the same conserved syntenic cluster ^46,47^. In our analysis of elasmobranch genomes, although we confirmed the presence of a TLR4 fragment in the elephant shark, no additional TLR4 hits were identified. These observations suggest that this gene was likely present in the ancestral jawed vertebrate but that it has been secondarily lost in elasmobranchs and is retained as a pseudogene in holocephalans. Additionally, no TLR4 orthologs were retrieved for coelacanth, Japanese pufferfish (*Takifugu rubripes*) or reedfish. Despite its absence from some basal bony fish representatives, TLR4 is present in other basal taxa like the lungfish and spotted gar (Supplementary Figure 1), suggesting that TLR4 gene may have undergone several independent losses during bony fish evolution. The lack of (or very impaired) LPS responses observed in ray-finned fish and cartilaginous fish may be tied to the general absence of the *TLR4*, although such reduced response was also observed in amphibians and teleosts with a functional TLR4 gene ^48,49^. Interestingly, cluster of differentiation 14 (CD14) and lymphocyte antigen 96 (LY96, also known as MD2), which are crucial for TLR4 signaling, have been reported as absent in elephant shark and teleost genomes ^47,50^ and also seem to be absent in the other elasmobranchs analyzed in this study.

#### TLR5 subfamily

In vertebrates, TLR5 occurs either as single- or multi-copy gene, where it plays a crucial role in detecting and responding to bacterial infections, specifically in recognizing flagellin ^51^. While TLR5 synteny is highly conserved among jawed vertebrates, recent studies have shown that TLR5 has undergone multiple independent events of pseudogenization and gene loss in teleost fish as well as birds and reptiles ^4,6,9^. In elasmobranchs, TLR5 is present as a single functional gene except in Rajiformes, for which no ortholog was identified (Supplementary Figure 1). Interestingly, additional incomplete TLR5-like sequences were detected in several shark species. These results suggest that the TLR5 subfamily in elasmobranchs has undergone both lineage-specific gene gain or loss and potential functional diversification that allowed its maintenance in some species.

#### TLR7 subfamily

The TLR7 subfamily includes TLR7, TLR8 and TLR9, which share structural and functional similarities, being essential for recognizing viral nucleic acids. Specifically, all three members are endosomal receptors detecting viral ssRNA, in the case of TLR7 and TLR8, or ssDNA, in the case of TLR9 ^4,44^. Several gene gain and loss events have been observed in the TLR7 subfamily, primarily in birds and ray-finned fish ^4,9^. Our results show a strongly supported TLR7 subfamily across jawed vertebrates (Figure 1; BS: 100%) and provide evidence for the presence of all TLR7 subfamily members in elasmobranchs (Supplementary Figure 1).

TLR7 and TLR8 are located in close vicinity in all available vertebrate genomes, reflecting their close evolutionary relationship ^52^. Previous studies in a few chondrichthyans showed that TLR7 and TLR8 are typically present as single-copy genes ^6,12^. Concordantly, our results show that TLR7 is consistently found as single-copy gene across all studied elasmobranchs, except in the blue shark *Prionace glauca* and in Rajiformes where additional pseudogenized copies were found. Likewise, TLR8 was also generally present as a single-copy gene across elasmobranchs, although all squalomorph and some galeoid shark taxa had two functional copies sharing 77-87% of amino acid (aa) identity (Figure 1 and Supplementary Figure 1). In some species with duplicated TLR8 copies (e.g., *Prionace glauca, Mustelus asterias, Isurus oxyrinchus*), one copy seems to be functional while the second is pseudogenized due to early stop codons (Supplementary Figure 1). Phylogenetic analysis further indicated that TLR7 and TLR8 form a monophyletic clade (BS: 62%); however, while all jawed vertebrate TLR8 sequences cluster together with high support (BS: 95%), the TLR7 sequences from teleosts were basal to other TLR7 and all TLR8 (Figure 1).

TLR9 occurs as a multicopy gene in non-teleost ray-finned fish and shows lineage-specific losses within teleost ^6^. Among lobe-finned fish, TLR9 is generally present as a single copy gene but it appears to have been lost in the archosaur and lepidosaur lineages ^6^. A previous analysis of the whale shark (*R. typus*) genome described two TLR9 sister clades and proposed a new nomenclature for distinct TLR9 genes ^12^. This classification included the mammalian TLR9 (TLR9a), two TLR9 copies in the whale shark, one of which renamed as TLR30 (or TLR9b), and TLR31 (or TLR9c) that also includes sequences found in amphibians and testudines ^12^. Our results do not support the existence of these proposed TLR9 sister clades or distinct paralogous genes (Figure 1, Supplementary Figure 2). Instead, cartilaginous and bony fish TLR9 sequences form a strongly supported monophyletic clade (BS: 100%), sister to the jawed vertebrate TLR7 and TLR8 clades. The existence of a single TLR9 gene shared by all jawed vertebrate taxa is further supported by the conserved synteny across species (Supplementary Figure 1). Regarding the new whale shark TLR30 ^12^, we did not find any evidence of its presence in any of the elasmobranch studied here, including in whale shark. All species possessed a single-copy TLR9 ortholog, except for Squaliformes. Notably, we identified four tandemly arranged TLR9 copies in the spiny dogfish (*Squalus acanthias*) with >94% of aa identity (∼84% identity in the TIR domain), while two copies were detected on separate scaffolds in the Pacific spiny dogfish (*S. suckleyi)* sharing 99% aa identity. These findings highlight recent lineage-specific duplication events within Squaliformes, contributing to TLR9 diversity.

#### TLR11 subfamily

The TLR11 subfamily is one of the most diverse of the vertebrate TLR subfamilies, with members involved in recognizing bacterial proteins (TLR11, TLR12, TLR20) and sensing nucleic acids (TLR13, TLR20, TLR21, TLR22 and TLR23) ^4^. Additionally, the genomic regions surrounding TLR11 subfamily members are notably less conserved compared to other TLRs (Supplementary Figure 1), suggesting a dynamic evolutionary history characterized by lineage-specific adaptations. Here, we detected the presence of TLR13, TLR21, TLR22, TLR29 and TLR30 in elasmobranchs.

TLR13 is detects bacterial 23S ribosomal RNA and ssRNA genomes ^53^. It has been identified in representatives of all major lineages of jawed vertebrates, including teleosts, coelacanth, tetrapods, and the white shark ^4,6^. Here, we confirm the presence of TLR13 orthologs in various shark species (Figure 1, Supplementary Figure 1) and while these orthologs occur mostly as pseudogenes (due to early stop codons), in a few species (*Sphyrna mokaran*, *Mustelus asterias*, *Cetorhinus maximus*, *Pristiophorus japonicus*) they code for full length proteins (Supplementary Figure 1). The synteny of TLR13 found in elasmobranchs is comparable to the one in mice and in coelacanth (Supplementary Figure 1).

Our analyses detected another TLR gene clustering within the TLR13 clade (BS: 92%), showing a distinct synteny and genomic location, and suggesting that these genes may result from gene duplication. This novel gene, designated as TLR30, is present as a full-length gene in the western clawed frog (*Xenopus tropicalis)*, coelacanth (*Latimeria chalumnae)*, and in a few shark species from different orders (Supplementary Figure 1). Interestingly, in shark species bearing both TLR13 and TLR30 genes, only one occurs as a full gene while the other is pseudogenized (Supplementary Figure 1). Remarkably, this complementary pattern between the TLR13 and TLR30 gene presence is consistent in bony vertebrates, except in the coelacanth where both occur as intact genes. No TLR13 or TLR30 orthologs were identified in any of the batoid species analyzed. One putative TLR13/30-like gene was detected in basal ray-finned fish, namely in spotted gar (*Lepisosteus oculatus*) and sterlet (*Acipenser ruthenus*), whose sequences clustered together in a clade basal to TLR13 and TLR30 (Figure 1). The synteny of this TLR13/30-like gene was shared by these two species but it was distinct from either of the two related TLRs (Supplementary Figure 1).

TLR21 is an endosomal receptor important for recognizing CpG oligodeoxynucleotides and is found in all vertebrates except mammals ^6,54–56^. Unlike the majority of TLR genes, the syntenic region surrounding TLR21 is not well conserved across vertebrates ^56^, displaying greater among-taxon variability. In this study, we identified TLR21 orthologs in most elasmobranch orders, except for Torpediniformes and Myliobatiformes (Supplementary Figure 1). Notably, Squaliformes is the only order where multiple TLR21 copies were detected in different scaffolds, namely 10 copies in the spiny dogfish and 16 in the Pacific spiny dogfish. However, only three and six copies encoded full-length proteins showing high sequence conservation from >84% to >96% of aa identity for spiny dogfish and Pacific spiny dogfish, respectively. These findings highlight the dynamic evolutionary history of TLR21 within elasmobranchs, with both gene gains (in Squaliformes) and losses (particularly among batoids) depending on lineage.

TLR22 plays a key role in immune defense against bacteria and, like TLR3, is important for recognizing dsRNA viruses. However, unlike the endosomal receptor TLR3, TLR22 is located at the cell surface ^57–60^. This ancient vertebrate TLR gene is found across jawless and jawed taxa, with representatives in all major lineages but secondarily lost in birds and mammals ^4,9^, and, likely in the coelacanth (Supplementary Figure 1). While it has been reported mostly as a single-copy gene, lineage-specific duplications have also been observed, particularly among ray-finned fish ^6^. TLR22 was detected as a single-copy gene in all the studied elasmobranchs although it has been pseudogenized in certain lineages due to early stop codons, such as in the family Hemiscyliidae of Orectolobiformes, in Torpediformes and in Myliobatiformes.

TLR29 was first described in the whale shark genome ^12^ as a TLR gene most closely related to TLR21. In this study, we report the presence of a TLR29 ortholog in all elasmobranch species analyzed, as well as in coelacanth and reedfish (Figure 1), suggesting it to be restricted to basal gnathostomes. Notably, we did not find any evidence of TLR29 presence in the elephant shark genome while in spiny dogfish it appears to be a pseudogene due to early stop codons.

Based on the above results, TLR gene diversity was generally higher in sharks compared to rays, both in terms of number of TLR genes and in gene copy number. For instance, TLR13 and TLR30 were exclusively found in sharks, while gene copy number was lower in batoids for TLR5 and TLR8. Among batoids, Myliobatiformes and Torpediniformes display the lowest TLR diversity, lacking functional copies of both TLR21 and TLR22. In contrast, squaliform sharks exhibit multiple duplications of viral-recognizing TLRs, particularly TLR8, TLR9, and TLR21. Additional variability in TLR gene presence/absence was confirmed between Elasmobranchs and Holocephalans, namely for TLR4, TLR13, TLR15, TLR29, and TLR30 (Supplementary Figure 1). These findings strengthen the dynamic nature of immune system evolution in these basal jawed vertebrates, likely as the result of the different ecological and environmental pressures faced by these species, as well as their long evolutionary history.

Within each TLR subfamily, the phylogenetic relationships among sequences largely reflected the evolutionary history of elasmobranchs. Overall, the phylogenetic placement of elasmobranch TLRs reveals a heterogeneous pattern across different gene families. Elasmobranch TLR sequences occupy basal positions in the phylogenies of TLR1, TLR5, TLR8, TLR9, TLR14, and TLR27, whereas for TLR2, TLR3, TLR21, TLR22, and TLR29, they appear in more derived positions. This variation suggests complex evolutionary trajectories and potential lineage-specific adaptations.

### Elasmobranch TLRs are under purifying selection

Previous studies have highlighted differences in the selective pressures acting on viral and non-viral TLRs across various vertebrate groups, including primates, rodents, carnivores, and birds. Non-viral TLRs (TLR1, TLR2, TLR4, TLR5 and TLR6) often exhibit stronger signals of positive selection compared to viral TLRs (TLR3, TLR7, TLR8, and TLR9), which are generally subject to strong purifying selection ^4,61–65^.

In contrast, studies in amphibians have failed to identify a consistent selective pattern ^10^, while investigations in cetaceans and birds have revealed comparable levels of positive selection acting on both viral and non-viral TLRs ^38,66^. To investigate whether elasmobranchs share this dichotomy, we analyzed partitioned data obtained from GARD, which detected recombinant events in all TLR genes studied. We applied five ML methods (M7/M8, SLAC, FEL, MEME and FUBAR), considering codons putatively under selection only if identified by at least two ML methods.

Our results indicate that TLR genes in elasmobranchs are predominantly evolving under strong purifying selection, as reflected by the low ω values (dN/dS ratios) detected under the M0 model in PAML (Table 2), which range from 0.15856 for TLR21 to 0.47030 for TLR27. These low ω values suggest that the majority of non-synonymous mutations in these genes are deleterious and efficiently removed by natural selection, thereby preserving the structural and functional integrity of these receptors, ensuring their essential roles in innate immunity. Notably, the lowest ω values, were observed in viral-sensing TLRs, particularly TLR21 (ω=0.15856) and TLR7 (ω =0.16063), indicating strong evolutionary constraints. Consistent with this pattern, these viral genes also exhibit the highest numbers of negatively selected sites (NSS), with 11,12% (384 sites) and 13,91% (450 sites) for TLR21 and TLR7, respectively, possibly underscoring the high degree of functional conservation necessary to maintain their role in pathogen detection. In contrast, non-viral TLRs displayed comparatively higher ω values, indicative of weaker purifying selection and higher tolerance for non-synonymous substitutions due to their higher redundancy.

**Table 2.**
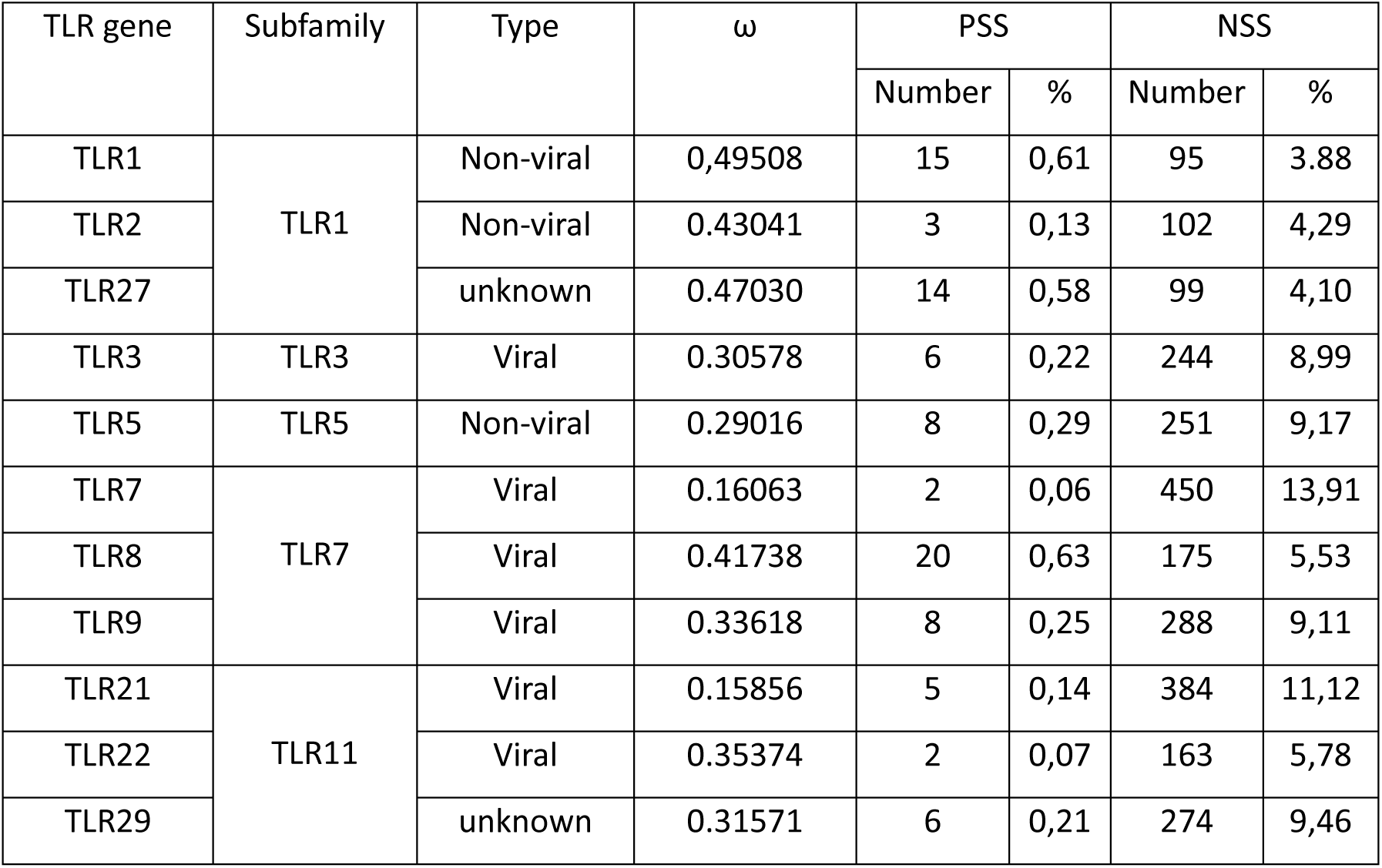
Summary of selective pressure analyses across studied genes.

| TLR gene | Subfamily | Type | $\omega$ | PSS | | NSS | |
| --- | --- | --- | --- | --- | --- | --- | --- |
|  |  |  |  | Number | % | Number | % |
| TLR1 | TLR1 | Non-viral | 0,49508 | 15 | 0,61 | 95 | 3,88 |
| TLR2 |  | Non-viral | 0.43041 | 3 | 0,13 | 102 | 4,29 |
| TLR27 |  | unknown | 0.47030 | 14 | 0,58 | 99 | 4,10 |
| TLR3 | TLR3 | Viral | 0.30578 | 6 | 0,22 | 244 | 8,99 |
| TLR5 | TLR5 | Non-viral | 0.29016 | 8 | 0,29 | 251 | 9,17 |
| TLR7 | TLR7 | Viral | 0.16063 | 2 | 0,06 | 450 | 13,91 |
| TLR8 |  | Viral | 0.41738 | 20 | 0,63 | 175 | 5,53 |
| TLR9 |  | Viral | 0.33618 | 8 | 0,25 | 288 | 9,11 |
| TLR21 | TLR11 | Viral | 0.15856 | 5 | 0,14 | 384 | 11,12 |
| TLR22 |  | Viral | 0.35374 | 2 | 0,07 | 163 | 5,78 |
| TLR29 |  | unknown | 0.31571 | 6 | 0,21 | 274 | 9,46 |

Despite the dominance of purifying selection, positively selected sites (PSS) were also identified in all TLR genes studied (Table 3, Supplementary Table 3), suggesting that certain residues have evolved to adapt to environmental or pathogen-driven pressures. The number of PSS varied across genes, with TLR8 showing the highest number of PSS (0,63%, 20 sites), all located in the extracellular domain, which likely reflects adaptation to recognize specific pathogen-derived ligands. Conversely, genes like TLR7 and TLR22 which recognize dsRNA, a key viral component, exhibit only two PSS (0,06% and 0,07%, respectively) in the extracellular domain.

**Table 3.** Chromosome location and association to MHC or MHC paralogous regions (1, 9 or 19) across representative taxa of vertebrate lineages. Lobe-finned fish: frog (*Xenopus laevis)*, coelacanth (*Latimeria chalumnae)*; ray-finned fish: reedfish (*Erpetoichthys calabaricus*), spotted gar (*Lepisosteus osseus*); cartilaginous fish: white shark (*Carcharodon carcharias*), sawfish (*Pristis pectinata*), elephant shark (*Callorhinchus milii*); agnathans: lamprey (*Petromyzon marinus*). MHC paralogs (MHCp) are highlighted in bold. For further details, please see Supplemental Figure 3.

| TLR gene | Sub-family | Frog | Coelacanth | Reedfish | Gar | White shark | Sawfish | Elephant shark | Lamprey |
| --- | --- | --- | --- | --- | --- | --- | --- | --- | --- |
| TLR1 | TLR1 | 1 | chr1 | chr5 | chr1 | chr1 | chr2 | NW_024704742.1 | - |
| TLR2 | TLR1 | 1 | chr1 | chr11 | chr1 | chr1 | chr2 | NW_024704742.1 | - |
| TLR18 | TLR1 | 8 **XNC** | chr6 (**MHCp9**) | chr2 [**MHCp14**] | **chr27 [MHCp14]** | - | chr40 | - | chr4/57/82 |
| TLR25 | TLR1 | - | - | chr1 | chr 7 | - | - | NW_024704746.1 | chr4/5/42 |
| TLR27 | TLR1 | - | chr6 (**MHCp1**) | chr10 [**MHCp1**] | chr9 [**MHCp1**] | **16 (MHCp1)** | chr3<br>(**MHCp1**) | NW_024704746.1 (**MHCp1**) | - |
| TLR13 | TLR11 | - | chr15(1) [**Xq equiv**] | - | **chr2 \*** | **9 [Xq equiv]** | - | - | - |
| TLR30 | TLR11 | 7 | chr6 |  |  | chr19 [**MHC**] | - | - | - |
| TLR21 | TLR11 | 7 | chr25 | chr17 | - | chr36 | chr40 | - | chr26/43/79 |
| TLR22 | TLR11 | 3 | chr16 | chr11 | chr11 | **8** | chr4 | NW_024704754.1 | chr61 |
| TLR29 | TLR11 | - | chr9 | chr12 [**b2m-MHC14**] | - | chr31 | chr33 |  | - |
| TLR30 | TLR3 | 1 | chr1 | chr5 | chr1 | chr4 | chr7 | unplaced scaffold | chr8 |
| TLR4 | TLR4 | pseudo? |  | - | **chr24 [MHCp9]** | - |  | NW_024704749.1 (**MHCp9**) | - |
| TLR5 | TLR5 | 5 | chr3 | chr1/chr15 | chr2 | chr5 | chr10 | - | chr6/29 |
| TLR7 | TLR7 | 2 | chr4 | chr4 | chr13 | chr18 | chr4 | NW_024704745.1 | chr3/24 |
| TLR8 | TLR7 | 2 | chr4 | chr4 | chr13/chr9 | chr18 | chr4 | NW_024704745.1 | chr3/24 |
| TLR9 | TLR7 | 4 (**MHCp1**) | chr14 | chr18 | chr4 | chr7 | chr6 | NW_024704753.1 | - |

\*next to MHC W-type syntenic genes.

The balance between positive and negative selective forces likely is a consequence of the evolutionary pressures faced by elasmobranchs in their aquatic environment, where exposure to a wide array of pathogens necessitates both conservation and adaptability in immune responses. Future research focusing on the functional implications of the positively selected sites could provide insights into the specific adaptations in elasmobranch immunity.

### TLR genes of gnathostomes map to the MHC and related paralogs

Most of the TLR genes shared by the main lineages of jawed vertebrates, including those shared also with Agnathans (i.e., lampreys and hagfish), consistently map to the same genomic regions in the cartilaginous and basal bony fish taxa (Supplementary Figure 1). Strikingly, there is a clear association of the vertebrate TLR18 gene to human chromosomes (huchr) 9/12/14 equivalent in bony vertebrates; and some gnathostome-specific TLRs shared by all main lineages also consistently mapped to other MHC paralogs (i.e., TLR4, TLR13/30, and TLR27). TLR13 maps to the huchr Xq paralog equivalent (*sensu* ^67^) in elasmobranchs and lobe-finned fish. This paralogous region has been suggested to be associated with the MHC proper (i.e., huchr 6 equivalent) based on their close location in *Xenopus* (i.e., chromosome 8L, MHC: 60-115 Mbp; Xq equivalent: 40-70 Mbp). Such association is supported by the close phylogenetic relationship between TLR13 and TLR30 shown here, and the location of TLR30 in the MHC of sharks. Moreover, while phylogenetically more diverged, TLR13 of spotted gar, a basal ray-finned fish, maps next to the MHC W-type syntenic genes that originally mapped to the MHC region ^68,69^. Further supporting the association of TLR13/30 and of the huchr Xq paralog equivalent to the MHC (Table 3). The reported TLR4 association to the MHC-paralog 9 equivalent is here confirmed across basal bony fish, with the TLR4 pseudogene of *C. milii* also maps to the huchr 9 equivalent. Finally, TLR27 consistently mapped to the MHC-paralog 1 equivalent in all gnathostome taxa surveyed here. The above results provide evidence that TLR diversification in vertebrates, and in gnathostomes in particular, took place in close physical association to the precursors of key adaptive immune genes (MHC, Igs and TCRs) and of genes involved in the emergence of adaptive immune responses.

### TLR evolution in vertebrates

A closer examination of TLR evolution across major vertebrate lineages, from cyclostomes to mammals (Figure 2), reveals a pattern of both deeply conserved and lineage-specific expansions. The most ancestral TLRs shared across vertebrate groups include members from all vertebrate TLR subfamilies that date back to at least the first round (1R) of whole genome duplication (WGD), i.e., prior to the split between jawless and jawed vertebrates (approximately, 500 million years ago). This set of nine TLR genes (TLR3, 5, 7, 8, 14, 21, 22, 24, 25; Figure 2) likely constitute a core set of innate immune genes crucial for recognizing conserved pathogen-associated molecular patterns. The emergence of jawed vertebrates (gnathostomes) after the second round (2R) of WGD was accompanied by a significant diversification of the TLR repertoire. In particular, chondrichthyans represent the earliest lineage to possess a broader TLR repertoire with 17 TLRs, highlighting a pivotal expansion event in vertebrate immune gene evolution. In addition to conserved and expanded TLRs, certain TLRs appear restricted to specific vertebrate clades. For instance, TLR24 is exclusive to cyclostomes ^70^, while TLR20 and TLR23 are unique to ray-finned fish ^71,72^, and TLR10 and TLR11 are specific to mammals, likely reflecting lineage-specific adaptations to terrestrial environments and distinct pathogen landscapes. TLR6 and TLR12 appear to be restricted to mammals and amphibians, suggesting possible lineage-specific adaptations or gene retention in these groups ^10,73^. Of particular interest is the extensive diversification observed within the TLR11 subfamily, especially in ray-finned fish, suggesting adaptive evolution driven by diverse microbial exposure in aquatic ecosystems. Altogether, this pattern underscores both the evolutionary conservation and innovation of immune recognition mechanisms across vertebrate lineages, with elasmobranchs playing a central role in the early expansion of the vertebrate TLR repertoire.

## Conclusions

This study is the first to investigate the baseline Toll-like receptor (TLR) repertoire in elasmobranchs, examining a wide range of species across most orders. Our findings provide new insights into the TLR repertoire in these ancient vertebrates and the evolutionary history. Elasmobranchs show a complex history of gains and losses of TLR genes, reflecting both conserved and lineage-specific adaptations to environmental and pathogenic pressures. We identified orthologs for several TLRs, including TLR1, TLR2, TLR3, TLR5, TLR5S, TLR7, TLR8, TLR9, TLR13, TLR14/18, TLR21, TLR22, TLR25, TLR27, and TLR29, across 28 species, uncovering diverse patterns of gene duplication and loss within subfamilies. Additionally, a pseudogene version of TLR4 was confirmed in the elephant shark genome. The most notable case of gene duplication occurs in Squaliformes, where duplications of TLR8, TLR9, and TLR21 were observed. Additionally, our analysis indicates that TLR genes in elasmobranchs are predominantly under strong purifying selection, reflecting the necessity of maintaining the structural and functional integrity of these immune receptors. Nevertheless, we also identified positively selected sites across all studied TLRs, suggesting that certain residues are adapting to pathogen-driven pressures. This study underscores the critical role of TLRs in immune function and the evolutionary relevance of elasmobranchs.

## Author contributions

FN, AV, and MJ conceived and designed the study. AMuñoz, FN, and AV performed the bioinformatic analyses. AMatos, JA, and RX analyzed the data and contributed to its interpretation. FN generated all figures and, together with AV, analyzed the data and drafted the manuscript. MJ and YO contributed to data interpretation and critically revised the manuscript. All authors edited the manuscript and approved the final version.

## Data availability

The datasets generated and analysed during the current study are available as supplementary data.

