## Supplementary Figure 1 for "Cartilaginous fish inform the lineage-specific evolution and MHC association of the TLR family"

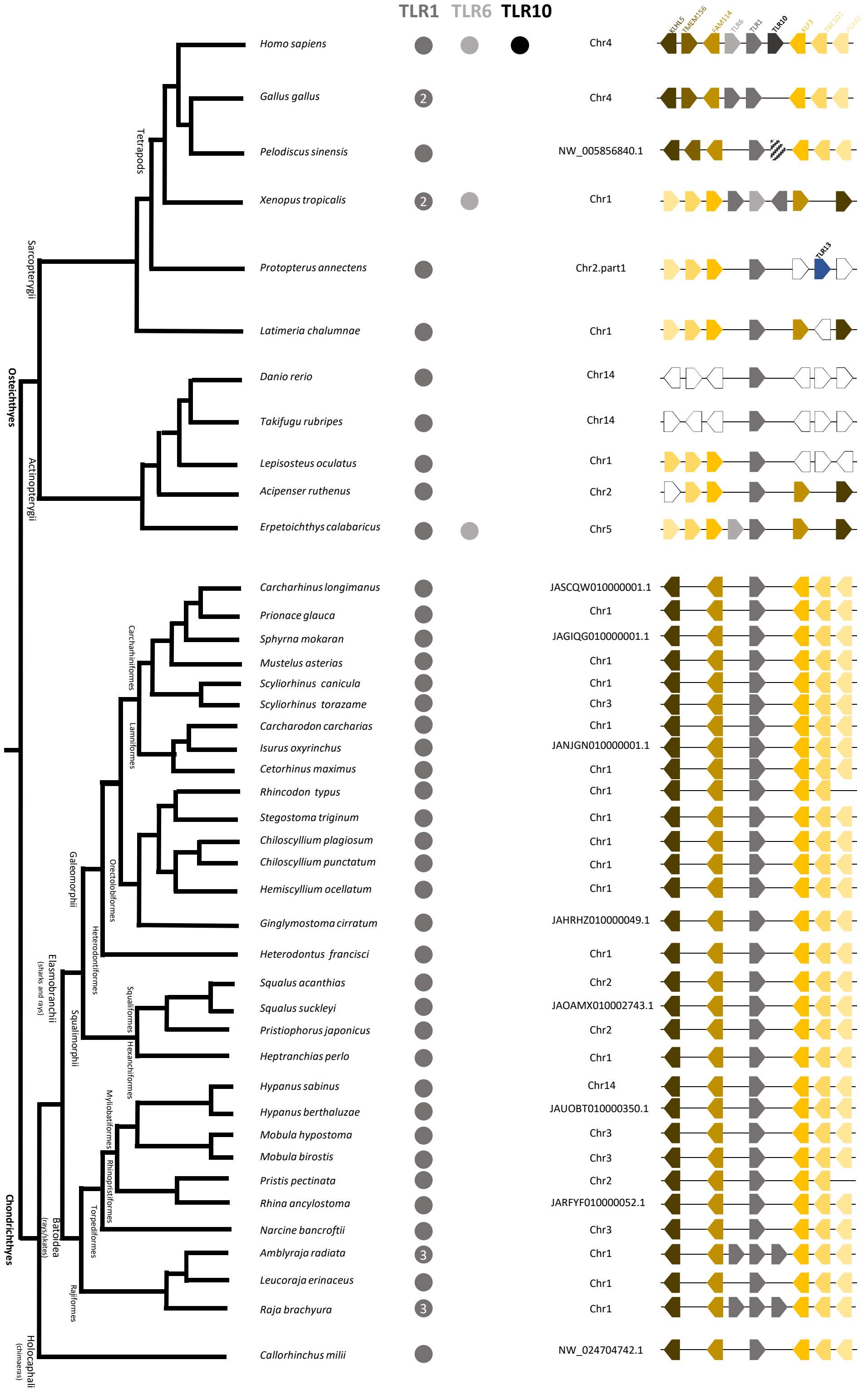
TLR1
TLR6
TLR10
Homo sapiens
Chr4
Gallus gallus
2
Chr4
Pelodiscus sinensis
NW_005856840.1
Xenopus tropicalis
2
Chr1
Protopterus annectens
Chr2.part1
TLR13
Latimeria chalumnae
Chr1
Danio rerio
Chr14
Takifugu rubripes
Chr14
Lepisosteus oculatus
Chr1
Acipenser ruthenus
Chr2
Erpetoichthys calabaricus
Chr5
Carcharhinus longimanus
JASCQW010000001.1
Prionace glauca
Chr1
Sphyrna mokaran
JAGIQG010000001.1
Mustelus asterias
Chr1
Scyliorhinus canicula
Chr1
Scyliorhinus torazame
Chr3
Carcharodon carcharias
Chr1
Isurus oxyrinchus
JANJGN010000001.1
Cetorhinus maximus
Chr1
Rhincodon typus
Chr1
Stegostoma triginum
Chr1
Chiloscyllium plagiosum
Chr1
Chiloscyllium punctatum
Chr1
Hemiscyllium ocellatum
Chr1
Ginglymostoma cirratum
JAHRHZ010000049.1
Heterodontus francisci
Chr1
Squalus acanthias
Chr2
Squalus suckleyi
JAOAMX010002743.1
Pristiophorus japonicus
Chr2
Heptranchias perlo
Chr1
Hypanus sabinus
Chr14
Hypanus berthaluzae
JAUOBT010000350.1
Mobula hypostoma
Chr3
Mobula birostis
Chr3
Pristis pectinata
Chr2
Rhina ancylostoma
JARFYF010000052.1
Narcine bancroftii
Chr3
Amblyraja radiata
3
Chr1
Leucoraja erinaceus
Chr1
Raja brachyura
3
Chr1
Callorhinchus milii
NW_024704742.1
Tetrapods
Sarcopterygii
Osteichthyes
Actinopterygii
Carcharhiniformes
Lamniformes
Galeomorphii
Orectolobiformes
Heterodontiformes
Elasmobranchii
(sharks and rays)
Squaliformes
Squalimorphii
Hexanchiformes
Myliobatiformes
Rhinopristiformes
Torpediformes
Batoidea
(rays/skates)
Rajiformes
Chondrichthyes
Holocephali
(chimaeras)

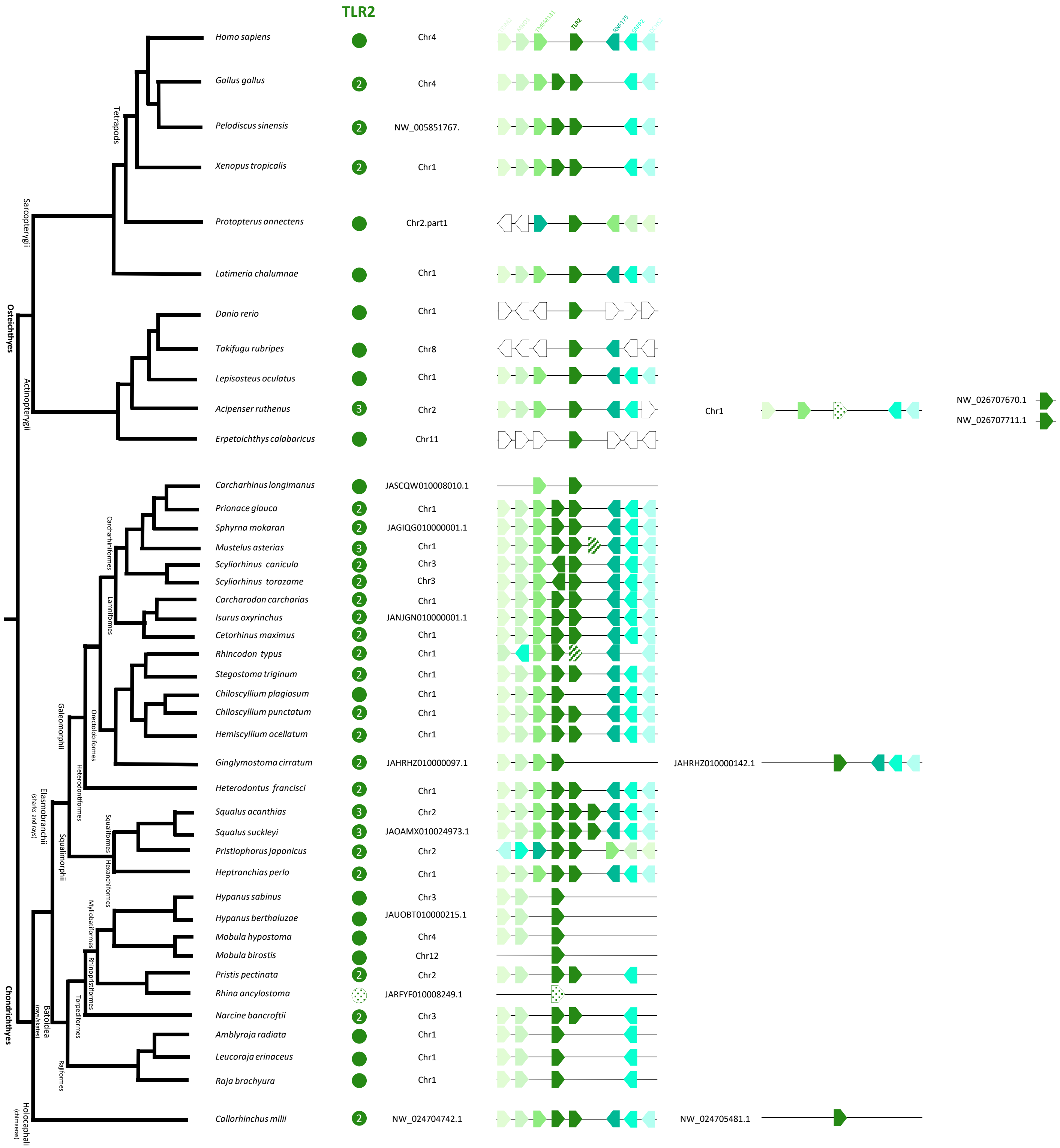
TLR2
Homo sapiens
Chr4
Gallus gallus
Chr4
Pelodiscus sinensis
NW_005851767.
Xenopus tropicalis
Chr1
Protopterus annectens
Chr2.part1
Latimeria chalumnae
Chr1
Danio rerio
Chr1
Takifugu rubripes
Chr8
Lepisosteus oculatus
Chr1
Acipenser ruthenus
Chr2
Chr1
NW_026707670.1
NW_026707711.1
Erpetoichthys calabaricus
Chr11
Carcharhinus longimanus
JASCQW010008010.1
Prionace glauca
Chr1
Sphyrna mokaran
JAGIQG010000001.1
Mustelus asterias
Chr1
Scyliorhinus canicula
Chr3
Scyliorhinus torazame
Chr3
Carcharodon carcharias
Chr1
Isurus oxyrinchus
JANJGN010000001.1
Cetorhinus maximus
Chr1
Rhincodon typus
Chr1
Stegostoma triginum
Chr1
Chiloscyllium plagiosum
Chr1
Chiloscyllium punctatum
Chr1
Hemiscyllium ocellatum
Chr1
Ginglymostoma cirratum
JAHRHZ010000097.1
JAHRHZ010000142.1
Heterodontus francisci
Chr1
Squalus acanthias
Chr2
Squalus suckleyi
JAOAMX010024973.1
Pristiophorus japonicus
Chr2
Heptranchias perlo
Chr1
Hypanus sabinus
Chr3
Hypanus berthaluzae
JAUOBT010000215.1
Mobula hypostoma
Chr4
Mobula birostis
Chr12
Pristis pectinata
Chr2
Rhina ancylostoma
JARFYF010008249.1
Narcine bancroftii
Chr3
Amblyraja radiata
Chr1
Leucoraja erinaceus
Chr1
Raja brachyura
Chr1
Callorhinchus milii
NW_024704742.1
NW_024705481.1
Tetrapods
Sarcopterygii
Osteichthyes
Actinopterygii
Carcharhiniformes
Lamniformes
Orectolobiformes
Galeomorphii
Heterodontiformes
Elasmobranchii
(sharks and rays)
Squalimorphii
Squaliformes
Hexanchiformes
Myliobatiformes
Rhinopristiformes
Torpediformes
Batoidea
(rays/skates)
Rajiformes
Chondrichthyes
Holocephali
(chimaeras)

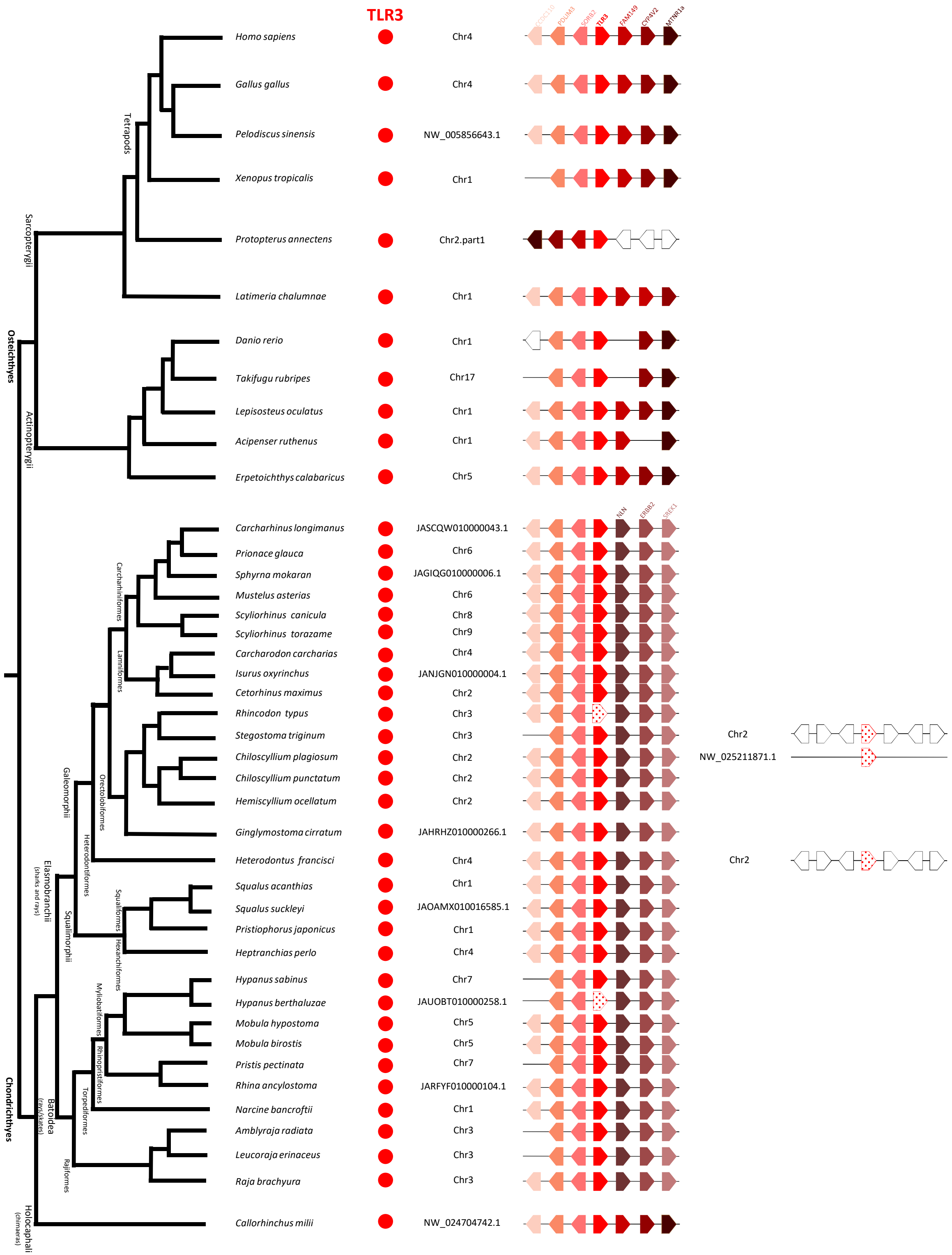
TLR3
CCDC110
PDLIM3
SORB2
TLR3
FAM149
CYP4V2
MTNR1a
NLN
ERBB2
SREK1
Homo sapiens Chr4
Gallus gallus Chr4
Pelodiscus sinensis NW_005856643.1
Xenopus tropicalis Chr1
Protopterus annectens Chr2.part1
Latimeria chalumnae Chr1
Danio rerio Chr1
Takifugu rubripes Chr17
Lepisosteus oculatus Chr1
Acipenser ruthenus Chr1
Erpetoichthys calabaricus Chr5
Carcharhinus longimanus JASCQW010000043.1
Prionace glauca Chr6
Sphyrna mokaran JAGIQG010000006.1
Mustelus asterias Chr6
Scyliorhinus canicula Chr8
Scyliorhinus torazame Chr9
Carcharodon carcharias Chr4
Isurus oxyrinchus JANJGN010000004.1
Cetorhinus maximus Chr2
Rhincodon typus Chr3
Stegostoma triginum Chr3
Chr2
Chiloscyllium plagiosum Chr2
NW_025211871.1
Chiloscyllium punctatum Chr2
Hemiscyllium ocellatum Chr2
Ginglymostoma cirratum JAHRHZ010000266.1
Heterodontus francisci Chr4
Chr2
Squalus acanthias Chr1
Squalus suckleyi JAOAMX010016585.1
Pristiophorus japonicus Chr1
Heptranchias perlo Chr4
Hypanus sabinus Chr7
Hypanus berthaluzae JAUOBT010000258.1
Mobula hypostoma Chr5
Mobula birostis Chr5
Pristis pectinata Chr7
Rhina ancylostoma JARFYF010000104.1
Narcine bancroftii Chr1
Amblyraja radiata Chr3
Leucoraja erinaceus Chr3
Raja brachyura Chr3
Callorhinchus milii NW_024704742.1
Osteichthyes
Sarcopterygii
Tetrapods
Actinopterygii
Chondrichthyes
Elasmobranchii
(sharks and rays)
Galeomorphii
Carcharhiniformes
Lamniformes
Orectolobiformes
Heterodontiformes
Squalomorphii
Squaliformes
Hexanchiformes
Batoidea
(rays/skates)
Myliobatiformes
Rhinopristiformes
Torpediformes
Rajiformes
Holocephali
(chimaeras)

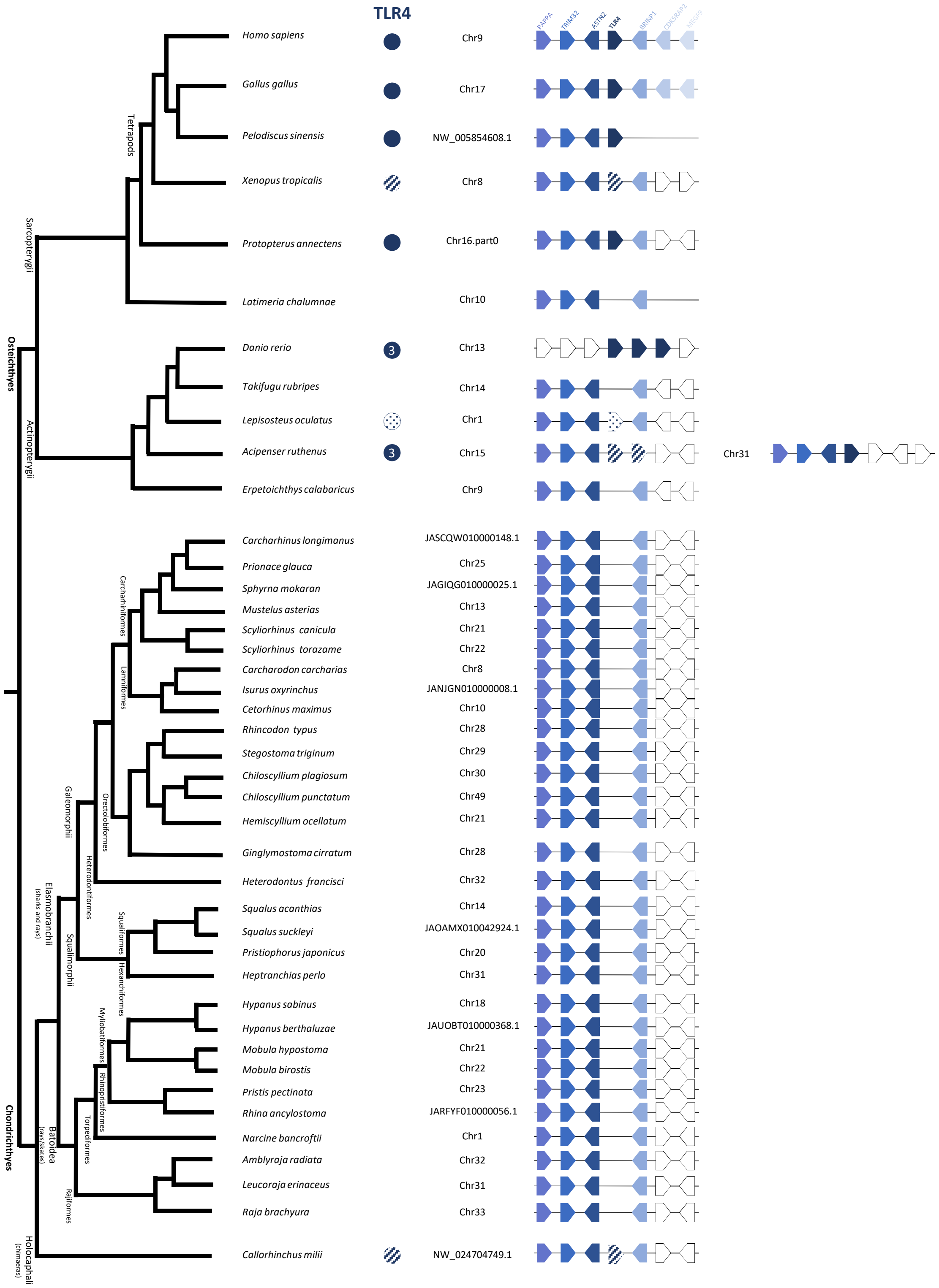

TLR4
PAPPA
TRIM32
ASTN2
TLR4
BRINP1
CDK5RAP2
MEGF9
Homo sapiens
Chr9
Gallus gallus
Chr17
Pelodiscus sinensis
NW_005854608.1
Xenopus tropicalis
Chr8
Protopterus annectens
Chr16.part0
Latimeria chalumnae
Chr10
Danio rerio
3
Chr13
Takifugu rubripes
Chr14
Lepisosteus oculatus
Chr1
Acipenser ruthenus
3
Chr15
Chr31
Erpetoichthys calabaricus
Chr9
Carcharhinus longimanus
JASCQW010000148.1
Prionace glauca
Chr25
Sphyrna mokaran
JAGIQG010000025.1
Mustelus asterias
Chr13
Scyliorhinus canicula
Chr21
Scyliorhinus torazame
Chr22
Carcharodon carcharias
Chr8
Isurus oxyrinchus
JANJGN010000008.1
Cetorhinus maximus
Chr10
Rhincodon typus
Chr28
Stegostoma triginum
Chr29
Chiloscyllium plagiosum
Chr30
Chiloscyllium punctatum
Chr49
Hemiscyllium ocellatum
Chr21
Ginglymostoma cirratum
Chr28
Heterodontus francisci
Chr32
Squalus acanthias
Chr14
Squalus suckleyi
JAOAMX010042924.1
Pristiophorus japonicus
Chr20
Heptranchias perlo
Chr31
Hypanus sabinus
Chr18
Hypanus berthaluzae
JAUOBT010000368.1
Mobula hypostoma
Chr21
Mobula birostis
Chr22
Pristis pectinata
Chr23
Rhina ancylostoma
JARFYF010000056.1
Narcine bancroftii
Chr1
Amblyraja radiata
Chr32
Leucoraja erinaceus
Chr31
Raja brachyura
Chr33
Callorhinchus milii
NW_024704749.1
Osteichthyes
Sarcopterygii
Tetrapods
Actinopterygii
Chondrichthyes
Elasmobranchii
(sharks and rays)
Galeomorphi
Carcharhiniformes
Lamniformes
Orectolobiformes
Heterodontiformes
Squalimorphi
Squaliformes
Hexanchiformes
Batoidea
(rays/skates)
Myliobatiformes
Rhinopristiformes
Torpediformes
Rajiformes
Holocephali
(chimaeras)

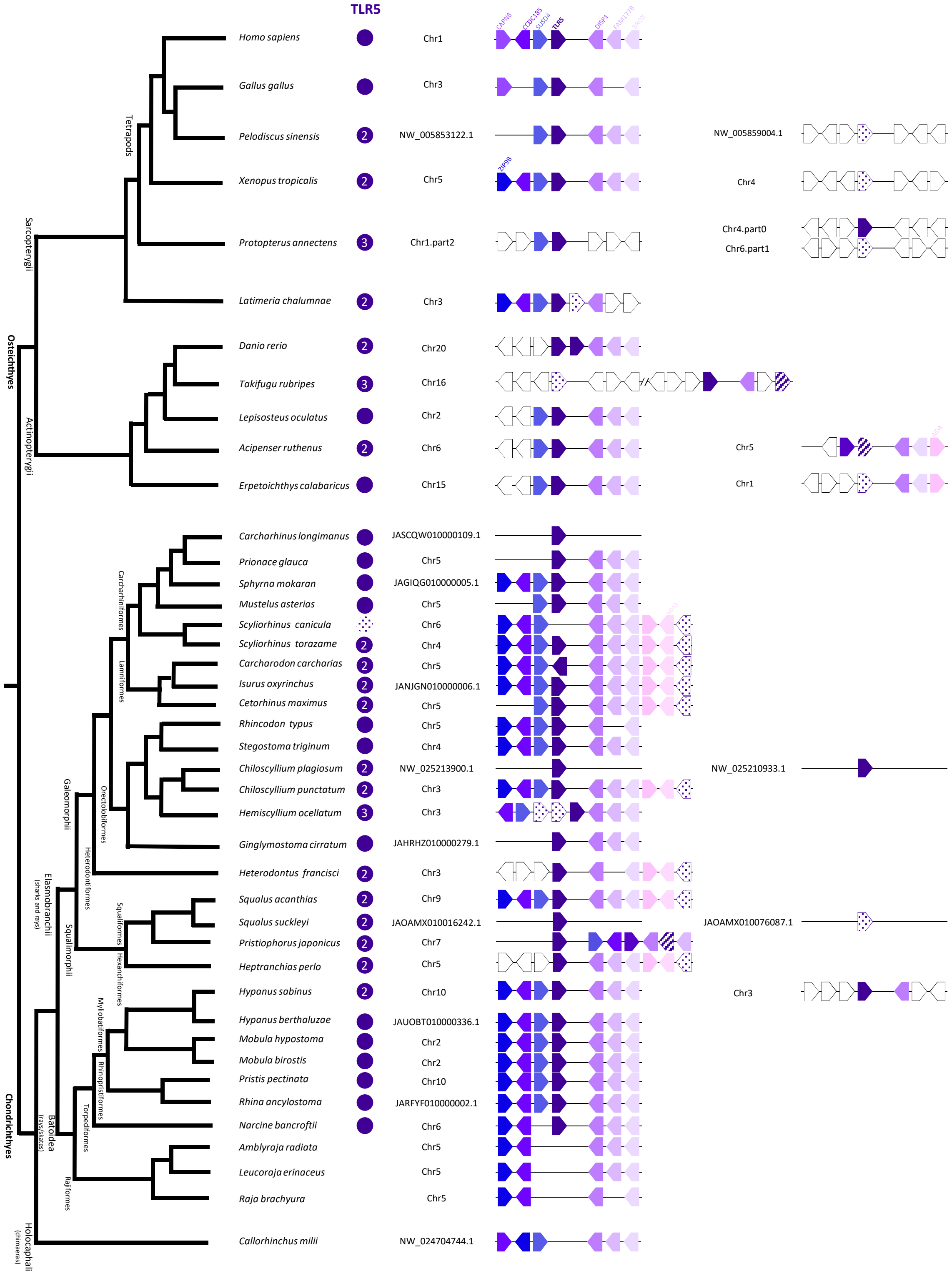

TLR5
Homo sapiens
Chr1
Gallus gallus
Chr3
Pelodiscus sinensis
NW_005853122.1
NW_005859004.1
Xenopus tropicalis
Chr5
Chr4
Protopterus annectens
Chr1.part2
Chr4.part0
Chr6.part1
Latimeria chalumnae
Chr3
Danio rerio
Chr20
Takifugu rubripes
Chr16
Lepisosteus oculatus
Chr2
Acipenser ruthenus
Chr6
Chr5
Erpetoichthys calabaricus
Chr15
Chr1
Carcharhinus longimanus
JASCQW010000109.1
Prionace glauca
Chr5
Sphyrna mokaran
JAGIQG010000005.1
Mustelus asterias
Chr5
Scyliorhinus canicula
Chr6
Scyliorhinus torazame
Chr4
Carcharodon carcharias
Chr5
Isurus oxyrinchus
JANJGN010000006.1
Cetorhinus maximus
Chr5
Rhincodon typus
Chr5
Stegostoma triginum
Chr4
Chiloscyllium plagiosum
NW_025213900.1
NW_025210933.1
Chiloscyllium punctatum
Chr3
Hemiscyllium ocellatum
Chr3
Ginglymostoma cirratum
JAHRHZ010000279.1
Heterodontus francisci
Chr3
Squalus acanthias
Chr9
Squalus suckleyi
JAOAMX010016242.1
JAOAMX010076087.1
Pristiophorus japonicus
Chr7
Heptranchias perlo
Chr5
Hypanus sabinus
Chr10
Chr3
Hypanus berthaluzae
JAUOBT010000336.1
Mobula hypostoma
Chr2
Mobula birostis
Chr2
Pristis pectinata
Chr10
Rhina ancylostoma
JARFYF010000002.1
Narcine bancroftii
Chr6
Amblyraja radiata
Chr5
Leucoraja erinaceus
Chr5
Raja brachyura
Chr5
Callorhinchus milii
NW_024704744.1
Tetrapods
Sarcopterygii
Osteichthyes
Actinopterygii
Carcharhiniformes
Lamniformes
Orectolobiformes
Heterodontiformes
Galeomorphii
Squaliformes
Hexanchiformes
Squalimorphii
Elasmobranchii
(sharks and rays)
Myliobatiformes
Rhinopristiformes
Torpediformes
Rajiformes
Batoidea
(rays/skates)
Chondrichthyes
Holocephali
(chimaeras)

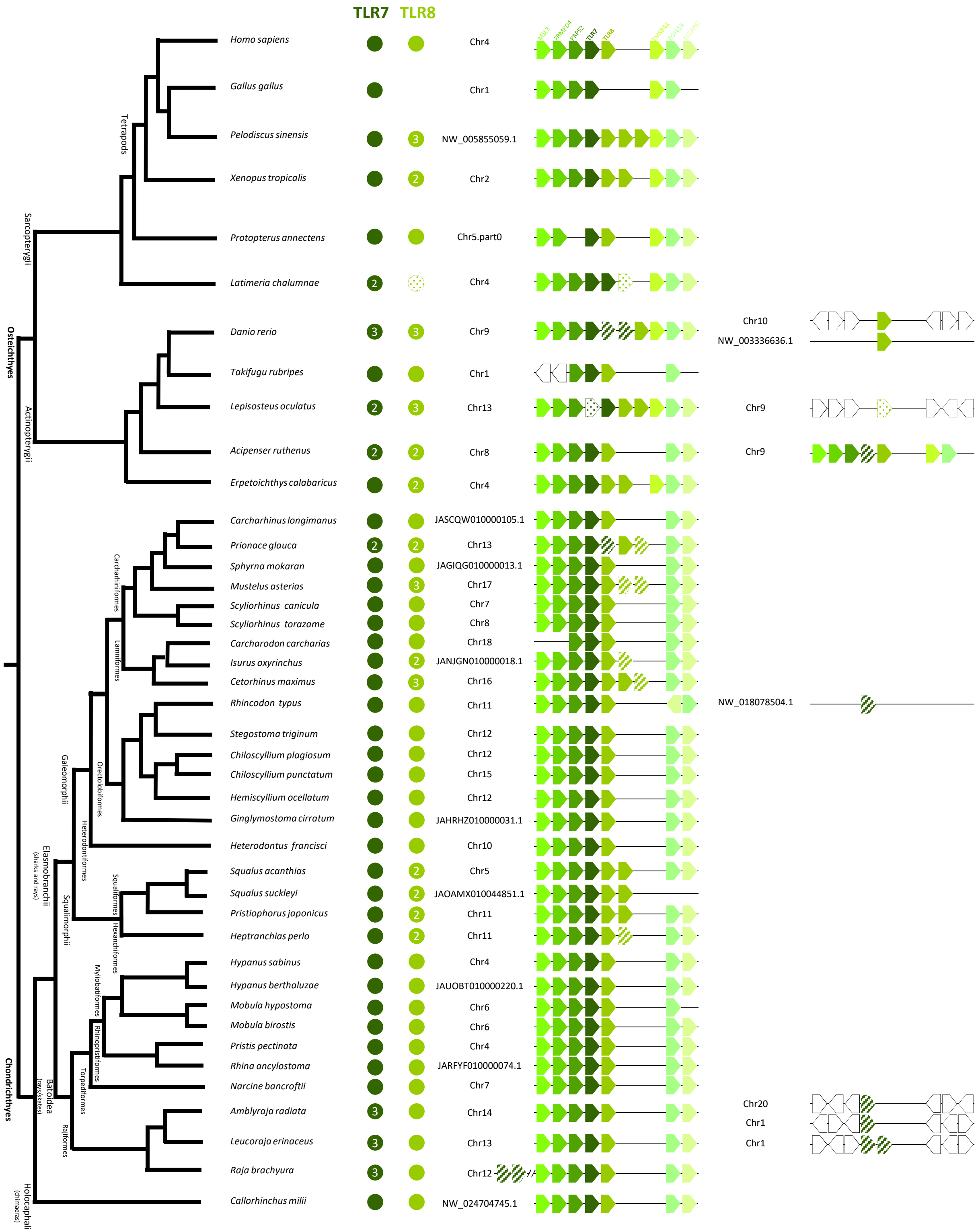
TLR7
TLR8
Osteichthyes
Sarcopterygii
Tetrapods
Actinopterygii
Chondrichthyes
Elasmobranchii
(sharks and rays)
Galeomorphii
Carcharhiniformes
Lamniformes
Orectolobiformes
Heterodontiformes
Squalimorphii
Squaliformes
Hexanchiformes
Batodea
(rays/skates)
Myliobatiformes
Rhinopristiformes
Torpediniformes
Rajiformes
Holocephali
(chimaeras)
Homo sapiens
Chr4
Gallus gallus
Chr1
Pelodiscus sinensis
3
NW_005855059.1
Xenopus tropicalis
2
Chr2
Protopterus annectens
Chr5.part0
Latimeria chalumnae
2
Chr4
Danio rerio
3
3
Chr9
Chr10
NW_003336636.1
Takifugu rubripes
Chr1
Lepisosteus oculatus
2
3
Chr13
Chr9
Acipenser ruthenus
2
2
Chr8
Chr9
Erpetoichthys calabaricus
2
Chr4
Carcharhinus longimanus
JASCQW010000105.1
Prionace glauca
2
2
Chr13
Sphyrna mokarran
JAGIQG010000013.1
Mustelus asterias
3
Chr17
Scyliorhinus canicula
Chr7
Scyliorhinus torazame
Chr8
Carcharodon carcharias
Chr18
Isurus oxyrinchus
2
JANJGN010000018.1
Cetorhinus maximus
3
Chr16
Rhincodon typus
Chr11
NW_018078504.1
Stegostoma triginum
Chr12
Chiloscyllium plagiosum
Chr12
Chiloscyllium punctatum
Chr15
Hemiscyllium ocellatum
Chr12
Ginglymostoma cirratum
JAHRHZ010000031.1
Heterodontus francisci
Chr10
Squalus acanthias
2
Chr5
Squalus suckleyi
2
JAOAMX010044851.1
Pristiophorus japonicus
2
Chr11
Heptranchias perlo
2
Chr11
Hypanus sabinus
Chr4
Hypanus berthaluzae
JAUOBT010000220.1
Mobula hypostoma
Chr6
Mobula birostis
Chr6
Pristis pectinata
Chr4
Rhina ancylostoma
JARFYF010000074.1
Narcine bancroftii
Chr7
Amblyraja radiata
3
Chr14
Chr20
Chr1
Leucoraja erinaceus
3
Chr13
Chr1
Raja brachyura
3
Chr12
Callorhinchus milii
NW_024704745.1

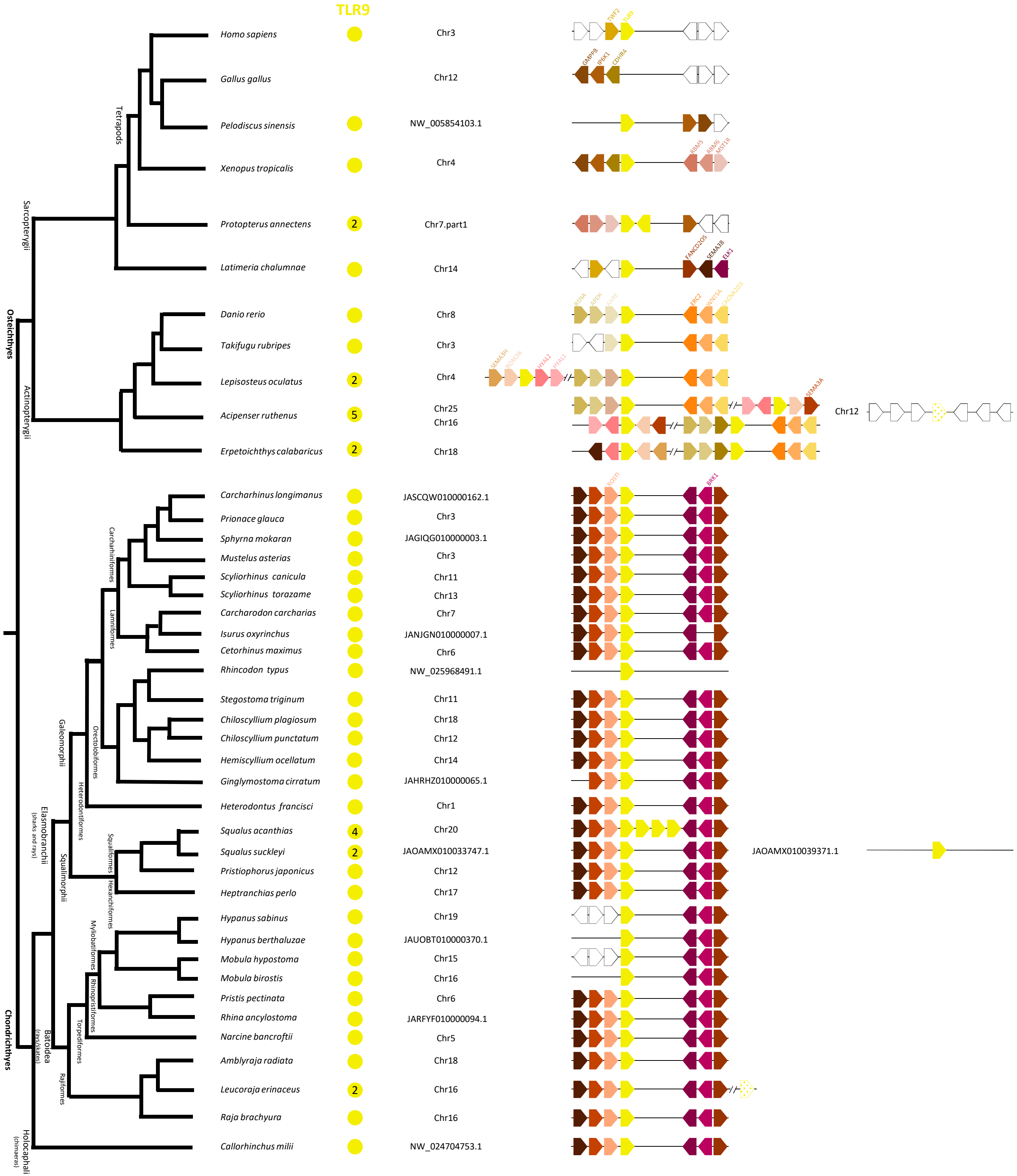
TLR9
Homo sapiens
Chr3
Gallus gallus
Chr12
Pelodiscus sinensis
NW_005854103.1
Xenopus tropicalis
Chr4
Protopterus annectens
2
Chr7.part1
Latimeria chalumnae
Chr14
Danio rerio
Chr8
Takifugu rubripes
Chr3
Lepisosteus oculatus
2
Chr4
Acipenser ruthenus
5
Chr25
Chr16
Chr12
Erpetoichthys calabaricus
2
Chr18
Carcharhinus longimanus
JASCQW010000162.1
Prionace glauca
Chr3
Sphyrna mokaran
JAGIQG010000003.1
Mustelus asterias
Chr3
Scyliorhinus canicula
Chr11
Scyliorhinus torazame
Chr13
Carcharodon carcharias
Chr7
Isurus oxyrinchus
JANJGN010000007.1
Cetorhinus maximus
Chr6
Rhincodon typus
NW_025968491.1
Stegostoma triginum
Chr11
Chiloscyllium plagiosum
Chr18
Chiloscyllium punctatum
Chr12
Hemiscyllium ocellatum
Chr14
Ginglymostoma cirratum
JAHRHZ010000065.1
Heterodontus francisci
Chr1
Squalus acanthias
4
Chr20
Squalus suckleyi
2
JAOAMX010033747.1
JAOAMX010039371.1
Pristiophorus japonicus
Chr12
Heptranchias perlo
Chr17
Hypanus sabinus
Chr19
Hypanus berthaluzae
JAUOBT010000370.1
Mobula hypostoma
Chr15
Mobula birostis
Chr16
Pristis pectinata
Chr6
Rhina ancylostoma
JARFYF010000094.1
Narcine bancroftii
Chr5
Amblyraja radiata
Chr18
Leucoraja erinaceus
2
Chr16
Raja brachyura
Chr16
Callorhinchus milii
NW_024704753.1
Tetrapods
Sarcopterygii
Osteichthyes
Actinopterygii
Carcharhiniformes
Lamniformes
Orectolobiformes
Galeomorphii
Heterodontiformes
Elasmobranchii
(sharks and rays)
Squalimorphii
Squaliformes
Hexanchiformes
Myliobatiformes
Rhinopristiformes
Torpediniformes
Batoidea
(rays/skates)
Rajiformes
Chondrichthyes
Holocephali
(chimaeras)

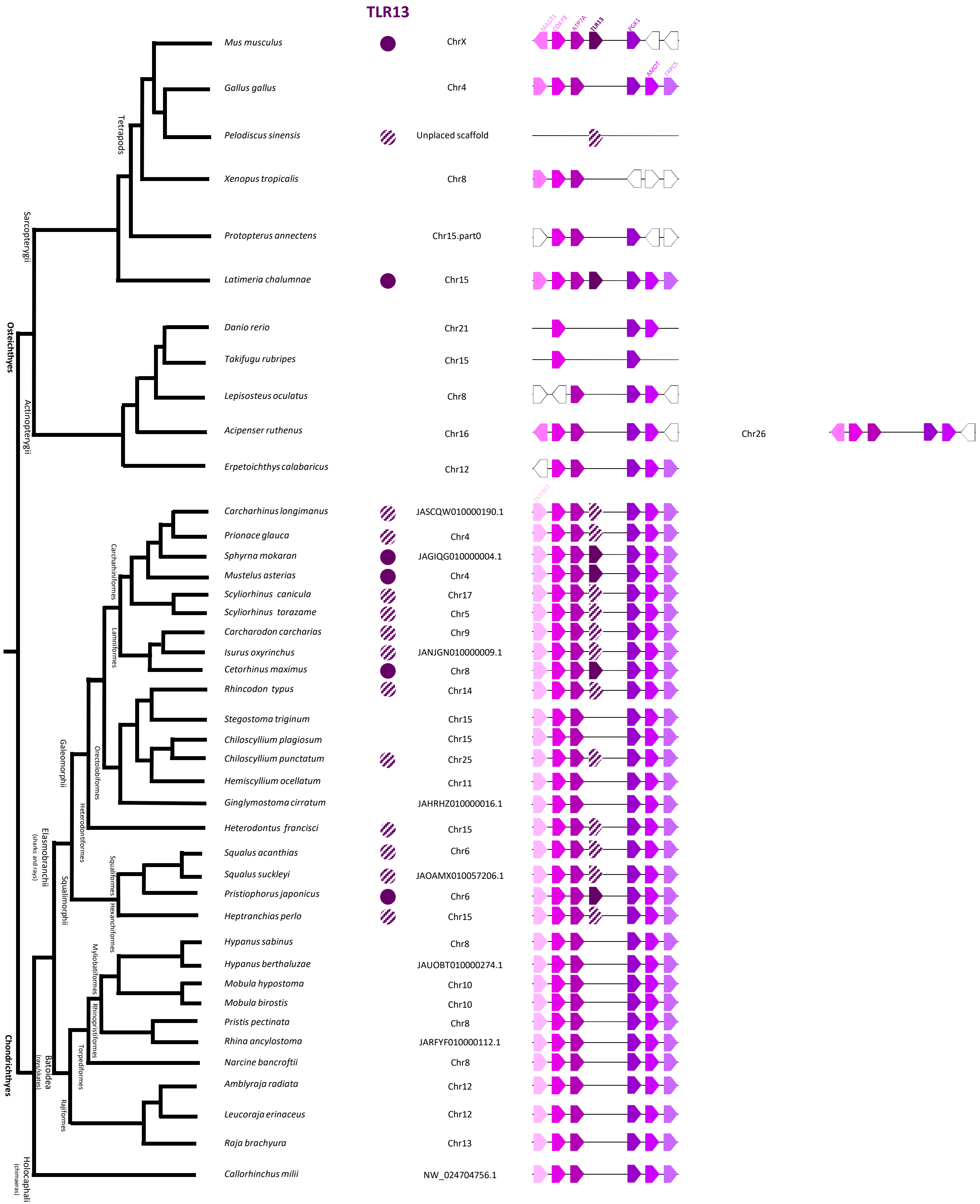
TLR13
MAGT1
COX7B
ATP7A
TLR13
PGK1
AMOT
TRPC5
Mus musculus
ChrX
Gallus gallus
Chr4
Pelodiscus sinensis
Unplaced scaffold
Xenopus tropicalis
Chr8
Protopterus annectens
Chr15.part0
Latimeria chalumnae
Chr15
Danio rerio
Chr21
Takifugu rubripes
Chr15
Lepisosteus oculatus
Chr8
Acipenser ruthenus
Chr16
Chr26
Erpetoichthys calabaricus
Chr12
Carcharhinus longimanus
JASCQW010000190.1
Prionace glauca
Chr4
Sphyrna mokaran
JAGIQG010000004.1
Mustelus asterias
Chr4
Scyliorhinus canicula
Chr17
Scyliorhinus torazame
Chr5
Carcharodon carcharias
Chr9
Isurus oxyrinchus
JANJGN010000009.1
Cetorhinus maximus
Chr8
Rhincodon typus
Chr14
Stegostoma triginum
Chr15
Chiloscyllium plagiosum
Chr15
Chiloscyllium punctatum
Chr25
Hemiscyllium ocellatum
Chr11
Ginglymostoma cirratum
JAHRHZ010000016.1
Heterodontus francisci
Chr15
Squalus acanthias
Chr6
Squalus suckleyi
JAOAMX010057206.1
Pristiophorus japonicus
Chr6
Heptranchias perlo
Chr15
Hypanus sabinus
Chr8
Hypanus berthaluzae
JAUOBT010000274.1
Mobula hypostoma
Chr10
Mobula birostis
Chr10
Pristis pectinata
Chr8
Rhina ancylostoma
JARFYF010000112.1
Narcine bancroftii
Chr8
Amblyraja radiata
Chr12
Leucoraja erinaceus
Chr12
Raja brachyura
Chr13
Callorhinchus milii
NW_024704756.1
Tetrapods
Sarcopterygii
Osteichthyes
Actinopterygii
Carcharhiniformes
Lamniformes
Orectolobiformes
Galeomorphii
Heterodontiformes
Elasmobranchii
(sharks and rays)
Squalimorphii
Squaliformes
Hexanchiformes
Myliobatiformes
Rhinopristiformes
Torpediformes
Batoidea
(rays/skates)
Rajiformes
Chondrichthyes
Holocephali
(chimaeras)

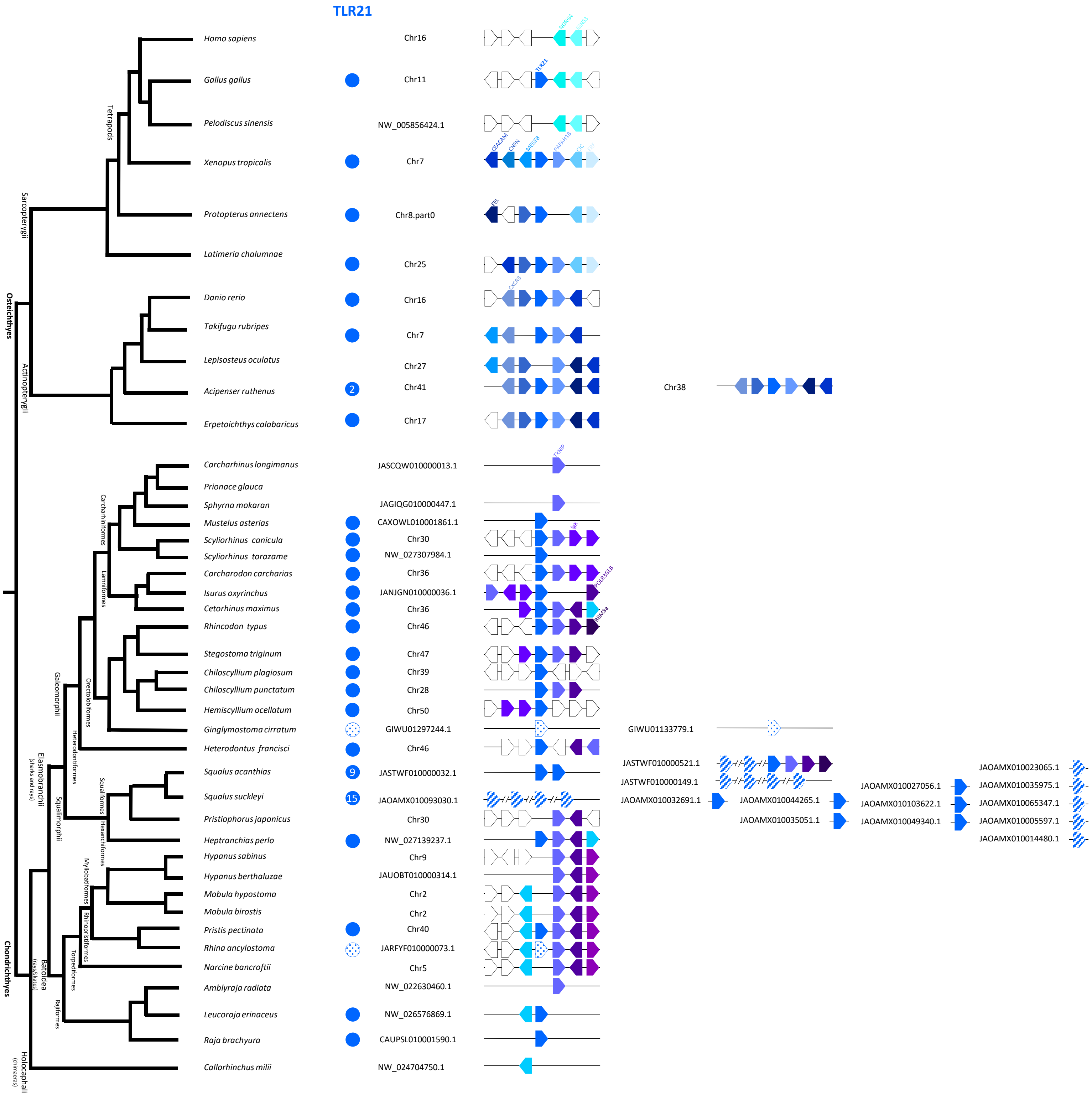

TLR21
Homo sapiens
Chr16
Gallus gallus
Chr11
Pelodiscus sinensis
NW_005856424.1
Xenopus tropicalis
Chr7
Protopterus annectens
Chr8.part0
Latimeria chalumnae
Chr25
Danio rerio
Chr16
Takifugu rubripes
Chr7
Lepisosteus oculatus
Chr27
Acipenser ruthenus
2
Chr41
Chr38
Erpetoichthys calabaricus
Chr17
Carcharhinus longimanus
JASCQW010000013.1
Prionace glauca
Sphyrna mokaran
JAGIQG010000447.1
Mustelus asterias
CAXOWL010001861.1
Scyliorhinus canicula
Chr30
Scyliorhinus torazame
NW_027307984.1
Carcharodon carcharias
Chr36
Isurus oxyrinchus
JANJGN010000036.1
Cetorhinus maximus
Chr36
Rhincodon typus
Chr46
Stegostoma triginum
Chr47
Chiloscyllium plagiosum
Chr39
Chiloscyllium punctatum
Chr28
Hemiscyllium ocellatum
Chr50
Ginglymostoma cirratum
GIWU01297244.1
GIWU01133779.1
Heterodontus francisci
Chr46
Squalus acanthias
9
JASTWF010000032.1
JASTWF010000521.1
JASTWF010000149.1
Squalus suckleyi
15
JAOAMX010093030.1
JAOAMX010032691.1
JAOAMX010044265.1
JAOAMX010035051.1
JAOAMX010027056.1
JAOAMX010103622.1
JAOAMX010049340.1
JAOAMX010023065.1
JAOAMX010035975.1
JAOAMX010065347.1
JAOAMX010005597.1
JAOAMX010014480.1
Pristiophorus japonicus
Chr30
Heptranchias perlo
NW_027139237.1
Hypanus sabinus
Chr9
Hypanus berthaluzae
JAUOBT010000314.1
Mobula hypostoma
Chr2
Mobula birostis
Chr2
Pristis pectinata
Chr40
Rhina ancylostoma
JARFYF010000073.1
Narcine bancroftii
Chr5
Amblyraja radiata
NW_022630460.1
Leucoraja erinaceus
NW_026576869.1
Raja brachyura
CAUPSL010001590.1
Callorhinchus milii
NW_024704750.1
Tetrapods
Sarcopterygii
Osteichthyes
Actinopterygii
Carcharhiniformes
Lamniformes
Orectolobiformes
Galeomorphii
Heterodontiformes
Elasmobranchii
(sharks and rays)
Squalomorphii
Squaliformes
Hexanchiformes
Myliobatiformes
Rhinopristiformes
Torpediniformes
Batoidea
(rays/skates)
Rajiformes
Chondrichthyes
Holocephali
(chimaeras)

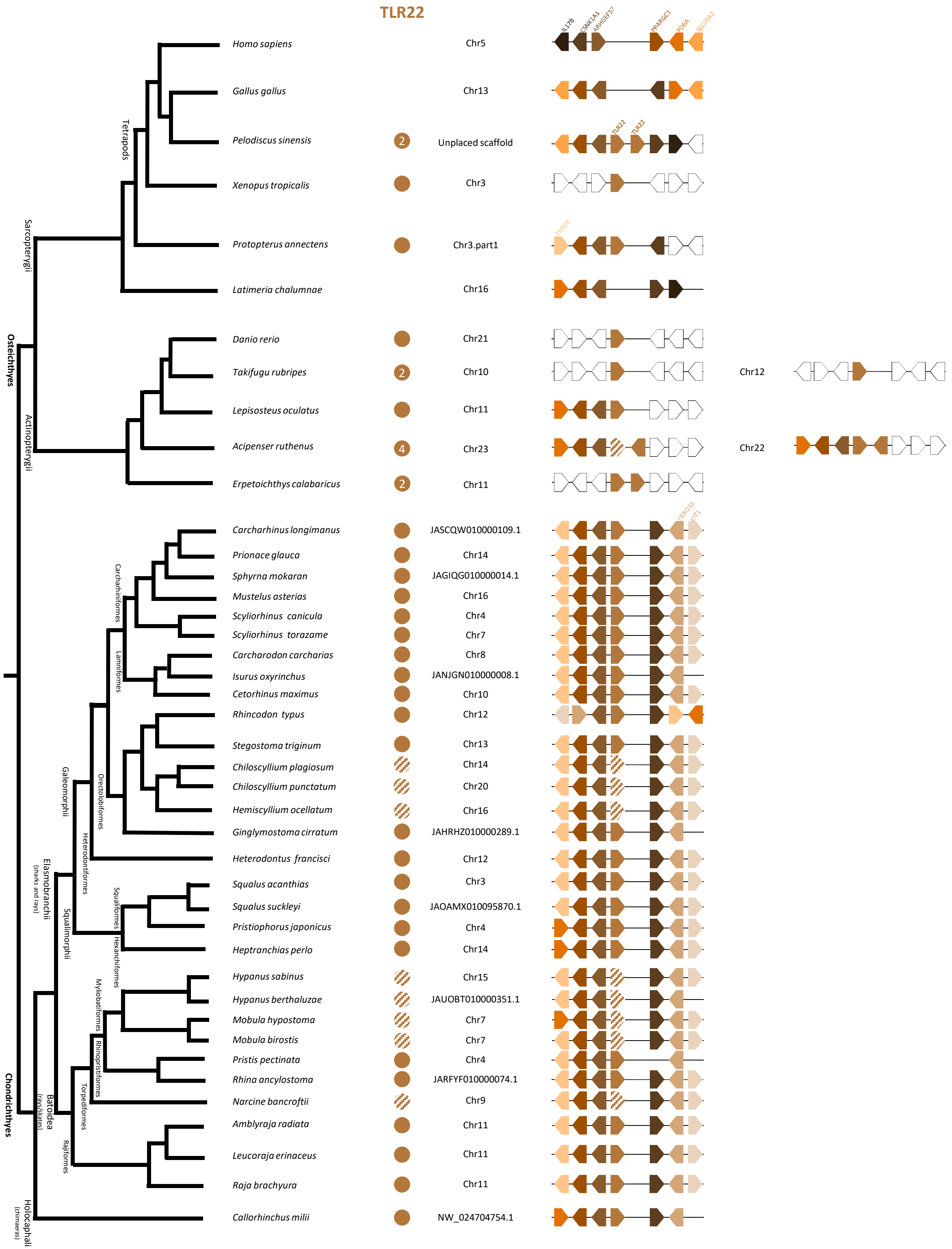

TLR22
Homo sapiens
Chr5
Gallus gallus
Chr13
Pelodiscus sinensis
2
Unplaced scaffold
Xenopus tropicalis
Chr3
Protopterus annectens
Chr3.part1
Latimeria chalumnae
Chr16
Danio rerio
Chr21
Takifugu rubripes
2
Chr10
Chr12
Lepisosteus oculatus
Chr11
Acipenser ruthenus
4
Chr23
Chr22
Erpetoichthys calabaricus
2
Chr11
Carcharhinus longimanus
JASCQW010000109.1
Prionace glauca
Chr14
Sphyrna mokaran
JAGIQG010000014.1
Mustelus asterias
Chr16
Scyliorhinus canicula
Chr4
Scyliorhinus torazame
Chr7
Carcharodon carcharias
Chr8
Isurus oxyrinchus
JANJGN010000008.1
Cetorhinus maximus
Chr10
Rhincodon typus
Chr12
Stegostoma triginum
Chr13
Chiloscyllium plagiosum
Chr14
Chiloscyllium punctatum
Chr20
Hemiscyllium ocellatum
Chr16
Ginglymostoma cirratum
JAHRHZ010000289.1
Heterodontus francisci
Chr12
Squalus acanthias
Chr3
Squalus suckleyi
JAOAMX010095870.1
Pristiophorus japonicus
Chr4
Heptranchias perlo
Chr14
Hypanus sabinus
Chr15
Hypanus berthaluzae
JAUOBT010000351.1
Mobula hypostoma
Chr7
Mobula birostis
Chr7
Pristis pectinata
Chr4
Rhina ancylostoma
JARFYF010000074.1
Narcine bancroftii
Chr9
Amblyraja radiata
Chr11
Leucoraja erinaceus
Chr11
Raja brachyura
Chr11
Callorhinchus milii
NW_024704754.1
Tetrapods
Sarcopterygii
Osteichthyes
Actinopterygii
Carcharhiniformes
Lamniformes
Orectolobiformes
Heterodontiformes
Galeomorphii
Squalomorphii
Squaliformes
Hexanchiformes
Elasmobranchii
(sharks and rays)
Myliobatiformes
Rhinopristiformes
Torpediniformes
Rajiformes
Batoidea
(rays/skates)
Chondrichthyes
Holocephali
(chimeras)

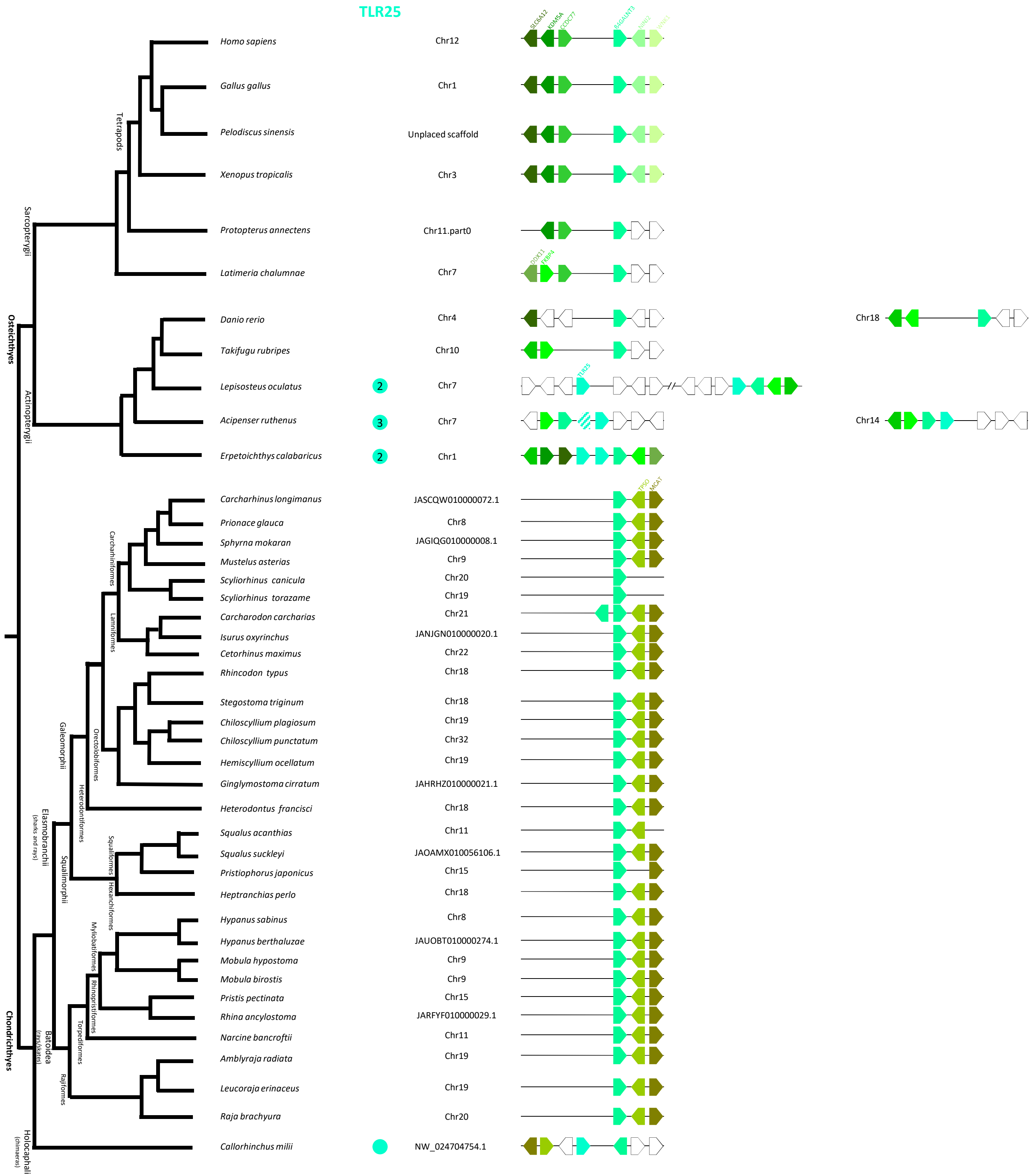

TLR25
Homo sapiens
Chr12
SLC6A12
KDM5A
CCDC77
B4GALNT3
NINJ2
WNK1
Gallus gallus
Chr1
Pelodiscus sinensis
Unplaced scaffold
Xenopus tropicalis
Chr3
Protopterus annectens
Chr11.part0
Latimeria chalumnae
Chr7
DDX11
FKBP4
Danio rerio
Chr4
Chr18
Takifugu rubripes
Chr10
Lepisosteus oculatus
2
Chr7
TLR25
Acipenser ruthenus
3
Chr7
Chr14
Erpetoichthys calabaricus
2
Chr1
Carcharhinus longimanus
JASCQW010000072.1
TPSO
MCAT
Prionace glauca
Chr8
Sphyrna mokaran
JAGIQG010000008.1
Mustelus asterias
Chr9
Scyliorhinus canicula
Chr20
Scyliorhinus torazame
Chr19
Carcharodon carcharias
Chr21
Isurus oxyrinchus
JANJGN010000020.1
Cetorhinus maximus
Chr22
Rhincodon typus
Chr18
Stegostoma triginum
Chr18
Chiloscyllium plagiosum
Chr19
Chiloscyllium punctatum
Chr32
Hemiscyllium ocellatum
Chr19
Ginglymostoma cirratum
JAHRHZ010000021.1
Heterodontus francisci
Chr18
Squalus acanthias
Chr11
Squalus suckleyi
JAOAMX010056106.1
Pristiophorus japonicus
Chr15
Heptranchias perlo
Chr18
Hypanus sabinus
Chr8
Hypanus berthaluzae
JAUOBT010000274.1
Mobula hypostoma
Chr9
Mobula birostis
Chr9
Pristis pectinata
Chr15
Rhina ancylostoma
JARFYF010000029.1
Narcine bancroftii
Chr11
Amblyraja radiata
Chr19
Leucoraja erinaceus
Chr19
Raja brachyura
Chr20
Callorhinchus milii
NW_024704754.1
Tetrapods
Sarcopterygii
Osteichthyes
Actinopterygii
Carcharhiniformes
Lamniformes
Orectolobiformes
Galeomorphii
Heterodontiformes
Elasmobranchii
(sharks and rays)
Squalimorphii
Squaliformes
Hexanchiformes
Myliobatiformes
Rhinopristiformes
Torpediformes
Batoidea
(rays/skates)
Rajiformes
Chondrichthyes
Holocephali
(chimaeras)

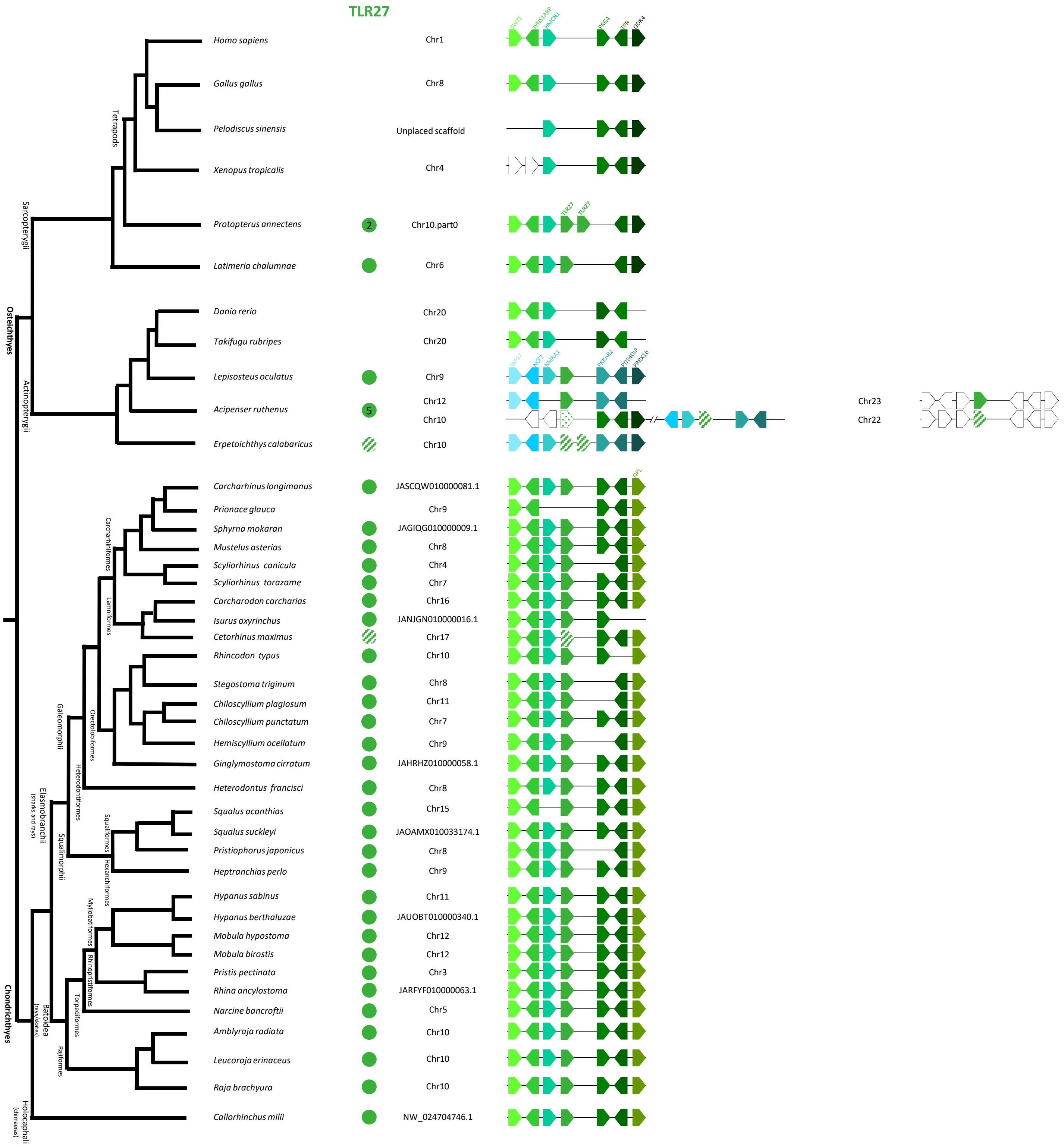
TLR27
Homo sapiens
Chr1
Gallus gallus
Chr8
Pelodiscus sinensis
Unplaced scaffold
Xenopus tropicalis
Chr4
Protopterus annectens
2
Chr10.part0
Latimeria chalumnae
Chr6
Danio rerio
Chr20
Takifugu rubripes
Chr20
Lepisosteus oculatus
Chr9
Acipenser ruthenus
5
Chr12
Chr10
Chr23
Chr22
Erpetoichthys calabaricus
Chr10
Carcharhinus longimanus
JASCQW010000081.1
Prionace glauca
Chr9
Sphyrna mokaran
JAGIQG010000009.1
Mustelus asterias
Chr8
Scyliorhinus canicula
Chr4
Scyliorhinus torazame
Chr7
Carcharodon carcharias
Chr16
Isurus oxyrinchus
JANJGN010000016.1
Cetorhinus maximus
Chr17
Rhincodon typus
Chr10
Stegostoma triginum
Chr8
Chiloscyllium plagiosum
Chr11
Chiloscyllium punctatum
Chr7
Hemiscyllium ocellatum
Chr9
Ginglymostoma cirratum
JAHRHZ010000058.1
Heterodontus francisci
Chr8
Squalus acanthias
Chr15
Squalus suckleyi
JAOAMX010033174.1
Pristiophorus japonicus
Chr8
Heptranchias perlo
Chr9
Hypanus sabinus
Chr11
Hypanus berthaluzae
JAUOBT010000340.1
Mobula hypostoma
Chr12
Mobula birostis
Chr12
Pristis pectinata
Chr3
Rhina ancylostoma
JARFYF010000063.1
Narcine bancroftii
Chr5
Amblyraja radiata
Chr10
Leucoraja erinaceus
Chr10
Raja brachyura
Chr10
Callorhinchus milii
NW_024704746.1
Tetrapods
Sarcopterygii
Osteichthyes
Actinopterygii
Carcharhiniformes
Lamniformes
Orectolobiformes
Galeomorphii
Heterodontiformes
Elasmobranchii
(sharks and rays)
Squalimorphii
Squaliformes
Hexanchiformes
Myliobatiformes
Rhinopristiformes
Torpediformes
Rajiformes
Batoidea
(rays/skates)
Chondrichthyes
Holocephali
(chimaeras)

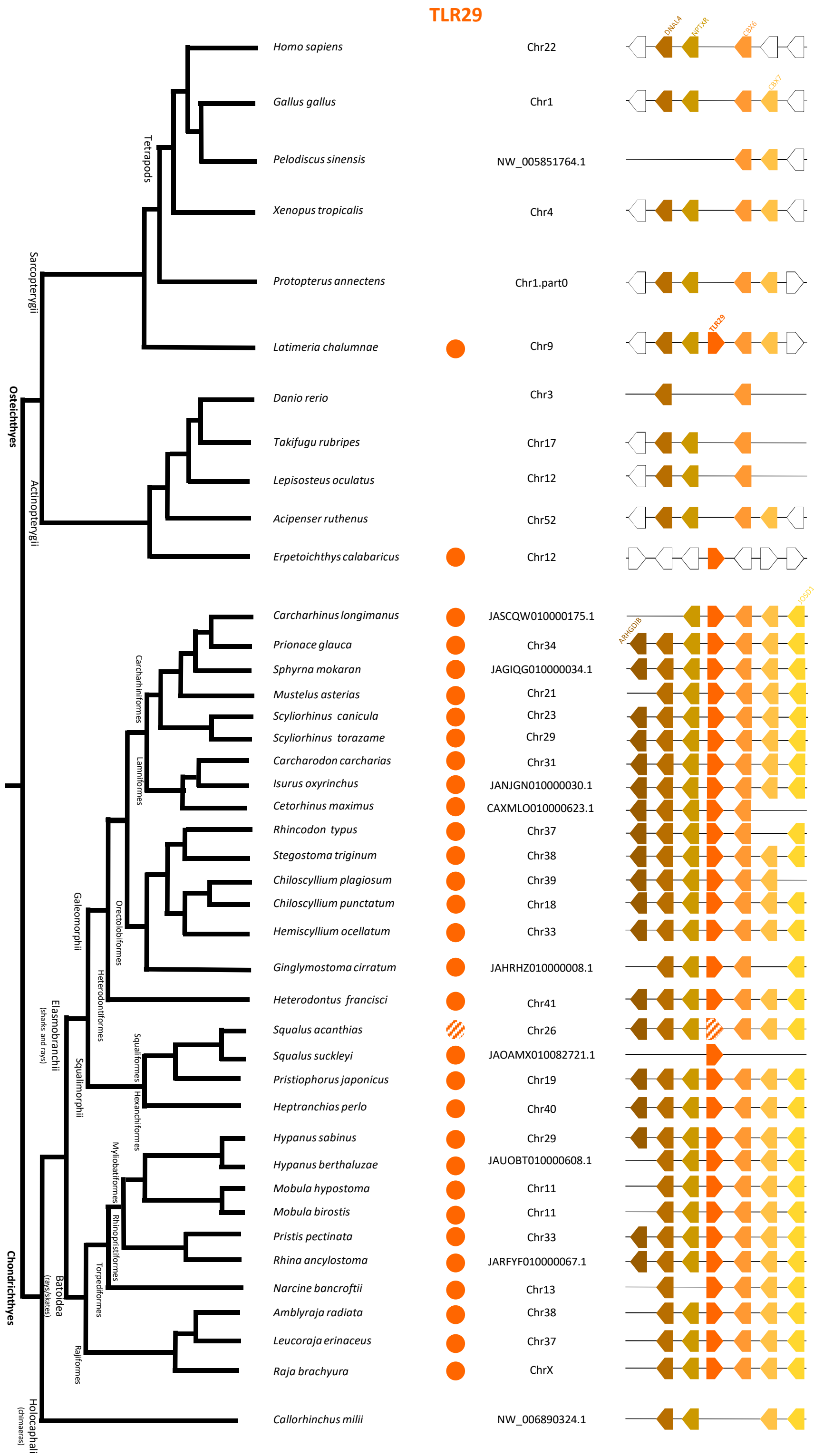
TLR29
DNAL4
NPTXR
CBX6
CBX7
TLR29
ARHGDIB
JOSD1
Homo sapiens
Chr22
Gallus gallus
Chr1
Pelodiscus sinensis
NW_005851764.1
Xenopus tropicalis
Chr4
Protopterus annectens
Chr1.part0
Latimeria chalumnae
Chr9
Danio rerio
Chr3
Takifugu rubripes
Chr17
Lepisosteus oculatus
Chr12
Acipenser ruthenus
Chr52
Erpetoichthys calabaricus
Chr12
Carcharhinus longimanus
JASCQW010000175.1
Prionace glauca
Chr34
Sphyrna mokaran
JAGIQG010000034.1
Mustelus asterias
Chr21
Scyliorhinus canicula
Chr23
Scyliorhinus torazame
Chr29
Carcharodon carcharias
Chr31
Isurus oxyrinchus
JANJGN010000030.1
Cetorhinus maximus
CAXMLO010000623.1
Rhincodon typus
Chr37
Stegostoma triginum
Chr38
Chiloscyllium plagiosum
Chr39
Chiloscyllium punctatum
Chr18
Hemiscyllium ocellatum
Chr33
Ginglymostoma cirratum
JAHRHZ010000008.1
Heterodontus francisci
Chr41
Squalus acanthias
Chr26
Squalus suckleyi
JAOAMX010082721.1
Pristiophorus japonicus
Chr19
Heptranchias perlo
Chr40
Hypanus sabinus
Chr29
Hypanus berthaluzae
JAUOBT010000608.1
Mobula hypostoma
Chr11
Mobula birostis
Chr11
Pristis pectinata
Chr33
Rhina ancylostoma
JARFYF010000067.1
Narcine bancroftii
Chr13
Amblyraja radiata
Chr38
Leucoraja erinaceus
Chr37
Raja brachyura
ChrX
Callorhinchus milii
NW_006890324.1
Osteichthyes
Sarcopterygii
Tetrapods
Actinopterygii
Chondrichthyes
Elasmobranchii
(sharks and rays)
Galeomorphii
Carcharhiniformes
Lamniformes
Orectolobiformes
Heterodontiformes
Squalomorphii
Squaliformes
Hexanchiformes
Batoidea
(rays/skates)
Myliobatiformes
Rhinopristiformes
Torpediniformes
Rajiformes
Holocephali
(chimaeras)

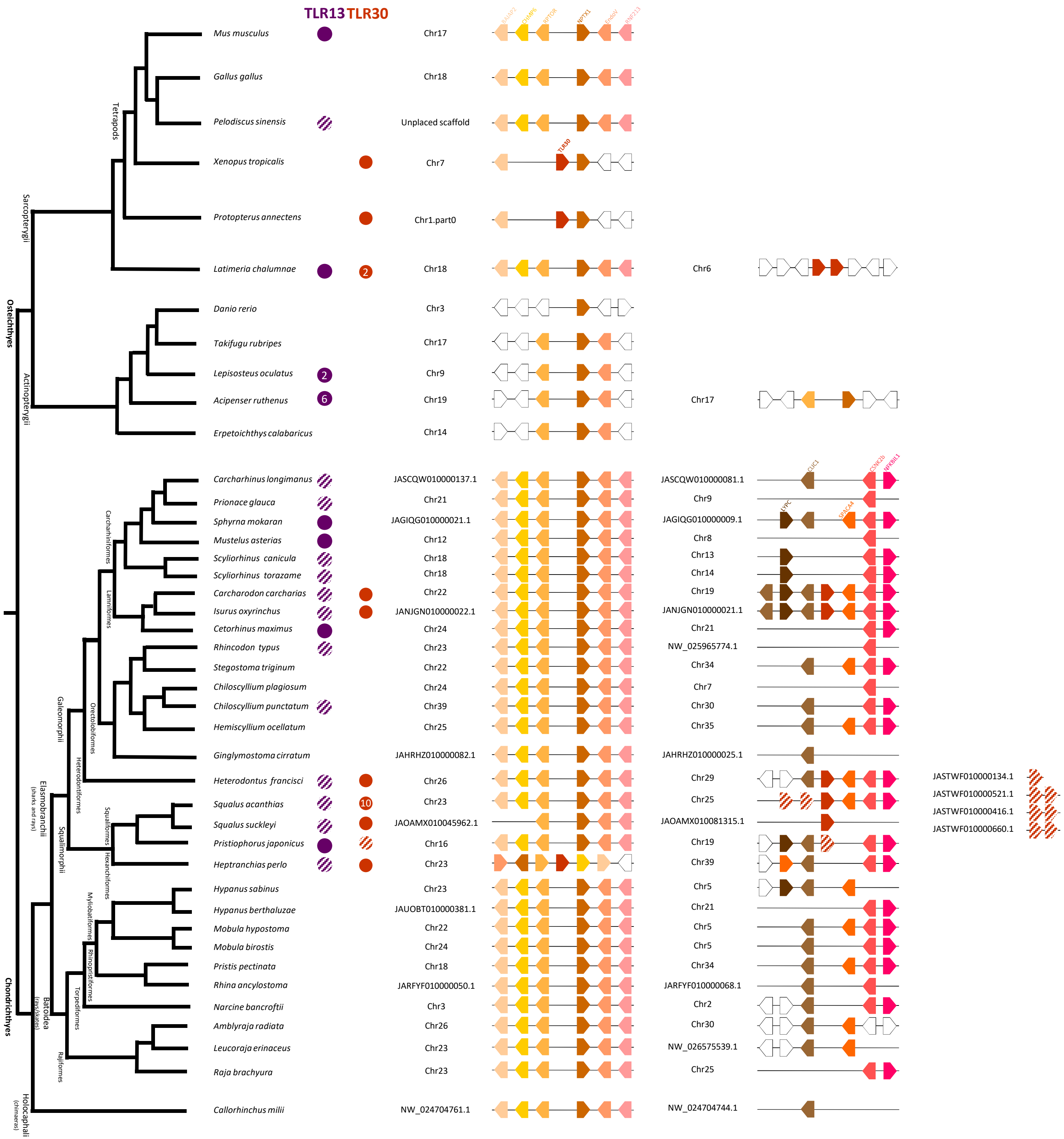
TLR13 TLR30
Mus musculus
Gallus gallus
Pelodiscus sinensis
Xenopus tropicalis
Protopterus annectens
Latimeria chalumnae
Danio rerio
Takifugu rubripes
Lepisosteus oculatus
Acipenser ruthenus
Erpetoichthys calabaricus
Carcharhinus longimanus
Prionace glauca
Sphyrna mokaran
Mustelus asterias
Scyliorhinus canicula
Scyliorhinus torazame
Carcharodon carcharias
Isurus oxyrinchus
Cetorhinus maximus
Rhincodon typus
Stegostoma triginum
Chiloscyllium plagiosum
Chiloscyllium punctatum
Hemiscyllium ocellatum
Ginglymostoma cirratum
Heterodontus francisci
Squalus acanthias
Squalus suckleyi
Pristiophorus japonicus
Heptranchias perlo
Hypanus sabinus
Hypanus berthaluzae
Mobula hypostoma
Mobula birostis
Pristis pectinata
Rhina ancylostoma
Narcine bancroftii
Amblyraja radiata
Leucoraja erinaceus
Raja brachyura
Callorhinchus milii
Tetrapods
Sarcopterygii
Osteichthyes
Actinopterygii
Carcharhiniformes
Lamniformes
Orectolobiformes
Galeomorphii
Heterodontiformes
Squaliformes
Hexanchiformes
Squalomorphii
Elasmobranchii (sharks and rays)
Myliobatiformes
Rhinopristiformes
Torpediniformes
Rajiformes
Batoidea
Chondrichthyes
Holocephali (chimaeras)
2
2
6
10
Chr17
Chr18
Unplaced scaffold
Chr7
Chr1.part0
Chr18
Chr3
Chr17
Chr9
Chr19
Chr14
JASCQW010000137.1
Chr21
JAGIQG010000021.1
Chr12
Chr18
Chr18
Chr22
JANJGN010000022.1
Chr24
Chr23
Chr22
Chr24
Chr39
Chr25
JAHRHZ010000082.1
Chr26
Chr23
JAOAMX010045962.1
Chr16
Chr23
Chr23
JAUOBT010000381.1
Chr22
Chr24
Chr18
JARFYF010000050.1
Chr3
Chr26
Chr23
Chr23
NW_024704761.1
TLR30
Chr6
Chr17
CLIC1
CSNK2b
NFKBIL1
LY6G
SLC44A4
JASCQW010000081.1
Chr9
JAGIQG010000009.1
Chr8
Chr13
Chr14
Chr19
JANJGN010000021.1
Chr21
NW_025965774.1
Chr34
Chr7
Chr30
Chr35
JAHRHZ010000025.1
Chr29
Chr25
JAOAMX010081315.1
Chr19
Chr39
Chr5
Chr21
Chr5
Chr5
Chr34
JARFYF010000068.1
Chr2
Chr30
NW_026575539.1
Chr25
NW_024704744.1
JASTWF010000134.1
JASTWF010000521.1
JASTWF010000416.1
JASTWF010000660.1

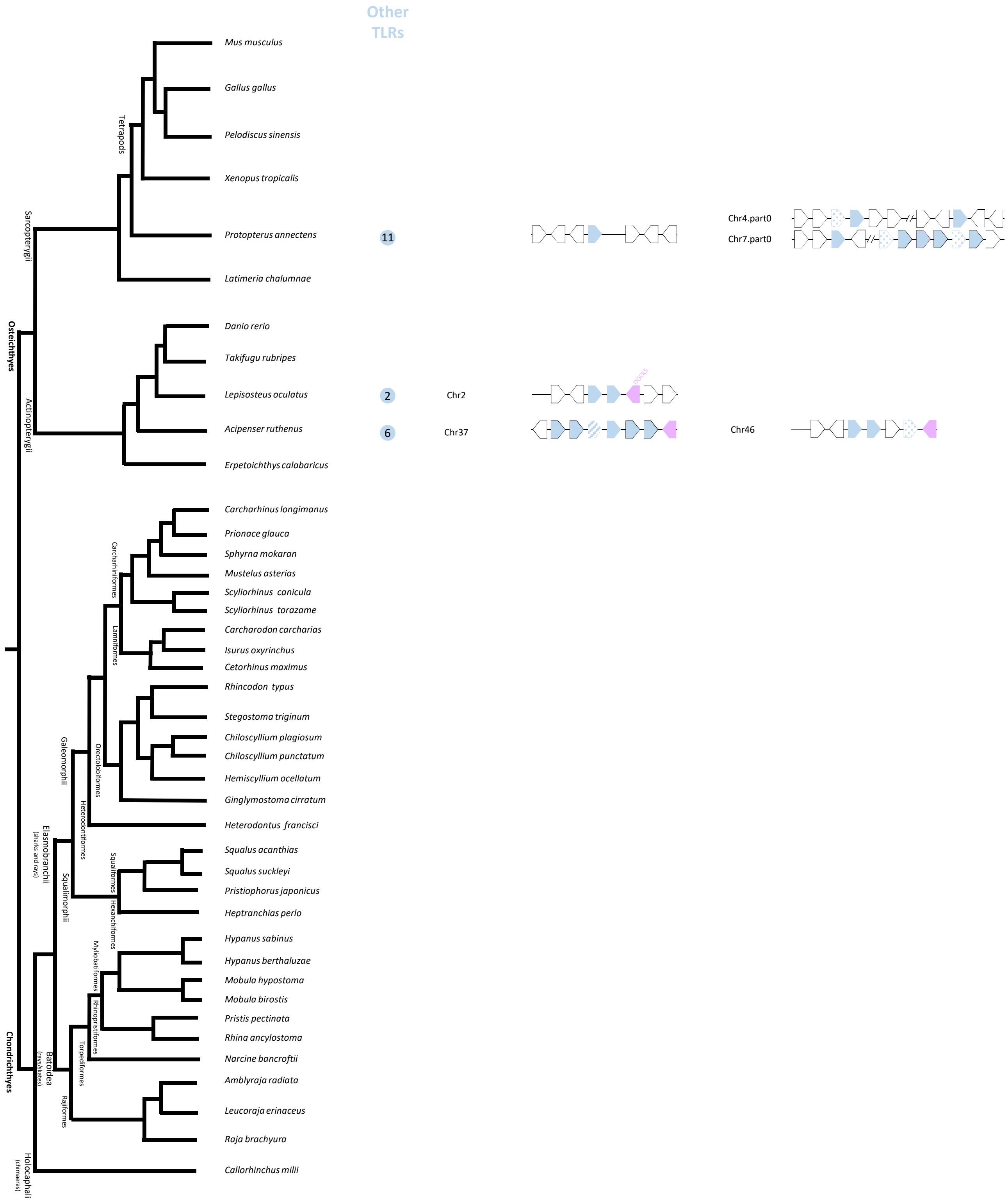

Other TLRs
Mus musculus
Gallus gallus
Pelodiscus sinensis
Xenopus tropicalis
Protopterus annectens
11
Chr4.part0
Chr7.part0
Latimeria chalumnae
Danio rerio
Takifugu rubripes
Lepisosteus oculatus
2
Chr2
Acipenser ruthenus
6
Chr37
Chr46
Erpetoichthys calabaricus
Carcharhinus longimanus
Prionace glauca
Sphyrna mokaran
Mustelus asterias
Scyliorhinus canicula
Scyliorhinus torazame
Carcharodon carcharias
Isurus oxyrinchus
Cetorhinus maximus
Rhincodon typus
Stegostoma triginum
Chiloscyllium plagiosum
Chiloscyllium punctatum
Hemiscyllium ocellatum
Ginglymostoma cirratum
Heterodontus francisci
Squalus acanthias
Squalus suckleyi
Pristiophorus japonicus
Heptranchias perlo
Hypanus sabinus
Hypanus berthaluzae
Mobula hypostoma
Mobula birostis
Pristis pectinata
Rhina ancylostoma
Narcine bancroftii
Amblyraja radiata
Leucoraja erinaceus
Raja brachyura
Callorhinchus milii
Tetrapods
Sarcopterygii
Osteichthyes
Actinopterygii
Carcharhiniformes
Lamniformes
Orectolobiformes
Galeomorphii
Heterodontiformes
Elasmobranchii
(sharks and rays)
Squaliformes
Hexanchiformes
Squalimorphii
Myliobatiformes
Rhinopristiformes
Torpediformes
Batoidea
(rays/skates)
Rajiformes
Chondrichthyes
Holocephali
(chimaeras)

**Supplementary Figure 1.** Detailed dendrogram of the *TLR* genes across different vertebrates. The presence of a *TLR* gene is marked with colored circles (dark gray - TLR1, light grey – TLR6, black – TLR10, green – TLR2, red – TLR3, dark blue – TLR4, purple – TLR5, dark green – TLR7, light green – TLR8, yellow – TLR9, dark purple – TLR13, blue – TLR21, brown – TLR22, green-blue – TLR25, green – TLR27, orange – TLR29, dark orange – TLR30). When multiple gene copies were found, the gene count is indicated inside the corresponding colored circle. Species in which these genes were not identified do not display a colored circle. These results were obtained through exhaustive BLAST searches against available vertebrate genomes. The figure is organized based on the evolutionary relationships among the analyzed vertebrates. Additionally, it incorporates a synteny analysis of each *TLR* gene, mapping their genomic locations across the same set of vertebrates. The analyzed genomic regions are represented by horizontal lines (not to scale), with predicted genes shown as arrows, where the arrowhead indicates gene orientation. Conserved genes shared among vertebrates are consistently color-coded across species, with pseudogenes indicated by a striped background, incomplete genes indicated by a dotted background and species-specific genes represented in white. Gene names are labeled in the first vertebrate species from top to bottom.
