## Supplementary Figure 2 for "Cartilaginous fish inform the lineage-specific evolution and MHC association of the TLR family"

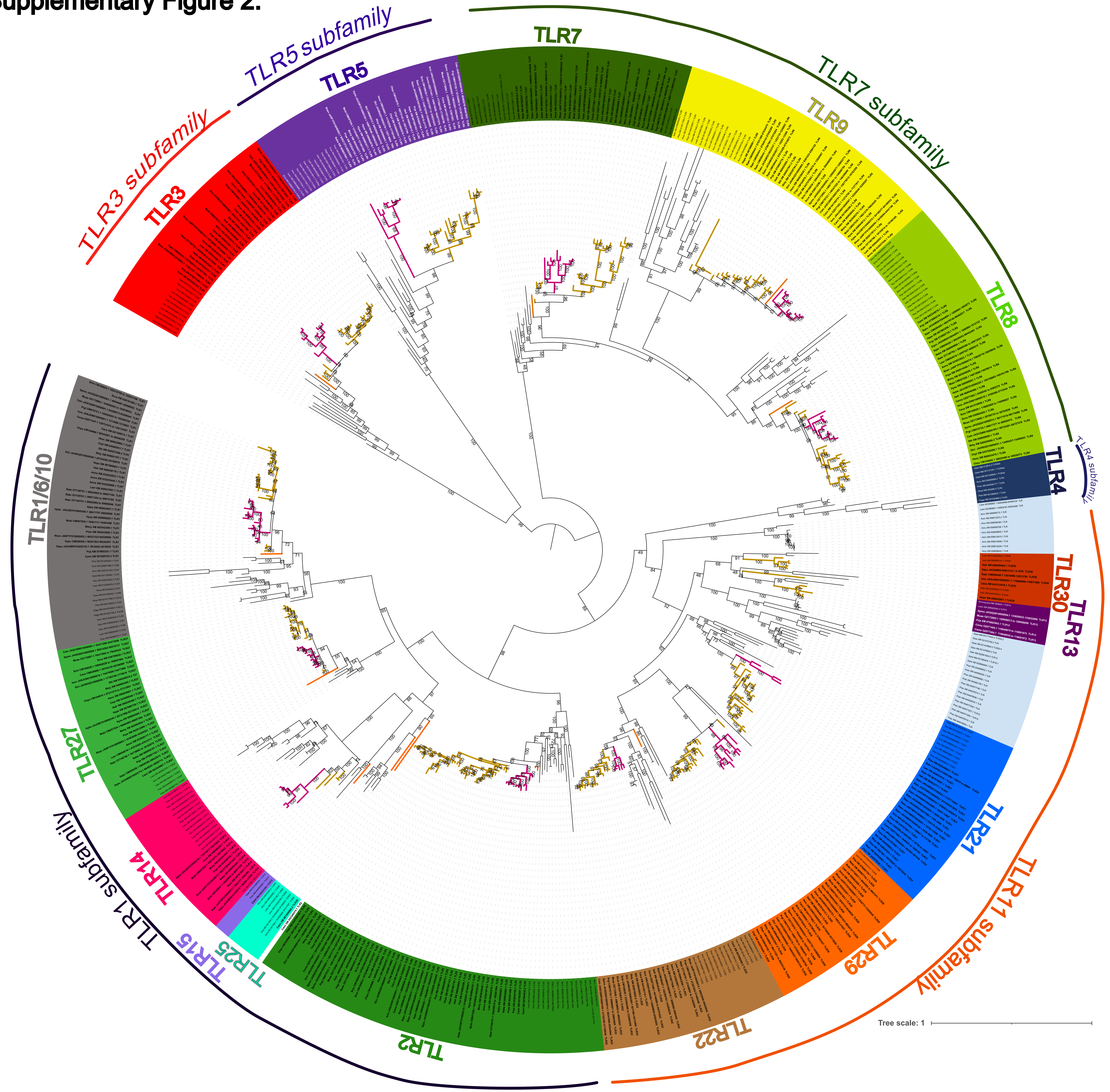

**Supplementary Figure 2.** Phylogenetic tree of Toll-like receptor (TLR) using the elasmobranch species studied, showing the division of TLRs into six subfamilies. The tree was constructed using the maximum likelihood (ML) method based on 572 TIR domain sequences, aligned across 220 amino acid sites, with the JTT+G4 substitution model. The tree is midpoint-rooted. Bootstrap support values are indicated at the internal nodes. Branch colors denote major chondrichthyan lineages: orange for chimaeras, pink for batoids, and brown for sharks.
