## Supplementary Table 1 for "Cartilaginous fish inform the lineage-specific evolution and MHC association of the TLR family"

| Class | Infraclass | Order | Family | Genus | Species | common name | Assembly | GenBank | RefSeq |
|---|---|---|---|---|---|---|---|---|---|
| Chondrichthyes | Selachii | Carcharhiniformes | Carcharhinidae | *Carcharhinus* | *Carcharhinus longimanus* | oceanic whitetip shark | ASM3026437v1 | GCA_030264375.1 | n.a. |
| | | | | *Prionace* | *Prionace glauca* | blue shark | ASM3797433v1 | GCA_037974335.1 | n.a. |
| | | | | *Negaprion* | *Negaprion brevirostris* | lemon shark | ASM3032400v1 | GCA_030324005.1 | n.a. |
| | | | Scyliorhinidae | *Scyliorhinus* | *Scyliorhinus torazame* | cloudy catshark | sScyTor2.1 | GCA_047496885.1 | n.a. |
| | | | | | *Scyliorhinus canicula* | smaller spotted catshark | sScyCan1.1 | GCA_902713615.1 | GCF_902713615.1 |
| | | | Sphyrnidae | *Sphyrna* | *Sphyrna mokarran* | great hammerhead | ASM2467906v1 | GCA_024679065.1 | n.a. |
| | | | Triakidae | *Mustelus* | *Mustelus asterias* | starry smooth-hound | sMusAst1.hap1.1 | GCA_964213995.1 | n.a. |
| | | Lamniformes | Lamnidae | *Carcharodon* | *Carcharodon carcharias* | white shark | sCarCar2.pri | GCA_017639515.1 | GCF_017639515.1 |
| | | | | *Isurus* | *Isurus oxyrinchus* | shortfin mako shark | ASM2677070v1 | GCA_026770705.1 | n.a. |
| | | | Cetorhinidae | *Cetorhinus* | *Cetorhinus maximus* | basking shark | sCetMax3.hap1.1 | GCA_964194155.1 | n.a. |
| | | Orectolobiformes | Hemiscylliidae | *Hemiscyllium* | *Hemiscyllium ocellatum* | epaulette shark | sHemOce1.pat.X.cur. | GCA_020745735.1 | GCF_020745735.1 |
| | | | | *Chiloscyllium* | *Chiloscyllium punctatum* | brownbanded bambooshark | sChiPun1.3 | GCA_047496795.1 | n.a. |
| | | | | | *Chiloscyllium plagiosum* | whitespotted bambooshark | ASM401019v1 | GCA_004010195.1 | GCF_004010195.1 |
| | | | Ginglymostomatidae | *Ginglymostoma* | *Ginglymostoma cirratum* | nurse shark | ASM2413778v1 | GCA_024137785.1 | n.a. |
| | | | Stegostomatidae | *Stegostoma* | *Stegostoma tigrinum* | zebra shark | sSteTig4.hap1 | GCA_030684315.1 | GCF_030684315.1 |
| | | | Rhincodontidae | *Rhincodon* | *Rhincodon typus* | whale shark | sRhiTyp1.1 | GCA_021869965.1 | GCF_021869965.1 |
| | | Heterodontiformes | Heterodontidae | *Heterodontus* | *Heterodontus francisci* | horn shark | sHetFra1.hap1 | GCA_036365525.1 | GCF_036365525.1 |
| | | Pristiophoriformes | Pristiophoridae | *Pristiophorus* | *Pristiophorus japonicus* | Japanese sawshark | sPriJap1.hap1 | GCA_044704955.1 | GCF_044704955.1 |
| | | Squaliformes | Squalidae | *Squalus* | *Squalus suckleyi* | Pacific spiny dogfish | GSC_Ssuck_1.0 | GCA_026260435.1 | n.a. |
| | | | | | *Squalus acanthias* | spiny dogfish | ASM3039002v1 | GCA_030390025.1 | n.a. |
| | | Hexanchiformes | Hexanchidae | *Heptranchias* | *Heptranchias perlo* | sharpnose sevengill shark | sHepPer1.hap1 | GCA_035084215.1 | GCF_035084215.1 |
| | Batoideia | Myliobatiformes | Dasyatidae | *Hypanus* | *Hypanus sabinus* | Atlantic stingray | sHypSab1.hap1 | GCA_030144855.1 | GCF_030144855.1 |
| | | | | | *Hypanus berthalutzae* | Lutz's stingray | ASM3236214v1 | GCA_032362145.1 | n.a. |
| | | | Myliobatidae | *Mobula* | *Mobula hypostoma* | lesser devil ray | sMobHyp1.1 | GCA_963921235.1 | GCA_963921235.1 |
| | | | | | *Mobula birostris* | giant manta | sMobBir1.hap2 | GCA_030035685.1 | n.a. |
| | | Rhinopristiformes | Rhinidae | *Rhina* | *Rhina ancylostomus* | bowmouth guitarfish | ASM3026539v1 | GCA_030265395.1 | n.a. |
| | | | Pristidae | *Pristis* | *Pristis pectinata* | smalltooth sawfish | sPriPec2.1.pri | GCA_009764475.2 | GCF_009764475.1 |
| | | Torpediformes | Narcinidae | *Narcine* | *Narcine bancroftii* | Caribbean electric ray | sNarBan1.hap1 | GCA_036971445.1 | GCF_036971445.1 |
| | | Rajiformes | Rajidae | *Amblyraja* | *Amblyraja radiata* | thorny skate | sAmbRad1.1.pri | GCA_010909765.2 | GCF_010909765.2 |
| | | | | *Raja* | *Raja brachyura* | blonde ray | sRajBra1.1 | GCA_963514005.1 | n.a. |
| | | | | *Leucoraja* | *Leucoraja erinaceus* | little skate | Leri_hhj_1 | GCA_028641065.1 | GCF_028641065.1 |
| | Holocephali | Chimaeriformes | Callorhinchidae | *Callorhinchus* | *Callorhinchus milii* | elephant shark | IMCB_Cmil_1.0 | GCA_018977255.1 | GCF_018977255.1 |
