## Supplementary Table 2 for "Cartilaginous fish inform the lineage-specific evolution and MHC association of the TLR family"

| Class | Infraclass | Order | Family | | Species | Common Name | chromossome | Gene bank location | NCBI Gene symbol | Acession (nt) | Accession (protein) | TLR Subfamily | NCBI Nomenclature | Suggested Nomenclature |
|---|---|---|---|---|---|---|---|---|---|---|---|---|---|---|
| Mammalia | | Primates | Hominidae | Hosa | Homo sapiens | Human | 4 | NC_000004.12 (38787569..38805644, complement) | TLR1 | NM_003263.4 | NP_003254.2 | TLR1 | TLR1 | TLR1 |
| Mammalia | | Primates | Hominidae | Hosa | Homo sapiens | Human | 4 | NC_000004.12 (38822897..38868390, complement) | TLR6 | NM_001394553.1 | NP_001381482.1 | TLR1 | TLR6 | TLR6 |
| Mammalia | | Primates | Hominidae | Hosa | Homo sapiens | Human | 4 | NC_000004.12 (38772238..38782990, complement) | TLR10 | NM_001017388.3 | NP_001017388.1 | TLR1 | TLR10 | TLR10 |
| Mammalia | | Primates | Hominidae | Hosa | Homo sapiens | Human | 4 | NC_000004.12 (153684280..153710637) | TLR2 | NM_001318787.2 | NP_001305716.1 | TLR1 | TLR2 | TLR2 |
| Mammalia | | Primates | Hominidae | Hosa | Homo sapiens | Human | 4 | NC_000004.12 (186069156..186088073) | tlr3 | NM_003265.3 | NP_003256.1 | tlr3 | TLR3 | TLR3 |
| Mammalia | | Primates | Hominidae | Hosa | Homo sapiens | Human | 9 | NC_000009.12 (117704403..117724735) | tlr4 | NM_003266.4 | NP_003257.1 | tlr4 | TLR4 | TLR4 |
| Mammalia | | Primates | Hominidae | Hosa | Homo sapiens | Human | 1 | NC_000001.11 (223109404..223143248, complement) | tlr5 | NM_003268.6 | NP_003259.2 | tlr5 | TLR5 | TLR5 |
| Mammalia | | Primates | Hominidae | Hosa | Homo sapiens | Human | X | NC_000023.11 (12867072..12890361) | TLR7 | NM_016562.4 | NP_057646.1 | TLR7 | TLR7 | TLR7 |
| Mammalia | | Primates | Hominidae | Hosa | Homo sapiens | Human | X | NC_000023.11 (12906620..12923169) | TLR8 | NM_016610.4 | NP_057694.2 | TLR7 | TLR8 | TLR8 |
| Mammalia | | Primates | Hominidae | Hosa | Homo sapiens | Human | 3 | NC_000003.12 (52221080..52225645, complement) | TLR9 | NM_017442.4 | NP_059138.1 | TLR9 | TLR9 | TLR9 |
| Mammalia | | Rodentia | Muridae | Mumu | Mus musculus | mouse | X | NC_000086.8 (105186881..105204099) | tlr13 | NM_205820.1 | NP_991389.1 | TLR11 | TLR13 | TLR13 |
| Birds | | Galliformes | Phasianidae | Gaga | Gallus gallus | Chicken | 4 | NC_052535.1 (69023529..69029651) | TLR1B | NM_001081709.4 | NP_001075178.4 | TLR1 | TLR6 | TLR1B |
| Birds | | Galliformes | Phasianidae | Gaga | Gallus gallus | Chicken | 4 | NC_052535.1 (69035725..69041254) | TLR1A | NM_001007488.5 | NP_001007489.5 | TLR1 | TLR1 | TLR1A |
| Birds | | Galliformes | Phasianidae | Gaga | Gallus gallus | Chicken | 4 | NC_052535.1 (19496158..19502594) | TLR2 | NM_001161650.3 | NP_001155122.3 | TLR1 | TLR2-type-2 | TLR2 |
| Birds | | Galliformes | Phasianidae | Gaga | Gallus gallus | Chicken | 4 | NC_052535.1 (19473260..19495377) | TLR2A | NM_001396826.1 | NP_001383755.1 | TLR1 | TLR2-type-1 | TLR2A |
| Birds | | Galliformes | Phasianidae | Gaga | Gallus gallus | Chicken | 4 | NC_052535.1 (60833481..60843093) | tlr3 | NM_001011691.4 | NP_001011691.4 | tlr3 | TLR3 | TLR3 |
| Birds | | Galliformes | Phasianidae | Gaga | Gallus gallus | Chicken | 17 | NC_052548.1 (4245618..4251245) | TLR4 | NM_001030693.2 | NP_001025864.2 | tlr4 | TLR4 | TLR4 |
| Birds | | Galliformes | Phasianidae | Gaga | Gallus gallus | Chicken | 3 | NC_052534.1 (17528808..17565492) | tlr5 | NM_001398059.1 | NP_001384988.1 | tlr5 | TLR5 | TLR5 |
| Birds | | Galliformes | Phasianidae | Gaga | Gallus gallus | Chicken | 1 | NC_052532.1 (123561394..123568153, complement) | TLR7 | NM_001011688.3 | NP_001011688.2 | TLR7 | TLR7 | TLR7 |
| Birds | | Galliformes | Phasianidae | Gaga | Gallus gallus | Chicken | 3 | NC_052534.1 (3199343..3203514, complement) | TLR15 | NM_001398238.1 | NP_001385167.1 | TLR1 | TLR15 | TLR15 |
| Birds | | Galliformes | Phasianidae | Gaga | Gallus gallus | Chicken | 11 | NC_052542.1 (308996..335580) | TLR21 | NM_001030558.3 | NP_001025729.3 | TLR11 | TLR21 | TLR21 |
| Sauropsida | | Testudines | Trionychidae | Pesi | Pelodiscus sinensis | Chinese soft-shelled turtle | Unplaced scaffold | NW_005856840.1 (1893154..1910131) | LOC102460455 | XM_006129523.3 | XP_006129585.1 | TLR1 | TLR1 | TLR1 |
| Sauropsida | | Testudines | Trionychidae | Pesi | Pelodiscus sinensis | Chinese soft-shelled turtle | Unplaced scaffold | NW_005856840.1 (1925756..1937839) | LOC102445089 | XM_025187956.1 | XP_025043741.1 | TLR1 | TLR10 | TLR10 |
| Sauropsida | | Testudines | Trionychidae | Pesi | Pelodiscus sinensis | Chinese soft-shelled turtle | Unplaced scaffold | NW_005851767.1 (809601..814945, complement) | LOC102457570 | XM_025185305.1 | XP_025041090.1 | TLR1 | TLR2-type-2 | TLR2 |
| Sauropsida | | Testudines | Trionychidae | Pesi | Pelodiscus sinensis | Chinese soft-shelled turtle | Unplaced scaffold | NW_005851767.1 (797139..814948, complement) | TLR2 | XM_025185298.1 | XP_025041083.1 | TLR1 | TLR2 | TLR2 |
| Sauropsida | | Testudines | Trionychidae | Pesi | Pelodiscus sinensis | Chinese soft-shelled turtle | Unplaced scaffold | NW_005856900.1 (78932..83893) | LOC102458059 | XM_025188077.1 | XP_025043862.1 | TLR1 | TLR2 | TLR2 |
| Sauropsida | | Testudines | Trionychidae | Pesi | Pelodiscus sinensis | Chinese soft-shelled turtle | Unplaced scaffold | NW_005856643.1 (5632855..5653455, complement) | tlr3 | XM_014576247.2 | XP_014431733.1 | tlr3 | TLR3 | TLR3 |
| Sauropsida | | Testudines | Trionychidae | Pesi | Pelodiscus sinensis | Chinese soft-shelled turtle | Unplaced scaffold | NW_005854608.1 (1746449..1761287) | tlr4 | NM_001286933.1 | NP_001273862.1 | tlr4 | TLR4 | TLR4 |
| Sauropsida | | Testudines | Trionychidae | Pesi | Pelodiscus sinensis | Chinese soft-shelled turtle | Unplaced scaffold | NW_005853122.1 (3093089..3128534, complement) | tlr5 | XM_006115600.3 | XP_006115662.2 | tlr5 | TLR5 | TLR5 |
| Sauropsida | | Testudines | Trionychidae | Pesi | Pelodiscus sinensis | Chinese soft-shelled turtle | Unplaced scaffold | NW_005859004.1 (591037..596901) | LOC102463208 | XM_014580301.2 | XP_014435787.1 | tlr5 | TLR5 | TLR5 |
| Sauropsida | | Testudines | Trionychidae | Pesi | Pelodiscus sinensis | Chinese soft-shelled turtle | Unplaced scaffold | NW_005855059.1 (831359..844040) | TLR7 | XM_014573459.2 | XP_014428945.1 | TLR7 | TLR7 | TLR7 |
| Sauropsida | | Testudines | Trionychidae | Pesi | Pelodiscus sinensis | Chinese soft-shelled turtle | Unplaced scaffold | NW_005855059.1 (856293..864093) | TLR8 | XM_006122845.3 | XP_006122907.1 | TLR7 | TLR8 | TLR8 |
| Sauropsida | | Testudines | Trionychidae | Pesi | Pelodiscus sinensis | Chinese soft-shelled turtle | Unplaced scaffold | NW_005855059.1 (868873..881735) | TLR8 | XM_014573452.2 | XP_014428938.1 | TLR7 | TLR8 | TLR8 |
| Sauropsida | | Testudines | Trionychidae | Pesi | Pelodiscus sinensis | Chinese soft-shelled turtle | Unplaced scaffold | NW_005855059.1 (886568..898015) | TLR8 | XM_006122794.3 | XP_006122856.1 | TLR7 | TLR8 | TLR8 |
| Sauropsida | | Testudines | Trionychidae | Pesi | Pelodiscus sinensis | Chinese soft-shelled turtle | Unplaced scaffold | NW_005854103.1 (67239..75858, complement) | LOC102444127 | XM_014571799.2 | XP_014427285.2 | TLR9 | TLR9 | TLR9 |
| Sauropsida | | Testudines | Trionychidae | Pesi | Pelodiscus sinensis | Chinese soft-shelled turtle | Unplaced scaffold | NW_005855764.1 (5767..8537, complement) | LOC102453419 | x | x | tlr11 | TLR13 | x |
| Sauropsida | | Testudines | Trionychidae | Pesi | Pelodiscus sinensis | Chinese soft-shelled turtle | Unplaced scaffold | NW_005854040.1 (98916..113445) | LOC102445063 | XM_014571663.2 | XP_014427149.1 | TLR1 | TLR2 | TLR18 |
| Sauropsida | | Testudines | Trionychidae | Pesi | Pelodiscus sinensis | Chinese soft-shelled turtle | Unplaced scaffold | NW_005871014.1 (4898304..4902059) | LOC102451718 | XM_006138102.3 | XP_006138164.2 | TLR11 | TLR13 | TLR22 |
| Sauropsida | | Testudines | Trionychidae | Pesi | Pelodiscus sinensis | Chinese soft-shelled turtle | Unplaced scaffold | NW_005871014.1 (4907944..4911699) | LOC102461996 | XM_006138146.3 | XP_006138208.2 | TLR11 | TLR13 | TLR22 |
| Amphibia | Lissamphibia | Anura | Pipidae | Xetr | Xenopus tropicalis | tropical clawed frog | 1 | NC_030677.2 (35759160..35766278) | TLR1 | XM_018095354.2 | XP_017950843.2 | TLR1 | TLR1 | TLR1 |
| Amphibia | Lissamphibia | Anura | Pipidae | Xetr | Xenopus tropicalis | tropical clawed frog | 1 | NC_030677.2 (35759161..35780429) | TLR6 | XM_018095356.2 | XP_017950845.1 | TLR1 | TLR6 | TLR6 |
| Amphibia | Lissamphibia | Anura | Pipidae | Xetr | Xenopus tropicalis | tropical clawed frog | 1 | NC_030677.2 (35787056..35792199, complement) | XB5889249<br>LOC100495326 | XM_018095359.2 | XP_017950848.2 | TLR1 | TLR1 | TLR1 |
| Amphibia | Lissamphibia | Anura | Pipidae | Xetr | Xenopus tropicalis | tropical clawed frog | 1 | NC_030677.2 (55006444..55009954, complement) | LOC100485927 | XM_018091005.2 | XP_017946494.1 | TLR1 | TLR2-type-2 | TLR2 |
| Amphibia | Lissamphibia | Anura | Pipidae | Xetr | Xenopus tropicalis | tropical clawed frog | 1 | NC_030677.2 (54994299..55002782, complement) | tlr2 | XM_002933491.5 | XP_002933537.2 | TLR1 | TLR2-type-2 | TLR2 |
| Amphibia | Lissamphibia | Anura | Pipidae | Xetr | Xenopus tropicalis | tropical clawed frog | 3 | NC_030679.2 (25058968..25060979, complement) | LOC116409470 | XM_031898125.1 | XP_031753985.1 | TLR1 | TLR2 | TLR2 |
| Amphibia | Lissamphibia | Anura | Pipidae | Xetr | Xenopus tropicalis | tropical clawed frog | 1 | NC_030677.2 (44170821..44189996, complement) | tlr3 | XM_002934402.4 | XP_002934448.3 | tlr3 | TLR3 | TLR3 |
| Amphibia | Lissamphibia | Anura | Pipidae | Xetr | Xenopus tropicalis | tropical clawed frog | 8 | NC_030684.2(33480842 to 33482529) | tlr4 | x | x | tlr4 | TLR4 | TLR4 |
| Amphibia | Lissamphibia | Anura | Pipidae | Xetr | Xenopus tropicalis | tropical clawed frog | 4 | NC_030680.2 (31584390..31588805, complement) | TLR5S | XM_002937504.5 | XP_002937550.2 | TLR5 | TLR5 | TLR5s |
| Amphibia | Lissamphibia | Anura | Pipidae | Xetr | Xenopus tropicalis | tropical clawed frog | 5 | NC_030681.2 (9338881..9428216) | tlr5 | NM_001078891.1 | NP_001072359.1 | x | TLR5 | SUSD4 |
| Amphibia | Lissamphibia | Anura | Pipidae | Xetr | Xenopus tropicalis | tropical clawed frog | 5 | NC_030681.2 (9433774..9449118) | LOC100496647 | XM_002940696.5 | XP_002940742.3 | tlr5 | uncharacterized | TLR5 |
| Amphibia | Lissamphibia | Anura | Pipidae | Xetr | Xenopus tropicalis | tropical clawed frog | 2 | NC_030678.2 (106068123..106076766, complement) | TLR7 | NM_001127411.1 | NP_001120883.1 | TLR7 | TLR7 | TLR7 |
| Amphibia | Lissamphibia | Anura | Pipidae | Xetr | Xenopus tropicalis | tropical clawed frog | 2 | NC_030678.2 (106044826..106048041, complement) | TLR8 | XM_002933813.2 | XP_002933859.2 | TLR7 | TLR8 | TLR8 |
| Amphibia | Lissamphibia | Anura | Pipidae | Xetr | Xenopus tropicalis | tropical clawed frog | 2 | NC_030678.2 (106030670..106041028, complement) | LOC100495962 | XM_004912345.2 | XP_004912402.2 | TLR7 | TLR8 | TLR8 |
| Amphibia | Lissamphibia | Anura | Pipidae | Xetr | Xenopus tropicalis | tropical clawed frog | 4 | NC_030680.2 (126164531..126168151) | tlr9 | XM_018093220.2 | XP_017948709.2 | TLR7 | TLR9 | TLR9 |
| Amphibia | Lissamphibia | Anura | Pipidae | Xetr | Xenopus tropicalis | tropical clawed frog | 7 | NC_030683.2 (103837497..103842669, complement) | LOC100498493 | XM_002941824.5 | XP_002941870.2 | TLR11 | TLR12 | TLR12 |
| Amphibia | Lissamphibia | Anura | Pipidae | Xetr | Xenopus tropicalis | tropical clawed frog | 8 | NC_030684.2 (141632539..141638022, complement) | tlr14.2 | XM_012954675.2 | XP_012810129.1 | TLR1 | TLR14.2 | TLR18 |
| Amphibia | Lissamphibia | Anura | Pipidae | Xetr | Xenopus tropicalis | tropical clawed frog | 8 | NC_030684.2 (141653294..141658347) | LOC100485358 | XM_002943050.5 | XP_002943096.1 | TLR1 | TLR2 | TLR18 |
| Amphibia | Lissamphibia | Anura | Pipidae | Xetr | Xenopus tropicalis | tropical clawed frog | 8 | NC_030684.2 (141676201..141685630) | TLR14.3 | XM_004920394.4 | XP_004920451.1 | TLR1 | TLR14.3 | TLR18 |
| Amphibia | Lissamphibia | Anura | Pipidae | Xetr | Xenopus tropicalis | tropical clawed frog | 8 | NC_030684.2 (142502798..142503779, complement) | LOC116406628 | XM_031891013.1 | XP_031746873.1 | TLR1 | TLR6 | TLR18 |
| Amphibia | Lissamphibia | Anura | Pipidae | Xetr | Xenopus tropicalis | tropical clawed frog | 8 | NC_030684.2 (142503762..142523565, complement) | LOC100487973 | XM_031891012.1 | XP_031746872.1 | TLR1 | TLR2 | TLR18 |
| Amphibia | Lissamphibia | Anura | Pipidae | Xetr | Xenopus tropicalis | tropical clawed frog | 7 | NC_030683.2 (107571316..107574360, complement) | LOC100497009 | XM_002936397.4 | XP_002936443.3 | TLR11 | TLR13 | TLR21 |
| Amphibia | Lissamphibia | Anura | Pipidae | Xetr | Xenopus tropicalis | tropical clawed frog | 3 | NC_030679.2 (42115679..42122247, complement) | LOC100485706 | XM_002942535.5 | XP_002942581.2 | TLR11 | TLR22 | TLR22 |
| Amphibia | Lissamphibia | Anura | Pipidae | Xetr | Xenopus tropicalis | tropical clawed frog | 7 | NC_030683.2 (114625278..114637558, complement) | LOC100485164 | XM_002935001.5 | XP_002935047.2 | TLR11 | TLR13 | TLR30 |
| Osteichthyes | Sarcopterygii | Lepidosireniformes | Protopteridae | Pran | Protopterus annectens | West African lungfish | 2.part1 | NC_056729.1 (294715822..294756417, complement) | LOC122794029 | XM_044062141.1 | XP_043918076.1 | TLR1 | TLR1 | TLR1 |
| Osteichthyes | Sarcopterygii | Lepidosireniformes | Protopteridae | Pran | Protopterus annectens | West African lungfish | 2.part1 | NC_056729.1 (291896176..291900237, complement) | LOC122794028 | XM_044062139.1 | XP_043918074.1 | TLR11 | TLR13 | TLR13 |
| Osteichthyes | Sarcopterygii | Lepidosireniformes | Protopteridae | Pran | Protopterus annectens | West African lungfish | 2.part1 | NC_056729.1 (664550331..664552989) | tlr2 | XM_044062406.1 | XP_043918341.1 | TLR1 | TLR2 | TLR2 |
| Osteichthyes | Sarcopterygii | Lepidosireniformes | Protopteridae | Pran | Protopterus annectens | West African lungfish | 4.part0 | NC_056733.1 (1080099481..1080215402, complement) | LOC122800166 | XM_044069680.1 | XP_043925615.1 | TLR1 | TLR2 | TLR2 |
| Osteichthyes | Sarcopterygii | Lepidosireniformes | Protopteridae | Pran | Protopterus annectens | West African lungfish | 2.part1 | NC_056729.1 (865559986..865869435) | tlr3 | XM_044062513.1 | XP_043918448.1 | tlr3 | TLR3 | TLR3 |
| Osteichthyes | Sarcopterygii | Lepidosireniformes | Protopteridae | Pran | Protopterus annectens | West African lungfish | 16.part0 | NC_056750.1 (737462855..737553413, complement) | tlr4 | XM_044058177.1 | XP_043914112.1 | tlr4 | TLR4 | TLR4 |
| Osteichthyes | Sarcopterygii | Lepidosireniformes | Protopteridae | Pran | Protopterus annectens | West African lungfish | 4.part0 | NC_056733.1 (1140202329..1140206079, complement) | LOC122799804 | XM_044069109.1 | XP_043925044.1 | tlr5 | TLR5 | TLR5 |
| Osteichthyes | Sarcopterygii | Lepidosireniformes | Protopteridae | Pran | Protopterus annectens | West African lungfish | 6.part1 | NC_056738.1 (401181496..401195844) | LOC122805591 | XM_044075816.1 | XP_043931751.1 | tlr5 | TLR5 | TLR5 |
| Osteichthyes | Sarcopterygii | Lepidosireniformes | Protopteridae | Pran | Protopterus annectens | West African lungfish | 1.part2 | NC_056727.1 (631889315..632539736) | LOC122790524 | XM_044058082.1 | XP_043914017.1 | tlr5 | TLR5 | TLR5 |
| Osteichthyes | Sarcopterygii | Lepidosireniformes | Protopteridae | Pran | Protopterus annectens | West African lungfish | 5.part0 | NC_056735.1 (693790959..693834385, complement) | LOC122802180 | XM_044071849.1 | XP_043927784.1 | TLR7 | TLR7 | TLR7 |
| Osteichthyes | Sarcopterygii | Lepidosireniformes | Protopteridae | Pran | Protopterus annectens | West African lungfish | 5.part0 | NC_056735.1 (693095100..693339419, complement) | LOC122802765 | XM_044072541.1 | XP_043928476.1 | TLR7 | TLR8 | TLR8 |
| Osteichthyes | Sarcopterygii | Lepidosireniformes | Protopteridae | Pran | Protopterus annectens | West African lungfish | 7.part1 | NC_056740.1 (491250595..491254491) | LOC122807839 | XM_044078937.1 | XP_043934872.1 | TLR7 | TLR7 | TLR9 |
| Osteichthyes | Sarcopterygii | Lepidosireniformes | Protopteridae | Pran | Protopterus annectens | West African lungfish | 7.part1 | NC_056740.1 (491442081..491568292, complement) | LOC122807840 | XM_044078938.1 | XP_043934873.1 | TLR7 | TLR7 | TLR9 |
| Osteichthyes | Sarcopterygii | Lepidosireniformes | Protopteridae | Pran | Protopterus annectens | West African lungfish | 14.part0 | NC_056748.1 (160889942..160894179) | LOC122788635 | XM_044055534.1 | XP_043911469.1 | TLR11 | TLR13 | |
| Osteichthyes | Sarcopterygii | Lepidosireniformes | Protopteridae | Pran | Protopterus annectens | West African lungfish | 4.part0 | NC_056733.1 (1484417340..1484419620, complement) | LOC122800183 | XM_044069690.1 | XP_043925625.1 | TLR11 | TLR13 | |
| Osteichthyes | Sarcopterygii | Lepidosireniformes | Protopteridae | Pran | Protopterus annectens | West African lungfish | 4.part0 | NC_056733.1 (1484028490..1484031755, complement) | LOC122799923 | XM_044069325.1 | XP_043925260.1 | TLR11 | TLR13 | |
| Osteichthyes | Sarcopterygii | Lepidosireniformes | Protopteridae | Pran | Protopterus annectens | West African lungfish | 4.part0 | NC_056733.1 (1727003117..1727005978) | LOC122800482 | XM_044069922.1 | XP_043925857.1 | TLR11 | TLR13 | |
| Osteichthyes | Sarcopterygii | Lepidosireniformes | Protopteridae | Pran | Protopterus annectens | West African lungfish | 7.part0 | NC_056739.1 (566460724..566466050, complement) | LOC122806594 | XM_044077146.1 | XP_043933081.1 | TLR11 | TLR13 | |
| Osteichthyes | Sarcopterygii | Lepidosireniformes | Protopteridae | Pran | Protopterus annectens | West African lungfish | 7.part0 | NC_056739.1 (693759599..693760640) | LOC122805838 | XM_044076092.1 | XP_043932027.1 | TLR11 | TLR13 | |
| Osteichthyes | Sarcopterygii | Lepidosireniformes | Protopteridae | Pran | Protopterus annectens | West African lungfish | 7.part0 | NC_056739.1 (694583371..694586100) | LOC122806076 | XM_044076314.1 | XP_043932249.1 | TLR11 | TLR13 | |
| Osteichthyes | Sarcopterygii | Lepidosireniformes | Protopteridae | Pran | Protopterus annectens | West African lungfish | 7.part0 | NC_056739.1 (695403432..695407344, complement) | LOC122806639 | XM_044077227.1 | XP_043933162.1 | TLR11 | TLR12 | |
| Osteichthyes | Sarcopterygii | Lepidosireniformes | Protopteridae | Pran | Protopterus annectens | West African lungfish | 7.part0 | NC_056739.1 (696910765..696913737) | LOC122806077 | XM_044076315.1 | XP_043932250.1 | TLR11 | TLR13 | |
| Osteichthyes | Sarcopterygii | Lepidosireniformes | Protopteridae | Pran | Protopterus annectens | West African lungfish | 7.part0 | NC_056739.1 (696916430..696917665) | LOC122805839 | XM_044076093.1 | XP_043932028.1 | TLR11 | TLR13 | |
| Osteichthyes | Sarcopterygii | Lepidosireniformes | Protopteridae | Pran | Protopterus annectens | West African lungfish | 7.part0 | NC_056739.1 (698741113..698744987) | LOC122806640 | XM_044077228.1 | XP_043933163.1 | TLR11 | TLR12 | |
| Osteichthyes | Sarcopterygii | Lepidosireniformes | Protopteridae | Pran | Protopterus annectens | West African lungfish | 7.part0 | NC_056739.1 (700602015..700604338) | LOC122806642 | XM_044077229.1 | XP_043933164.1 | TLR11 | TLR13 | |
| Osteichthyes | Sarcopterygii | Lepidosireniformes | Protopteridae | Pran | Protopterus annectens | West African lungfish | 1.part2 | NC_056727.1 (106739928..106743970) | LOC122788141 | XM_044055095.1 | XP_043911030.1 | TLR1 | TLR2 type-2 | TLR15 |
| Osteichthyes | Sarcopterygii | Lepidosireniformes | Protopteridae | Pran | Protopterus annectens | West African lungfish | 8.part0 | NC_056741.1 (458009525..458525788) | LOC122808872 | XM_044079973.1 | XP_043935908.1 | TLR1 | TLR2 | TLR18 |
| Osteichthyes | Sarcopterygii | Lepidosireniformes | Protopteridae | Pran | Protopterus annectens | West African lungfish | 8.part0 | NC_056741.1 (458851206..458941543) | LOC122808873 | XM_044079976.1 | XP_043935911.1 | TLR1 | TLR2 | TLR18 |
| Osteichthyes | Sarcopterygii | Lepidosireniformes | Protopteridae | Pran | Protopterus annectens | West African lungfish | 8.part0 | NC_056741.1 (1287040208..1287418842, complement) | LOC122809365 | XM_044080803.1 | XP_043936738.1 | tlr11 | TLR13 | TLR21 |
| Osteichthyes | Sarcopterygii | Lepidosireniformes | Protopteridae | Pran | Protopterus annectens | West African lungfish | 3.part1 | NC_056732.1 (1741264050..1741267294, complement) | LOC122798906 | XM_044068004.1 | XP_043923939.1 | TLR11 | TLR13 | TLR22 |
| Osteichthyes | Sarcopterygii | Lepidosireniformes | Protopteridae | Pran | Protopterus annectens | West African lungfish | 10.part0 | NC_056744.1 (404478878..404481971) | LOC122814164 | XM_044086776.1 | XP_043942711.1 | TLR1 | TLR2 type-2 | TLR27 |
| Osteichthyes | Sarcopterygii | Lepidosireniformes | Protopteridae | Pran | Protopterus annectens | West African lungfish | 10.part0 | NC_056744.1 (404605651..404665367) | LOC122813159 | XM_044085315.1 | XP_043941250.1 | TLR1 | TLR2 | TLR27 |
| Osteichthyes | Sarcopterygii | Lepidosireniformes | Protopteridae | Pran | Protopterus annectens | West African lungfish | 1.part0 | NC_056726.1 (1012048901..1012446397, complement) | LOC122808725 | XM_044079734.1 | XP_043935669.1 | TLR11 | TLR13 | TLR30 |
| Osteichthyes | Actinistia | Coelacanthiformes | Latimeriidae | Lach | Latimeria chalumnae | coelacanth | 1 | NC_088139.1 (347493021..347495354, complement) | TLR1 | XM_006004367.3 | XP_006004429.3 | TLR1 | TLR1 | TLR1 |
| Osteichthyes | Actinistia | Coelacanthiformes | Latimeriidae | Lach | Latimeria chalumnae | coelacanth | 1 | NC_088139.1 (295627723..295634482) | tlr2 | XM_014493815.1 | XP_014349301.1 | TLR1 | TLR2 | TLR2 |
| Osteichthyes | Actinistia | Coelacanthiformes | Latimeriidae | Lach | Latimeria chalumnae | coelacanth | 1 | NC_088139.1 (323302249..323348159) | tlr3 | XM_064562465.1 | XP_064418535.1 | tlr3 | TLR3 | TLR3 |
| Osteichthyes | Actinistia | Coelacanthiformes | Latimeriidae | Lach | Latimeria chalumnae | coelacanth | 3 | NC_088141.1 (266199641..266201563, complement) | x | x | x | tlr5 | x | TLR5 |
| Osteichthyes | Actinistia | Coelacanthiformes | Latimeriidae | Lach | Latimeria chalumnae | coelacanth | 3 | NC_088141.1 (266233821..266236415, complement) | LOC102366872 | XM_064555280.1 | XP_064411350.1 | tlr5 | tlr5 | TLR5 |
| Osteichthyes | Actinistia | Coelacanthiformes | Latimeriidae | Lach | Latimeria chalumnae | coelacanth | 4 | NC_088142.1 (80755256..80766476, complement) | TLR7 | XM_064556896.1 | XP_064412966.1 | TLR7 | TLR7 | TLR7 |
| Osteichthyes | Actinistia | Coelacanthiformes | Latimeriidae | Lach | Latimeria chalumnae | coelacanth | 4 | NC_088142.1 (80739380..80742373, complement) | LOC102356471 | XM_064556895.1 | XP_064412965.1 | TLR7 | TLR7 | TLR8 |
| Osteichthyes | Actinistia | Coelacanthiformes | Latimeriidae | Lach | Latimeria chalumnae | coelacanth | 4 | NC_088142.1 (80689958..80728011, complement) | LOC135356425 | XM_064556894.1 | XP_064412964.1 | TLR7 | TLR8 | TLR8 |

| | | | | | | | | | | | | | |
|---|---|---|---|---|---|---|---|---|---|---|---|---|---|
| Osteichthyes | Actinistia | Coelacanthiformes | Latimeriidae | Lach | Latimeria chalumnae | coelacanth | 14 | NC_088152.1 (14194418..14197320, complement) | tlr9 | XM_005987092.1 | XP_005987154.1 | TLR7 | TLR9 | TLR9 |
| Osteichthyes | Actinistia | Coelacanthiformes | Latimeriidae | Lach | Latimeria chalumnae | coelacanth | 15 | NC_088153.1 (7331059..7336364, complement) | LOC102347180 | XM_006000240.3 | XP_006000302.1 | TLR11 | TLR13 | TLR13 |
| Osteichthyes | Actinistia | Coelacanthiformes | Latimeriidae | Lach | Latimeria chalumnae | coelacanth | 6 | NC_088144.1 (138572809..138575862) | LOC102359898 | XM_064560214.1 | XP_064416284.1 | TLR11 | TLR13 | TLR13 |
| Osteichthyes | Actinistia | Coelacanthiformes | Latimeriidae | Lach | Latimeria chalumnae | coelacanth | 6 | NC_088144.1 (138597020..138599773, complement) | LOC102359627 | XM_014493952.2 | XP_014349438.2 | TLR11 | TLR13 | TLR13 |
| Osteichthyes | Actinistia | Coelacanthiformes | Latimeriidae | Lach | Latimeria chalumnae | coelacanth | 6 | NC_088144.1 (112830128..112869666, complement) | LOC102355189 | XM_005991679.3 | XP_005991741.2 | TLR1 | TLR18 | TLR18 |
| Osteichthyes | Actinistia | Coelacanthiformes | Latimeriidae | Lach | Latimeria chalumnae | coelacanth | 25 | NC_088163.1 (24621432..24624479, complement) | TLR21 | XM_005997988.1 | XP_005998050.1 | tlr11 | TLR21 | TLR21 |
| Osteichthyes | Actinistia | Coelacanthiformes | Latimeriidae | Lach | Latimeria chalumnae | coelacanth | 6 | NC_088144.1 (46056506..46058923) | LOC102350366 | XM_005997279.1 | XP_005997341.1 | TLR1 | TLR2 type-2 | TLR27 |
| Osteichthyes | Actinistia | Coelacanthiformes | Latimeriidae | Lach | Latimeria chalumnae | coelacanth | 9 | NC_088147.1 (11130103..11154098, complement) | LOC102349691 | XM_064563319.1 | XP_064419389.1 | TLR11 | TLR13 | TLR29 |
| Actinopterygii | Teleostei | Cypriniformes | Cyprinidae | Dare | Danio rerio | zebrafish | 14 | NC_007125.7 (732606..736575, complement) | tlr1 | NM_001130593.1 | NP_001124065.1 | TLR1 | TLR1 | TLR1 |
| Actinopterygii | Teleostei | Cypriniformes | Cyprinidae | Dare | Danio rerio | zebrafish | 1 | NC_007112.7 (8983965..8990541) | tlr2 | NM_212812.1 | NP_997977.1 | TLR1 | TLR2 | TLR2 |
| Actinopterygii | Teleostei | Cypriniformes | Cyprinidae | Dare | Danio rerio | zebrafish | 1 | NC_007112.7 (17892944..17898227) | tlr3 | NM_001013269.3 | NP_001013287.2 | tlr3 | TLR3 | TLR3 |
| Actinopterygii | Teleostei | Cypriniformes | Cyprinidae | Dare | Danio rerio | zebrafish | 13 | NC_007124.7 (18514663..18517495) | tlr4ba | NM_001131051.1 | NP_001124523.1 | tlr4 | TLR4 | TLR4 |
| Actinopterygii | Teleostei | Cypriniformes | Cyprinidae | Dare | Danio rerio | zebrafish | 13 | NC_007124.7 (18520293..18523630) | TLR4al | NM_001328605.1 | NP_001315534.1 | tlr4 | TLR4 | TLR4 |
| Actinopterygii | Teleostei | Cypriniformes | Cyprinidae | Dare | Danio rerio | zebrafish | 13 | NC_007124.7 (18523654..18528859) | tlr4bb | NM_212813.2 | NP_997978.2 | tlr4 | TLR4 | TLR4 |
| Actinopterygii | Teleostei | Cypriniformes | Cyprinidae | Dare | Danio rerio | zebrafish | 20 | NC_007131.7 (51471979..51476180) | tlr5b | NM_001130595.2 | NP_001124067.2 | tlr5 | TLR5b | TLR5b |
| Actinopterygii | Teleostei | Cypriniformes | Cyprinidae | Dare | Danio rerio | zebrafish | 20 | NC_007131.7 (51478965..51485659) | tlr5a | XM_001919017.7 | XP_001919052.2 | tlr5 | TLR5a | TLR5a |
| Actinopterygii | Teleostei | Cypriniformes | Cyprinidae | Dare | Danio rerio | zebrafish | 9 | NC_007120.7 (54114154..54147139) | TLR7 | XM_021479060.2 | XP_021334735.2 | TLR7 | TLR7 | TLR7 |
| Actinopterygii | Teleostei | Cypriniformes | Cyprinidae | Dare | Danio rerio | zebrafish | 9 | NC_007120.7 (54147212..54147973) | LOC100332713 | x | x | tlr7 | x | |
| Actinopterygii | Teleostei | Cypriniformes | Cyprinidae | Dare | Danio rerio | zebrafish | 9 | NC_007120.7 (54147973..54148739) | LOC137496515 | x | x | tlr7 | x | |
| Actinopterygii | Teleostei | Cypriniformes | Cyprinidae | Dare | Danio rerio | zebrafish | 9 | NC_007120.7 (54151062..54163965) | LOC100332583 | XM_002665908.7 | XP_002665954.4 | TLR7 | TLR8 | TLR8 |
| Actinopterygii | Teleostei | Cypriniformes | Cyprinidae | Dare | Danio rerio | zebrafish | 10 | NC_007121.7 (43838297..43850090, complement) | tlr8b | NM_001386709.1 | NP_001373638.1 | TLR7 | TLR8b | TLR8 |
| Actinopterygii | Teleostei | Cypriniformes | Cyprinidae | Dare | Danio rerio | zebrafish | Unplaced Scaffold | NW_003336636.1 (6147..13421, complement) | TLR8a | XM_001920559.7 | XP_001920594.5 | TLR7 | TLR8a | TLR8 |
| Actinopterygii | Teleostei | Cypriniformes | Cyprinidae | Dare | Danio rerio | zebrafish | 8 | NC_007119.7 (53730368..53733659, complement) | tlr9 | NM_001130594.1 | NP_001124066.1 | TLR7 | | TLR9 |
| Actinopterygii | Teleostei | Cypriniformes | Cyprinidae | Dare | Danio rerio | zebrafish | 9 | NC_007120.7 (27346408..27349337) | LOC103911657 | XM_021472192.2 | XP_021327867.1 | TLR11 | TLR13 | TLR13 |
| Actinopterygii | Teleostei | Cypriniformes | Cyprinidae | Dare | Danio rerio | zebrafish | 9 | NC_007120.7 (27368886..27371771) | LOC572462 | XM_021478955.2 | XP_021334630.1 | TLR11 | TLR13 | TLR13 |
| Actinopterygii | Teleostei | Cypriniformes | Cyprinidae | Dare | Danio rerio | zebrafish | 16 | NC_007127.7 (22739010..22749551, complement) | LOC137487195 | XM_002664846.7 | XP_002664892.4 | TLR11 | TLR13 | TLR13 |
| Actinopterygii | Teleostei | Cypriniformes | Cyprinidae | Dare | Danio rerio | zebrafish | 16 | NC_007127.7 (29391791..29405133, complement) | tlr18 | NM_001089350.1 | NP_001082819.1 | tlr1 | TLR18 | TLR18 |
| Actinopterygii | Teleostei | Cypriniformes | Cyprinidae | Dare | Danio rerio | zebrafish | 16 | NC_007127.7 (22722196..22730375, complement) | tlr19.1 | NM_001365424.1 | NP_001352353.1 | TLR11 | TLR19 | TLR19 |
| Actinopterygii | Teleostei | Cypriniformes | Cyprinidae | Dare | Danio rerio | zebrafish | 9 | NC_007120.7 (27352298..27355294) | tlr20.2 | NM_001177443.2 | NP_001170914.2 | TLR11 | TLR20.2 | TLR20.2 |
| Actinopterygii | Teleostei | Cypriniformes | Cyprinidae | Dare | Danio rerio | zebrafish | 9 | NC_007120.7 (27376776..27379777) | tlr20.4 | XM_021478954.2 | XP_021334629.1 | TLR11 | TLR20.4 | TLR20.4 |
| Actinopterygii | Teleostei | Cypriniformes | Cyprinidae | Dare | Danio rerio | zebrafish | 16 | NC_007127.7 (11785898..11789108, complement) | tlr21 | NM_001199335.1 | NP_001186264.1 | tlr11 | TLR21 | TLR21 |
| Actinopterygii | Teleostei | Cypriniformes | Cyprinidae | Dare | Danio rerio | zebrafish | 21 | NC_007132.7 (38849851..38853737, complement) | TLR22 | NM_001128675.2 | NP_001122147.2 | TLR11 | TLR22 | TLR22 |
| Actinopterygii | Teleostei | Tetraodontiformes | Tetraodontidae | Taru | Takifugu rubripes | torafugu | 14 | NC_042298.1 (6131312..6134591) | tlr1 | XM_003970363.3 | XP_003970412.2 | TLR1 | | TLR1 |
| Actinopterygii | Teleostei | Tetraodontiformes | Tetraodontidae | Taru | Takifugu rubripes | torafugu | 8 | NC_042292.1 (19585212..19589837, complement) | tlr2 | XM_011617904.2 | XP_011616206.2 | TLR1 | | TLR2 |
| Actinopterygii | Teleostei | Tetraodontiformes | Tetraodontidae | Taru | Takifugu rubripes | torafugu | 17 | NC_042301.1 (12191536..12196342) | tlr3 | XM_003972308.3 | XP_003972357.2 | tlr3 | | TLR3 |
| Actinopterygii | Teleostei | Tetraodontiformes | Tetraodontidae | Taru | Takifugu rubripes | torafugu | 16 | NC_042300.1 (12521135..12525025, complement) | LOC101062562 | XM_029849771.1 | XP_029705631.1 | tlr5 | | |
| Actinopterygii | Teleostei | Tetraodontiformes | Tetraodontidae | Taru | Takifugu rubripes | torafugu | 16 | NC_042300.1 (1726017..1729565, complement) | LOC115253071 | x | x | tlr5 | | |
| Actinopterygii | Teleostei | Tetraodontiformes | Tetraodontidae | Taru | Takifugu rubripes | torafugu | 16 | NC_042300.1 (1809679..1815008, complement) | tlr5 | XM_011611738.2 | XP_011610040.2 | tlr5 | | TLR5 |
| Actinopterygii | Teleostei | Tetraodontiformes | Tetraodontidae | Taru | Takifugu rubripes | torafugu | 1 | NC_042285.1 (18374593..18378156, complement) | tlr7 | XM_011609151.2 | XP_011607453.2 | tlr7 | | TLR7 |
| Actinopterygii | Teleostei | Tetraodontiformes | Tetraodontidae | Taru | Takifugu rubripes | torafugu | 1 | NC_042285.1 (18370495..18374479, complement) | LOC115246422 | XM_029847010.1 | XP_029702870.1 | TLR7 | | TLR8 |
| Actinopterygii | Teleostei | Tetraodontiformes | Tetraodontidae | Taru | Takifugu rubripes | torafugu | 3 | NC_042287.1(11254719..11251509, complement) | x | AC156439.1 | x | tlr7 | x | TLR9 |
| Actinopterygii | Teleostei | Tetraodontiformes | Tetraodontidae | Taru | Takifugu rubripes | torafugu | 7 | NC_042291.1 (5379626..5390401, complement) | LOC101068527 | XM_003965816.3 | XP_003965865.2 | tlr1 | TLR2 | TLR18 |
| Actinopterygii | Teleostei | Tetraodontiformes | Tetraodontidae | Taru | Takifugu rubripes | torafugu | 7 | NC_042291.1 (16653038..16659017, complement) | tlr21 | NM_001032579.1 | NP_001027751.1 | tlr11 | TLR21 | TLR21 |
| Actinopterygii | Teleostei | Tetraodontiformes | Tetraodontidae | Taru | Takifugu rubripes | torafugu | 10 | NC_042294.1 (9765148..9769486, complement) | TLR22 | NM_001113193.1 | NP_001106664.1 | TLR11 | TLR22 | TLR22 |
| Actinopterygii | Teleostei | Tetraodontiformes | Tetraodontidae | Taru | Takifugu rubripes | torafugu | 12 | NC_042296.1 (8504969..8512021) | TLR13 | XM_029844550.1 | XP_029700410.1 | TLR11 | TLR13 | TLR22 |
| Actinopterygii | Holostei | Lepisosteiformes | Lepisosteidae | Leoc | Lepisostelus oculatus | spotted gar | 1 | NC_090696.1 (67633212..67640311) | tlr1 | XM_015345878.2 | XP_015201364.2 | TLR1 | | TLR1 |
| Actinopterygii | Holostei | Lepisosteiformes | Lepisosteidae | Leoc | Lepisostelus oculatus | spotted gar | 1 | NC_090696.1 (24817621..24822920, complement) | tlr2 | XM_015344720.2 | XP_015200206.2 | TLR1 | | TLR2 |
| Actinopterygii | Holostei | Lepisosteiformes | Lepisosteidae | Leoc | Lepisostelus oculatus | spotted gar | 1 | NC_090696.1 (75846200..75852884) | tlr3 | XM_006630108.3 | XP_006630171.2 | tlr3 | | TLR3 |
| Actinopterygii | Holostei | Lepisosteiformes | Lepisosteidae | Leoc | Lepisostelus oculatus | spotted gar | 24 | NC_090719.1 (10491708..10494754, complement) | LOC102693850 | XM_069183148.1 | XP_069039249.1 | tlr4 | | TLR4 |
| Actinopterygii | Holostei | Lepisosteiformes | Lepisosteidae | Leoc | Lepisostelus oculatus | spotted gar | 2 | NC_090697.1 (42883566..42902389, complement) | LOC102693268 | XM_006625986.3 | XP_006626049.1 | tlr5 | | TLR5 |
| Actinopterygii | Holostei | Lepisosteiformes | Lepisosteidae | Leoc | Lepisostelus oculatus | spotted gar | 13 | NC_090708.1 (21823227..21827032) | LOC107079175 | XM_015361454.2 | XP_015216940.2 | tlr7 | | TLR7 |
| Actinopterygii | Holostei | Lepisosteiformes | Lepisosteidae | Leoc | Lepisostelus oculatus | spotted gar | 13 | NC_090708.1(21825933 to 21829100) | x | x | x | tlr7 | | TLR7 |
| Actinopterygii | Holostei | Lepisosteiformes | Lepisosteidae | Leoc | Lepisostelus oculatus | spotted gar | 13 | NC_090708.1 (21831536..21836748) | LOC102685072 | XM_015361455.2 | XP_015216941.2 | TLR7 | | TLR8 |
| Actinopterygii | Holostei | Lepisosteiformes | Lepisosteidae | Leoc | Lepisostelus oculatus | spotted gar | 13 | NC_090708.1 (21839044..21854679) | LOC102682393 | XM_006637902.3 | XP_006637965.2 | TLR7 | TMBX | TLR8 |
| Actinopterygii | Holostei | Lepisosteiformes | Lepisosteidae | Leoc | Lepisostelus oculatus | spotted gar | 9 | NC_090704.1 (4675430..4683968) | LOC107078350 | XM_015355233.2 | XP_015210719.2 | tlr7 | TLR8 | TLR8 |
| Actinopterygii | Holostei | Lepisosteiformes | Lepisosteidae | Leoc | Lepisostelus oculatus | spotted gar | 4 | NC_090699.1 (65776471..65781412) | LOC102686134 | XM_015348292.2 | XP_015203778.2 | TLR7 | TLR9 | TLR9 |
| Actinopterygii | Holostei | Lepisosteiformes | Lepisosteidae | Leoc | Lepisostelus oculatus | spotted gar | 4 | NC_090699.1 (64273352..64278288, complement) | | XM_015348046.2 | XP_015203532.2 | TLR7 | TLR9 | TLR9 |
| Actinopterygii | Holostei | Lepisosteiformes | Lepisosteidae | Leoc | Lepisostelus oculatus | spotted gar | 2 | NC_090697.1(64922761 to 64924290) | x | x | x | TLR11 | x | TLR13 |
| Actinopterygii | Holostei | Lepisosteiformes | Lepisosteidae | Leoc | Lepisostelus oculatus | spotted gar | 2 | NC_090697.1(64934566-64936052) | x | x | x | TLR11 | x | TLR13 |
| Actinopterygii | Holostei | Lepisosteiformes | Lepisosteidae | Leoc | Lepisostelus oculatus | spotted gar | 27 | NC_090722.1 (4786762..4814024, complement) | | XM_015336729.2 | XP_015192215.2 | tlr1 | TLR18 | TLR18 |
| Actinopterygii | Holostei | Lepisosteiformes | Lepisosteidae | Leoc | Lepisostelus oculatus | spotted gar | 11 | NC_090706.1 (39380963..39434786) | tlr22 | XM_069195281.1 | XP_069051382.1 | TLR11 | TLR22 | TLR22 |
| Actinopterygii | Holostei | Lepisosteiformes | Lepisosteidae | Leoc | Lepisostelus oculatus | spotted gar | 7 | NC_090702.1 (25307176..25315071) | LOC102689191 | XM_015352785.2 | XP_015208271.1 | TLR1 | TLR1 | TLR25 |
| Actinopterygii | Holostei | Lepisosteiformes | Lepisosteidae | Leoc | Lepisostelus oculatus | spotted gar | 7 | NC_090702.1 (11326658..11334652) | LOC102684141 | XM_015353150.2 | XP_015208636.2 | TLR1 | TLR1 | TLR25 |
| Actinopterygii | Holostei | Lepisosteiformes | Lepisosteidae | Leoc | Lepisostelus oculatus | spotted gar | 9 | NC_090704.1 (7854745..7857544) | LOC138241259 | XM_069194433.1 | XP_069050534.1 | TLR1 | TLR2 type-2 | TLR27 |
| Actinopterygii | Chondrostei | Acipenseriformes | Acipenseridae | Acru | Acipenser ruthenus | Sterlet | 2 | NC_081190.1 (111827747..111833508, complement) | TLR1 | XM_034015138.3 | XP_033871029.3 | TLR1 | | TLR1 |
| Actinopterygii | Chondrostei | Acipenseriformes | Acipenseridae | Acru | Acipenser ruthenus | Sterlet | 1 | NC_081189.1 (88035019..88037183, complement) | LOC131736893 | XM_059022529.1 | XP_058878512.1 | TLR2 | | |
| Actinopterygii | Chondrostei | Acipenseriformes | Acipenseridae | Acru | Acipenser ruthenus | Sterlet | 2 | NC_081190.1 (79946167..79950531) | LOC117408874 | XM_034014252.3 | XP_033870143.3 | TLR2 | | |
| Actinopterygii | Chondrostei | Acipenseriformes | Acipenseridae | Acru | Acipenser ruthenus | Sterlet | Unplaced scaffold | NW_026707711.1 (3824..8591) | LOC131728095 | XM_059019269.1 | XP_058875252.1 | | TLR2 | |
| Actinopterygii | Chondrostei | Acipenseriformes | Acipenseridae | Acru | Acipenser ruthenus | Sterlet | | NW_026707670.1 (4216..15016, complement) | LOC131728026 | XM_059019234.1 | XP_058875217.1 | | TLR2 | |
| Actinopterygii | Chondrostei | Acipenseriformes | Acipenseridae | Acru | Acipenser ruthenus | Sterlet | 1 | NC_081189.1 (62826033..62844992, complement) | tlr3 | XM_034035991.3 | XP_033891882.3 | tlr3 | | |
| Actinopterygii | Chondrostei | Acipenseriformes | Acipenseridae | Acru | Acipenser ruthenus | Sterlet | 15 | NC_081203.1 (13095020..13096260, complement) | LOC117422512 | XM_034927943.2 | XP_034783834.1 | tlr4 | | |
| Actinopterygii | Chondrostei | Acipenseriformes | Acipenseridae | Acru | Acipenser ruthenus | Sterlet | 15 | NC_081203.1 (13096998..13102279, complement) | LOC131697593 | x | x | tlr4 | TLR4 | |
| Actinopterygii | Chondrostei | Acipenseriformes | Acipenseridae | Acru | Acipenser ruthenus | Sterlet | 31 | NC_081219.1 (1501131..1506771) | LOC117396808 | XM_034909094.2 | XP_034764985.1 | tlr4 | TLR4 | TLR4 |
| Actinopterygii | Chondrostei | Acipenseriformes | Acipenseridae | Acru | Acipenser ruthenus | Sterlet | 5 | NC_081193.1 (45894391..45900392) | LOC117403155 | x | x | tlr5 | TLR5 | |
| Actinopterygii | Chondrostei | Acipenseriformes | Acipenseridae | Acru | Acipenser ruthenus | Sterlet | 6 | NC_081194.1 (26796455..26805945) | LOC117410752 | XM_034017534.3 | XP_033873425.3 | tlr5 | TLR5 | |
| Actinopterygii | Chondrostei | Acipenseriformes | Acipenseridae | Acru | Acipenser ruthenus | Sterlet | 8 | NC_081196.1 (41010650..41023386) | tlr7 | XM_034012390.3 | XP_033868281.1 | tlr7 | TLR7 | TLR7 |
| Actinopterygii | Chondrostei | Acipenseriformes | Acipenseridae | Acru | Acipenser ruthenus | Sterlet | 8 | NC_081196.1 (41023290..41050832) | LOC117406516 | XM_059029661.1 | XP_058885644.1 | tlr7 | TLR8 | TLR8 |
| Actinopterygii | Chondrostei | Acipenseriformes | Acipenseridae | Acru | Acipenser ruthenus | Sterlet | 9 | NC_081197.1 (47557830..47559121) | LOC117971113 | x | x | tlr7 | TLR7 | TLR7 |
| Actinopterygii | Chondrostei | Acipenseriformes | Acipenseridae | Acru | Acipenser ruthenus | Sterlet | 9 | NC_081197.1 (47568188..47573603) | LOC117405786 | XM_059030397.1 | XP_058886380.1 | tlr7 | TLR8 | TLR8 |
| Actinopterygii | Chondrostei | Acipenseriformes | Acipenseridae | Acru | Acipenser ruthenus | Sterlet | 25 | NC_081213.1 (27858997..27862646) | LOC117412109 | XM_059000511.1 | XP_058856494.1 | tlr7 | TLR9 | |
| Actinopterygii | Chondrostei | Acipenseriformes | Acipenseridae | Acru | Acipenser ruthenus | Sterlet | 25 | NC_081213.1 (26888463..26891879, complement) | LOC117412533 | XM_034020989.3 | XP_033876880.3 | tlr7 | TLR9 | |
| Actinopterygii | Chondrostei | Acipenseriformes | Acipenseridae | Acru | Acipenser ruthenus | Sterlet | 16 | NC_081204.1 (1391781..1394570, complement) | LOC131697865 | XM_058989596.1 | XP_058845579.1 | tlr7 | TLR9 | |
| Actinopterygii | Chondrostei | Acipenseriformes | Acipenseridae | Acru | Acipenser ruthenus | Sterlet | 16 | NC_081204.1 (2373121..2376603) | LOC117963561 | XM_034904158.2 | XP_034760049.2 | tlr7 | TLR9 | |
| Actinopterygii | Chondrostei | Acipenseriformes | Acipenseridae | Acru | Acipenser ruthenus | Sterlet | 12 | NC_081200.1 (35068956..35076787, complement) | si:ch211-106k21.5 | XM_034029641.3 | XP_033885532.3 | tlr7 | TLR9 | |
| Actinopterygii | Chondrostei | Acipenseriformes | Acipenseridae | Acru | Acipenser ruthenus | Sterlet | 37 | NC_081225.1 (3514079..3519578, complement) | LOC131706875 | XM_059008772.1 | XP_058864755.1 | | TLR13 | |
| Actinopterygii | Chondrostei | Acipenseriformes | Acipenseridae | Acru | Acipenser ruthenus | Sterlet | 37 | NC_081225.1 (3544136..3545332, complement) | LOC131706753 | XM_059008438.1 | XP_058864421.1 | | TLR13 | |
| Actinopterygii | Chondrostei | Acipenseriformes | Acipenseridae | Acru | Acipenser ruthenus | Sterlet | 37 | NC_081225.1 (3574978..3576846, complement) | LOC131706807 | XM_059008605.1 | XP_058864588.1 | | TLR13 | |
| Actinopterygii | Chondrostei | Acipenseriformes | Acipenseridae | Acru | Acipenser ruthenus | Sterlet | 37 | NC_081225.1 (3606798..3626597, complement) | LOC131706754 | x | x | | TLR13 | |
| Actinopterygii | Chondrostei | Acipenseriformes | Acipenseridae | Acru | Acipenser ruthenus | Sterlet | 37 | NC_081225.1 (3639128..3642753, complement) | LOC131706868 | XM_059008756.1 | XP_058864739.1 | | TLR13 | |
| Actinopterygii | Chondrostei | Acipenseriformes | Acipenseridae | Acru | Acipenser ruthenus | Sterlet | 46 | NC_081234.1 (2511159..2527720, complement) | LOC131720979 | XM_059013613.1 | XP_058869596.1 | | TLR2 | |
| Actinopterygii | Chondrostei | Acipenseriformes | Acipenseridae | Acru | Acipenser ruthenus | Sterlet | 46 | NC_081234.1 (2469316..2473376, complement) | LOC131720972 | XM_059013573.1 | XP_058869556.1 | | TLR13 | |
| Actinopterygii | Chondrostei | Acipenseriformes | Acipenseridae | Acru | Acipenser ruthenus | Sterlet | 46 | NC_081234.1 (2410297..2410951, complement) | LOC131721000 | XM_059013740.1 | XP_058869723.1 | | TLR13 | |
| Actinopterygii | Chondrostei | Acipenseriformes | Acipenseridae | Acru | Acipenser ruthenus | Sterlet | 37 | NC_081225.1 (3682803..3702251, complement) | LOC131706789 | XM_059008542.1 | XP_058864525.1 | TLR11 | TLR2 | |
| Actinopterygii | Chondrostei | Acipenseriformes | Acipenseridae | Acru | Acipenser ruthenus | Sterlet | 35 | NC_081223.1 (11003751..11015444, complement) | LOC117395620 | XM_033994631.3 | XP_033850522.3 | tlr1 | TLR2 | TLR18 |
| Actinopterygii | Chondrostei | Acipenseriformes | Acipenseridae | Acru | Acipenser ruthenus | Sterlet | 32 | NC_081220.1 (13607191..13618952, complement) | LOC117965134 | XM_034910085.2 | XP_034765976.2 | tlr1 | TLR2 | TLR18 |
| Actinopterygii | Chondrostei | Acipenseriformes | Acipenseridae | Acru | Acipenser ruthenus | Sterlet | 41 | NC_081229.1 (1524738..1530172) | LOC131709107 | XM_059011025.1 | XP_058867008.1 | TLR11 | TLR13 | TLR21 |
| Actinopterygii | Chondrostei | Acipenseriformes | Acipenseridae | Acru | Acipenser ruthenus | Sterlet | 38 | NC_081226.1 (61806..66406) | LOC131707138 | XM_059009360.1 | XP_058865343.1 | TLR11 | TLR13 | TLR21 |
| Actinopterygii | Chondrostei | Acipenseriformes | Acipenseridae | Acru | Acipenser ruthenus | Sterlet | 23 | NC_081211.1 (29238143..29247847) | LOC117412777 | x | x | TLR11 | TLR6 | TLR22 |
| Actinopterygii | Chondrostei | Acipenseriformes | Acipenseridae | Acru | Acipenser ruthenus | Sterlet | 23 | NC_081211.1 (29250610..29254298, complement) | LOC117964560 | XM_058997192.1 | XP_058853175.1 | TLR11 | TLR13 | TLR22 |
| Actinopterygii | Chondrostei | Acipenseriformes | Acipenseridae | Acru | Acipenser ruthenus | Sterlet | 22 | NC_081210.1 (30415371..30423891) | LOC131699494 | XM_058996508.1 | XP_058852491.1 | TLR11 | TLR13 | TLR22 |
| Actinopterygii | Chondrostei | Acipenseriformes | Acipenseridae | Acru | Acipenser ruthenus | Sterlet | 22 | NC_081210.1 (30425199..30428060, complement) | LOC117431738 | XM_058996626.1 | XP_058852609.1 | TLR11 | TLR13 | TLR22 |
| Actinopterygii | Chondrostei | Acipenseriformes | Acipenseridae | Acru | Acipenser ruthenus | Sterlet | 7 | NC_081195.1 (38100294..38110100, complement) | LOC117415230 | XM_059026903.1 | XP_058882886.1 | TLR1 | TLR1 | TLR25 |
| Actinopterygii | Chondrostei | Acipenseriformes | Acipenseridae | Acru | Acipenser ruthenus | Sterlet | 7 | NC_081195.1 (38110900..38113596) | LOC117415385 | x | x | tlr1 | TLR1 | TLR25 |
| Actinopterygii | Chondrostei | Acipenseriformes | Acipenseridae | Acru | Acipenser ruthenus | Sterlet | 14 | NC_081202.1 (31401214..31405020) | LOC117419919 | XM_034033304.2 | XP_033889195.2 | tlr1 | TLR1 | TLR25 |
| Actinopterygii | Chondrostei | Acipenseriformes | Acipenseridae | Acru | Acipenser ruthenus | Sterlet | 12 | NC_081200.1 (30137248..30149792) | LOC117416631 | XM_034027898.3 | XP_033883789.1 | TLR1 | TLR2 type-2 | TLR27 |
| Actinopterygii | Chondrostei | Acipenseriformes | Acipenseridae | Acru | Acipenser ruthenus | Sterlet | 10 | NC_081198.1 (11726416..11728513, complement) | LOC131738105 | x | x | TLR1 | TLR2 | TLR27 |
| Actinopterygii | Chondrostei | Acipenseriformes | Acipenseridae | Acru | Acipenser ruthenus | Sterlet | 23 | NC_081211.1 (18320867..18326525, complement) | LOC131699621 | XM_058996749.1 | XP_058852732.1 | TLR1 | TLR2 | TLR27 |
| Actinopterygii | Chondrostei | Acipenseriformes | Acipenseridae | Acru | Acipenser ruthenus | Sterlet | 23 | NC_081211.1 (18307678..18314142, complement) | LOC117413139 | XM_058996748.1 | XP_058852731.1 | TLR1 | TLR2 type-2 | TLR27 |
| Actinopterygii | Chondrostei | Acipenseriformes | Acipenseridae | Acru | Acipenser ruthenus | Sterlet | 23 | NC_081211.1(18308357 to 18310180) | x | x | x | tlr1 | x | TLR27 |
| Actinopterygii | Chondrostei | Acipenseriformes | Acipenseridae | Acru | Acipenser ruthenus | Sterlet | 22 | NC_081210.1 (19457849..19465165, complement) | LOC131699437 | x | x | TLR1 | TLR2 type-2 | TLR27 |
| Actinopterygii | Chondrostei | Acipenseriformes | Acipenseridae | Acru | Acipenser ruthenus | Sterlet | 10 | NC_081198.1 (11164693..11167367) | LOC131738104 | XM_059031359.1 | XP_058887342.1 | | TLR6 | TLR27 |
| Actinopterygii | Cladistia | Polypteriformes | Polypteridae | Erca | Erpetoichthys calabaricus | reedfish | 5 | NC_041398.2 (114325267..114339946, complement) | LOC114651904 | XM_028801980.2 | XP_028657813.2 | TLR1 | TLR6 | |

| | | | | | | | | | | | | | | |
|---|---|---|---|---|---|---|---|---|---|---|---|---|---|---|
| Actinopterygii | Cladistia | Polypteriformes | Polypteridae | Erca | Erpetoichthys calabaricus | reedfish | 5 | NC_041398.2 (114313493..114317632, complement) | tlr1 | XM_051928200.1 | XP_051784160.1 | TLR1 | TLR1 | |
| Actinopterygii | Cladistia | Polypteriformes | Polypteridae | Erca | Erpetoichthys calabaricus | reedfish | 11 | NC_041404.2 (46162166..46211731, complement) | LOC114660386 | XM_051934214.1 | XP_051790174.1 | TLR2 | | |
| Actinopterygii | Cladistia | Polypteriformes | Polypteridae | Erca | Erpetoichthys calabaricus | reedfish | 5 | NC_041398.2 (141164857..141249724, complement) | tlr3 | XM_051928505.1 | XP_051784465.1 | tlr3 | | |
| Actinopterygii | Cladistia | Polypteriformes | Polypteridae | Erca | Erpetoichthys calabaricus | reedfish | 1 | NC_041394.2 (100031992..100037546) | LOC114653759 | XM_028804260.2 | XP_028660093.1 | tlr5 | TLR5 | |
| Actinopterygii | Cladistia | Polypteriformes | Polypteridae | Erca | Erpetoichthys calabaricus | reedfish | 15 | NC_041408.2 (3886408..3903432, complement) | tlr5a | XM_028820179.2 | XP_028676012.2 | tlr5 | TLR5a | |
| Actinopterygii | Cladistia | Polypteriformes | Polypteridae | Erca | Erpetoichthys calabaricus | reedfish | 4 | NC_041397.2 (276211440..276232710) | LOC114650905 | XM_028800827.2 | XP_028656660.1 | tlr7 | TLR7 | TLR7 |
| Actinopterygii | Cladistia | Polypteriformes | Polypteridae | Erca | Erpetoichthys calabaricus | reedfish | 4 | NC_041397.2 (276272184..276285486) | LOC114650906 | XM_028800828.2 | XP_028656661.1 | tlr7 | TLR8 | TLR8 |
| Actinopterygii | Cladistia | Polypteriformes | Polypteridae | Erca | Erpetoichthys calabaricus | reedfish | 4 | NC_041397.2 (276312181..276315369) | LOC114650907 | XM_028800830.2 | XP_028656663.2 | tlr7 | TLR8 | TLR8 |
| Actinopterygii | Cladistia | Polypteriformes | Polypteridae | Erca | Erpetoichthys calabaricus | reedfish | 18 | NC_041411.2 (29272654..29293565) | tlr9 | XM_028824602.2 | XP_028680435.1 | tlr7 | TLR9 | TLR9 |
| Actinopterygii | Cladistia | Polypteriformes | Polypteridae | Erca | Erpetoichthys calabaricus | reedfish | 18 | NC_041411.2 (26892595..26901311) | LOC114669108 | XM_051921395.1 | XP_051777355.1 | tlr7 | TLR9 | |
| Actinopterygii | Cladistia | Polypteriformes | Polypteridae | Erca | Erpetoichthys calabaricus | reedfish | 2 | NC_041395.2 (81702203..81802830) | tlr18 | XM_028793972.2 | XP_028649805.2 | tlr1 | TLR18 | TLR18 |
| Actinopterygii | Cladistia | Polypteriformes | Polypteridae | Erca | Erpetoichthys calabaricus | reedfish | 17 | NC_041410.2 (31387575..31397218, complement) | LOC114667918 | XM_028823439.2 | XP_028679272.2 | tlr11 | TLR21 | TLR21 |
| Actinopterygii | Cladistia | Polypteriformes | Polypteridae | Erca | Erpetoichthys calabaricus | reedfish | 11 | NC_041404.2 (60378350..60427506, complement) | LOC114660449 | XM_028813148.2 | XP_028668981.2 | TLR11 | TLR13 | TLR22 |
| Actinopterygii | Cladistia | Polypteriformes | Polypteridae | Erca | Erpetoichthys calabaricus | reedfish | 11 | NC_041404.2 (60329315..60347053, complement) | LOC114642289 | XM_028791245.2 | XP_028647078.2 | TLR11 | TLR13 | TLR22 |
| Actinopterygii | Cladistia | Polypteriformes | Polypteridae | Erca | Erpetoichthys calabaricus | reedfish | 1 | NC_041394.2(181009489 to 181011957, complement) | x | x | x | tlr1 | x | TLR25 |
| Actinopterygii | Cladistia | Polypteriformes | Polypteridae | Erca | Erpetoichthys calabaricus | reedfish | 1 | NC_041394.2(181054460 to 181056925, complement) | x | x | x | tlr1 | x | TLR25 |
| Actinopterygii | Cladistia | Polypteriformes | Polypteridae | Erca | Erpetoichthys calabaricus | reedfish | 10 | NC_041403.2 (66792953..66795091) | LOC127529318 | XM_051932433.1 | XP_051788393.1 | TLR1 | TLR2 type-2 | TLR27 |
| Actinopterygii | Cladistia | Polypteriformes | Polypteridae | Erca | Erpetoichthys calabaricus | reedfish | 10 | NC_041403.2 (66812575..66828880) | LOC114658383 | XM_051932681.1 | XP_051788641.1 | TLR1 | TLR2 | TLR27 |
| Actinopterygii | Cladistia | Polypteriformes | Polypteridae | Erca | Erpetoichthys calabaricus | reedfish | 12 | NC_041405.2 (10201884..10209475) | LOC114661827 | XM_028815033.2 | XP_028670866.1 | TLR11 | TLR13 | TLR29 |
| Chondrichthyes | Elasmobranchii | Carcharhiniformes | Carcharhinidae | Calo | Carcharhinus longimanus | oceanic whitetip shark | JASCQW010000001.1 | JASCQW010000001.1(82286073..82288544) | x | x | x | TLR1 | x | |
| Chondrichthyes | Elasmobranchii | Carcharhiniformes | Carcharhinidae | Calo | Carcharhinus longimanus | oceanic whitetip shark | JASCQW010008010.1 | JASCQW010008010.1(27-end) | x | x | x | TLR1 | x | |
| Chondrichthyes | Elasmobranchii | Carcharhiniformes | Carcharhinidae | Calo | Carcharhinus longimanus | oceanic whitetip shark | JASCQW010000043.1 | JASCQW010000043.1(24301400 to 24314263) | x | x | x | tlr3 | x | |
| Chondrichthyes | Elasmobranchii | Carcharhiniformes | Carcharhinidae | Calo | Carcharhinus longimanus | oceanic whitetip shark | JASCQW010000109.1 | JASCQW010000109.1(48465276 to 48467933) | x | x | x | tlr5 | x | |
| Chondrichthyes | Elasmobranchii | Carcharhiniformes | Carcharhinidae | Calo | Carcharhinus longimanus | oceanic whitetip shark | JASCQW010000105.1 | JASCQW010000105.1(58671174 to 58674341) | x | x | x | TLR7 | x | |
| Chondrichthyes | Elasmobranchii | Carcharhiniformes | Carcharhinidae | Calo | Carcharhinus longimanus | oceanic whitetip shark | JASCQW010000105.1 | JASCQW010000105.1(58724381 to 58727476) | x | x | x | TLR7 | x | |
| Chondrichthyes | Elasmobranchii | Carcharhiniformes | Carcharhinidae | Calo | Carcharhinus longimanus | oceanic whitetip shark | JASCQW010000162.1 | JASCQW010000162.1(126779212 to 126782355) | x | x | x | TLR7 | x | TLR9 |
| Chondrichthyes | Elasmobranchii | Carcharhiniformes | Carcharhinidae | Calo | Carcharhinus longimanus | oceanic whitetip shark | JASCQW010000190.1 | JASCQW010000190.1( 130524969 to 130527808) | x | x | x | TLR11 | x | TLR13 |
| Chondrichthyes | Elasmobranchii | Carcharhiniformes | Carcharhinidae | Calo | Carcharhinus longimanus | oceanic whitetip shark | JASCQW010065522.1 | JASCQW010065522.1(0-523) | x | x | x | TLR1 | x | TLR18 |
| Chondrichthyes | Elasmobranchii | Carcharhiniformes | Carcharhinidae | Calo | Carcharhinus longimanus | oceanic whitetip shark | JASCQW010000109.1 | JASCQW010000109.1(31722188 to 31725046) | x | x | x | TLR11 | x | TLR22 |
| Chondrichthyes | Elasmobranchii | Carcharhiniformes | Carcharhinidae | Calo | Carcharhinus longimanus | oceanic whitetip shark | JASCQW010000081.1 | JASCQW010000081.1(53471586 to 53474059) | x | x | x | TLR1 | x | TLR27 |
| Chondrichthyes | Elasmobranchii | Carcharhiniformes | Carcharhinidae | Calo | Carcharhinus longimanus | oceanic whitetip shark | JASCQW010000175.1 | JASCQW010000175.1(12185053 to 12188788) | x | x | x | TLR11 | x | TLR29 |
| Chondrichthyes | Elasmobranchii | Carcharhiniformes | Carcharhinidae | Prgl | Prionace glauca | blue shark | 1 | CM075721.1(102054283 to 102056754) | x | x | x | TLR1 | x | TLR1 |
| Chondrichthyes | Elasmobranchii | Carcharhiniformes | Carcharhinidae | Prgl | Prionace glauca | blue shark | 1 | CM075721.1(146102480 to 146104846) | x | x | x | TLR1 | x | TLR2A |
| Chondrichthyes | Elasmobranchii | Carcharhiniformes | Carcharhinidae | Prgl | Prionace glauca | blue shark | 1 | CM075721.1(146141482 to 146143878) | x | x | x | TLR1 | x | TLR2B |
| Chondrichthyes | Elasmobranchii | Carcharhiniformes | Carcharhinidae | Prgl | Prionace glauca | blue shark | 6 | CM075726.1(30900000 to 30913082) | x | x | x | tlr3 | x | TLR3 |
| Chondrichthyes | Elasmobranchii | Carcharhiniformes | Carcharhinidae | Prgl | Prionace glauca | blue shark | 5 | CM075725.1(12134066 to 12136723) | x | x | x | tlr5 | x | TLR5 |
| Chondrichthyes | Elasmobranchii | Carcharhiniformes | Carcharhinidae | Prgl | Prionace glauca | blue shark | 13 | CM075733.1(66549015 to 66552182) | x | x | x | tlr7 | x | TLR7 |
| Chondrichthyes | Elasmobranchii | Carcharhiniformes | Carcharhinidae | Prgl | Prionace glauca | blue shark | 13 | CM075733.1(66546989 to 66547973) | x | x | x | tlr7 | x | |
| Chondrichthyes | Elasmobranchii | Carcharhiniformes | Carcharhinidae | Prgl | Prionace glauca | blue shark | 13 | CM075733.1(66617377 to 66620472) | x | x | x | tlr7 | x | TLR8 |
| Chondrichthyes | Elasmobranchii | Carcharhiniformes | Carcharhinidae | Prgl | Prionace glauca | blue shark | 13 | CM075733.1(66627292 to 66629999) | x | x | x | tlr7 | x | |
| Chondrichthyes | Elasmobranchii | Carcharhiniformes | Carcharhinidae | Prgl | Prionace glauca | blue shark | 3 | CM075723.1(146347525 to 146350668) | x | x | x | tlr7 | x | TLR9 |
| Chondrichthyes | Elasmobranchii | Carcharhiniformes | Carcharhinidae | Prgl | Prionace glauca | blue shark | 4 | CM075724.1(153270512 to 153272017) | | | | TLR11 | x | TLR13 |
| Chondrichthyes | Elasmobranchii | Carcharhiniformes | Carcharhinidae | Prgl | Prionace glauca | blue shark | 14 | CM075734.1(39990621 to 39993569) | x | x | x | TLR11 | x | TLR22 |
| Chondrichthyes | Elasmobranchii | Carcharhiniformes | Carcharhinidae | Prgl | Prionace glauca | blue shark | 34 | CM075754.1(19861782 to 19864670) | x | x | x | TLR11 | x | TLR29 |
| Chondrichthyes | Elasmobranchii | Carcharhiniformes | Scyliorhinidae | Scca | Scyliorhinus canicula | catshark | 3 | NC_052148.1 (135767018..135804752) | LOC119962987 | XM_038791409.1 | XP_038647337.1 | TLR1 | TLR1 | |
| Chondrichthyes | Elasmobranchii | Carcharhiniformes | Scyliorhinidae | Scca | Scyliorhinus canicula | catshark | 3 | NC_052148.1 (203309975..203341850, complement) | LOC119963231 | XM_038792028.1 | XP_038647956.1 | TLR1 | TLR2 | |
| Chondrichthyes | Elasmobranchii | Carcharhiniformes | Scyliorhinidae | Scca | Scyliorhinus canicula | catshark | 3 | NC_052148.1 (203400747..203480559) | LOC119963232 | XM_038792032.1 | XP_038647960.1 | TLR1 | TLR2 | |
| Chondrichthyes | Elasmobranchii | Carcharhiniformes | Scyliorhinidae | Scca | Scyliorhinus canicula | catshark | 8 | NC_052153.1 (164657593..164692002, complement) | tlr3 | XM_038805636.1 | XP_038661564.1 | tlr3 | TLR3 | |
| Chondrichthyes | Elasmobranchii | Carcharhiniformes | Scyliorhinidae | Scca | Scyliorhinus canicula | catshark | 6 | NC_052151.1 (12662283..12676168) | LOC119966991 | XM_038798967.1 | XP_038654895.1 | tlr5 | TLR5 | |
| Chondrichthyes | Elasmobranchii | Carcharhiniformes | Scyliorhinidae | Scca | Scyliorhinus canicula | catshark | 7 | NC_052152.1 (92186419..92236835) | LOC119969192 | XM_038802452.1 | XP_038658380.1 | TLR7 | TLR7 | |
| Chondrichthyes | Elasmobranchii | Carcharhiniformes | Scyliorhinidae | Scca | Scyliorhinus canicula | catshark | 7 | NC_052152.1 (92291736..92398310) | LOC119969193 | XM_038802453.1 | XP_038658381.1 | TLR7 | TLR8 | |
| Chondrichthyes | Elasmobranchii | Carcharhiniformes | Scyliorhinidae | Scca | Scyliorhinus canicula | catshark | 11 | NC_052156.1 (13583298..13599197) | TLR9 | XM_038812296.1 | XP_038668224.1 | TLR7 | TLR9 | |
| Chondrichthyes | Elasmobranchii | Carcharhiniformes | Scyliorhinidae | Scca | Scyliorhinus canicula | catshark | 17 | NC_052162.1 (30646334..30656388, complement) | LOC119952319 | XM_038775972.1 | XP_038631900.1 | TLR11 | TLR13 | TLR13 |
| Chondrichthyes | Elasmobranchii | Carcharhiniformes | Scyliorhinidae | Scca | Scyliorhinus canicula | catshark | 30 | NC_052175.1 (10676270..10684340, complement) | tlr21 | XM_038787067.1 | XP_038642995.1 | TLR11 | TLR21 | TLR21 |
| Chondrichthyes | Elasmobranchii | Carcharhiniformes | Scyliorhinidae | Scca | Scyliorhinus canicula | catshark | 4 | NC_052149.1 (174838734..174842563) | tlr22 | XM_038795574.1 | XP_038651502.1 | TLR11 | TLR22 | TLR22 |
| Chondrichthyes | Elasmobranchii | Carcharhiniformes | Scyliorhinidae | Scca | Scyliorhinus canicula | catshark | 4 | NC_052149.1 (83069616..83084292, complement) | LOC119964722 | XM_038794620.1 | XP_038650548.1 | TLR1 | TLR2 | TLR27 |
| Chondrichthyes | Elasmobranchii | Carcharhiniformes | Scyliorhinidae | Scca | Scyliorhinus canicula | catshark | 23 | NC_052168.1 (2627510..2631735) | LOC119956591 | XM_038783918.1 | XP_038639846.1 | TLR11 | TLR13 | TLR29 |
| Chondrichthyes | Elasmobranchii | Carcharhiniformes | Scyliorhinidae | Scto | Scyliorhinus torazame | cloudy catshark | 3 | CM105024.1(185218709 to 185221186) | x | x | x | TLR1 | x | TLR1 |
| Chondrichthyes | Elasmobranchii | Carcharhiniformes | Scyliorhinidae | Scto | Scyliorhinus torazame | cloudy catshark | 3 | CM105024.1(97224645 - 97227047) | x | x | x | TLR1 | x | TLR2 |
| Chondrichthyes | Elasmobranchii | Carcharhiniformes | Scyliorhinidae | Scto | Scyliorhinus torazame | cloudy catshark | 3 | CM105024.1(97054032 - 97051679) | x | x | x | TLR1 | x | TLR2 |
| Chondrichthyes | Elasmobranchii | Carcharhiniformes | Scyliorhinidae | Scto | Scyliorhinus torazame | cloudy catshark | 9 | CM105030.1( 54932405 to 54967062) | x | x | x | tlr3 | x | TLR3 |
| Chondrichthyes | Elasmobranchii | Carcharhiniformes | Scyliorhinidae | Scto | Scyliorhinus torazame | cloudy catshark | 5 | CM105025.1( 338221601 to 338224255) | x | x | x | tlr5 | x | TLR5 |
| Chondrichthyes | Elasmobranchii | Carcharhiniformes | Scyliorhinidae | Scto | Scyliorhinus torazame | cloudy catshark | 8 | CM105029.1(118506644 to 118509796) | x | x | x | tlr7 | x | TLR7 |
| Chondrichthyes | Elasmobranchii | Carcharhiniformes | Scyliorhinidae | Scto | Scyliorhinus torazame | cloudy catshark | 8 | CM105029.1(118592505 to 118595627) | x | x | x | tlr7 | x | TLR8 |
| Chondrichthyes | Elasmobranchii | Carcharhiniformes | Scyliorhinidae | Scto | Scyliorhinus torazame | cloudy catshark | 13 | CM105034.1(206456917 to 206460063) | x | x | x | tlr7 | x | TLR9 |
| Chondrichthyes | Elasmobranchii | Carcharhiniformes | Scyliorhinidae | Scto | Scyliorhinus torazame | cloudy catshark | 26 | CM105047.1(40828883 to 40831330) | x | x | x | tlr1 | x | TLR18 |
| Chondrichthyes | Elasmobranchii | Carcharhiniformes | Scyliorhinidae | Scto | Scyliorhinus torazame | cloudy catshark | NW_027307984.1 | NW_027307984.1(109947 to 112796) | x | x | x | tlr11 | x | TLR21 |
| Chondrichthyes | Elasmobranchii | Carcharhiniformes | Scyliorhinidae | Scto | Scyliorhinus torazame | cloudy catshark | 7 | CM105028.1(233892486 to 233895359) | x | x | x | TLR11 | x | TLR22 |
| Chondrichthyes | Elasmobranchii | Carcharhiniformes | Scyliorhinidae | Scto | Scyliorhinus torazame | cloudy catshark | 7 | CM105028.1(109855529 to 109857964) | x | x | x | TLR1 | x | TLR27 |
| Chondrichthyes | Elasmobranchii | Carcharhiniformes | Scyliorhinidae | Scto | Scyliorhinus torazame | cloudy catshark | 29 | CM105050.1(37679545 to 37682427) | x | x | x | TLR11 | x | TLR29 |
| Chondrichthyes | Elasmobranchii | Carcharhiniformes | Scyliorhinidae | Scto | Scyliorhinus torazame | cloudy catshark | | | | | | | | |
| Chondrichthyes | Elasmobranchii | Carcharhiniformes | Scyliorhinidae | Scto | Scyliorhinus torazame | cloudy catshark | | | | | | | | |
| Chondrichthyes | Elasmobranchii | Carcharhiniformes | Scyliorhinidae | Scto | Scyliorhinus torazame | cloudy catshark | | | | | | | | |
| Chondrichthyes | Elasmobranchii | Carcharhiniformes | Scyliorhinidae | Scto | Scyliorhinus torazame | cloudy catshark | | | | | | | | |
| Chondrichthyes | Elasmobranchii | Carcharhiniformes | Sphyrnidae | Spmo | Sphyrna mokarran | great hammerhead | JAGIQG010000001.1 | JAGIQG010000001.1(88094091 to 88096562) | x | x | x | TLR1 | x | TLR1 |
| Chondrichthyes | Elasmobranchii | Carcharhiniformes | Sphyrnidae | Spmo | Sphyrna mokarran | great hammerhead | JAGIQG010000001.1 | JAGIQG010000001.1( 128131714 to 128134110) | x | x | x | TLR1 | x | TLR2 |
| Chondrichthyes | Elasmobranchii | Carcharhiniformes | Sphyrnidae | Spmo | Sphyrna mokarran | great hammerhead | JAGIQG010000001.1 | JAGIQG010000001.1( 128093191 to 128095557) | x | x | x | TLR1 | x | TLR2 |
| Chondrichthyes | Elasmobranchii | Carcharhiniformes | Sphyrnidae | Spmo | Sphyrna mokarran | great hammerhead | JAGIQG010000006.1 | JAGIQG010000006.1(27966700 to 27978856) | x | x | x | tlr3 | x | TLR3 |
| Chondrichthyes | Elasmobranchii | Carcharhiniformes | Sphyrnidae | Spmo | Sphyrna mokarran | great hammerhead | JAGIQG010000005.1 | JAGIQG010000005.1(10822293 to 10824941) | x | x | x | tlr5 | x | TLR5 |
| Chondrichthyes | Elasmobranchii | Carcharhiniformes | Sphyrnidae | Spmo | Sphyrna mokarran | great hammerhead | JAGIQG010000013.1 | JAGIQG010000013.1(59114306 to 59117464) | x | x | x | TLR7 | x | TLR7 |
| Chondrichthyes | Elasmobranchii | Carcharhiniformes | Sphyrnidae | Spmo | Sphyrna mokarran | great hammerhead | JAGIQG010000013.1 | JAGIQG010000013.1(59171018 to 59174098) | x | x | x | TLR7 | x | TLR8 |
| Chondrichthyes | Elasmobranchii | Carcharhiniformes | Sphyrnidae | Spmo | Sphyrna mokarran | great hammerhead | JAGIQG010000003.1 | JAGIQG010000003.1(129800577 to 129803711) | x | x | x | tlr7 | x | TLR9 |
| Chondrichthyes | Elasmobranchii | Carcharhiniformes | Sphyrnidae | Spmo | Sphyrna mokarran | great hammerhead | JAGIQG010000014.1 | JAGIQG010000014.1(38241686 to 38244544) | x | x | x | TLR11 | x | TLR22 |
| Chondrichthyes | Elasmobranchii | Carcharhiniformes | Sphyrnidae | Spmo | Sphyrna mokarran | great hammerhead | JAGIQG010000009.1 | JAGIQG010000009.1(55514262 to 55516721) | x | x | x | TLR1 | x | TLR27 |
| Chondrichthyes | Elasmobranchii | Carcharhiniformes | Sphyrnidae | Spmo | Sphyrna mokarran | great hammerhead | JAGIQG010000034.1 | JAGIQG010000034.1(14257761 to 14260604) | x | x | x | TLR11 | x | TLR29 |
| Chondrichthyes | Elasmobranchii | Carcharhiniformes | Triakidae | Muas | Mustelus asterias | starry smooth-hound | 1 | OZ172890.1(104936552 to 104939023) | x | x | x | TLR1 | x | TLR1 |
| Chondrichthyes | Elasmobranchii | Carcharhiniformes | Triakidae | Muas | Mustelus asterias | starry smooth-hound | 1 | OZ172890.1(59543483 to 59545855) | x | x | x | TLR1 | x | TLR2 |
| Chondrichthyes | Elasmobranchii | Carcharhiniformes | Triakidae | Muas | Mustelus asterias | starry smooth-hound | 1 | OZ172890.1(59482628 to 59485030) | x | x | x | TLR1 | x | TLR2 |
| Chondrichthyes | Elasmobranchii | Carcharhiniformes | Triakidae | Muas | Mustelus asterias | starry smooth-hound | 6 | OZ172895.1(117772029 to 117796976) | x | x | x | tlr3 | x | TLR3 |
| Chondrichthyes | Elasmobranchii | Carcharhiniformes | Triakidae | Muas | Mustelus asterias | starry smooth-hound | 5 | OZ172894.1(12558926 to 12561583) | x | x | x | tlr5 | x | TLR5 |
| Chondrichthyes | Elasmobranchii | Carcharhiniformes | Triakidae | Muas | Mustelus asterias | starry smooth-hound | 17 | OZ172906.1(26848455 to 26851623) | x | x | x | tlr7 | x | TLR7 |
| Chondrichthyes | Elasmobranchii | Carcharhiniformes | Triakidae | Muas | Mustelus asterias | starry smooth-hound | 17 | OZ172906.1(26780733 to 26783828) | x | x | x | tlr7 | x | TLR8 |
| Chondrichthyes | Elasmobranchii | Carcharhiniformes | Triakidae | Muas | Mustelus asterias | starry smooth-hound | 17 | OZ172906.1(26770000 to 26772690) | x | x | x | tlr7 | x | TLR8 |
| Chondrichthyes | Elasmobranchii | Carcharhiniformes | Triakidae | Muas | Mustelus asterias | starry smooth-hound | 17 | OZ172906.1(26761599 to 26764990) | x | x | x | tlr7 | x | TLR8 |
| Chondrichthyes | Elasmobranchii | Carcharhiniformes | Triakidae | Muas | Mustelus asterias | starry smooth-hound | 3 | OZ172892.1(148977561 to 148980704) | x | x | x | tlr7 | x | TLR9 |
| Chondrichthyes | Elasmobranchii | Carcharhiniformes | Triakidae | Muas | Mustelus asterias | starry smooth-hound | 21 | OZ172910.1(7634402 to 7637341) | x | x | x | TLR11 | x | TLR13 |
| Chondrichthyes | Elasmobranchii | Carcharhiniformes | Triakidae | Muas | Mustelus asterias | starry smooth-hound | | CAXOWL010002521.1(19010 to 21353) | x | x | x | TLR1 | x | TLR18 |
| Chondrichthyes | Elasmobranchii | Carcharhiniformes | Triakidae | Muas | Mustelus asterias | starry smooth-hound | 16 | OZ172905.1(49639665 to 49642523) | x | x | x | TLR11 | x | TLR22 |
| Chondrichthyes | Elasmobranchii | Carcharhiniformes | Triakidae | Muas | Mustelus asterias | starry smooth-hound | 8 | OZ172897.1(79417855 to 79420323) | x | x | x | TLR1 | x | TLR27 |
| Chondrichthyes | Elasmobranchii | Carcharhiniformes | Triakidae | Muas | Mustelus asterias | starry smooth-hound | 4 | OZ172893.1(150056013 to 150058859) | x | x | x | TLR11 | x | TLR29 |
| Chondrichthyes | Elasmobranchii | Lamniformes | Lamnidae | Caca | Carcharodon carcharias | white shark | 1 | NC_054467.1 (137198148..137217514, complement) | LOC121276509 | XM_041185014.1 | XP_041040948.1 | TLR1 | | TLR1 |
| Chondrichthyes | Elasmobranchii | Lamniformes | Lamnidae | Caca | Carcharodon carcharias | white shark | 1 | NC_054467.1 (71562210..71568482) | LOC121279250 | XM_041190335.1 | XP_041046269.1 | TLR1 | | TLR2 |
| Chondrichthyes | Elasmobranchii | Lamniformes | Lamnidae | Caca | Carcharodon carcharias | white shark | 1 | NC_054467.1 (71637114..71646334) | LOC121283582 | XM_041198334.1 | XP_041054268.1 | TLR1 | | TLR2 |
| Chondrichthyes | Elasmobranchii | Lamniformes | Lamnidae | Caca | Carcharodon carcharias | white shark | 4 | NC_054470.1 (166673395..166708115, complement) | tlr3 | XM_041186292.1 | XP_041042226.1 | tlr3 | | TLR3 |
| Chondrichthyes | Elasmobranchii | Lamniformes | Lamnidae | Caca | Carcharodon carcharias | white shark | 5 | NC_054471.1 (22817794..22903855) | LOC121277609 | XM_041187153.1 | XP_041043087.1 | tlr5 | | TLR5 |
| Chondrichthyes | Elasmobranchii | Lamniformes | Lamnidae | Caca | Carcharodon carcharias | white shark | 5 | NC_054471.1 (21346624 to 21347661) | x | x | x | tlr5 | | |
| Chondrichthyes | Elasmobranchii | Lamniformes | Lamnidae | Caca | Carcharodon carcharias | white shark | 18 | NC_054484.1 (109949888..109969553, complement) | LOC121290929 | XM_041211980.1 | XP_041067914.1 | TLR7 | | |
| Chondrichthyes | Elasmobranchii | Lamniformes | Lamnidae | Caca | Carcharodon carcharias | white shark | 18 | NC_054484.1 (109789534..109794383, complement) | LOC121290500 | XM_041210938.1 | XP_041066872.1 | TLR7 | | |
| Chondrichthyes | Elasmobranchii | Lamniformes | Lamnidae | Caca | Carcharodon carcharias | white shark | 7 | NC_054473.1 (16045285..16052920) | LOC121279600 | XM_041190751.1 | XP_041046685.1 | tlr7 | TLR7 | TLR9 |
| Chondrichthyes | Elasmobranchii | Lamniformes | Lamnidae | Caca | Carcharodon carcharias | white shark | 9 | NC_054475.1 (143717843..143720699) | LOC121282183 | XM_041195743.1 | XP_041051677.1 | tlr11 | TLR13 | TLR13 |
| Chondrichthyes | Elasmobranchii | Lamniformes | Lamnidae | Caca | Carcharodon carcharias | white shark | 36 | NW_024470757.1 (294917..298196) | LOC121274470 | XM_041181809.1 | XP_041037743.1 | tlr11 | TLR13 | TLR21 |
| Chondrichthyes | Elasmobranchii | Lamniformes | Lamnidae | Caca | Carcharodon carcharias | white shark | 8 | NC_054474.1 (63337867..63340867, complement) | tlr22 | XM_041193110.1 | XP_041049044.1 | TLR11 | TLR22 | TLR22 |

| | | | | | | | | | | | | | | |
|---|---|---|---|---|---|---|---|---|---|---|---|---|---|---|
| Chondrichthyes | Elasmobranchii | Lamniformes | Lamnidae | Caca | Carcharodon carcharias | white shark | 16 | NC_054482.1 (45704120 to 45706588) | x | x | x | TLR1 | x | TLR27 |
| Chondrichthyes | Elasmobranchii | Lamniformes | Lamnidae | Caca | Carcharodon carcharias | white shark | 31 | NC_054497.1 (15071367..15075141, complement) | LOC121271756 | XM_041177992.1 | XP_041033926.1 | TLR11 | x | TLR29 |
| Chondrichthyes | Elasmobranchii | Lamniformes | Lamnidae | Caca | Carcharodon carcharias | white shark | 19 | NC_054485.1 (111810514..111813273, complement) | LOC121291350 | XM_041212419.1 | XP_041068353.1 | TLR11 | TLR13 | TLR30 |
| Chondrichthyes | Elasmobranchii | Lamniformes | Lamnidae | Isox | Isurus oxyrinchus | shortfin mako shark | JANJGN010000001.1 | JANJGN010000001.1(7213086 to 7215563) | x | x | x | TLR1 | x | TLR1 |
| Chondrichthyes | Elasmobranchii | Lamniformes | Lamnidae | Isox | Isurus oxyrinchus | shortfin mako shark | JANJGN010000001.1 | JANJGN010000001.1(72861169 to 72863544) | x | x | x | TLR1 | x | TLR2 |
| Chondrichthyes | Elasmobranchii | Lamniformes | Lamnidae | Isox | Isurus oxyrinchus | shortfin mako shark | JANJGN010000001.1 | JANJGN010000001.1(72946442 to 72948814) | x | x | x | TLR1 | x | TLR2 |
| Chondrichthyes | Elasmobranchii | Lamniformes | Lamnidae | Isox | Isurus oxyrinchus | shortfin mako shark | JANJGN010000004.1 | JANJGN010000004.1(165662800 to 165682761) | x | x | x | tlr3 | x | TLR3 |
| Chondrichthyes | Elasmobranchii | Lamniformes | Lamnidae | Isox | Isurus oxyrinchus | shortfin mako shark | JANJGN010000006.1 | JANJGN010000006.1(177264142 to 177266790) | x | x | x | tlr5 | x | TLR5 |
| Chondrichthyes | Elasmobranchii | Lamniformes | Lamnidae | Isox | Isurus oxyrinchus | shortfin mako shark | JANJGN010000006.1 | JANJGN010000006.1(178250980 to 178252992) | x | x | x | tlr5 | x | TLR5 |
| Chondrichthyes | Elasmobranchii | Lamniformes | Lamnidae | Isox | Isurus oxyrinchus | shortfin mako shark | JANJGN010000018.1 | JANJGN010000018.1(3954426 to 3957575) | x | x | x | TLR7 | x | TLR7 |
| Chondrichthyes | Elasmobranchii | Lamniformes | Lamnidae | Isox | Isurus oxyrinchus | shortfin mako shark | JANJGN010000018.1 | JANJGN010000018.1(3709306 to 3712434) | x | x | x | TLR7 | x | TLR8 |
| Chondrichthyes | Elasmobranchii | Lamniformes | Lamnidae | Isox | Isurus oxyrinchus | shortfin mako shark | JANJGN010000018.1 | JANJGN010000018.1(3695888 to 3696478) | x | x | x | TLR7 | x | TLR8 |
| Chondrichthyes | Elasmobranchii | Lamniformes | Lamnidae | Isox | Isurus oxyrinchus | shortfin mako shark | JANJGN010000007.1 | JANJGN010000007.1(15205049 to 15208207) | x | x | x | tlr7 | x | TLR9 |
| Chondrichthyes | Elasmobranchii | Lamniformes | Lamnidae | Isox | Isurus oxyrinchus | shortfin mako shark | JANJGN010000036.1 | JANJGN010000036.1(464854 to 467418) | x | x | x | tlr11 | x | TLR21 |
| Chondrichthyes | Elasmobranchii | Lamniformes | Lamnidae | Isox | Isurus oxyrinchus | shortfin mako shark | JANJGN010000008.1 | JANJGN010000008.1(63050683 to 63053502) | x | x | x | TLR11 | x | TLR22 |
| Chondrichthyes | Elasmobranchii | Lamniformes | Lamnidae | Isox | Isurus oxyrinchus | shortfin mako shark | JANJGN010000016.1 | JANJGN010000016.1(74475500 to 74477968) | x | x | x | TLR1 | x | TLR27 |
| Chondrichthyes | Elasmobranchii | Lamniformes | Lamnidae | Isox | Isurus oxyrinchus | shortfin mako shark | JANJGN010000030.1 | JANJGN010000030.1(16176040 to 16178928) | x | x | x | TLR11 | x | TLR29 |
| Chondrichthyes | Elasmobranchii | Lamniformes | Lamnidae | Isox | Isurus oxyrinchus | shortfin mako shark | JANJGN010000021.1 | JANJGN010000021.1(116668809-116671568) | x | x | x | TLR11 | x | TLR30 |
| Chondrichthyes | Elasmobranchii | Lamniformes | Cetorhinidae | Cema | Cetorhinus maximus | basking shark | 1 | OZ077447.1(129730974 to 129733487) | x | x | x | TLR1 | x | TLR1 |
| Chondrichthyes | Elasmobranchii | Lamniformes | Cetorhinidae | Cema | Cetorhinus maximus | basking shark | 1 | OZ077447.1(65743022 to 65745397) | x | x | x | TLR1 | x | TLR2 |
| Chondrichthyes | Elasmobranchii | Lamniformes | Cetorhinidae | Cema | Cetorhinus maximus | basking shark | 1 | OZ077447.1( 65656054 to 65658426) | x | x | x | TLR1 | x | TLR2 |
| Chondrichthyes | Elasmobranchii | Lamniformes | Cetorhinidae | Cema | Cetorhinus maximus | basking shark | 2 | OZ077448.1(32840987 to 32860900) | x | x | x | tlr3 | x | TLR3 |
| Chondrichthyes | Elasmobranchii | Lamniformes | Cetorhinidae | Cema | Cetorhinus maximus | basking shark | 5 | OZ077451.1(164874928 to 164877576) | x | x | x | tlr5 | x | TLR5 |
| Chondrichthyes | Elasmobranchii | Lamniformes | Cetorhinidae | Cema | Cetorhinus maximus | basking shark | 16 | OZ077462.1(8352127 to 8355276) | x | x | x | tlr7 | x | TLR7 |
| Chondrichthyes | Elasmobranchii | Lamniformes | Cetorhinidae | Cema | Cetorhinus maximus | basking shark | 16 | OZ077462.1(8389242 to 8392370) | x | x | x | tlr7 | x | TLR8 |
| Chondrichthyes | Elasmobranchii | Lamniformes | Cetorhinidae | Cema | Cetorhinus maximus | basking shark | 16 | OZ077462.1(8352127 to 8355276) | x | x | x | tlr7 | x | TLR8 |
| Chondrichthyes | Elasmobranchii | Lamniformes | Cetorhinidae | Cema | Cetorhinus maximus | basking shark | 16 | OZ077462.1(8403900 to 8406755) | x | x | x | tlr7 | x | TLR8 |
| Chondrichthyes | Elasmobranchii | Lamniformes | Cetorhinidae | Cema | Cetorhinus maximus | basking shark | 6 | OZ077452.1(11274750 to 11277911) | x | x | x | tlr7 | x | TLR9 |
| Chondrichthyes | Elasmobranchii | Lamniformes | Cetorhinidae | Cema | Cetorhinus maximus | basking shark | 8 | OZ077454.1( 119849035 to 119851872) | x | x | x | tlr11 | x | TLR13 |
| Chondrichthyes | Elasmobranchii | Lamniformes | Cetorhinidae | Cema | Cetorhinus maximus | basking shark | 36 | OZ077483.1(1649541 to 1652102) | x | x | x | tlr11 | x | TLR21 |
| Chondrichthyes | Elasmobranchii | Lamniformes | Cetorhinidae | Cema | Cetorhinus maximus | basking shark | 10 | OZ077456.1(84748698 to 84751622) | x | x | x | TLR11 | x | TLR22 |
| Chondrichthyes | Elasmobranchii | Lamniformes | Cetorhinidae | Cema | Cetorhinus maximus | basking shark | 17 | OZ077463.1(68787569 to 68790236) | x | x | x | TLR1 | x | TLR27 |
| Chondrichthyes | Elasmobranchii | Lamniformes | Cetorhinidae | Cema | Cetorhinus maximus | basking shark | 8 | OZ077454.1(119848975 to 119851872) | x | x | x | TLR11 | x | TLR29 |
| Chondrichthyes | Elasmobranchii | Squaliformes | Squalidae | Sqac | Squalus acanthias | Spiny dogfish | 2 | CM059426.1(96351923 to 96354307) | x | x | x | TLR1 | x | TLR1 |
| Chondrichthyes | Elasmobranchii | Squaliformes | Squalidae | Sqac | Squalus acanthias | Spiny dogfish | 2 | CM059426.1(150280000-150284000, complement) | x | x | x | TLR1 | x | TLR2 |
| Chondrichthyes | Elasmobranchii | Squaliformes | Squalidae | Sqac | Squalus acanthias | Spiny dogfish | 2 | CM059426.1(150323950 to 150326375, ) | x | x | x | TLR1 | x | TLR2 |
| Chondrichthyes | Elasmobranchii | Squaliformes | Squalidae | Sqac | Squalus acanthias | Spiny dogfish | 2 | CM059426.1(150400000 to 150403319) | x | x | x | TLR1 | x | TLR2 |
| Chondrichthyes | Elasmobranchii | Squaliformes | Squalidae | Sqac | Squalus acanthias | Spiny dogfish | 1 | CM059425.1(237565181 to 237577881, complement) | x | x | x | tlr3 | x | TLR3 |
| Chondrichthyes | Elasmobranchii | Squaliformes | Squalidae | Sqac | Squalus acanthias | Spiny dogfish | 9 | CM059433.1(134678239 to 134680899) | x | x | x | tlr5 | x | TLR5 |
| Chondrichthyes | Elasmobranchii | Squaliformes | Squalidae | Sqac | Squalus acanthias | Spiny dogfish | 9 | CM059433.1(135461065-135465500) | x | x | x | TLR5 | x | TLR5 |
| Chondrichthyes | Elasmobranchii | Squaliformes | Squalidae | Sqac | Squalus acanthias | Spiny dogfish | 5 | CM059429.1(99397283 to 99400423) | x | x | x | TLR7 | x | TLR7 |
| Chondrichthyes | Elasmobranchii | Squaliformes | Squalidae | Sqac | Squalus acanthias | Spiny dogfish | 5 | CM059429.1( 99439353 to 99442493) | x | x | x | TLR7 | x | TLR8 |
| Chondrichthyes | Elasmobranchii | Squaliformes | Squalidae | Sqac | Squalus acanthias | Spiny dogfish | 5 | CM059429.1(99479749 to 99482817) | x | x | x | TLR7 | x | TLR8 |
| Chondrichthyes | Elasmobranchii | Squaliformes | Squalidae | Sqac | Squalus acanthias | Spiny dogfish | 20 | CM059444.1(57013429 to 57016572) | x | x | x | TLR7 | x | TLR9 |
| Chondrichthyes | Elasmobranchii | Squaliformes | Squalidae | Sqac | Squalus acanthias | Spiny dogfish | 20 | CM059444.1(57031396 to 57034470) | x | x | x | TLR7 | x | TLR9 |
| Chondrichthyes | Elasmobranchii | Squaliformes | Squalidae | Sqac | Squalus acanthias | Spiny dogfish | 20 | CM059444.1(57049142 to 57052285) | x | x | x | TLR7 | x | TLR9 |
| Chondrichthyes | Elasmobranchii | Squaliformes | Squalidae | Sqac | Squalus acanthias | Spiny dogfish | 20 | CM059444.1(57067061 to 57070204) | x | x | x | TLR7 | x | TLR9 |
| Chondrichthyes | Elasmobranchii | Squaliformes | Squalidae | Sqac | Squalus acanthias | Spiny dogfish | 6 | CM059430.1(153185041-153187878) | x | x | x | TLR11 | x | TLR13 |
| Chondrichthyes | Elasmobranchii | Squaliformes | Squalidae | Sqac | Squalus acanthias | Spiny dogfish | JASTWF010000070.1 | JASTWF010000070.1 (1602101 to 1604569) | x | x | x | tlr1 | x | TLR18 |
| Chondrichthyes | Elasmobranchii | Squaliformes | Squalidae | Sqac | Squalus acanthias | Spiny dogfish | JASTWF010000032.1 | JASTWF010000032.1( 911329 to 913620) | x | x | x | TLR11 | x | TLR21 |
| Chondrichthyes | Elasmobranchii | Squaliformes | Squalidae | Sqac | Squalus acanthias | Spiny dogfish | JASTWF010000032.1 | JASTWF010000032.1(3950574 to 3952874) | x | x | x | TLR11 | x | TLR21 |
| Chondrichthyes | Elasmobranchii | Squaliformes | Squalidae | Sqac | Squalus acanthias | Spiny dogfish | JASTWF010000149.1 | JASTWF010000149.1(48101 to 50663) | x | x | x | TLR11 | x | TLR21 |
| Chondrichthyes | Elasmobranchii | Squaliformes | Squalidae | Sqac | Squalus acanthias | Spiny dogfish | JASTWF010000149.1 | JASTWF010000149.1(155017 to 157576) | x | x | x | TLR11 | x | TLR21 |
| Chondrichthyes | Elasmobranchii | Squaliformes | Squalidae | Sqac | Squalus acanthias | Spiny dogfish | JASTWF010000149.1 | JASTWF010000149.1(725712 to 728274) | x | x | x | TLR11 | x | TLR21 |
| Chondrichthyes | Elasmobranchii | Squaliformes | Squalidae | Sqac | Squalus acanthias | Spiny dogfish | JASTWF010000149.1 | JASTWF010000149.1( 930045 to 932607) | x | x | x | TLR11 | x | TLR21 |
| Chondrichthyes | Elasmobranchii | Squaliformes | Squalidae | Sqac | Squalus acanthias | Spiny dogfish | JASTWF010000521.1 | JASTWF010000521.1(34946-36694) | x | x | x | TLR11 | x | TLR21 |
| Chondrichthyes | Elasmobranchii | Squaliformes | Squalidae | Sqac | Squalus acanthias | Spiny dogfish | JASTWF010000521.1 | JASTWF010000521.1(155670-158235) | x | x | x | TLR11 | x | TLR21 |
| Chondrichthyes | Elasmobranchii | Squaliformes | Squalidae | Sqac | Squalus acanthias | Spiny dogfish | JASTWF010000521.1 | JASTWF010000521.1(232119 to 234524) | x | x | x | TLR11 | x | TLR21 |
| Chondrichthyes | Elasmobranchii | Squaliformes | Squalidae | Sqac | Squalus acanthias | Spiny dogfish | 3 | CM059427.1( 142289161 to 142292034) | x | x | x | TLR11 | x | TLR22 |
| Chondrichthyes | Elasmobranchii | Squaliformes | Squalidae | Sqac | Squalus acanthias | Spiny dogfish | 15 | CM059439.1(59820390 to 59822813) | x | x | x | TLR1 | x | TLR27 |
| Chondrichthyes | Elasmobranchii | Squaliformes | Squalidae | Sqac | Squalus acanthias | Spiny dogfish | 26 | CM059450.1(2349430 to 2352323) | x | x | x | TLR11 | x | TLR29 |
| Chondrichthyes | Elasmobranchii | Squaliformes | Squalidae | Sqac | Squalus acanthias | Spiny dogfish | 25 | CM059449.1(12561774-12564516) | x | x | x | TLR11 | x | TLR30 |
| Chondrichthyes | Elasmobranchii | Squaliformes | Squalidae | Sqac | Squalus acanthias | Spiny dogfish | 25 | CM059449.1(12810986-12813754) | x | x | x | TLR11 | x | TLR30 |
| Chondrichthyes | Elasmobranchii | Squaliformes | Squalidae | Sqac | Squalus acanthias | Spiny dogfish | JASTWF010000660.1 | JASTWF010000660.1(129595-131343) | x | x | x | TLR11 | x | TLR30 |
| Chondrichthyes | Elasmobranchii | Squaliformes | Squalidae | Sqac | Squalus acanthias | Spiny dogfish | JASTWF010000660.1 | JASTWF010000660.1(234113-235861) | x | x | x | TLR11 | x | TLR30 |
| Chondrichthyes | Elasmobranchii | Squaliformes | Squalidae | Sqac | Squalus acanthias | Spiny dogfish | JASTWF010000134.1 | JASTWF010000134.1(980285-982033) | x | x | x | TLR11 | x | TLR30 |
| Chondrichthyes | Elasmobranchii | Squaliformes | Squalidae | Sqac | Squalus acanthias | Spiny dogfish | JASTWF010000416.1 | JASTWF010000416.1(100208-101956) | x | x | x | TLR11 | x | TLR30 |
| Chondrichthyes | Elasmobranchii | Squaliformes | Squalidae | Sqsu | Squalus suckleyi | Puget sound dogfish | JAOAMX010002743.1 | JAOAMX010002743.1(3816554 to 3818938) | x | x | x | TLR1 | x | TLR1 |
| Chondrichthyes | Elasmobranchii | Squaliformes | Squalidae | Sqsu | Squalus suckleyi | Puget sound dogfish | JAOAMX010024973.1 | JAOAMX010024973.1(2365424-2366821, complement) | x | x | x | TLR1 | x | TLR2 |
| Chondrichthyes | Elasmobranchii | Squaliformes | Squalidae | Sqsu | Squalus suckleyi | Puget sound dogfish | JAOAMX010024973.2 | JAOAMX010024973.1(2408840 to 2411218, complement) | x | x | x | TLR1 | x | TLR2 |
| Chondrichthyes | Elasmobranchii | Squaliformes | Squalidae | Sqsu | Squalus suckleyi | Puget sound dogfish | JAOAMX010024973.3 | JAOAMX010024973.1(2480200 to 2482654, complement) | x | x | x | TLR1 | x | TLR2 |
| Chondrichthyes | Elasmobranchii | Squaliformes | Squalidae | Sqsu | Squalus suckleyi | Puget sound dogfish | JAOAMX010016585.1 | JAOAMX010016585.1(11951727 to 11964430) | x | x | x | tlr3 | x | TLR3 |
| Chondrichthyes | Elasmobranchii | Squaliformes | Squalidae | Sqsu | Squalus suckleyi | Puget sound dogfish | JAOAMX010016242.1 | JAOAMX010016242.1(18515 to 21175.) | x | x | x | tlr5 | x | TLR5 |
| Chondrichthyes | Elasmobranchii | Squaliformes | Squalidae | Sqsu | Squalus suckleyi | Puget sound dogfish | JAOAMX010076087.1 | JAOAMX010076087.1(57000-61181) | x | x | x | tlr5 | x | TLR5 |
| Chondrichthyes | Elasmobranchii | Squaliformes | Squalidae | Sqsu | Squalus suckleyi | Puget sound dogfish | JAOAMX010044851.1 | JAOAMX010044851.1(1466916 to 1470038) | x | x | x | TLR7 | x | TLR7 |
| Chondrichthyes | Elasmobranchii | Squaliformes | Squalidae | Sqsu | Squalus suckleyi | Puget sound dogfish | JAOAMX010044851.1 | JAOAMX010044851.1( 1550004 to 1553072) | x | x | x | TLR7 | x | TLR8 |
| Chondrichthyes | Elasmobranchii | Squaliformes | Squalidae | Sqsu | Squalus suckleyi | Puget sound dogfish | JAOAMX010044851.1 | JAOAMX010044851.1(1508992 to 1512132) | x | x | x | TLR7 | x | TLR8 |
| Chondrichthyes | Elasmobranchii | Squaliformes | Squalidae | Sqsu | Squalus suckleyi | Puget sound dogfish | JAOAMX010039371.1 | JAOAMX010039371.1(15205049 to 15208207) | x | x | x | TLR7 | x | TLR9 |
| Chondrichthyes | Elasmobranchii | Squaliformes | Squalidae | Sqsu | Squalus suckleyi | Puget sound dogfish | JAOAMX010033747.1 | JAOAMX010033747.1(10253 to 13396) | x | x | x | TLR7 | x | TLR9 |
| Chondrichthyes | Elasmobranchii | Squaliformes | Squalidae | Sqsu | Squalus suckleyi | Puget sound dogfish | JAOAMX010057206.1 | JAOAMX010057206.1(6524858-6527671) | x | x | x | TLR11 | x | TLR13 |
| Chondrichthyes | Elasmobranchii | Squaliformes | Squalidae | Sqsu | Squalus suckleyi | Puget sound dogfish | JAOAMX010100510.1 | JAOAMX010100510.1 (38594 to 41059) | x | x | x | tlr1 | x | TLR18 |
| Chondrichthyes | Elasmobranchii | Squaliformes | Squalidae | Sqsu | Squalus suckleyi | Puget sound dogfish | JAOAMX010103622.1 | JAOAMX010103622.1(31377 to 33941) | x | x | x | TLR11 | x | TLR21 |
| Chondrichthyes | Elasmobranchii | Squaliformes | Squalidae | Sqsu | Squalus suckleyi | Puget sound dogfish | JAOAMX010032691.1 | JAOAMX010032691.1(2655 to 4805) | x | x | x | TLR11 | x | TLR21 |
| Chondrichthyes | Elasmobranchii | Squaliformes | Squalidae | Sqsu | Squalus suckleyi | Puget sound dogfish | JAOAMX010044265.1 | JAOAMX010044265.1(10457 to 12607) | x | x | x | TLR11 | x | TLR21 |
| Chondrichthyes | Elasmobranchii | Squaliformes | Squalidae | Sqsu | Squalus suckleyi | Puget sound dogfish | JAOAMX010027056.1 | JAOAMX010027056.1(1185 to 3590) | x | x | x | TLR11 | x | TLR21 |
| Chondrichthyes | Elasmobranchii | Squaliformes | Squalidae | Sqsu | Squalus suckleyi | Puget sound dogfish | JAOAMX010049340.1 | JAOAMX010049340.1(2820 to 4958) | x | x | x | TLR11 | x | TLR21 |
| Chondrichthyes | Elasmobranchii | Squaliformes | Squalidae | Sqsu | Squalus suckleyi | Puget sound dogfish | JAOAMX010035051.1 | JAOAMX010035051.1(12782 to 15250) | x | x | x | TLR11 | x | TLR21 |
| Chondrichthyes | Elasmobranchii | Squaliformes | Squalidae | Sqsu | Squalus suckleyi | Puget sound dogfish | JAOAMX010093030.1 | JAOAMX010093030.1(37152 to 39714) | x | x | x | TLR11 | x | TLR21 |
| Chondrichthyes | Elasmobranchii | Squaliformes | Squalidae | Sqsu | Squalus suckleyi | Puget sound dogfish | JAOAMX010093030.1 | JAOAMX010093030.1(238690 to 241252) | x | x | x | TLR11 | x | TLR21 |
| Chondrichthyes | Elasmobranchii | Squaliformes | Squalidae | Sqsu | Squalus suckleyi | Puget sound dogfish | JAOAMX010093030.1 | JAOAMX010093030.1(137475 to 140033) | x | x | x | TLR11 | x | TLR21 |
| Chondrichthyes | Elasmobranchii | Squaliformes | Squalidae | Sqsu | Squalus suckleyi | Puget sound dogfish | JAOAMX010093030.1 | JAOAMX010093030.1(702510 to 705070) | x | x | x | TLR11 | x | TLR21 |
| Chondrichthyes | Elasmobranchii | Squaliformes | Squalidae | Sqsu | Squalus suckleyi | Puget sound dogfish | JAOAMX010023065.1 | JAOAMX010023065.1(18233 to 20743) | x | x | x | TLR11 | x | TLR21 |
| Chondrichthyes | Elasmobranchii | Squaliformes | Squalidae | Sqsu | Squalus suckleyi | Puget sound dogfish | JAOAMX010035975.1 | JAOAMX010035975.1(10460 to 13025) | x | x | x | TLR11 | x | TLR21 |
| Chondrichthyes | Elasmobranchii | Squaliformes | Squalidae | Sqsu | Squalus suckleyi | Puget sound dogfish | JAOAMX010065347.1 | JAOAMX010065347.1( 2847 to 5412) | x | x | x | TLR11 | x | TLR21 |
| Chondrichthyes | Elasmobranchii | Squaliformes | Squalidae | Sqsu | Squalus suckleyi | Puget sound dogfish | JAOAMX010005597.1 | JAOAMX010005597.1(32319 to 34942) | x | x | x | TLR11 | x | TLR21 |
| Chondrichthyes | Elasmobranchii | Squaliformes | Squalidae | Sqsu | Squalus suckleyi | Puget sound dogfish | JAOAMX010014480.1 | JAOAMX010014480.1(24457 to 26205) | x | x | x | TLR11 | x | TLR21 |
| Chondrichthyes | Elasmobranchii | Squaliformes | Squalidae | Sqsu | Squalus suckleyi | Puget sound dogfish | JAOAMX010095870.1 | JAOAMX010095870.1(3060569 to 3063442) | x | x | x | TLR11 | x | TLR22 |
| Chondrichthyes | Elasmobranchii | Squaliformes | Squalidae | Sqsu | Squalus suckleyi | Puget sound dogfish | JAOAMX010033174.1 | JAOAMX010033174.1(6648291 to 6650714) | x | x | x | TLR1 | x | TLR27 |
| Chondrichthyes | Elasmobranchii | Squaliformes | Squalidae | Sqsu | Squalus suckleyi | Puget sound dogfish | JAOAMX010082721.1 | JAOAMX010082721.1(18933 to 21827) | x | x | x | TLR11 | x | TLR29 |
| Chondrichthyes | Elasmobranchii | Squaliformes | Squalidae | Sqsu | Squalus suckleyi | Puget sound dogfish | JAOAMX010081315.1 | JAOAMX010081315.1(3-1076) | x | x | x | TLR11 | x | TLR30 |
| Chondrichthyes | Elasmobranchii | Pristiophoriformes | Pristiophoridae | Prja | Pristiophorus japonicus | Japanese sawshark | 2 | NC_091978.1 (176923914..176943442) | LOC139232836 | XM_070863331.1 | XP_070719432.1 | TLR1 | TLR1 | TLR1 |
| Chondrichthyes | Elasmobranchii | Pristiophoriformes | Pristiophoridae | Prja | Pristiophorus japonicus | Japanese sawshark | 2 | NC_091978.1 (278011621..278018213, complement) | LOC139230462 | XM_070862286.1 | XP_070718387.1 | TLR1 | TLR2 | TLR2 |
| Chondrichthyes | Elasmobranchii | Pristiophoriformes | Pristiophoridae | Prja | Pristiophorus japonicus | Japanese sawshark | 2 | NC_091978.1 (278285417..278324304) | LOC139246533 | XM_070871420.1 | XP_070727521.1 | TLR1 | TLR2 | TLR2 |
| Chondrichthyes | Elasmobranchii | Pristiophoriformes | Pristiophoridae | Prja | Pristiophorus japonicus | Japanese sawshark | 1 | NC_091977.1 (47274240..47302689) | LOC139263353 | XM_070879300.1 | XP_070735401.1 | TLR3 | TLR3 | TLR3 |
| Chondrichthyes | Elasmobranchii | Pristiophoriformes | Pristiophoridae | Prja | Pristiophorus japonicus | Japanese sawshark | 7 | NC_091983.1 (216594868..216627877, complement) | LOC139267453 | XM_070885798.1 | XP_070741899.1 | TLR5 | TLR5 | TLR5 |
| Chondrichthyes | Elasmobranchii | Pristiophoriformes | Pristiophoridae | Prja | Pristiophorus japonicus | Japanese sawshark | 11 | NC_091987.1 (113081061..113086206) | TLR7 | XM_070892734.1 | XP_070748835.1 | TLR7 | TLR7 | TLR7 |
| Chondrichthyes | Elasmobranchii | Pristiophoriformes | Pristiophoridae | Prja | Pristiophorus japonicus | Japanese sawshark | 11 | NC_091987.1 (113160289..113163099) | LOC139275565 | XM_070892735.1 | XP_070748836.1 | TLR7 | TLR8 | TLR8 |
| Chondrichthyes | Elasmobranchii | Pristiophoriformes | Pristiophoridae | Prja | Pristiophorus japonicus | Japanese sawshark | 11 | NC_091987.1 (113208670..113213501) | LOC139276355 | XM_070894210.1 | XP_070750311.1 | TLR7 | TLR8 | TLR8 |
| Chondrichthyes | Elasmobranchii | Pristiophoriformes | Pristiophoridae | Prja | Pristiophorus japonicus | Japanese sawshark | 12 | NC_091988.1 (62407150..62417616, complement) | tlr9 | XM_070894779.1 | XP_070750880.1 | TLR7 | TLR9 | TLR9 |
| Chondrichthyes | Elasmobranchii | Pristiophoriformes | Pristiophoridae | Prja | Pristiophorus japonicus | Japanese sawshark | 6 | NC_091982.1 (33174365..33177202) | LOC139266093 | XM_070883943.1 | XP_070740044.1 | TLR11 | TLR13 | TLR13 |
| Chondrichthyes | Elasmobranchii | Pristiophoriformes | Pristiophoridae | Prja | Pristiophorus japonicus | Japanese sawshark | 4 | NC_091980.1 (99893406..99896294) | LOC139262471 | XM_070877656.1 | XP_070733757.1 | TLR11 | TLR13 | TLR22 |
| Chondrichthyes | Elasmobranchii | Pristiophoriformes | Pristiophoridae | Prja | Pristiophorus japonicus | Japanese sawshark | 8 | NC_091984.1 (98135049..98141773, complement) | LOC139268652 | XM_070887153.1 | XP_070743254.1 | TLR1 | TLR2 | TLR27 |

| | | | | | | | | | | | | | | |
|---|---|---|---|---|---|---|---|---|---|---|---|---|---|---|
| Chondrichthyes | Elasmobranchii | Pristiophoriformes | Pristiophoridae | Prja | Pristiphorus japonicus | Japanese sawshark | 19 | NC_091995.1 (98695541..98698423) | tlr21 | XM_070862135.1 | XP_070718236.1 | TLR11 | TLR21 | TLR29 |
| Chondrichthyes | Elasmobranchii | Pristiophoriformes | Pristiophoridae | Prja | Pristiphorus japonicus | Japanese sawshark | 19 | NC_091995.1 (15220733..15221768, complement) | LOC139230129 | XM_070861973.1 | XP_070718074.1 | TLR11 | TLR13 | TLR30 |
| Chondrichthyes | Elasmobranchii | Hexanchiformes | Hexanchidae | Hepe | Heptranchias perlo | Sharpnose sevengill shark | 1 | NC_090325.1 (94041310..94061939) | TLR1 | XM_067988803.1 | XP_067844904.1 | TLR1 | | TLR1 |
| Chondrichthyes | Elasmobranchii | Hexanchiformes | Hexanchidae | Hepe | Heptranchias perlo | Sharpnose sevengill shark | 1 | NC_090325.1 (139577555..139593231, complement) | LOC137326544 | XM_067991747.1 | XP_067847848.1 | TLR1 | | TLR2 |
| Chondrichthyes | Elasmobranchii | Hexanchiformes | Hexanchidae | Hepe | Heptranchias perlo | Sharpnose sevengill shark | 1 | NC_090325.1 (139456359..139509349, complement) | LOC137326548 | XM_067991761.1 | XP_067847862.1 | TLR1 | | TLR2 |
| Chondrichthyes | Elasmobranchii | Hexanchiformes | Hexanchidae | Hepe | Heptranchias perlo | Sharpnose sevengill shark | 4 | NC_090328.1 (24645497..24671414) | tlr3 | XM_067982610.1 | XP_067838711.1 | tlr3 | | TLR3 |
| Chondrichthyes | Elasmobranchii | Hexanchiformes | Hexanchidae | Hepe | Heptranchias perlo | Sharpnose sevengill shark | 5 | NC_090329.1 (129121441..129153568) | LOC137322107 | XM_067985194.1 | XP_067841295.1 | tlr5 | | TLR5 |
| Chondrichthyes | Elasmobranchii | Hexanchiformes | Hexanchidae | Hepe | Heptranchias perlo | Sharpnose sevengill shark | 5 | NC_090329.1(129906514 to 129907833, complement) | x | x | x | TLR5 | x | TLR5 |
| Chondrichthyes | Elasmobranchii | Hexanchiformes | Hexanchidae | Hepe | Heptranchias perlo | Sharpnose sevengill shark | 11 | NC_090335.1 (24574183..24592963, complement) | TLR7 | XM_067992207.1 | XP_067848308.1 | TLR7 | | TLR7 |
| Chondrichthyes | Elasmobranchii | Hexanchiformes | Hexanchidae | Hepe | Heptranchias perlo | Sharpnose sevengill shark | 11 | NC_090335.1 (24528247..24567288, complement) | LOC137326826 | XM_067992206.1 | XP_067848307.1 | TLR7 | | TLR8 |
| Chondrichthyes | Elasmobranchii | Hexanchiformes | Hexanchidae | Hepe | Heptranchias perlo | Sharpnose sevengill shark | 11 | NC_090335.1 (24500637..24524100, complement) | LOC137326824 | x | x | TLR7 | | TLR8 |
| Chondrichthyes | Elasmobranchii | Hexanchiformes | Hexanchidae | Hepe | Heptranchias perlo | Sharpnose sevengill shark | 17 | NC_090341.1 (9703019..9707751) | tlr9 | XM_067999081.1 | XP_067855182.1 | tlr7 | TLR9 | TLR9 |
| Chondrichthyes | Elasmobranchii | Hexanchiformes | Hexanchidae | Hepe | Heptranchias perlo | Sharpnose sevengill shark | 15 | NC_090339.1 (46119763..46134124) | LOC137333058 | XM_067997153.1 | XP_067853254.1 | TLR11 | TLR13 | TLR13 |
| Chondrichthyes | Elasmobranchii | Hexanchiformes | Hexanchidae | Hepe | Heptranchias perlo | Sharpnose sevengill shark | 44 | NC_090368.1 (3288586..3328942, complement) | tlr18 | XM_067975901.1 | XP_067832002.1 | tlr1 | TLR18 | TLR18 |
| Chondrichthyes | Elasmobranchii | Hexanchiformes | Hexanchidae | Hepe | Heptranchias perlo | Sharpnose sevengill shark | Unplaced Scaffold | NW_027139237.1 (499956..543717, complement) | tlr21 | XM_067978097.1 | XP_067834198.1 | tlr11 | TLR21 | TLR21 |
| Chondrichthyes | Elasmobranchii | Hexanchiformes | Hexanchidae | Hepe | Heptranchias perlo | Sharpnose sevengill shark | 14 | NC_090338.1 (46377965..46383075, complement) | tlr22 | XM_067996393.1 | XP_067852494.1 | TLR11 | TLR22 | TLR22 |
| Chondrichthyes | Elasmobranchii | Hexanchiformes | Hexanchidae | Hepe | Heptranchias perlo | Sharpnose sevengill shark | 9 | NC_090333.1 (55484919..55488583, complement) | LOC137325027 | XM_067989729.1 | XP_067845830.1 | TLR1 | TLR2 | TLR27 |
| Chondrichthyes | Elasmobranchii | Hexanchiformes | Hexanchidae | Hepe | Heptranchias perlo | Sharpnose sevengill shark | 40 | NC_090364.1 (2141464..2145981) | LOC137305476 | XM_067974313.1 | XP_067830414.1 | TLR11 | TLR13 | TLR29 |
| Chondrichthyes | Elasmobranchii | Hexanchiformes | Hexanchidae | Hepe | Heptranchias perlo | Sharpnose sevengill shark | 23 | NC_090347.1 (9850274..9865385, complement) | LOC137341078 | XM_068003987.1 | XP_067860088.1 | TLR11 | TLR13 | TLR30 |
| Chondrichthyes | Elasmobranchii | Orectolobiformes | Rhincodontidae | Rhty | Rhincodon typus | whale shark | 1 | NC_063332.1 (85857050..85869569) | LOC109936549 | XM_048601197.1 | XP_048457154.1 | TLR1 | | TLR1 |
| Chondrichthyes | Elasmobranchii | Orectolobiformes | Rhincodontidae | Rhty | Rhincodon typus | whale shark | 1 | NC_063332.1 (127696005..127782180, complement) | LOC109912800 | XM_048597995.1 | XP_048453952.1 | TLR1 | | TLR2 |
| Chondrichthyes | Elasmobranchii | Orectolobiformes | Rhincodontidae | Rhty | Rhincodon typus | whale shark | 1 | NC_063332.1 127797715..127797280 complement) | tlr2 | x | x | tlr1 | x | TLR2 |
| Chondrichthyes | Elasmobranchii | Orectolobiformes | Rhincodontidae | Rhty | Rhincodon typus | whale shark | 3 | NC_063334.1 (103888221..103909173, complement) | LOC109919526 | XM_048593896.1 | XP_048449853.1 | tlr3 | | TLR3 |
| Chondrichthyes | Elasmobranchii | Orectolobiformes | Rhincodontidae | Rhty | Rhincodon typus | whale shark | 3 | NC_063334.1 (103551999..103554547) | LOC125481922 | XM_048593874.1 | XP_048449831.1 | tlr3 | | TLR3 |
| Chondrichthyes | Elasmobranchii | Orectolobiformes | Rhincodontidae | Rhty | Rhincodon typus | whale shark | 5 | NC_063336.1 (6566560..6608755, complement) | LOC125483092 | XM_048596643.1 | XP_048452600.1 | tlr5 | | TLR5 |
| Chondrichthyes | Elasmobranchii | Orectolobiformes | Rhincodontidae | Rhty | Rhincodon typus | whale shark | 11 | NC_063342.1 (60297234..60303913) | tlr7 | XM_020520648.2 | XP_020376237.1 | TLR7 | | TLR7 |
| Chondrichthyes | Elasmobranchii | Orectolobiformes | Rhincodontidae | Rhty | Rhincodon typus | whale shark | 11 | NC_063342.1 (60328504..60353253) | LOC102685072 | XM_020520658.2 | XP_020376247.1 | TLR7 | | TLR8 |
| Chondrichthyes | Elasmobranchii | Orectolobiformes | Rhincodontidae | Rhty | Rhincodon typus | whale shark | Unplaced Scaffold | NW_025968491.1 (2677..21290) | tlr9 | XM_020535784.2 | XM_020535784.2 | tlr7 | TLR9 | TLR9 |
| Chondrichthyes | Elasmobranchii | Orectolobiformes | Rhincodontidae | Rhty | Rhincodon typus | whale shark | 14 | NC_063345.1(23473521 to 23473771, complement) | x | x | x | TLR11 | x | TLR13 |
| Chondrichthyes | Elasmobranchii | Orectolobiformes | Rhincodontidae | Rhty | Rhincodon typus | whale shark | 46 | NC_063377.1 (5887164..5922543) | LOC109923055 | XM_048620331.1 | XP_048476288.1 | tlr11 | TLR13 | TLR21 |
| Chondrichthyes | Elasmobranchii | Orectolobiformes | Rhincodontidae | Rhty | Rhincodon typus | whale shark | 12 | NC_063343.1 (27552376..27557431, complement) | tlr22 | XM_020518663.2 | XP_020374252.2 | TLR11 | TLR22 | TLR22 |
| Chondrichthyes | Elasmobranchii | Orectolobiformes | Rhincodontidae | Rhty | Rhincodon typus | whale shark | 10 | NC_063341.1 (25419572..25427159) | LOC109928617 | XM_048602405.1 | XP_048458362.1 | TLR1 | TLR2 | TLR27 |
| Chondrichthyes | Elasmobranchii | Orectolobiformes | Rhincodontidae | Rhty | Rhincodon typus | whale shark | 37 | NC_063368.1 (19963677..19967897, complement) | LOC109934146 | XM_020532957.2 | XP_020388546.1 | TLR11 | TLR13 | TLR29 |
| Chondrichthyes | Elasmobranchii | Orectolobiformes | Hemiscylliidae | Heoc | Hemiscyllium ocellatum | epaulette shark | 1 | NC_083401.1 (106701153..106713543) | TLR1 | XM_060825664.1 | XP_060681647.1 | TLR1 | | TLR1 |
| Chondrichthyes | Elasmobranchii | Orectolobiformes | Hemiscylliidae | Heoc | Hemiscyllium ocellatum | epaulette shark | 1 | NC_083401.1 (166313309..166341556) | LOC132819313 | XM_060830797.1 | XP_060686780.1 | tlr1 | | TLR2 |
| Chondrichthyes | Elasmobranchii | Orectolobiformes | Hemiscylliidae | Heoc | Hemiscyllium ocellatum | epaulette shark | 1 | NC_083401.1 (166405652..166407946) | LOC132819394 | XM_060830914.1 | XP_060686897.1 | tlr1 | | TLR2 |
| Chondrichthyes | Elasmobranchii | Orectolobiformes | Hemiscylliidae | Heoc | Hemiscyllium ocellatum | epaulette shark | 2 | NC_083402.1 (31414771..31451164) | tlr3 | XM_060835432.1 | XP_060691415.1 | tlr3 | | TLR3 |
| Chondrichthyes | Elasmobranchii | Orectolobiformes | Hemiscylliidae | Heoc | Hemiscyllium ocellatum | epaulette shark | 3 | NC_083403.1 (11706048..11710646, complement) | tlr5b | XM_060854358.1 | XP_060710341.1 | tlr5 | | TLR5 |
| Chondrichthyes | Elasmobranchii | Orectolobiformes | Hemiscylliidae | Heoc | Hemiscyllium ocellatum | epaulette shark | 3 | CM036661.1(11728442 to 11731091.) | x | x | x | tlr5 | | TLR5 |
| Chondrichthyes | Elasmobranchii | Orectolobiformes | Hemiscylliidae | Heoc | Hemiscyllium ocellatum | epaulette shark | 3 | CM036661.1(11752538 to 11755189.) | x | x | x | tlr5 | | TLR5 |
| Chondrichthyes | Elasmobranchii | Orectolobiformes | Hemiscylliidae | Heoc | Hemiscyllium ocellatum | epaulette shark | 12 | NC_083412.1 (31204219..31215503, complement) | tlr7 | XM_060833266.1 | XP_060689249.1 | TLR7 | | TLR7 |
| Chondrichthyes | Elasmobranchii | Orectolobiformes | Hemiscylliidae | Heoc | Hemiscyllium ocellatum | epaulette shark | 12 | NC_083412.1 (31141901..31163339, complement) | LOC132821007 | XM_060833475.1 | XP_060689458.1 | TLR7 | | TLR8 |
| Chondrichthyes | Elasmobranchii | Orectolobiformes | Hemiscylliidae | Heoc | Hemiscyllium ocellatum | epaulette shark | 14 | NC_083414.1 (12870575..12895805) | tlr9 | XM_060835830.1 | XP_060691813.1 | tlr7 | TLR9 | TLR9 |
| Chondrichthyes | Elasmobranchii | Orectolobiformes | Hemiscylliidae | Heoc | Hemiscyllium ocellatum | epaulette shark | 50 | NC_083450.1 (9875630..9878986, complement) | LOC132805550 | XM_060820661.1 | XP_060676644.1 | tlr11 | TLR13 | TLR21 |
| Chondrichthyes | Elasmobranchii | Orectolobiformes | Hemiscylliidae | Heoc | Hemiscyllium ocellatum | epaulette shark | 16 | CM036674.1(34295036 to 34296737) | x | x | x | TLR11 | x | TLR22 |
| Chondrichthyes | Elasmobranchii | Orectolobiformes | Hemiscylliidae | Heoc | Hemiscyllium ocellatum | epaulette shark | 9 | NC_083409.1 (51920598..51923032) | LOC132818507 | XM_060829567.1 | XP_060685550.1 | TLR1 | TLR2 | TLR27 |
| Chondrichthyes | Elasmobranchii | Orectolobiformes | Hemiscylliidae | Heoc | Hemiscyllium ocellatum | epaulette shark | 33 | NC_083433.1 (2690622..2694648) | LOC132831174 | XM_060849185.1 | XP_060705168.1 | TLR11 | TLR13 | TLR29 |
| Chondrichthyes | Elasmobranchii | Orectolobiformes | Hemiscylliidae | Chpl | Chiloscyllium plagiosum | whitespotted bambooshark | 1 | NC_057710.1 (98200367..98247090) | TLR1 | XM_043693011.1 | XP_043548946.1 | TLR1 | TLR1 | TLR1 |
| Chondrichthyes | Elasmobranchii | Orectolobiformes | Hemiscylliidae | Chpl | Chiloscyllium plagiosum | whitespotted bambooshark | 1 | NC_057710.1 (155917373..155984972) | tlr2 | XM_043700155.1 | XP_043556090.1 | tlr1 | TLR2 | TLR2 |
| Chondrichthyes | Elasmobranchii | Orectolobiformes | Hemiscylliidae | Chpl | Chiloscyllium plagiosum | whitespotted bambooshark | 2 | NC_057711.1 (27890600..27899952) | tlr3 | XM_043702664.1 | XP_043558599.1 | tlr3 | TLR3 | TLR3 |
| Chondrichthyes | Elasmobranchii | Orectolobiformes | Hemiscylliidae | Chpl | Chiloscyllium plagiosum | whitespotted bambooshark | Unplaced Scaffold | NW_025211871.1 (1608..22513, complement) | LOC122547166 | XM_043685765.1 | XP_043541700.1 | tlr3 | TLR3 | TLR3 |
| Chondrichthyes | Elasmobranchii | Orectolobiformes | Hemiscylliidae | Chpl | Chiloscyllium plagiosum | whitespotted bambooshark | Unplaced Scaffold | NW_025213900.1 (14166..22189) | LOC122547751 | XM_043686334.1 | XP_043542269.1 | tlr5 | TLR5 | TLR5 |
| Chondrichthyes | Elasmobranchii | Orectolobiformes | Hemiscylliidae | Chpl | Chiloscyllium plagiosum | whitespotted bambooshark | Unplaced Scaffold | NW_025210933.1 (2285..11281, complement) | LOC122546976 | XM_043685578.1 | XP_043541513.1 | tlr5 | TLR5 | TLR5 |
| Chondrichthyes | Elasmobranchii | Orectolobiformes | Hemiscylliidae | Chpl | Chiloscyllium plagiosum | whitespotted bambooshark | 12 | NC_057721.1 (16944458..16966275, complement) | LOC122555017 | XM_043700596.1 | XP_043556531.1 | TLR7 | TLR7 | TLR7 |
| Chondrichthyes | Elasmobranchii | Orectolobiformes | Hemiscylliidae | Chpl | Chiloscyllium plagiosum | whitespotted bambooshark | 12 | NC_057721.1 (17003496..17035093, complement) | tlr7 | XM_043700597.1 | XP_043556532.1 | TLR7 | TLR7 | TLR8 |
| Chondrichthyes | Elasmobranchii | Orectolobiformes | Hemiscylliidae | Chpl | Chiloscyllium plagiosum | whitespotted bambooshark | 18 | NC_057727.1 (10569011..10594995) | tlr9 | XM_043707663.1 | XP_043563598.1 | tlr7 | TLR9 | TLR9 |
| Chondrichthyes | Elasmobranchii | Orectolobiformes | Hemiscylliidae | Chpl | Chiloscyllium plagiosum | whitespotted bambooshark | 39 | NC_057748.1 (15833916..15848688, complement) | LOC122542273 | XM_043679848.1 | XP_043535783.1 | tlr11 | TLR13 | TLR21 |
| Chondrichthyes | Elasmobranchii | Orectolobiformes | Hemiscylliidae | Chpl | Chiloscyllium plagiosum | whitespotted bambooshark | 14 | NC_057723.1 (49340168 to 49342004, complement) | tlr22 | x | x | tlr11 | x | TLR22 |
| Chondrichthyes | Elasmobranchii | Orectolobiformes | Hemiscylliidae | Chpl | Chiloscyllium plagiosum | whitespotted bambooshark | 11 | NC_057720.1 (63314446..63323592, complement) | LOC122554467 | XM_043699540.1 | XP_043555475.1 | TLR1 | TLR2 | TLR27 |
| Chondrichthyes | Elasmobranchii | Orectolobiformes | Hemiscylliidae | Chpl | Chiloscyllium plagiosum | whitespotted bambooshark | 3 | NC_057748.1 (1693903..1698155) | LOC122542095 | XM_043679456.1 | XP_043535391.1 | TLR11 | TLR13 | TLR29 |
| Chondrichthyes | Elasmobranchii | Orectolobiformes | Hemiscylliidae | Chpu | Chiloscyllium punctatum | brownbanded bambooshark | 1 | CM104968.1(66343851 to 66346292) | x | x | x | TLR1 | x | TLR1 |
| Chondrichthyes | Elasmobranchii | Orectolobiformes | Hemiscylliidae | Chpu | Chiloscyllium punctatum | brownbanded bambooshark | 1 | CM104968.1(3885508 to 3887889) | x | x | x | tlr2 | x | TLR2 |
| Chondrichthyes | Elasmobranchii | Orectolobiformes | Hemiscylliidae | Chpu | Chiloscyllium punctatum | brownbanded bambooshark | 1 | CM104968.1(3831288 to 3833663) | x | x | x | tlr2 | x | TLR2 |
| Chondrichthyes | Elasmobranchii | Orectolobiformes | Hemiscylliidae | Chpu | Chiloscyllium punctatum | brownbanded bambooshark | 2 | CM104969.1(35408476 to 35430595) | x | x | x | tlr3 | x | TLR3 |
| Chondrichthyes | Elasmobranchii | Orectolobiformes | Hemiscylliidae | Chpu | Chiloscyllium punctatum | brownbanded bambooshark | 3 | CM104970.1(18072143 to 18074794, complement) | x | x | x | tlr5 | x | TLR5 |
| Chondrichthyes | Elasmobranchii | Orectolobiformes | Hemiscylliidae | Chpu | Chiloscyllium punctatum | brownbanded bambooshark | 3 | CM104970.1(18049242 to 18051170) | x | x | x | tlr5 | x | TLR5 |
| Chondrichthyes | Elasmobranchii | Orectolobiformes | Hemiscylliidae | Chpu | Chiloscyllium punctatum | brownbanded bambooshark | 15 | CM104982.1(30684258 to 30687461, complement) | x | x | x | tlr7 | x | TLR7 |
| Chondrichthyes | Elasmobranchii | Orectolobiformes | Hemiscylliidae | Chpu | Chiloscyllium punctatum | brownbanded bambooshark | 15 | CM104982.1(30625450 to 30628575) | x | x | x | tlr7 | x | TLR8 |
| Chondrichthyes | Elasmobranchii | Orectolobiformes | Hemiscylliidae | Chpu | Chiloscyllium punctatum | brownbanded bambooshark | 12 | CM104979.1( 13916870 to 13920007) | x | x | x | tlr7 | x | TLR9 |
| Chondrichthyes | Elasmobranchii | Orectolobiformes | Hemiscylliidae | Chpu | Chiloscyllium punctatum | brownbanded bambooshark | 25 | NC_092763.1(59939242 to 59939458) | x | x | x | tlr11 | x | TLR13 |
| Chondrichthyes | Elasmobranchii | Orectolobiformes | Hemiscylliidae | Chpu | Chiloscyllium punctatum | brownbanded bambooshark | 28 | NC_092766.1(18270800 to 18273640, complement) | x | x | x | tlr11 | x | TLR21 |
| Chondrichthyes | Elasmobranchii | Orectolobiformes | Hemiscylliidae | Chpu | Chiloscyllium punctatum | brownbanded bambooshark | 20 | CM104987.1(54428152 to 54430731) | x | x | x | tlr11 | x | TLR22 |
| Chondrichthyes | Elasmobranchii | Orectolobiformes | Hemiscylliidae | Chpu | Chiloscyllium punctatum | brownbanded bambooshark | 7 | CM104974.1(57311372 to 57313801) | x | x | x | TLR1 | x | TLR27 |
| Chondrichthyes | Elasmobranchii | Orectolobiformes | Hemiscylliidae | Chpu | Chiloscyllium punctatum | brownbanded bambooshark | 18 | CM104985.1(99317261 to 99320146) | x | x | x | TLR11 | x | TLR29 |
| Chondrichthyes | Elasmobranchii | Orectolobiformes | Stegostomatidae | Stti | Stegostoma tigrinum | zebra shark | 1 | NC_081354.1 (97608657..97663701, complement) | LOC125450897 | XM_048527266.2 | XP_048383223.2 | TLR1 | | TLR1 |
| Chondrichthyes | Elasmobranchii | Orectolobiformes | Stegostomatidae | Stti | Stegostoma tigrinum | zebra shark | 1 | NC_081354.1 (53296145..53319546) | LOC125452751 | XM_059645414.1 | XP_059501397.1 | TLR1 | | TLR2 |
| Chondrichthyes | Elasmobranchii | Orectolobiformes | Stegostomatidae | Stti | Stegostoma tigrinum | zebra shark | 1 | NC_081354.1 (53333256..53487752) | LOC132209546 | XM_059645426.1 | XP_059501409.1 | TLR1 | | TLR2 |
| Chondrichthyes | Elasmobranchii | Orectolobiformes | Stegostomatidae | Stti | Stegostoma tigrinum | zebra shark | 3 | NC_081356.1 (111755446..111782187, complement) | tlr3 | XM_059644834.1 | XP_059500817.1 | tlr3 | | TLR3 |
| Chondrichthyes | Elasmobranchii | Orectolobiformes | Stegostomatidae | Stti | Stegostoma tigrinum | zebra shark | 2 | NC_081355.1 (136453319..136599594, complement) | LOC125466711 | XM_048561639.2 | XP_048417596.1 | tlr3 | | TLR3 |
| Chondrichthyes | Elasmobranchii | Orectolobiformes | Stegostomatidae | Stti | Stegostoma tigrinum | zebra shark | 4 | CM060512.1(8652085 to 8654742) | tlr5 | x | x | tlr5 | | TLR5 |
| Chondrichthyes | Elasmobranchii | Orectolobiformes | Stegostomatidae | Stti | Stegostoma tigrinum | zebra shark | 12 | NC_081365.1 (63989306..63994030) | TLR7 | XM_059649999.1 | XP_059505982.1 | TLR7 | | TLR7 |
| Chondrichthyes | Elasmobranchii | Orectolobiformes | Stegostomatidae | Stti | Stegostoma tigrinum | zebra shark | 12 | NC_081365.1 (64041750..64044563) | LOC132206004 | XM_059650000.1 | XP_059505983.1 | TLR7 | TLR7 | TLR8 |
| Chondrichthyes | Elasmobranchii | Orectolobiformes | Stegostomatidae | Stti | Stegostoma tigrinum | zebra shark | 11 | NC_081364.1 (11764490..11782683) | | XM_048547841.2 | XP_048403798.1 | tlr7 | TLR9 | TLR9 |
| Chondrichthyes | Elasmobranchii | Orectolobiformes | Stegostomatidae | Stti | Stegostoma tigrinum | zebra shark | 47 | NC_081400.1 (7725474..7748001, complement) | LOC125449773 | XM_048525663.2 | XP_048381620.2 | tlr11 | TLR13 | TLR21 |
| Chondrichthyes | Elasmobranchii | Orectolobiformes | Stegostomatidae | Stti | Stegostoma tigrinum | zebra shark | 13 | NC_081366.1 (48344660..48349091, complement) | tlr22 | XM_048543920.2 | XP_048399877.2 | tlr11 | TLR22 | TLR22 |
| Chondrichthyes | Elasmobranchii | Orectolobiformes | Stegostomatidae | Stti | Stegostoma tigrinum | zebra shark | 8 | NC_081361.1 (44606651..44614924) | LOC125456697 | XM_048540076.2 | XP_048396033.1 | TLR1 | TLR2 | TLR27 |
| Chondrichthyes | Elasmobranchii | Orectolobiformes | Stegostomatidae | Stti | Stegostoma tigrinum | zebra shark | 38 | NC_081391.1 (25000049..25003225, complement) | LOC132206424 | XM_059640548.1 | XP_059496531.1 | TLR11 | TLR13 | TLR29 |
| Chondrichthyes | Elasmobranchii | Orectolobiformes | Ginglymostomatidae | Gici | Ginglymostoma cirratum | nurse shark | JAHRHZ010000049.1 | JAHRHZ010000049.1(20754208 to 20756676) | x | x | x | TLR1 | x | TLR1 |
| Chondrichthyes | Elasmobranchii | Orectolobiformes | Ginglymostomatidae | Gici | Ginglymostoma cirratum | nurse shark | JAHRHZ010000142.1 | JAHRHZ010000142.1(1385509 to 1387884) | x | x | x | tlr1 | x | TLR2 |
| Chondrichthyes | Elasmobranchii | Orectolobiformes | Ginglymostomatidae | Gici | Ginglymostoma cirratum | nurse shark | JAHRHZ010000097.1 | JAHRHZ010000097.1(32281 to 34662) | x | x | x | tlr1 | x | TLR2 |
| Chondrichthyes | Elasmobranchii | Orectolobiformes | Ginglymostomatidae | Gici | Ginglymostoma cirratum | nurse shark | JAHRHZ010000266.1 | JAHRHZ010000266.1(20427578 to 20441380) | x | x | x | tlr3 | x | TLR3 |
| Chondrichthyes | Elasmobranchii | Orectolobiformes | Ginglymostomatidae | Gici | Ginglymostoma cirratum | nurse shark | JAHRHZ010000279.1 | JAHRHZ010000279.1(7207 to 9864) | x | x | x | tlr5 | x | TLR5 |
| Chondrichthyes | Elasmobranchii | Orectolobiformes | Ginglymostomatidae | Gici | Ginglymostoma cirratum | nurse shark | JAHRHZ010000031.1 | JAHRHZ010000031.1(13447145 to 13450330) | x | x | x | tlr7 | x | TLR7 |
| Chondrichthyes | Elasmobranchii | Orectolobiformes | Ginglymostomatidae | Gici | Ginglymostoma cirratum | nurse shark | JAHRHZ010000031.1 | JAHRHZ010000031.1( 13492437 to 13495553) | x | x | x | tlr7 | x | TLR8 |
| Chondrichthyes | Elasmobranchii | Orectolobiformes | Ginglymostomatidae | Gici | Ginglymostoma cirratum | nurse shark | JAHRHZ010000065.1 | JAHRHZ010000065.1(9381033 to 9384182) | x | x | x | tlr7 | x | TLR9 |
| Chondrichthyes | Elasmobranchii | Orectolobiformes | Ginglymostomatidae | Gici | Ginglymostoma cirratum | nurse shark | GIWU01133779.1 | GIWU01133779.1(1..585) | x | x | x | tlr11 | x | TLR21 |
| Chondrichthyes | Elasmobranchii | Orectolobiformes | Ginglymostomatidae | Gici | Ginglymostoma cirratum | nurse shark | GIWU01297244.1 | GIWU01297244.1(1..351) | x | x | x | tlr11 | x | TLR21 |
| Chondrichthyes | Elasmobranchii | Orectolobiformes | Ginglymostomatidae | Gici | Ginglymostoma cirratum | nurse shark | JAHRHZ010000289.1 | JAHRHZ010000289.1(6211661 to 6214498) | x | x | x | tlr11 | x | TLR22 |
| Chondrichthyes | Elasmobranchii | Orectolobiformes | Ginglymostomatidae | Gici | Ginglymostoma cirratum | nurse shark | JAHRHZ010000058.1 | JAHRHZ010000058.1(17782758 to 17785187) | x | x | x | TLR1 | x | TLR27 |
| Chondrichthyes | Elasmobranchii | Orectolobiformes | Ginglymostomatidae | Gici | Ginglymostoma cirratum | nurse shark | JAHRHZ010000008.1 | JAHRHZ010000008.1(19494322 to 19497210) | x | x | x | TLR11 | x | TLR29 |
| Chondrichthyes | Elasmobranchii | Heterodontiformes | Heterodontidae | Hefr | Heterodontus francisci | horn shark | 1 | NC_090371.1 (139072833..139109352) | TLR1 | XM_068036779.1 | XP_067892880.1 | TLR1 | | TLR1 |
| Chondrichthyes | Elasmobranchii | Heterodontiformes | Heterodontidae | Hefr | Heterodontus francisci | horn shark | 1 | NC_090371.1 (205179104..205200902, complement) | LOC137374491 | XM_068040675.1 | XP_067896776.1 | tlr1 | | TLR2 |
| Chondrichthyes | Elasmobranchii | Heterodontiformes | Heterodontidae | Hefr | Heterodontus francisci | horn shark | 1 | NC_090371.1 (205106722..205127279, complement) | LOC137374479 | XM_068040663.1 | XP_067896764.1 | tlr1 | | TLR2 |
| Chondrichthyes | Elasmobranchii | Heterodontiformes | Heterodontidae | Hefr | Heterodontus francisci | horn shark | 2 | NC_090372.1 (191090753..191269973, complement) | LOC137349404 | XM_068014936.1 | XP_067871037.1 | tlr3 | | TLR3 |
| Chondrichthyes | Elasmobranchii | Heterodontiformes | Heterodontidae | Hefr | Heterodontus francisci | horn shark | 4 | NC_090374.1 (42522784..42544166) | cd180 | XM_068029423.1 | XP_067885524.1 | tlr3 | CD180 | TLR3 |
| Chondrichthyes | Elasmobranchii | Heterodontiformes | Heterodontidae | Hefr | Heterodontus francisci | horn shark | 3 | NC_090373.1 (195507891..195590446) | LOC137365608 | XM_068028012.1 | XP_067884113.1 | tlr5 | | TLR5 |
| Chondrichthyes | Elasmobranchii | Heterodontiformes | Heterodontidae | Hefr | Heterodontus francisci | horn shark | 3 | NC_090373.1 (196912706 to 196914016) | x | x | x | TLR5 | x | TLR5 |
| Chondrichthyes | Elasmobranchii | Heterodontiformes | Heterodontidae | Hefr | Heterodontus francisci | horn shark | 10 | NC_090380.1 (38544164..38569117, complement) | TLR7 | XM_068040471.1 | XP_067896572.1 | TLR7 | | TLR7 |
| Chondrichthyes | Elasmobranchii | Heterodontiformes | Heterodontidae | Hefr | Heterodontus francisci | horn shark | 10 | NC_090380.1 (38459099..38518809, complement) | LOC137374405 | XM_068040466.1 | XP_067896567.1 | TLR7 | | TLR8 |
| Chondrichthyes | Elasmobranchii | Heterodontiformes | Heterodontidae | Hefr | Heterodontus francisci | horn shark | 1 | NC_090389.1 (80126456..80137612, complement) | tlr9 | XM_068052085.1 | XP_067908186.1 | TLR7 | TLR9 | TLR9 |
| Chondrichthyes | Elasmobranchii | Heterodontiformes | Heterodontidae | Hefr | Heterodontus francisci | horn shark | 15 | NC_090385.1 (29910688..29923196, complement) | LOC137377403 | XM_068046982.1 | XP_067903083.1 | TLR11 | TLR13 | TLR13 |
| Chondrichthyes | Elasmobranchii | Heterodontiformes | Heterodontidae | Hefr | Heterodontus francisci | horn shark | 46 | NC_090416.1 (15355434..15377005) | TLR18 | XM_068022720.1 | XP_067878821.1 | TLR1 | TLR18 | TLR18 |

| | | | | | | | | | | | | | | |
|---|---|---|---|---|---|---|---|---|---|---|---|---|---|---|
| Chondrichthyes | Elasmobranchii | Heterodontiformes | Heterodontidae | Hefr | Heterodontus francisci | horn shark | 46 | NC_090416.1 (15373740 to 15375089) | x | x | x | tlr1 | x | TLR18 |
| Chondrichthyes | Elasmobranchii | Heterodontiformes | Heterodontidae | Hefr | Heterodontus francisci | horn shark | 46 | NC_090416.1 (15358472 to 15360634) | x | x | x | tlr1 | x | TLR18 |
| Chondrichthyes | Elasmobranchii | Heterodontiformes | Heterodontidae | Hefr | Heterodontus francisci | horn shark | 46 | NC_090416.1 (9041921..9080613) | tlr21 | XM_068022777.1 | XP_067878878.1 | tlr11 | TLR21 | TLR21 |
| Chondrichthyes | Elasmobranchii | Heterodontiformes | Heterodontidae | Hefr | Heterodontus francisci | horn shark | 12 | NC_090382.1 (54439830..54462431) | tlr22 | XM_068043430.1 | XP_067899531.1 | tlr11 | TLR22 | TLR22 |
| Chondrichthyes | Elasmobranchii | Heterodontiformes | Heterodontidae | Hefr | Heterodontus francisci | horn shark | 8 | NC_090378.1 (77595762..77603065, complement) | LOC137372554 | XM_068036468.1 | XP_067892569.1 | TLR1 | TLR2 | TLR27 |
| Chondrichthyes | Elasmobranchii | Heterodontiformes | Heterodontidae | Hefr | Heterodontus francisci | horn shark | 41 | NC_090411.1 (35233160..35238219, complement) | LOC137353480 | XM_068019807.1 | XP_067875908.1 | TLR11 | TLR13 | TLR29 |
| Chondrichthyes | Elasmobranchii | Heterodontiformes | Heterodontidae | Hefr | Heterodontus francisci | horn shark | 29 | NC_090399.1 (14399286..14429820, complement) | LOC137346147 | XM_068009483.1 | XP_067865584.1 | TLR11 | TLR13 | TLR30 |
| Chondrichthyes | Elasmobranchii | Pristiophoriformes | Pristiophoridae | Prja | Pristiophorus japonicus | Japanese sawshark | 2 | NC_091978.1 (176923914..176943442) | LOC139232836 | XM_070863331.1 | XP_070719432.1 | TLR1 | TLR1 | TLR1 |
| Chondrichthyes | Elasmobranchii | Pristiophoriformes | Pristiophoridae | Prja | Pristiophorus japonicus | Japanese sawshark | 2 | NC_091978.1 (278011621..278018213, complement) | LOC139230462 | XM_070862286.1 | XP_070718387.1 | TLR2 | TLR2 | TLR2 |
| Chondrichthyes | Elasmobranchii | Pristiophoriformes | Pristiophoridae | Prja | Pristiophorus japonicus | Japanese sawshark | 2 | NC_091978.1 (278285417..278324304) | LOC139246533 | XM_070871420.1 | XP_070727521.1 | TLR2 | TLR2 | TLR2 |
| Chondrichthyes | Elasmobranchii | Pristiophoriformes | Pristiophoridae | Prja | Pristiophorus japonicus | Japanese sawshark | 1 | NC_091977.1 (47274240..47302689) | LOC139263353 | XM_070879300.1 | XP_070735401.1 | tlr3 | | TLR3 |
| Chondrichthyes | Elasmobranchii | Pristiophoriformes | Pristiophoridae | Prja | Pristiophorus japonicus | Japanese sawshark | 7 | NC_091983.1 (216594868..216627877, complement) | LOC139267453 | XM_070885798.1 | XP_070741899.1 | tlr5 | | TLR5 |
| Chondrichthyes | Elasmobranchii | Pristiophoriformes | Pristiophoridae | Prja | Pristiophorus japonicus | Japanese sawshark | 7 | NC_091983.1( 219106038 to 219110738) | x | x | x | TLR5 | x | TLR5 |
| Chondrichthyes | Elasmobranchii | Pristiophoriformes | Pristiophoridae | Prja | Pristiophorus japonicus | Japanese sawshark | 11 | NC_091987.1 (113081061..113086206) | tlr7 | XM_070892734.1 | XP_070748835.1 | tlr7 | TLR7 | TLR7 |
| Chondrichthyes | Elasmobranchii | Pristiophoriformes | Pristiophoridae | Prja | Pristiophorus japonicus | Japanese sawshark | 11 | NC_091987.1 (113208670..113213501) | LOC139276355 | XM_070894210.1 | XP_070750311.1 | tlr8 | TLR8 | TLR8 |
| Chondrichthyes | Elasmobranchii | Pristiophoriformes | Pristiophoridae | Prja | Pristiophorus japonicus | Japanese sawshark | 11 | NC_091987.1 (113160289..113163099) | LOC139275565 | XM_070892735.1 | XP_070748836.1 | tlr8 | TLR8 | TLR8 |
| Chondrichthyes | Elasmobranchii | Pristiophoriformes | Pristiophoridae | Prja | Pristiophorus japonicus | Japanese sawshark | 12 | NC_091988.1 (62407150..62417616, complement) | tlr9 | XM_070894779.1 | XP_070750880.1 | tlr7 | TLR9 | TLR9 |
| Chondrichthyes | Elasmobranchii | Pristiophoriformes | Pristiophoridae | Prja | Pristiophorus japonicus | Japanese sawshark | 6 | NC_091982.1 (33174365..33177202) | LOC139266093 | XM_070883943.1 | XP_070740044.1 | tlr11 | TLR13 | TLR13 |
| Chondrichthyes | Elasmobranchii | Pristiophoriformes | Pristiophoridae | Prja | Pristiophorus japonicus | Japanese sawshark | 4 | NC_091980.1 (99893406..99896294) | LOC139262471 | XM_070877656.1 | XP_070733757.1 | tlr11 | TLR13 | TLR22 |
| Chondrichthyes | Elasmobranchii | Pristiophoriformes | Pristiophoridae | Prja | Pristiophorus japonicus | Japanese sawshark | 8 | NC_091984.1 (98135049..98141773, complement) | LOC139268652 | XM_070887153.1 | XP_070743254.1 | TLR1 | TLR2 | TLR27 |
| Chondrichthyes | Elasmobranchii | Pristiophoriformes | Pristiophoridae | Prja | Pristiophorus japonicus | Japanese sawshark | 19 | NC_091995.1 (98695541..98698423) | tlr21 | XM_070862135.1 | XP_070718236.1 | tlr11 | TLR21 | TLR29 |
| Chondrichthyes | Elasmobranchii | Pristiophoriformes | Pristiophoridae | Prja | Pristiophorus japonicus | Japanese sawshark | 19 | NC_091995.1 (15220733..15221768, complement) | LOC139230129 | XM_070861973.1 | XP_070718074.1 | TLR11 | TLR13 | TLR30 |
| Chondrichthyes | Batoidea | Rajiformes | Rajidae | Leer | Leucoraja erinaceus | little skate | 1 | NC_073377.1 (74065689..74101920) | LOC129698488 | XM_055637607.1 | XP_055493582.1 | TLR1 | TLR6 | |
| Chondrichthyes | Batoidea | Rajiformes | Rajidae | Leer | Leucoraja erinaceus | little skate | 1 | NC_073377.1 (78300159..78305999, complement) | LOC129706901 | x | x | | TLR7 | |
| Chondrichthyes | Batoidea | Rajiformes | Rajidae | Leer | Leucoraja erinaceus | little skate | 1 | NC_073377.1 (139103721..139136893) | tlr2 | XM_055644370.1 | XP_055500345.1 | tlr1 | TLR2 | |
| Chondrichthyes | Batoidea | Rajiformes | Rajidae | Leer | Leucoraja erinaceus | little skate | 3 | NC_073379.1 (7566529..7586880, complement) | tlr3 | XM_055631988.1 | XP_055487963.1 | tlr3 | TLR3 | |
| Chondrichthyes | Batoidea | Rajiformes | Rajidae | Leer | Leucoraja erinaceus | little skate | 1 | NC_073377.1 (126249347..126253514, complement) | LOC129695899 | x | x | TLR7 | TLR7 | |
| Chondrichthyes | Batoidea | Rajiformes | Rajidae | Leer | Leucoraja erinaceus | little skate | 1 | NC_073377.1 (127501396..127503202) | LOC129696004 | XM_055633442.1 | XP_055489417.1 | TLR7 | TLR7 | |
| Chondrichthyes | Batoidea | Rajiformes | Rajidae | Leer | Leucoraja erinaceus | little skate | 13 | NC_073389.1 (8590181..8644928, complement) | LOC129703103 | XM_055645301.1 | XP_055501276.1 | tlr7 | uncharacterized | TLR7 |
| Chondrichthyes | Batoidea | Rajiformes | Rajidae | Leer | Leucoraja erinaceus | little skate | 13 | NC_073389.1( 8597027 to 8600510) | x | x | x | tlr7 | x | TLR8 |
| Chondrichthyes | Batoidea | Rajiformes | Rajidae | Leer | Leucoraja erinaceus | little skate | 16 | NC_073392.1 (814222..818674) | LOC129704440 | XM_055647507.1 | XP_055503482.1 | tlr7 | TLR7 | TLR9 |
| Chondrichthyes | Batoidea | Rajiformes | Rajidae | Leer | Leucoraja erinaceus | little skate | 16 | NC_073392.1 (29314978..29319881) | LOC129704712 | XM_055648021.1 | XP_055503996.1 | tlr7 | TLR9 | TLR9 |
| Chondrichthyes | Batoidea | Rajiformes | Rajidae | Leer | Leucoraja erinaceus | little skate | Unplaced Scaffold | NW_026576869.1 (10599..14294, complement) | LOC129695050 | XM_055631810.1 | XP_055487785.1 | tlr11 | TLR13 | TLR21 |
| Chondrichthyes | Batoidea | Rajiformes | Rajidae | Leer | Leucoraja erinaceus | little skate | 11 | NC_073387.1 (36367718..36374259) | tlr22 | XM_055643028.1 | XP_055499003.1 | tlr11 | TLR22 | TLR22 |
| Chondrichthyes | Batoidea | Rajiformes | Rajidae | Leer | Leucoraja erinaceus | little skate | 10 | NC_073386.1 (47812747..47810303, complement) | x | x | x | TLR1 | x | TLR27 |
| Chondrichthyes | Batoidea | Rajiformes | Rajidae | Leer | Leucoraja erinaceus | little skate | 37 | NC_073413.1 (3991618..3996465) | LOC129713793 | XM_055663095.1 | XP_055519070.1 | TLR11 | TLR13 | TLR29 |
| Chondrichthyes | Batoidea | Rajiformes | Rajidae | Rabr | Raja brachyura | blonde ray | 1 | OY740781.1(89824803 -89827184) | x | x | x | TLR1 | x | TLR1 |
| Chondrichthyes | Batoidea | Rajiformes | Rajidae | Rabr | Raja brachyura | blonde ray | 1 | OY740781.1(89871384 -89873765) | x | x | x | TLR1 | x | TLR1 |
| Chondrichthyes | Batoidea | Rajiformes | Rajidae | Rabr | Raja brachyura | blonde ray | 1 | OY740781.1(89902665 -89905046) | x | x | x | TLR1 | x | TLR1 |
| Chondrichthyes | Batoidea | Rajiformes | Rajidae | Rabr | Raja brachyura | blonde ray | 1 | OY740781.1(156259219 to 156261594) | x | x | x | tlr1 | x | TLR2 |
| Chondrichthyes | Batoidea | Rajiformes | Rajidae | Rabr | Raja brachyura | blonde ray | 3 | OY740783.1(9456948 to 9475399) | x | x | x | tlr3 | x | TLR3 |
| Chondrichthyes | Batoidea | Rajiformes | Rajidae | Rabr | Raja brachyura | blonde ray | 12 | OY740792.1(22625707 to 22628962) | x | x | x | tlr7 | x | x |
| Chondrichthyes | Batoidea | Rajiformes | Rajidae | Rabr | Raja brachyura | blonde ray | 12 | OY740792.1(24925074 to 24928332) | x | x | x | tlr7 | x | x |
| Chondrichthyes | Batoidea | Rajiformes | Rajidae | Rabr | Raja brachyura | blonde ray | 12 | OY740792.1( 58036991 to 58040131) | x | x | x | tlr7 | x | TLR7 |
| Chondrichthyes | Batoidea | Rajiformes | Rajidae | Rabr | Raja brachyura | blonde ray | 12 | OY740792.1(58069386 to 58072523) | x | x | x | tlr7 | x | TLR8 |
| Chondrichthyes | Batoidea | Rajiformes | Rajidae | Rabr | Raja brachyura | blonde ray | 16 | OY740796.1(19103014 to 19106169) | x | x | x | tlr7 | x | TLR9 |
| Chondrichthyes | Batoidea | Rajiformes | Rajidae | Rabr | Raja brachyura | blonde ray | CAUPSL010000959.1 | CAUPSL010000959.1(3105 to 5201) | x | x | x | tlr1 | x | TLR18 |
| Chondrichthyes | Batoidea | Rajiformes | Rajidae | Rabr | Raja brachyura | blonde ray | CAUPSL010001590.1 | CAUPSL010001590.1(58319 to 60661) | x | x | x | tlr11 | x | TLR21 |
| Chondrichthyes | Batoidea | Rajiformes | Rajidae | Rabr | Raja brachyura | blonde ray | 11 | OY740791.1(27449991 to 27452894) | x | x | x | tlr11 | x | TLR22 |
| Chondrichthyes | Batoidea | Rajiformes | Rajidae | Rabr | Raja brachyura | blonde ray | 10 | OY740790.1(16179662 to 16182070) | x | x | x | TLR1 | x | TLR27 |
| Chondrichthyes | Batoidea | Rajiformes | Rajidae | Rabr | Raja brachyura | blonde ray | X | OY740817.1(9944585 to 9947473) | x | x | x | TLR11 | x | TLR29 |
| Chondrichthyes | Batoidea | Rajiformes | Rajidae | Amra | Amblyraja radiata | thorny skate | 1 | NC_045956.1 (99972225..99987274, complement) | LOC116976921 | XM_033026986.1 | XP_032882877.1 | TLR1 | TLR6 | TLR1 |
| Chondrichthyes | Batoidea | Rajiformes | Rajidae | Amra | Amblyraja radiata | thorny skate | 1 | NC_045956.1 (99957453..99963774, complement) | LOC116980894 | XM_033033523.1 | XP_032889414.1 | TLR1 | TLR6 | TLR1 |
| Chondrichthyes | Batoidea | Rajiformes | Rajidae | Amra | Amblyraja radiata | thorny skate | 1 | NC_045956.1 (99938093..99940550, complement) | LOC116980807 | XM_033033440.1 | XP_032889331.1 | TLR1 | TLR6 | TLR1 |
| Chondrichthyes | Batoidea | Rajiformes | Rajidae | Amra | Amblyraja radiata | thorny skate | 1 | NC_045956.1 (31682556..31713597, complement) | tlr2 | XM_033019779.1 | XP_032875670.1 | tlr1 | TLR2 | TLR2 |
| Chondrichthyes | Batoidea | Rajiformes | Rajidae | Amra | Amblyraja radiata | thorny skate | 3 | NC_045958.1 (109258812..109276930) | tlr3 | XM_033018573.1 | XP_032874464.1 | tlr3 | TLR3 | TLR3 |
| Chondrichthyes | Batoidea | Rajiformes | Rajidae | Amra | Amblyraja radiata | thorny skate | 14 | NC_045969.1 (50664790..50680730) | LOC116980762 | XM_033033333.1 | XP_032889224.1 | TLR7 | TLR7 | TLR7 |
| Chondrichthyes | Batoidea | Rajiformes | Rajidae | Amra | Amblyraja radiata | thorny skate | 14 | NC_045969.1 (50724377..50727162) | LOC116980925 | XM_033033522.1 | XP_032889413.1 t | TLR7 | TLR8 | TLR8 |
| Chondrichthyes | Batoidea | Rajiformes | Rajidae | Amra | Amblyraja radiata | thorny skate | 20 | NC_045975.1 (35259416..35264016) | LOC116984411 | x | x | TLR7 | TLR7 | |
| Chondrichthyes | Batoidea | Rajiformes | Rajidae | Amra | Amblyraja radiata | thorny skate | 1 | NC_045956.1 (150026135..150031025, complement) | LOC116984305 | x | x | TLR7 | TLR7 | |
| Chondrichthyes | Batoidea | Rajiformes | Rajidae | Amra | Amblyraja radiata | thorny skate | 18 | NC_045973.1 (30995027..31000260) | tlr9 | XM_033037038.1 | XP_032892929.1 | tlr7 | TLR9 | TLR9 |
| Chondrichthyes | Batoidea | Rajiformes | Rajidae | Amra | Amblyraja radiata | thorny skate | 11 | NC_045966.1 (25418014..25422921, complement) | LOC116978441 | XM_033029571.1 | XP_032885462.1 | tlr11 | TLR13 | TLR22 |
| Chondrichthyes | Batoidea | Rajiformes | Rajidae | Amra | Amblyraja radiata | thorny skate | 10 | NC_045965.1 (16235607..16255977) | LOC116977601 | XM_033028177.1 | XP_032884068.1 | TLR1 | x | TLR27 |
| Chondrichthyes | Batoidea | Rajiformes | Rajidae | Amra | Amblyraja radiata | thorny skate | 38 | NC_045993.1 (9424092..9427023, complement) | LOC116967007 | XM_033013428.1 | XP_032869319.1 | TLR11 | TLR13 | TLR29 |
| Chondrichthyes | Batoidea | Rhinopristiformes | Rhinidae | Rhan | Rhina ancylostoma | bowmouth guitarfish | JARFYF010000052.1 | JARFYF010000052.1(66227222 to 66229606) | x | x | x | TLR1 | x | TLR1 |
| Chondrichthyes | Batoidea | Rhinopristiformes | Rhinidae | Rhan | Rhina ancylostoma | bowmouth guitarfish | JARFYF010008249.1 | JARFYF010008249.1(all) | x | x | x | x | x | TLR2 |
| Chondrichthyes | Batoidea | Rhinopristiformes | Rhinidae | Rhan | Rhina ancylostoma | bowmouth guitarfish | JARFYF010000104.1 | JARFYF010000104.1(19736577 to 19738430) | x | x | x | tlr3 | x | TLR3 |
| Chondrichthyes | Batoidea | Rhinopristiformes | Rhinidae | Rhan | Rhina ancylostoma | bowmouth guitarfish | JARFYF010000002.1 | JARFYF010000002.1(92921373 to 92923991) | x | x | x | tlr5 | x | TLR5 |
| Chondrichthyes | Batoidea | Rhinopristiformes | Rhinidae | Rhan | Rhina ancylostoma | bowmouth guitarfish | JARFYF010000074.1 | JARFYF010000074.1(70661179 to 70664322) | x | x | x | TLR7 | x | TLR7 |
| Chondrichthyes | Batoidea | Rhinopristiformes | Rhinidae | Rhan | Rhina ancylostoma | bowmouth guitarfish | JARFYF010000074.1 | JARFYF010000074.1(70636778 to 70639930) | x | x | x | TLR7 | x | TLR8 |
| Chondrichthyes | Batoidea | Rhinopristiformes | Rhinidae | Rhan | Rhina ancylostoma | bowmouth guitarfish | JARFYF010000094.1 | JARFYF010000094.1(8438594 to 8441749) | x | x | x | tlr7 | x | TLR9 |
| Chondrichthyes | Batoidea | Rhinopristiformes | Rhinidae | Rhan | Rhina ancylostoma | bowmouth guitarfish | JARFYF010000073.1 | JARFYF010000073.1(6428739 to 6431042) | x | x | x | tlr1 | x | TLR18 |
| Chondrichthyes | Batoidea | Rhinopristiformes | Rhinidae | Rhan | Rhina ancylostoma | bowmouth guitarfish | JARFYF010000073.1 | JARFYF010000073.1(1132813 to 1135300) | x | x | x | tlr11 | x | TLR21 |
| Chondrichthyes | Batoidea | Rhinopristiformes | Rhinidae | Rhan | Rhina ancylostoma | bowmouth guitarfish | JARFYF010000074.1 | JARFYF010000074.1(36022949 to 36025842) | x | x | x | tlr11 | x | TLR22 |
| Chondrichthyes | Batoidea | Rhinopristiformes | Rhinidae | Rhan | Rhina ancylostoma | bowmouth guitarfish | JARFYF010000063.1 | JARFYF010000063.1(43383056 to 43385521) | x | x | x | TLR1 | x | TLR27 |
| Chondrichthyes | Batoidea | Rhinopristiformes | Rhinidae | Rhan | Rhina ancylostoma | bowmouth guitarfish | JARFYF010000067.1 | JARFYF010000067.1(11542711 to 11545548) | x | x | x | TLR11 | x | TLR29 |
| Chondrichthyes | Batoidea | Rhinopristiformes | Pristidae | Prpe | Pristis pectinata | smalltooth sawfish | 2 | NC_067406.1 (69143095..69158511) | LOC127586075 | XM_052043898.1 | XP_051899858.1 | TLR1 | | |
| Chondrichthyes | Batoidea | Rhinopristiformes | Pristidae | Prpe | Pristis pectinata | smalltooth sawfish | 2 | NC_067406.1 (96749373..96771627) | LOC127582338 | XM_052037498.1 | XP_051893458.1 | tlr1 | | |
| Chondrichthyes | Batoidea | Rhinopristiformes | Pristidae | Prpe | Pristis pectinata | smalltooth sawfish | 2 | NC_067406.1 (96828913..96831290) | LOC127584118 | XM_052040705.1 | XP_051896665.1 | tlr1 | | |
| Chondrichthyes | Batoidea | Rhinopristiformes | Pristidae | Prpe | Pristis pectinata | smalltooth sawfish | 7 | NC_067411.1 (20818827..20838657) | tlr3 | XM_052020093.1 | XP_051876053.1 | tlr3 | | |
| Chondrichthyes | Batoidea | Rhinopristiformes | Pristidae | Prpe | Pristis pectinata | smalltooth sawfish | 10 | NC_067414.1 (96758245..96773890) | LOC127574781 | XM_052024120.1 | XP_051880080.1 | tlr5 | | |
| Chondrichthyes | Batoidea | Rhinopristiformes | Pristidae | Prpe | Pristis pectinata | smalltooth sawfish | 4 | NC_067408.1 (72849621..72883624, complement) | LOC127569651 | XM_052014451.1 | XP_051870411.1 | TLR7 | | |
| Chondrichthyes | Batoidea | Rhinopristiformes | Pristidae | Prpe | Pristis pectinata | smalltooth sawfish | 4 | NC_067408.1 (72824512..72839020, complement) | LOC127569650 | XM_052014450.1 | XP_051870410.1 | TLR7 | | |
| Chondrichthyes | Batoidea | Rhinopristiformes | Pristidae | Prpe | Pristis pectinata | smalltooth sawfish | 6 | NC_067410.1 (9638155..9641688) | LOC127571301 | XM_052017573.1 | XP_051873533.1 | tlr7 | TLR7 | TLR9 |
| Chondrichthyes | Batoidea | Rhinopristiformes | Pristidae | Prpe | Pristis pectinata | smalltooth sawfish | 40 | NC_067443.1 (8226903..8229460) | tlr18 | XM_052045743.1 | XP_051901703.1 | tlr1 | TLR18 | TLR18 |
| Chondrichthyes | Batoidea | Rhinopristiformes | Pristidae | Prpe | Pristis pectinata | smalltooth sawfish | 40 | NC_067443.1 (2241853..2252603) | tlr21 | XM_052045853.1 | XP_051901813.1 | tlr11 | TLR21 | TLR21 |
| Chondrichthyes | Batoidea | Rhinopristiformes | Pristidae | Prpe | Pristis pectinata | smalltooth sawfish | 4 | CM019858.1(37397552 to 37400446, complement) | x | x | x | tlr11 | x | TLR22 |
| Chondrichthyes | Batoidea | Rhinopristiformes | Pristidae | Prpe | Pristis pectinata | smalltooth sawfish | 3 | NC_067407.1 (45355081..45365580, complement) | LOC127568397 | XM_052012080.1 | XP_051868040.1 | TLR1 | TLR2 | TLR27 |
| Chondrichthyes | Batoidea | Rhinopristiformes | Pristidae | Prpe | Pristis pectinata | smalltooth sawfish | 33 | NC_067437.1 (11674849..11677671, complement) | LOC127585417 | XM_052042841.1 | XP_051898801.1 | TLR11 | TLR13 | TLR29 |
| Chondrichthyes | Batoidea | Torpediniformes | Narcinidae | Naba | Narcine bancroftii | Caribbean electric ray | 3 | NC_091471.1 (113221207..113246598) | tlr1 | XM_069924941.1 | XP_069781042.1 | TLR1 | | TLR1 |
| Chondrichthyes | Batoidea | Torpediniformes | Narcinidae | Naba | Narcine bancroftii | Caribbean electric ray | 3 | NC_091471.1 (220945607..220951229, complement) | LOC138758158 | XM_069926682.1 | XP_069782783.1 | tlr1 | | TLR2 |
| Chondrichthyes | Batoidea | Torpediniformes | Narcinidae | Naba | Narcine bancroftii | Caribbean electric ray | 3 | NC_091471.1 (220869063..220878992, complement) | LOC138758157 | XM_069926681.1 | XP_069782782.1 | tlr1 | | TLR2 |
| Chondrichthyes | Batoidea | Torpediniformes | Narcinidae | Naba | Narcine bancroftii | Caribbean electric ray | 1 | NC_091469.1 (129471530..129491452, complement) | tlr3 | XM_069939237.1 | XP_069795338.1 | tlr3 | | TLR3 |
| Chondrichthyes | Batoidea | Torpediniformes | Narcinidae | Naba | Narcine bancroftii | Caribbean electric ray | 6 | NC_091474.1 (244180957..244218930) | LOC138737202 | XM_069887861.1 | XP_069743962.1 | tlr5 | | TLR5 |
| Chondrichthyes | Batoidea | Torpediniformes | Narcinidae | Naba | Narcine bancroftii | Caribbean electric ray | 7 | NC_091475.1 (46605207..46617102, complement) | TLR7 | XM_069889972.1 | XP_069746073.1 | TLR7 | | TLR7 |
| Chondrichthyes | Batoidea | Torpediniformes | Narcinidae | Naba | Narcine bancroftii | Caribbean electric ray | 7 | NC_091475.1 (46578571..46591717, complement) | LOC138738858 | XM_069889969.1 | XP_069746070.1 | TLR7 | | TLR8 |
| Chondrichthyes | Batoidea | Torpediniformes | Narcinidae | Naba | Narcine bancroftii | Caribbean electric ray | 5 | NC_091473.1 (5515006..5527945, complement) | tlr9 | XM_069936642.1 | XP_069792743.1 | TLR7 | TLR9 | TLR9 |
| Chondrichthyes | Batoidea | Torpediniformes | Narcinidae | Naba | Narcine bancroftii | Caribbean electric ray | 5 | NC_091473.1 (192581364..192608069, complement) | tlr18 | XM_069940450.1 | XP_069796551.1 | tlr1 | TLR18 | TLR18 |
| Chondrichthyes | Batoidea | Torpediniformes | Narcinidae | Naba | Narcine bancroftii | Caribbean electric ray | 9 | NC_091477.1 (34396562 to 34397339, complement) | x | x | x | tlr11 | x | TLR22 |
| Chondrichthyes | Batoidea | Torpediniformes | Narcinidae | Naba | Narcine bancroftii | Caribbean electric ray | 5 | NC_091473.1 (85486798..85499881) | LOC138763660 | XM_069938187.1 | XP_069794288.1 | TLR1 | TLR2 | TLR27 |
| Chondrichthyes | Batoidea | Torpediniformes | Narcinidae | Naba | Narcine bancroftii | Caribbean electric ray | 13 | NC_091481.1 (53151824..53157297) | LOC138748045 | XM_069907742.1 | XP_069763843.1 | TLR11 | TLR13 | TLR29 |
| Chondrichthyes | Batoidea | Myliobatiformes | Dasyatidae | Hysa | Hypanus sabinus | Atlantic stingray | 14 | NC_082719.1 (43537470..43544918, complement) | tlr1 | XM_059988629.1 | XP_059844612.1 | TLR1 | | TLR1 |
| Chondrichthyes | Batoidea | Myliobatiformes | Dasyatidae | Hysa | Hypanus sabinus | Atlantic stingray | 3 | NC_082708.1 (94116254..94137502) | tlr2 | XM_059964797.1 | XP_059820780.1 | tlr1 | | TLR2 |
| Chondrichthyes | Batoidea | Myliobatiformes | Dasyatidae | Hysa | Hypanus sabinus | Atlantic stingray | 7 | NC_082712.1 (32495427..32515230) | tlr3 | XM_059974391.1 | XP_059830374.1 | tlr3 | | TLR3 |
| Chondrichthyes | Batoidea | Myliobatiformes | Dasyatidae | Hysa | Hypanus sabinus | Atlantic stingray | 10 | NC_082715.1 (113360857..113390587, complement) | LOC132400994 | XM_059982676.1 | XP_059838659.1 | tlr5 | | TLR5 |
| Chondrichthyes | Batoidea | Myliobatiformes | Dasyatidae | Hysa | Hypanus sabinus | Atlantic stingray | 3 | NC_082708.1 (137082663..137091591) | LOC132390935 | XM_059963471.1 | XP_059819454.1 | tlr5 | | TLR5 |
| Chondrichthyes | Batoidea | Myliobatiformes | Dasyatidae | Hysa | Hypanus sabinus | Atlantic stingray | 4 | NC_082709.1 (121500843..121552263, complement) | tlr7 | XM_059968054.1 | XP_059824037.1 | TLR7 | | TLR7 |
| Chondrichthyes | Batoidea | Myliobatiformes | Dasyatidae | Hysa | Hypanus sabinus | Atlantic stingray | 4 | NC_082709.1 (121484719..121494114, complement) | LOC132393159 | XM_059968051.1 | XP_059824034.1 | TLR7 | | TLR8 |
| Chondrichthyes | Batoidea | Myliobatiformes | Dasyatidae | Hysa | Hypanus sabinus | Atlantic stingray | 19 | NC_082724.1 (69212651..69237849, complement) | tlr9 | XM_059944612.1 | XP_059800595.1 | tlr7 | TLR9 | TLR9 |
| Chondrichthyes | Batoidea | Myliobatiformes | Dasyatidae | Hysa | Hypanus sabinus | Atlantic stingray | 9 | NC_082714.1 (95520545..95576010, complement) | tlr18 | XM_059980282.1 | XP_059836265.1 | tlr1 | TLR18 | TLR18 |
| Chondrichthyes | Batoidea | Myliobatiformes | Dasyatidae | Hysa | Hypanus sabinus | Atlantic stingray | 15 | NC_082720.1 (36661534..36662443) | LOC132405214 | x | x | tlr11 | TLR13 | TLR22 |
| Chondrichthyes | Batoidea | Myliobatiformes | Dasyatidae | Hysa | Hypanus sabinus | Atlantic stingray | 11 | NC_082716.1 (63397312..63415916, complement) | LOC132402030 | XM_059984505.1 | XP_059840488.1 | TLR1 | TLR2 | TLR27 |

| | | | | | | | | | | | | | | |
|---|---|---|---|---|---|---|---|---|---|---|---|---|---|---|
| Chondrichthyes | Batoidea | Myliobatiformes | Dasyatidae | Hysa | Hypanus sabinus | Atlantic stingray | 29 | NC_082734.1 (10712772..10722178, complement) | LOC132382971 | XM_059953589.1 | XP_059809572.1 | TLR11 | TLR13 | TLR29 |
| Chondrichthyes | Batoidea | Myliobatiformes | Dasyatidae | Hybe | Hypanus berthalutzae | Lutz's stingray | JAUOBT010000350.1 | JAUOBT010000350.1(38617707 to 38620088) | x | x | x | TLR1 | x | TLR1 |
| Chondrichthyes | Batoidea | Myliobatiformes | Dasyatidae | Hybe | Hypanus berthalutzae | Lutz's stingray | JAUOBT010000215.1 | JAUOBT010000215.1(87493881 to 87496271) | x | x | x | tlr1 | x | TLR2 |
| Chondrichthyes | Batoidea | Myliobatiformes | Dasyatidae | Hybe | Hypanus berthalutzae | Lutz's stingray | JAUOBT010000258.1 | JAUOBT010000258.1(27609880 to 27621086) | x | x | x | tlr3 | x | TLR3 |
| Chondrichthyes | Batoidea | Myliobatiformes | Dasyatidae | Hybe | Hypanus berthalutzae | Lutz's stingray | JAUOBT010000336.1 | JAUOBT010000336.1(108979548 to 108982235) | x | x | x | tlr5 | x | TLR5 |
| Chondrichthyes | Batoidea | Myliobatiformes | Dasyatidae | Hybe | Hypanus berthalutzae | Lutz's stingray | JAUOBT010000220.1 | JAUOBT010000220.1( 105184979 to 105188098) | x | x | x | TLR7 | x | TLR7 |
| Chondrichthyes | Batoidea | Myliobatiformes | Dasyatidae | Hybe | Hypanus berthalutzae | Lutz's stingray | JAUOBT010000220.1 | JAUOBT010000220.1(105158683 to 105161799) | x | x | x | TLR7 | x | TLR8 |
| Chondrichthyes | Batoidea | Myliobatiformes | Dasyatidae | Hybe | Hypanus berthalutzae | Lutz's stingray | JAUOBT010000370.1 | JAUOBT010000370.1(64156377 to 64159532) | x | x | x | tlr7 | x | TLR9 |
| Chondrichthyes | Batoidea | Myliobatiformes | Dasyatidae | Hybe | Hypanus berthalutzae | Lutz's stingray | JAUOBT010000276.1 | JAUOBT010000276.1(73670071 to 73672563) | x | x | x | tlr1 | x | TLR18 |
| Chondrichthyes | Batoidea | Myliobatiformes | Dasyatidae | Hybe | Hypanus berthalutzae | Lutz's stingray | JAUOBT010000351.1 | JAUOBT010000351.1(30141037 to 30146395) | x | x | x | tlr11 | x | TLR22 |
| Chondrichthyes | Batoidea | Myliobatiformes | Dasyatidae | Hybe | Hypanus berthalutzae | Lutz's stingray | JAUOBT010000340.1 | JAUOBT010000340.1(62121966 to 62124512) | x | x | x | TLR1 | x | TLR27 |
| Chondrichthyes | Batoidea | Myliobatiformes | Dasyatidae | Hybe | Hypanus berthalutzae | Lutz's stingray | JAUOBT010000608.1 | JAUOBT010000608.1(9001531 to 9004414) | x | x | x | TLR11 | x | TLR29 |
| Chondrichthyes | Batoidea | Myliobatiformes | Mobulidae | Mohy | Mobula hypostoma | lesser devil ray | 3 | NC_086099.1 (70602974..70621451) | tlr1 | XM_063042459.1 | XP_062898529.1 | TLR1 | | TLR1 |
| Chondrichthyes | Batoidea | Myliobatiformes | Mobulidae | Mohy | Mobula hypostoma | lesser devil ray | 4 | NC_086100.1 (98613785..98649179) | tlr2 | XM_063046245.1 | XP_062902315.1 | tlr1 | | TLR2 |
| Chondrichthyes | Batoidea | Myliobatiformes | Mobulidae | Mohy | Mobula hypostoma | lesser devil ray | 5 | NC_086101.1 (168536688..168568427, complement) | tlr3 | XM_063049378.1 | XP_062905448.1 | tlr3 | | TLR3 |
| Chondrichthyes | Batoidea | Myliobatiformes | Mobulidae | Mohy | Mobula hypostoma | lesser devil ray | 2 | NC_086098.1 (122198773..122229931, complement) | LOC134342444 | XM_063040577.1 | XP_062896647.1 | tlr5 | | TLR5 |
| Chondrichthyes | Batoidea | Myliobatiformes | Mobulidae | Mohy | Mobula hypostoma | lesser devil ray | 6 | NC_086102.1 (74557884..74605498) | tlr7 | XM_063051004.1 | XP_062907074.1 | TLR7 | | TLR7 |
| Chondrichthyes | Batoidea | Myliobatiformes | Mobulidae | Mohy | Mobula hypostoma | lesser devil ray | 6 | NC_086102.1 (74623342..74631125) | LOC134347573 | XM_063049926.1 | XP_062905996.1 | TLR7 | | TLR8 |
| Chondrichthyes | Batoidea | Myliobatiformes | Mobulidae | Mohy | Mobula hypostoma | lesser devil ray | 15 | NC_086111.1 (586068..611816, complement) | tlr9 | XM_063068341.1 | XP_062924411.1 | tlr7 | TLR9 | TLR9 |
| Chondrichthyes | Batoidea | Myliobatiformes | Mobulidae | Mohy | Mobula hypostoma | lesser devil ray | 2 | NC_086098.1 (236082466..236143277) | tlr18 | XM_063041657.1 | XP_062897727.1 | tlr1 | TLR18 | TLR18 |
| Chondrichthyes | Batoidea | Myliobatiformes | Mobulidae | Mohy | Mobula hypostoma | lesser devil ray | 7 | NC_086103.1(58670000 to 58676199, complement) | x | x | x | tlr11 | x | TLR22 |
| Chondrichthyes | Batoidea | Myliobatiformes | Mobulidae | Mohy | Mobula hypostoma | lesser devil ray | 12 | NC_086108.1 (48588574..48611482) | LOC134355160 | XM_063064934.1 | XP_062921004.1 | TLR1 | TLR2 | TLR27 |
| Chondrichthyes | Batoidea | Myliobatiformes | Mobulidae | Mohy | Mobula hypostoma | lesser devil ray | 11 | NC_086107.1 (106266589..106277644) | LOC134353523 | XM_063061522.1 | XP_062917592.1 | TLR11 | TLR13 | TLR29 |
| Chondrichthyes | Batoidea | Myliobatiformes | Mobulidae | Mobi | Mobula birostris | giant manta | 3 | CM057525.1(70401117 to 70403498) | x | x | x | TLR1 | x | |
| Chondrichthyes | Batoidea | Myliobatiformes | Mobulidae | Mobi | Mobula birostris | giant manta | 12 | CM057534.1(98329806 to 98332187) | x | x | x | tlr1 | x | |
| Chondrichthyes | Batoidea | Myliobatiformes | Mobulidae | Mobi | Mobula birostris | giant manta | 5 | CM057527.1( 28619928 to 28630562) | x | x | x | tlr3 | x | |
| Chondrichthyes | Batoidea | Myliobatiformes | Mobulidae | Mobi | Mobula birostris | giant manta | 2 | CM057524.1(121844977 to 121847589) | x | x | x | tlr5 | x | |
| Chondrichthyes | Batoidea | Myliobatiformes | Mobulidae | Mobi | Mobula birostris | giant manta | 6 | CM057528.1(74946870 to 74950013) | x | x | x | TLR7 | x | |
| Chondrichthyes | Batoidea | Myliobatiformes | Mobulidae | Mobi | Mobula birostris | giant manta | 6 | CM057528.1(74973698 to 74976814) | x | x | x | TLR7 | x | |
| Chondrichthyes | Batoidea | Myliobatiformes | Mobulidae | Mobi | Mobula birostris | giant manta | 16 | CM057538.1(79621276 to 79624425) | x | x | x | tlr7 | x | TLR9 |
| Chondrichthyes | Batoidea | Myliobatiformes | Mobulidae | Mobi | Mobula birostris | giant manta | 2 | CM057524.1(16072570 to 16075002) | x | x | x | tlr1 | x | TLR18 |
| Chondrichthyes | Batoidea | Myliobatiformes | Mobulidae | Mobi | Mobula birostris | giant manta | 7 | CM057529.1(129372096 to 129378075) | x | x | x | tlr11 | x | TLR22 |
| Chondrichthyes | Batoidea | Myliobatiformes | Mobulidae | Mobi | Mobula birostris | giant manta | 12 | CM057534.1(66984902 to 66987364) | x | x | x | TLR1 | x | TLR27 |
| Chondrichthyes | Batoidea | Myliobatiformes | Mobulidae | Mobi | Mobula birostris | giant manta | 11 | CM057533.1(9243123 to 9246020) | x | x | x | TLR11 | x | TLR29 |
| Chondrichthyes | Holocephali | Chimaeriformes | Callorhinchidae | Cami | Callorhinchus milii | Australian ghostshark | Unplaced scaffold | NW_024704742.1 (47504656..47508771, complement) | tlr1 | XM_007889182.2 | XP_007887373.1 | TLR1 | TLR1 | |
| Chondrichthyes | Holocephali | Chimaeriformes | Callorhinchidae | Cami | Callorhinchus milii | Australian ghostshark | Unplaced scaffold | NW_024704742.1 (24618905..24621403, complement) | LOC103185499 | XM_007904038.2 | XP_007902229.2 | tlr1 | TLR2 | |
| Chondrichthyes | Holocephali | Chimaeriformes | Callorhinchidae | Cami | Callorhinchus milii | Australian ghostshark | Unplaced scaffold | NW_024704742.1 (24654804..24658131) | LOC103185476 | XM_007903987.2 | XP_007902178.1 | tlr1 | TLR2 | |
| Chondrichthyes | Holocephali | Chimaeriformes | Callorhinchidae | Cami | Callorhinchus milii | Australian ghostshark | Unplaced scaffold | NW_024704742.1 (56560128..56570300, complement) | tlr3 | XM_007892819.2 | XP_007891010.2 | tlr3 | TLR3 | |
| Chondrichthyes | Holocephali | Chimaeriformes | Callorhinchidae | Cami | Callorhinchus milii | Australian ghostshark | Unplaced scaffold | NW_024704749.1 (14722543..14736431) | tlr4 | x | x | TLR4 | TLR4 | |
| Chondrichthyes | Holocephali | Chimaeriformes | Callorhinchidae | Cami | Callorhinchus milii | Australian ghostshark | Unplaced scaffold | NW_024704745.1 (17716849..17723218) | tlr7 | XM_007891618.2 | XP_007889809.1 | tlr7 | TLR7 | TLR7 |
| Chondrichthyes | Holocephali | Chimaeriformes | Callorhinchidae | Cami | Callorhinchus milii | Australian ghostshark | Unplaced scaffold | NW_024704745.1 (17722990..17747488) | LOC103177589 | XM_042334075.1 | XP_042190009.1 | tlr7 | TLR8 | TLR8 |
| Chondrichthyes | Holocephali | Chimaeriformes | Callorhinchidae | Cami | Callorhinchus milii | Australian ghostshark | Unplaced scaffold | NW_024704753.1 (13230390..13235963, complement) | LOC103176640 | XM_007890298.2 | XP_007888489.2 | tlr7 | TLR7 | TLR9 |
| Chondrichthyes | Holocephali | Chimaeriformes | Callorhinchidae | Cami | Callorhinchus milii | Australian ghostshark | | NW_024704744.1 (84043092..84046879, complement) | LOC103177980 | XM_042333155.1 | XP_042189089.1 | TLR1 | TLR6 | TLR15 |
| Chondrichthyes | Holocephali | Chimaeriformes | Callorhinchidae | Cami | Callorhinchus milii | Australian ghostshark | Unplaced scaffold | NW_024704754.1 (4346245..4349106) | tlr22 | XM_007906265.2 | XP_007904456.1 | tlr11 | TLR22 | TLR22 |
| Chondrichthyes | Holocephali | Chimaeriformes | Callorhinchidae | Cami | Callorhinchus milii | Australian ghostshark | Unplaced scaffold | NW_024704746.1 (48708996..48711652) | LOC103181886 | XM_007898660.2 | XP_007896851.2 | tlr1 | TLR2 | TLR25 |
| Chondrichthyes | Holocephali | Chimaeriformes | Callorhinchidae | Cami | Callorhinchus milii | Australian ghostshark | Unplaced scaffold | NW_024704746.1 (9528107..9530521, complement) | LOC103180074 | XM_042334967.1 | XP_042190901.1 | TLR1 | TLR2 type-2 | TLR27 |
| Chondrichthyes | Holocephali | Chimaeriformes | Callorhinchidae | Cami | Callorhinchus milii | Australian ghostshark | Unplaced scaffold | NW_024705481.1 (28064..30505) | LOC121852316 | XM_042346415.1 | XP_042202349.1 | tlr1 | TLR2 | |
