## Supplementary Table 3 for "Cartilaginous fish inform the lineage-specific evolution and MHC association of the TLR family"

| Gene | No. aa residues | No. of species | ω (M0) | lmL M7 | lmL M8 | (-2lnaL) | Sgnificance | PAML | SLAC | FEL | MEME | FUBAR | Total no. of sites | |
|---|---|---|---|---|---|---|---|---|---|---|---|---|---|---|
| **TLR1** | **2448** | 23 | 0,49508 | -15540,879930 | -15491,503619 | | | 5, 9, 10, 12, 15, 16, 17, 18, 20, 25, 27, 30, 39, 45, 46, 68, 70, 71, 79, 87, 89, 107, 115, 127, 157, 170, 180, 185, 190, 198, 205, 206, 207, 208, 211, 212, 234, 237, 238, 239, 240, 241, 242, 246, 247, 48, 251, 252, 253, 254, 255, 256, 257, 258, 260, 261, 263, 264, 267, 271, 273,291, 293, 294, 295, 297, 298, 299, 302, 304, 308, 310, 313, 314, 317, 319, 320, 321, 322,323, 324, 325, 326, 327, 328, 329, 330, 336, 338, 339, 340, 341, 342, 343, 346, 347, 348, 349, 350, 356, 357, 364, 369,370, 373, 381, 384, 400, 402, 403, 406, 418, 428, 429, 431, 435, 449, 488, 495, 509, 586, 588, 622, 654,659, 660, 667, 680, 755, 793, 803, 816 | x | 79, 291, 299,304, 323, 329, 340, 423, 659, 667, 816 | 70, 79, 107, 165, 177, 198, 207, 276, 299, 323, 340, 367, 370, 379, 473, 522, 659, 666, 799, 816 | x | 15 | **Diversifying selection** |
| | | | | x | x | x | x | x | 26, 34, 65, 78, 82, 86, 100, 102, 109, 120, 136, 162, 176, 262, 359, 368, 380, 396, 414, 427, 441, 456, 461, 463, 465, 474, 483, 503, 508, 511, 518, 529, 532, 539, 541, 551, 556, 560, 566, 580, 584, 589, 600, 604, 630, 636, 642, 674, 684, 702, 711, 715, 733, 734, 738, 743, 747, 758, 772, 787, 801, 806, 814 | 26, 29, 33, 34, 39, 65, 72, 78, 82, 86, 96, 100, 102, 109, 119, 120, 122, 134, 136, 147, 149, 153, 162, 166, 175, 176, 186, 194, 217, 227, 244, 245, 262, 281, 337, 345, 352, 359, 363, 368, 380, 388, 389, 395, 396, 401, 408, 414, 427, 434, 441, 451, 454, 456, 457, 459, 461, 463, 465, 474, 475, 476, 480, 483, 487, 492, 503, 506, 508, 511, 515, 518, 529, 530, 532, 538, 541, 545, 547, 551, 553, 556, 560, 562, 566, 577, 580, 582, 584, 587, 589, 590, 595, 598, 600, 601, 604, 611, 619, 630, 636, 640, 642, 651, 657, 673, 674, 684, 687, 693, 698, 699, 702, 705, 708, 711, 715, 730, 731, 732, 733, 734, 737, 738, 740, 741, 743, 744, 747, 748, 751, 758, 761, 764, 767, 772, 774, 776, 779, 787, 795, 801, 805, 806, 807, 814 | x | 1, 26, 34, 65, 78, 82, 86, 88, 96, 100, 102, 109, 119, 120, 136, 147, 153, 162, 166, 175, 176, 186, 194, 245, 262, 345, 352, 359, 368, 380, 389, 396, 401, 414, 427, 434, 441, 451, 456, 461, 463, 465, 474, 480, 483, 508, 511, 518, 529, 530, 532, 541, 545, 551, 553, 556, 560, 562, 566, 580, 582, 584, 589, 590, 595, 600, 604, 630, 636, 640, 642, 674, 684, 687, 702, 708, 711, 715, 731, 732, 733, 734, 738, 743, 744, 747, 758, 772, 779, 787, 801, 805, 806, 814 | 95 | **Purifying selection** |

| Gene | No. aa residues | No. of species | ω (M0) | lmL M7 | lmL M8 | (-2lnaL) | Sgnificanc e | PAML | SLAC | FEL | MEME | FUBAR | Total no. of sites | |
|---|---|---|---|---|---|---|---|---|---|---|---|---|---|---|
| TLR2 | 2376 | 18 | 0.43041 | -14582,254597 | -14558,784112 | | | 26, 59, 101, 254, 302, 303, 334, 341, 382, 552, 607, 788 | x | 21, 39, 214, 552 | 21, 79, 99, 214, 240, 266, 365, 399, 456, 485, 521, 552, 571 | x | 3 | **Diversifying selection** |
| | | | | x | x | x | x | x | 43, 61, 63, 73, 81, 98, 107, 111, 113, 116, 132, 137, 165, 204, 227, 243, 265, 289, 292, 320, 322, 329, 345, 378, 401, 414, 427, 439, 446, 471, 517, 539, 542, 545, 546, 581, 586, 592, 611, 615, 632, 643, 645, 647, 654, 671, 679, 684, 687, 708, 717, 724, 727, 735 | 5, 10, 13, 15, 29, 43, 51, 53, 61, 63, 67, 68, 73, 78, 81, 93, 98, 107, 111, 113, 114, 115, 116, 132, 136, 137, 161, 164, 165, 174, 202, 204, 227, 231, 238, 243, 265, 270, 286, 287, 289, 292, 299, 320, 322, 329, 345, 351, 356, 376, 378, 392, 394, 401, 414, 423, 425, 427, 439, 446, 450, 452, 469, 471, 486, 491, 492, 517, 529, 539, 541, 542, 543, 545, 546, 555, 561, 564, 566, 568, 581, 583, 586, 592, 611, 615, 619, 629, 632, 643, 644, 645, 647, 648, 649, 650, 654, 662, 669, 670, 671, 679, 684, 687, 691, 692, 696, 708, 710, 711, 712, 717, 719, 721, 724, 727, 732, 735, 742, 746, 754, 755, 761, 767, 771, 778 | x | 5, 10, 43, 51, 53, 61, 63, 67, 68, 73, 78, 81, 93, 98, 107, 111, 113, 114, 115, 116, 132, 136, 137, 161, 164, 165, 174, 202, 204, 227, 231, 238, 243, 265, 270, 286, 289, 292, 299, 320, 322, 329, 345, 356, 376, 378, 392, 394, 401, 414, 427, 439, 446, 471, 486, 492, 517, 529, 539, 542, 543, 545, 546, 564, 568, 581, 583, 586, 592, 611, 615, 619, 629, 632, 643, 645, 647, 648, 649, 654, 662, 670, 671, 679, 684, 687, 691, 692, 696, 708, 710, 711, 717, 719, 721, 724, 727, 735, 742, 754, 761, 767, 771 | 102 | **Purifying selection** |

| Gene | No. aa residues | No. of species | ω (M0) | lmL M7 | lmL M8 | (-2lnaL) | Sgnificance | PAML | SLAC | FEL | MEME | FUBAR | Total no. of sites | |
|---|---|---|---|---|---|---|---|---|---|---|---|---|---|---|
| **TLR3** | **2715** | 20 | 0.30578 | -17331,254754 | -17327,255488 | | | 212, 466 | 212, 529 | 2, 68, 212, 357, 483, 529, 616, 879 | 2, 21, 66, 158, 170, 199, 209, 212, 249, 312, 357, 390, 466, 483, 529, 603 | 212, 357, 466 | 6 | **Diversifying selection** |
| | | | | x | x | x | x | x | 48, 52, 58, 67, 81, 83, 85, 103, 107, 131, 151, 163, 177, 192, 205, 220, 236, 243, 246, 256, 264, 265, 271, 298, 299, 302, 317, 345, 354, 355, 363, 368, 372, 375, 382, 394, 399, 402, 404, 413, 424, 425, 438, 446, 447, 450, 453, 454, 457, 463, 474, 477, 478, 480, 486, 488, 493, 498, 499, 500, 502, 510, 511, 512, 518, 528, 550, 551, 553, 556, 563, 572, 574, 580, 581, 583, 587, 590, 598, 604, 607, 620, 622, 623, 644, 664, 666, 678, 679, 686, 697, 701, 704, 708, 709, 732, 734, 742, 755, 759, 770, 775, 790, 792, 804, 830, 835, 845, 848, 851, 874, 897 | 5, 26, 32, 33, 34, 35, 41, 44, 46, 47, 48, 52, 58, 67, 69, 78, 80, 81, 83, 85, 90, 92, 97, 99, 102, 103, 107, 112, 116, 118, 120, 124, 126, 131, 139, 151, 159, 163, 174, 176, 177, 180, 181, 184, 192, 201, 203, 205, 220, 229, 231, 236, 239, 243, 246, 248, 251, 253, 256, 264, 265, 271, 273, 276, 279, 294, 298, 299, 300, 302, 314, 317, 318, 333, 345, 354, 355, 360, 363, 368, 372, 375, 381, 382, 387, 391, 394, 399, 400, 402, 404, 407, 408, 411, 412, 413, 419, 420, 424, 425, 426, 436, 438, 442, 446, 447, 450, 451, 453, 454, 457, 463, 474, 477, 478, 480, 484, 486, 488, 493, 494, 498, 499, 500, 501, 502, 506, 508, 510, 511, 512, 518, 520, 523, 528, 536, 543, 544, 546, 549, 550, 551, 552, 553, 555, 556, 557, 563, 566, 567, 572, 573, 574, 577, 580, 581, 583, 584, 587, 590, 593, 597, 598, 601, 604, 607, 610, 612, 615, 620, 622, 623, 624, 635, 639, 643, 644, 646, 647, 648, 649, 657, 664, 666, 668, 674, 678, 679, 686, 687, 691, 697, 700, 701, 704, 705, 708, 709, 717, 724, 726, 732, 734, 737, 738, 742, 743, 745, 746, 747, 748, 755, 756, 757, 759, 770, 773, 775, 780, 789, 790, 791, 792, 793, 794, 795, 797, 804, 807, 811, 816, 817, 819, 823, 826, 830, 833, 835, 836, 842, 845, 848, 851, 855, 859, 861, 865, 873, 874, 878, 889, 890, 893, 897, 898, 899, 904, 905 | x | 1, 5, 26, 33, 34, 35, 37, 41, 44, 47, 48, 50, 52, 58, 67, 69, 78, 80, 81, 83, 85, 90, 92, 97, 99, 102, 103, 107, 112, 118, 120, 124, 126, 131, 141, 151, 155, 159, 163, 174, 176, 177, 179, 180, 181, 184, 189, 192, 200, 203, 205, 220, 229, 231, 236, 239, 243, 246, 248, 251, 253, 256, 264, 265, 271, 273, 276, 277, 279, 280, 281, 294, 298, 300, 302, 314, 317, 318, 320, 333, 345, 346, 349, 354, 355, 360, 363, 368, 372, 375, 381, 382, 383, 387, 390, 394, 397, 399, 402, 404, 407, 408, 411, 412, 413, 417, 419, 420, 424, 425, 426, 436, 438, 440, 442, 446, 447, 450, 451, 453, 454, 457, 463, 474, 477, 478, 480, 484, 485, 486, 488, 493, 494, 498, 499, 500, 501, 502, 506, 508, 510, 511, 512, 515, 516, 518, 519, 520, 523, 525, 528, 536, 539, 542, 543, 544, 546, 549, 550, 551, 552, 553, 555, 556, 557, 563, 566, 567, 572, 573, 574, 577, 580, 581, 583, 584, 587, 590, 593, 595, 597, 598, 601, 604, 607, 610, 612, 619, 620, 622, 623, 624, 633, 635, 639, 641, 642, 643, 644, 645, 646, 647, 648, 649, 650, 652, 664, 666, 668, 674, 677, 678, 679, 681, 686, 687, 691, 693, 697, 700, 701, 704, 705, 708, 709, 716, 724, 726, 731, 732, 734, 737, 738, 742, 743, 745, 746, 747, 748, 756, 757, 759, 761, 762, 765, 770, 773, 775, 780, 788, 789, 790, 791, 792, 793, 795, 797, 799, 804, 811, 812, 816, 817, 819, 823, 826, 830, 833, 835, 836, 842, 845, 848, 851, 855, 859, 861, 865, 873, 874, 876, 878, 886, 889, 890, 893, 897, 898, 899, 904, 905 | 244 | **Purifying selection** |

| Gene | No. aa residues | No. of species | ω (M0) | lmL M7 | lmL M8 | (-2lnaL) | Sgnificance | Paml | SLAC | FEL | MEME | FUBAR | Total no. of site | |
|---|---|---|---|---|---|---|---|---|---|---|---|---|---|---|
| **TLR5** | **2736** | 15 | 0.29016 | -16787,50806 | -16784,93301 | | | 236, 237, 261, 425 | x | 41, 55, 323, 340, 348, 425, 447, 612, 634, 688, 862, 884 | 41, 52, 55, 61, 190, 193, 199, 204, 261, 317, 323, 326, 339, 340, 350, 425, 447, 451, 478, 488, 634, 708, 883 | 55, 425 | 8 | **Diversifying selection** |
| | | | | x | x | x | x | x | 13, 46, 54, 63, 66, 94, 113, 118, 124, 126, 131, 137, 138, 147, 152, 153, 159, 162, 164, 173, 186, 189, 192, 212, 227, 245, 246, 256, 260, 275, 279, 293, 300, 316, 335, 352, 354, 358, 369, 371, 374 376, 383, 390, 393, 398, 407, 410, 415, 419, 421, 424, 432, 448, 459, 460, 463, 465, 473, 474, 512, 514, 515, 519, 533, 539, 552, 571, 576, 584, 592, 593, 594, 596, 603, 604, 607, 618, 624, 638, 670, 678, 681, 682, 698, 715, 729, 731, 766, 767, 770, 771, 773, 774, 780, 785, 788, 789, 790, 796, 799, 802, 812, 815, 816, 822, 826, 827, 834, 841, 843, 853, 856, 909 | 8, 9, 10, 13, 44, 45, 46, 54, 57, 63, 66, 69, 77, 82, 86, 88, 94, 99, 100, 109, 113, 116, 117, 118, 121, 123, 124, 126, 128, 131, 137, 138, 141, 142, 145, 147, 150, 151, 152, 153, 154, 159, 161, 162, 164, 173, 175, 178, 180, 186, 188, 189, 192, 195, 212, 226, 227, 231, 232, 235, 245, 246, 254, 256, 258, 259, 260, 264, 275, 279, 281, 282, 289, 292, 293, 294, 295, 297, 299, 300, 303, 304, 306, 308, 311, 312, 316, 317, 324, 328, 330, 335, 341, 344, 352, 354, 357, 358, 359, 362, 365, 369, 371, 374, 375, 376, 381, 383, 390, 393, 394, 396, 397, 398, 400, 403, 407, 409, 410, 412, 415, 417, 419, 421, 424, 432, 441, 444, 448, 449, 459, 460, 463, 465, 466, 470, 473, 474, 490, 493, 496, 498, 501, 504, 507, 510, 511, 512, 514, 515, 516, 517, 519, 525, 526, 532, 533, 534, 537, 539, 540, 541, 542, 543, 546, 548, 551, 552, 553, 558, 559, 562, 564, 566, 567, 568, 571, 576, 578, 581, 584, 587, 588, 589, 592, 593, 594, 596, 597, 599, 602, 603, 604, 607, 614, 615, 617, 618, 624, 628, 631, 638, 644, 647, 650, 658, 660, 663, 670, 673, 674, 678, 681, 682, 684,687, 691, 692, 698, 706, 715, 719, 725, 729, 731, 732, 733, 734, 737, 745, 747, 749, 757, 760, 762, 766, 767, 770, 771, 772, 773, 774, 775, 776, 779, 780, 781, 785, 787, 788, 789, 790, 792, 796, 799, 800, 802, 805, 808, 809, 812, 813, 815, 816, 818, 822, 824, 826, 827, 833, 834, 841, 842, 843, 853, 855, 856, 858, 867, 872, 873, 898, 899, 904, 908, 909, 911 | x | 3, 9, 10, 13, 44, 45, 46, 54, 63, 66, 77, 86, 87, 88, 94, 99, 100, 105, 113, 114, 116, 117, 118, 119, 123, 124, 126, 127, 128, 131, 137, 138, 140, 141, 147, 150, 152, 153, 154, 159, 161, 162, 164, 172, 173, 175, 177, 178, 186, 188, 189, 192, 200, 203, 209, 212, 221, 226, 227, 231, 232, 235, 245, 246, 247, 248, 252, 254, 256, 258, 259, 260, 264, 272, 275, 279, 281, 289, 292, 293, 295, 296, 297, 300, 303, 304, 308, 311, 312, 315, 316, 328, 330, 335, 341, 344, 352, 354, 358, 359, 362, 365, 369, 371, 373, 374, 375, 376, 379, 381, 383, 390, 393, 394, 396, 397, 398, 400, 403, 407, 409, 410, 412, 415, 417, 419, 421, 424, 432, 433, 441, 444, 445, 448, 449, 459, 460, 463, 465, 466, 470, 473, 474, 477, 490, 493, 496, 498, 504, 507, 511, 512, 514, 515, 516, 517, 519, 525, 532, 533, 534, 537, 539, 541, 542, 546, 548, 551, 552, 553, 558, 562, 564, 566, 567, 568, 571, 576, 578, 580, 581, 584, 587, 588, 589, 592, 593, 594, 596, 597, 599, 602, 603, 604, 607, 611, 614, 616, 617, 618, 619, 624, 631, 638, 644, 647, 658, 663, 670, 673, 674, 675, 678, 681, 682, 692, 698, 706, 715, 719, 725, 729, 731, 732, 733, 734, 737, 745, 747, 749, 760, 762, 766, 767, 770, 771, 772, 773, 774, 775, 776, 778, 779, 780, 781, 782, 785, 787, 788, 789, 790, 792, 796, 799, 800, 802, 804, 805, 807, 808, 809, 811, 812, 813, 815, 816, 818, 822, 824, 826, 827, 830, 833, 834, 841, 842, 843, 847, 848, 853, 856, 858, 867, 873, 897, 899, 904, 906, 909, 911 | 251 | **Purifying selection** |

| Gene | No. aa residues | No. of species | ω (M0) | lmL M7 | lmL M8 | (-2lnaL) | Sgnificance | PAML | SLAC | FEL | MEME | FUBAR | Total no. of sites | |
|---|---|---|---|---|---|---|---|---|---|---|---|---|---|---|
| **TLR7** | 3234 | 21 | 0.16063 | -23862, 031766 | -23854, 542755 | | | x | x | 335, 738 | 14, 41, 42, 53, 335, 375, 505, 508, 699, 705, 714, 738, 747, 827, 846, 1043 | x | 2 | **Diversifying selection** |
| | | | | x | x | x | x | x | 35, 36, 37, 49, 50, 52, 57, 58, 61, 62, 64, 73, 78, 82, 83, 84, 85, 89, 90, 92, 93, 95, 96, 98, 100, 104, 106, 107, 126, 127, 128, 129, 130, 131, 133, 135, 137, 141, 145, 149, 152, 154, 161, 164, 172, 176, 180, 191, 194, 196, 199, 208, 210, 211, 213, 215, 216, 222, 232, 243, 249, 250, 252, 256, 257, 258, 260, 263, 264, 265, 268, 271, 274, 279, 284, 290, 294, 295, 297, 299, 306, 311, 319, 320, 323, 324, 330, 333, 334, 336, 340, 343, 344, 347, 349, 350, 351, 352, 354, 359, 364, 367, 369, 370, 371, 373, 374, 376, 378, 379, 380, 390, 400, 402, 404, 406, 408, 427, 429, 430, 432, 433, 435, 446, 463, 464, 467, 473, 475, 476, 478, 483, 496, 511, 528, 531, 532, 534, 542, 545, 549, 552, 566, 568, 571, 575, 581, 582, 585, 587, 590, 591, 595, 597, 608, 611, 617, 622, 623, 624, 631, 635, 638, 648, 657, 666, 667, 668, 669, 671, 675, 681, 685, 686, 726, 727, 749, 764, 765, 779, 795, 804, 806, 809, 811, 813, 814, 819, 824, 826, 831, 835, 839, 842, 849, 858, 867, 879, 882, 899, 904, 908, 909, 911, 922, 923, 924, 927, 928, 945, 949, 951, 955, 956, 957, 958, 960, 961, 964, 965, 968, 970, 971, 973, 976, 980, 981, 982, 990, 997, 1003, 1009, 1013, 1023, 1029, 1031, 1035, 1042, 1047, 1049, 1050, 1051, 1053, 1059, 1066, 1075 | 34, 35, 36, 37, 44, 45, 48, 49, 50, 51, 52, 57, 58, 59, 61, 62, 64, 66, 67, 69, 70, 71, 72, 73, 78, 82, 83, 84, 85, 86, 89, 90, 92, 93, 94, 95, 96, 97, 98, 100, 104, 106, 107, 111, 114, 115, 120, 122, 123, 124, 126, 127, 128, 129, 130, 131, 132, 133, 134, 135, 136, 137, 141, 143, 145, 148, 149, 151, 152, 154, 156, 158, 161, 164, 165, 167, 171, 172, 175, 176, 178, 179, 180, 182, 183, 186, 190, 191, 193, 194, 196, 198, 199, 200, 203, 204, 207, 208, 210, 211, 212, 213, 214, 215, 216, 218, 221, 222, 224, 231, 232, 233, 235, 242, 243, 244, 249, 250, 252, 253, 255, 256, 257, 258, 260, 261, 263, 264, 265, 268, 269, 271, 274, 275, 279, 284, 287, 289, 290, 294, 295, 297, 299, 300, 303, 305, 306, 311, 318, 319, 320, 323, 324, 326, 330, 331, 333, 334, 340, 342, 343, 344, 347, 349, 350, 351, 354, 359, 360, 364, 367, 369, 370, 371, 373, 374, 376, 378, 379, 380, 390, 391, 400, 401, 402, 404, 406, 408, 411, 412, 426, 427, 428, 429, 430, 432, 434, 435, 445, 446, 449, 461, 463, 464, 465, 466, 467, 472, 473, 475, 476, 478, 480, 481, 483, 496, 500, 504, 511, 512, 513, 515, 517, 519, 521, 527, 528, 529, 531, 532, 533, 534, 535, 536, 537, 538, 539, 542, 547, 548, 549, 552, 555, 557, 558, 559, 562, 563, 565, 566, 567, 568, 569, 571, 572, 575, 581, 582, 583, 585, 587, 589, 590, 591, 594, 595, 597, 598, 600, 601, 602, 603, 605, 607, 608, 609, 611, 612, 613, 615, 617, 619, 621, 622, 623, 624, 631, 632, 634, 635, 636, 638, 639, 643, 647, 648, 653, 657, 658, 660, 662, 666, 667, 668, 669, 671, 673, 675, 681, 684, 685, 686, 689, 691, 694, 696, 698, 702, 704, 706, 709, 717, 722, 724, 726, 727, 736, 737, 744, 749, 756, 760, 761, 763, 764, 765, 768, 771, 776, 777, 779, 781, 792, 795, 804, 806, 809, 811, 812, 813, 814, 815, 819, 824, 826, 831, 832, 833, 834, 835, 836, 837, 838, 839, 841, 842, 844, 846, 849, 850, 853, 857, 858, 863, 867, 871, 876, 879, 882, 884, 891, 892, 894, 896, 899, 904, 907, 908, 909, 911, 920, 922, 923, 924, 926, 927, 928, 941, 942, 945, 948, 949, 950, 951, 952, 955, 956, 957, 958, 960, 961, 964, 965, 966, 968, 970, 971, 972, 973, 975, 976, 980, 981, 982, 984, 990, 991, 993, 996, 997, 999, 1003, 1006, 1007, 1009, 1013, 1015, 1022, 1023, 1029, 1030, 1031, 1033, 1035, 1038, 1040, 1041, 1042, 1047, 1048, 1049, 1050, 1051, 1053, 1059, 1060, 1062, 1066, 1069, 1075 | x | 2, 8, 29, 33, 34, 35, 36, 37, 40, 44, 45, 48, 49, 50, 51, 52, 57, 58, 59, 60, 61, 62, 64, 66, 67, 69, 70, 71, 72, 73, 78, 82, 83, 84, 85, 86, 89, 90, 92, 93, 94, 95, 96, 97, 98, 100, 104, 105, 106, 107, 111, 113, 114, 115, 116, 119, 120, 122, 123, 124, 125, 126, 127, 128, 129, 130, 131, 132, 133, 134, 135, 136, 137, 141, 143, 145, 148, 149, 151, 152, 154, 155, 156, 158, 160, 161, 164, 165, 167, 171, 172, 175, 176, 177, 178, 179, 180, 182, 183, 184, 186, 187, 190, 191, 193, 194, 196, 197, 198, 199, 203, 204, 207, 208, 210, 211, 212, 213, 214, 215, 216, 217, 218, 219, 221, 222, 224, 225, 230, 231, 232, 233, 235, 242, 243, 244, 246, 247, 249, 250, 252, 253, 254, 255, 256, 257, 258, 259, 260, 261, 263, 264, 265, 266, 268, 269, 271, 274, 275, 276, 277, 279, 283, 284, 287, 289, 290, 292, 294, 295, 296, 297, 299, 300, 301, 302, 303, 305, 306, 307, 310, 311, 317, 318, 319, 320, 323, 324, 326, 328, 330, 331, 333, 334, 336, 337, 340, 342, 343, 344, 347, 349, 350, 351, 352, 354, 359, 360, 364, 365, 366, 367, 369, 370, 371, 372, 373, 374, 376, 377, 378, 379, 380, 381, 388, 390, 391, 394, 396, 397, 399, 400, 401, 402, 403, 404, 406, 408, 410, 411, 412, 418, 422, 423, 426, 427, 428, 429, 430, 432, 433, 435, 441, 445, 446, 449, 450, 457, 458, 459, 460, 461, 463, 464, 465, 466, 467, 472, 473, 475, 476, 478, 480, 481, 482, 483, 487, 496, 500, 502, 504, 507, 511, 512, 513, 515, 517, 519, 521, 527, 528, 529, 531, 532, 533, 534, 535, 536, 537, 538, 539, 542, 543, 545, 546, 547, 548, 549, 552, 555, 556, 557, 558, 559, 560, 563, 564, 565, 566, 567, 568, 569, 571, 572, 575, 577, 579, 581, 582, 583, 584, 585, 587, 589, 590, 591, 594, 595, 596, 597, 598, 600, 602, 603, 605, 607, 608, 609, 611, 612, 615, 617, 618, 619, 621, 622, 623, 624, 631, 632, 635, 636, 638, 639, 642, 645, 646, 647, 648, 649, 653, 654, 657, 658, 660, 662, 665, 666, 667, 668, 669, 670, 671, 673, 675, 676, 677, 679, 680, 681, 684, 685, 686, 689, 691, 694, 696, 697, 698, 703, 706, 707, 709, 717, 722, 724, 726, 727, 735, 736, 737, 741, 744, 749, 755, 756, 760, 761, 763, 764, 765, 768, 771, 773, 776, 777, 779, 781, 792, 794, 795, 799, 804, 806, 809, 811, 812, 813, 814, 815, 818, 819, 821, 822, 823, 824, 826, 831, 832, 833, 834, 835, 838, 839, 840, 841, 842, 843, 844, 846, 849, 853, 857, 858, 863, 867, 871, 876, 879, 882, 883, 884, 886, 890, 891, 892, 893, 894, 896, 899, 901, 904, 905, 907, 908, 909, 911, 914, 920, 921, 922, 923, 924, 925, 926, 927, 928, 933, 934, 936, 937, 938, 941, 945, 946, 948, 949, 950, 951, 952, 955, 956, 957, 958, 960, 961, 962, 964, 965, 966, 967, 968, 970, 971, 972, 973, 975, 976, 980, 981, 982, 984, 987, 989, 990, 991, 993, 994, 995, 996, 997, 999, 1003, 1006, 1007, 1009, 1012, 1013, 1015, 1019, 1022, 1023, 1025, 1027, 1028, 1029, 1030, 1031, 1033, 1035, 1038, 1040, 1041, 1042, 1044, 1045, 1047, 1048, 1049, 1050, 1051, 1053, 1055, 1057, 1059, 1060, 1062, 1066, 1069, 1070, 1075 | 450 | **Purifying selection** |

| Gene | No. aa residues | No. of species | ω (M0) | lmL M7 | lmL M8 | (-2lnaL) | Sgnificance | Paml >95% | SLAC | FEL | MEME | FUBAR | Total no. of site | |
|---|---|---|---|---|---|---|---|---|---|---|---|---|---|---|
| **TLR8** | **3165** | 18 | 0.41738 | -19003,057429 | -18967,837654 | | | 3, 5, 6, 9, 24,39, 40, 41, 42, 53, 57, 58,61, 74, 76, 78, 83, 87, 92, 98, 100, 101, 103, 104, 106, 107,108, 111, 112, 116, 126,139, 140, 147, 150, 186, 191, 196, 203, 204, 208, 212, 213, 222, 225, 226, 231, 233, 234, 242, 247,249, 276, 277, 288, 290, 294, 301, 303, 320, 322, 328, 354, 355, 357, 359, 361, 364, 376, 387, 388, 395, 397, 419, 466, 470, 472, 482, 485, 486, 487, 490, 492, 493, 494,496, 530, 543609, 634, 641, 644, 648, 663, 665, 667, 668,671, 679, 690, 692, 693, 695, 697, 700, 712, 721,736, 738, 740, 764, 765, 767,769, 776, 778, 819, 845, 856, 881, 884, 886, 888, 929, 940, 955, 1029, 1044, 1052 | 140, 671 | 108, 139, 140,203,327, 609, 641, 648, 671, 679, 736, 769, 778, 845 | 12, 39, 61, 86, 108, 139, 140, 158, 176, 186, 246, 249, 394, 461, 570,609, 618, 648, 671, 740, 769, 775, 778, 845, 884, 886 | 139 | 20 | **Diversifying selection** |
| | | | | x | x | x | x | x | 47, 51, 60, 62, 75, 99, 142, 188, 197, 259, 260, 262, 268, 281, 307, 319, 326, 334, 365, 373, 379, 405, 406, 410, 446, 498, 499, 507, 509, 532, 534, 544, 567, 568, 578, 581, 583, 591, 594, 595, 600, 617, 624, 643, 662, 664, 669, 711, 719, 727, 743, 745, 746, 751, 759, 771, 781, 804, 853, 864, 870, 900, 903, 906, 912, 938, 946, 948, 952, 953, 954, 975, 1018, 1037 | 8, 25, 30, 31, 35, 36, 46, 47, 51, 59, 60, 62, 69, 75, 77, 79, 88, 90, 96, 99,121, 123, 131, 136,138, 142, 145, 153, 161, 168, 181, 184, 188, 190, 195, 197, 199, 210, 214, 217, 224, 230, 237, 259, 260, 262, 265, 267, 268, 269, 272, 274, 278, 281, 283, 302, 307, 318, 319, 326, 333, 334, 339, 342, 351, 356, 362, 365, 373, 379, 380, 384, 396, 400, 402, 405, 406, 410, 411, 416, 428, 430, 432, 439, 446, 458, 462, 498, 507, 509, 512, 516, 532, 534, 536, 537, 539, 544, 546, 554, 556, 563, 567, 568, 572, 574, 578, 581, 583, 591, 594, 595, 600, 608, 613, 614,617, 624, 628, 643, 645, 659, 662, 664, 666, 669, 676, 678, 681, 687, 691, 699, 707, 709, 711, 713, 719, 722, 723, 727, 728, 733, 734, 739, 743, 745, 746, 748, 751, 752, 757, 759, 771, 781, 788, 800, 804, 805, 813, 814, 815, 818, 826, 833, 837, 853, 857, 864, 865, 868, 870, 898, 899, 900, 903, 906, 907, 912, 921, 925, 926, 932, 933, 938, 941, 942, 946, 948, 950, 952, 953, 954, 964, 965, 967, 972, 975, 984, 986, 990, 991, 992, 998, 1002, 1006, 1009, 1012, 1017, 1018, 1019, 1022, 1034, 1037, 1052, 1053 | | 25, 30, 31, 35, 36, 46, 47, 51, 60, 62, 69, 75, 77, 99, 121, 123, 138, 142, 145, 168, 184, 188, 197, 199, 214, 217, 224, 230, 255, 259, 260, 262, 265, 267, 268, 269, 272, 274, 278, 281, 283, 307, 319, 326, 333, 334, 339, 348, 356, 362, 363, 365, 373, 379, 380, 396, 400, 405, 406, 410, 411, 416, 428, 430, 432, 439, 446, 458, 495, 498, 499, 507, 509, 512, 516, 532, 534, 536, 539, 544, 552, 554, 563, 567, 568, 572, 574, 578, 583, 591, 594, 595, 600, 617, 624, 643, 659, 662, 664, 669, 678, 681, 687, 691, 699, 707, 709, 711, 713, 719, 722, 723, 727, 733, 734, 739, 743, 745, 746, 748, 751, 752, 757, 759, 771, 781, 787, 793,800, 805, 813, 814, 815, 818, 828, 833, 837, 847, 857, 864, 865, 868, 870, 898, 899, 900, 903, 906, 907, 912, 925, 926, 933, 935, 938, 941, 942, 946, 948, 950, 952, 953, 954, 964, 965, 972, 975, 984, 986, 990, 991, 992, 998, 1002, 1006, 1009, 1012, 1017, 1018, 1019, 1022, 1034, 1037, 1046 | 175 | **Purifying selection** |

| Gene | No. aa residues | No. of species | ω (M0) | lmL M7 | lmL M8 | (-2lnaL) | Sgnificance | Paml >95% | SLAC | FEL | MEME | FUBAR | Total no. of site | |
|---|---|---|---|---|---|---|---|---|---|---|---|---|---|---|
| **TLR9** | **3162** | 23 | 0.33618 | -23168,548096 | -23148,160434 | | | 33, 194, 281, 315, 341, 387, 449, 547, 746, 765, 836 | x | 194, 236, 428, 444, 473, 714, 836, 1054 | 4, 21, 36, 37, 38, 112, 123, 135, 189, 194, 220, 291, 304, 305, 327, 335, 426, 428, 444, 456, 473, 547, 553, 582, 603, 701, 706, 714, 717, 738, 767, 792, 873, 1046, 1050, 1054 | x | 8 | **Diversifying selection** |
| | | | | | | | | x | 13, 30, 48, 54, 75, 90, 92, 96, 104, 106, 114, 128, 134, 147, 160, 169, 170, 176, 196, 206, 209, 210, 211, 223, 233, 244, 249, 250, 251, 254, 257, 258, 270, 273, 282, 288, 289, 309, 313, 321, 325, 329, 338, 349, 357, 364, 366, 369, 377, 389, 391, 395, 396, 413, 417, 418, 498, 509, 510, 518, 522, 523, 525, 549, 556, 564, 565, 572, 574, 587, 591, 594, 596, 597, 600, 604, 616, 626, 629, 639, 641, 644, 646, 652, 654, 662, 664, 673, 681, 683, 688, 694, 697, 699, 713, 716, 721, 736, 741, 743, 747, 754, 757, 771, 773, 774, 775, 779, 788, 789, 790, 798, 802, 816, 827, 833, 841, 843, 844, 849, 857, 859, 861, 864, 878, 879, 880, 887, 891, 892, 893, 896, 898, 899, 902, 914, 915, 922, 923, 932, 934, 940, 951, 955, 962, 967, 971, 980, 987, 998, 999, 1005, 1014, 1015, 1035, 1039 | 13, 25, 29, 30, 41, 48, 54, 58, 75, 81, 90, 92, 93, 95, 96, 102, 104, 106, 114, 117, 122, 128, 133, 134, 137, 142, 147, 149, 153, 160, 165, 169, 170, 172, 175, 176, 196, 206, 207, 209, 210, 211, 213, 219, 223, 224, 227, 228, 233, 244, 249, 250, 251, 254, 255, 257, 258, 259, 260, 265, 270, 273, 282, 288, 289, 292, 300, 306, 307, 309, 313, 321, 325, 326, 329, 338, 349, 350, 357, 358, 364, 365, 366, 369, 372, 373, 377, 380, 385, 388, 389, 391, 394, 395, 396, 401, 403, 409, 413, 415, 416, 417, 418, 421, 422, 423, 455, 460, 469, 480, 487, 488, 491, 498, 501, 503, 504, 509, 510, 518, 521, 522, 523, 525, 532, 534, 540, 541, 542, 545, 549, 550, 551, 556, 564, 565, 572, 574, 587, 589, 591, 592, 594, 596, 597, 598, 600, 604, 609, 610, 612, 616, 619, 624, 626, 629, 639, 641, 644, 646, 648, 652, 654, 662, 664, 669, 673, 675, 679, 681, 683, 686, 688, 689, 694, 695, 697, 698, 699, 703, 713, 716, 721, 728, 732, 736, 741, 742, 743, 747, 750, 751, 754, 757, 759, 766, 768, 771, 772, 773, 774, 775, 776, 779, 788, 789, 790, 793, 796, 798, 801, 802, 805, 806, 807, 810, 815, 816, 818, 821, 823, 825, 827, 833, 841, 843, 844, 846, 848, 849, 856, 857, 859, 861, 862, 863, 864, 867, 871, 878, 879, 880, 881, 887, 891, 892, 893, 895, 896, 897, 898, 899, 900, 901, 902, 904, 907, 913, 914, 915, 922, 923, 931, 932, 933, 934, 940, 942, 951, 953, 955, 957, 962, 966, 967, 971, 975, 978, 980, 985, 987, 990, 993, 994, 996, 998, 999, 1003, 1005, 1006, 1014, 1015, 1022, 1033, 1035, 1039 | x | 11, 12, 13, 25, 29, 30, 32, 41, 44, 47, 48, 54, 56, 58, 60, 63, 65, 69, 75, 81, 87, 90, 92, 93, 95, 96, 98, 102, 104, 106, 114, 117, 122, 125, 126, 127, 128, 133, 134, 137, 142, 145, 146, 147, 153, 159, 160, 165, 167, 169, 170, 172, 174, 175, 176, 190, 196, 201, 206, 207, 209, 210, 211, 213, 219, 222, 223, 224, 227, 228, 232, 233, 244, 246, 247, 249, 250, 251, 252, 254, 255, 257, 258, 259, 260, 264, 265, 270, 273, 276, 282, 285, 287, 288, 289, 292, 300, 306, 307, 309, 310, 313, 321, 325, 326, 329, 337, 338, 349, 350, 357, 358, 363, 364, 365, 366, 368, 369, 372, 373, 377, 380, 385, 388, 389, 391, 392, 393, 394, 395, 396, 403, 409, 413, 415, 416, 417, 418, 421, 422, 423, 455, 487, 488, 491, 492, 498, 501, 503, 504, 505, 509, 510, 518, 521, 522, 523, 525, 526, 530, 532, 533, 534, 540, 541, 542, 543, 545, 549, 550, 551, 555, 556, 557, 560, 564, 565, 570, 572, 573, 574, 579, 586, 587, 589, 591, 592, 594, 596, 597, 600, 601, 604, 606, 609, 610, 612, 614, 616, 619, 622, 624, 626, 629, 634, 639, 641, 644, 646, 648, 652, 653, 654, 662, 664, 669, 673, 675, 679, 681, 683, 686, 688, 689, 692, 693, 694, 695, 697, 698, 699, 703, 713, 715, 716, 721, 723, 728, 731, 732, 736, 741, 742, 743, 745, 747, 750, 754, 757, 759, 766, 768, 771, 772, 773, 774, 775, 776, 777, 779, 785, 788, 789, 790, 793, 796, 797, 798, 801, 802, 805, 806, 807, 810, 815, 816, 818, 821, 823, 826, 827, 833, 834, 841, 843, 844, 846, 848, 849, 851, 856, 857, 859, 861, 862, 863, 864, 867, 871, 878, 879, 880, 881, 884, 885, 887, 891, 892, 893, 895, 896, 898, 899, 900, 901, 902, 904, 907, 913, 914, 915, 916, 922, 923, 925, 931, 932, 933, 934, 937, 940, 942, 951, 953, 955, 957, 958, 962, 966, 967, 971, 973, 975, 978, 980, 983, 985, 987, 990, 991, 993, 994, 996, 998, 999, 1003, 1005, 1006, 1009, 1010, 1012, 1014, 1022, 1029, 1032, 1033, 1035, 1039, 1040, 1045 | 288 | **Purifying selection** |

| Gene | No. aa residues | No. of species | lmL M7 | ω (M0) | lmL M8 | (-2lnaL) | Sgnificance | Paml >95% | SLAC | FEL | MEME | FUBAR | Total no. of site | |
|---|---|---|---|---|---|---|---|---|---|---|---|---|---|---|
| **TLR21** | **3453** | 14 | -20272,219816 | 0.15856 | -20271,638116 | | | 222 | x | 213, 235, 263, 307, 321, 326, 376, 990 | 78, 96, 146, 213, 218, 235, 263, 307, 321, 332, 608, 676, 694, 752, 819, 832, 911, 921 | x | 5 | **Diversifying selection** |
| | | | x | | x | x | x | x | 28, 46, 62, 83, 97, 105, 109, 123, 147, 152, 160, 164, 169, 171, 175, 186, 187, 200, 208, 211, 221, 231, 238, 240, 261, 268, 269, 273, 280, 284, 288, 293, 301, 304, 305, 309, 314, 317, 319, 331, 337, 344, 351, 355, 360, 364, 366, 585, 611, 617, 624, 634, 635, 639, 641, 651, 655, 671, 672, 684, 695, 698, 699, 705, 708, 712, 723, 726, 728, 730, 734, 737, 747, 754, 758, 762, 764, 767, 769, 770, 771, 772, 774, 787, 789, 790, 792, 799, 801, 805, 806, 812, 813, 815, 816, 823, 837, 838, 850, 858, 862, 872, 876, 877, 881, 883, 885, 888, 891, 912, 919, 929, 930, 933, 934, 939, 943, 948, 969, 970, 972, 975, 976, 984, 993, 994, 995, 996, 998, 999, 1001, 1003, 1004, 1007, 1008, 1012, 1013, 1019, 1027, 1031, 1033, 1034, 1037, 1042, 1043, 1052, 1055, 1064, 1065, 1071, 1075, 1078, 1080, 1081, 1082, 1084, 1095, 1098, 1100, 1102, 1113, 1114, 1118, 1125, 1127, 1130, 1134 | 6, 7, 28, 30, 42, 45, 46, 47, 48, 49, 58, 60, 62, 67, 73, 81, 83, 87, 89, 90, 93, 95, 97, 99, 102, 105, 109, 110, 116, 117, 118, 123, 128, 130, 133, 134, 137, 140, 145, 147, 149, 152, 153, 155, 156, 160, 161, 162, 164, 167, 169, 171, 173, 174, 175, 178, 185, 186, 187, 195, 196, 197, 200, 207, 208, 209, 210, 211, 214, 216, 217, 220, 221, 224, 225, 230, 231, 237, 238, 240, 241, 247, 250, 254, 256, 257, 258, 260, 261, 268, 269, 272, 273, 274, 280, 284, 287, 288, 289, 291, 293, 297, 300, 301, 303, 304, 305, 308, 311, 312, 314, 317, 319, 330, 331, 337, 338, 339, 341, 344, 351, 355, 358, 360, 361, 362, 363, 364, 366, 367, 368, 371, 374, 377, 389, 473, 481, 489, 551, 561, 576, 579, 580, 582, 585, 588, 590, 591, 592, 595, 599, 600, 603, 609, 611, 613, 617, 619, 624, 630, 634, 635, 638, 639, 640, 641, 651, 654, 655, 658, 659, 660, 662, 663, 665, 666, 671, 672, 675, 676, 678, 679, 681, 682, 684, 685, 687, 688, 689, 690, 691, 695, 698, 699, 701, 705, 706, 707, 708, 709, 710, 712, 714, 718, 719, 720, 723, 726, 727, 728, 729, 730, 731, 733, 734, 737, 739, 740, 741, 747, 748, 751, 753, 754, 757, 758, 759, 762, 763, 764, 765, 766, 767, 769, 770, 771, 772, 774, 778, 781, 784, 785, 787, 789, 790, 792, 799, 801, 802, 804, 805, 806, 808, 811, 812, 813, 815, 816, 818, 820, 822, 823, 825, 828, 834, 835, 836, 837, 838, 839, 840, 841, 845, 847, 850, 852, 857, 858, 860, 861, 862, 863, 864, 865, 869, 870, 871, 872, 874, 876, 877, 881, 883, 884, 885, 886, 887, 888, 891, 893, 894, 896, 898, 902, 903, 908, 909, 910, 912, 913, 918, 919, 922, 925, 926, 927, 928, 929, 930, 932, 933, 934, 935, 937, 939, 943, 948, 954, 957, 960, 963, 966, 969, 970, 972, 975, 976, 978, 982, 983, 984, 988, 993, 994, 995, 996, 998, 999, 1000, 1001, 1003, 1004, 1007, 1008, 1012, 1013, 1016, 1019, 1022, 1024, 1025, 1026, 1027, 1028, 1029, 1031, 1032, 1033, 1034, 1036, 1037, 1038, 1040, 1041, 1042, 1043, 1047, 1052, 1053, 1055, 1056, 1061, 1062, 1064, 1065, 1068, 1070, 1071, 1073, 1074, 1075, 1076, 1077, 1078, 1079, 1080, 1081, 1082, 1083, 1084, 1087, 1088, 1091, 1094, 1095, 1097, 1098, 1099, 1100, 1102, 1103, 1106, 1107, 1108, 1113, 1114, 1118, 1120, 1124, 1125, 1126, 1127, 1129, 1130, 1132, 1134, 1135, 1137, 1139, 1142, 1145, 1147 | x | 6, 7, 22, 26, 28, 30, 42, 45, 46, 47, 48, 49, 55, 57, 58, 60, 61, 62, 67, 73, 81, 83, 85, 87, 89, 90, 93, 95, 97, 102, 104, 105, 108, 109, 116, 117, 118, 119, 123, 125, 128, 133, 134, 137, 140, 143, 145, 147, 148, 149, 151, 152, 153, 155, 156, 158, 160, 161, 162, 164, 167, 169, 171, 173, 174, 175, 178, 184, 185, 186, 187, 195, 196, 197, 200, 206, 208, 209, 210, 211, 214, 217, 221, 224, 225, 229, 230, 231, 236, 237, 238, 240, 243, 247, 254, 256, 257, 258, 261, 268, 269, 271, 272, 273, 274, 275, 280, 284, 287, 288, 289, 291, 293, 297, 300, 301, 302, 304, 305, 306, 308, 309, 311, 314, 317, 319, 330, 331, 336, 337, 338, 339, 341, 344, 345, 346, 350, 351, 352, 355, 358, 360, 361, 362, 364, 366, 367, 371, 374, 377, 389, 391, 473, 481, 489, 547, 551, 559, 561, 576, 580, 585, 588, 590, 591, 592, 595, 596, 599, 600, 603, 609, 611, 612, 616, 617, 622, 623, 624, 626, 630, 634, 635, 638, 639, 641, 651, 654, 655, 659, 660, 662, 663, 665, 671, 672, 675, 681, 682, 684, 685, 687, 688, 689, 690, 693, 695, 698, 699, 703, 705, 706, 708, 709, 710, 712, 714, 718, 719, 720, 721, 723, 725, 726, 727, 728, 729, 730, 733, 734, 735, 737, 740, 741, 747, 748, 753, 754, 757, 758, 761, 762, 763, 764, 765, 766, 767, 769, 770, 771, 772, 773, 774, 777, 778, 779, 781, 784, 787, 789, 790, 791, 792, 796, 799, 801, 802, 805, 806, 808, 810, 812, 813, 815, 816, 818, 820, 822, 823, 825, 828, 829, 830, 834, 835, 836, 837, 838, 839, 840, 842, 845, 846, 850, 852, 857, 858, 860, 861, 862, 863, 864, 865, 867, 869, 870, 871, 872, 874, 876, 877, 881, 883, 884, 885, 886, 887, 888, 891, 893, 894, 896, 898, 899, 901, 902, 903, 908, 909, 910, 912, 918, 919, 922, 927, 929, 930, 931, 932, 933, 934, 935, 937, 939, 941, 943, 946, 947, 948, 954, 955, 957, 958, 960, 963, 965, 966, 969, 970, 972, 975, 976, 977, 982, 983, 984, 988, 993, 994, 995, 996, 998, 999, 1000, 1001, 1003, 1004, 1006, 1007, 1008, 1012, 1013, 1016, 1019, 1022, 1024, 1025, 1026, 1027, 1028, 1029, 1031, 1032, 1033, 1034, 1036, 1037, 1038, 1041, 1042, 1043, 1047, 1051, 1052, 1053, 1055, 1056, 1061, 1062, 1064, 1065, 1068, 1069, 1070, 1071, 1074, 1075, 1076, 1077, 1078, 1079, 1080, 1081, 1082, 1083, 1084, 1085, 1087, 1088, 1091, 1095, 1097, 1098, 1099, 1100, 1102, 1103, 1106, 1108, 1113, 1114, 1118, 1120, 1122, 1123, 1124, 1125, 1127, 1129, 1130, 1132, 1134, 1135, 1137, 1147, 1151 | 384 | **Purifying selection** |

| Gene | No. aa residues | No. of species | ω (M0) | lmL M7 | lmL M8 | (-2lnaL) | Sgnificance | Paml >95% | SLAC | FEL | MEME | FUBAR | Total no. of site | |
|---|---|---|---|---|---|---|---|---|---|---|---|---|---|---|
| **TLR22** | **2820** | 15 | 0.35374 | -15271,495892 | -15270,975822 | | | x | x | 121, 353, 739 | 87, 110, 159, 202, 261, 340, 353, 489, 562, 575, 739, 870, 884, 899 | x | 2 | **Diversifying selection** |
| | | | | x | x | x | x | x | 21, 32, 65, 95, 150, 156, 164, 236, 264, 275, 276, 279, 290, 297, 320, 365, 368, 370, 371, 389, 404, 408, 434, 436, 437, 448, 454, 478, 481, 509, 542, 546, 560, 563, 576, 588, 601, 658, 660, 686, 703, 712, 714, 770, 797, 802, 826, 843, 844, 854, 892 | 8, 9, 21, 32, 33, 41, 43, 44, 65, 67, 72, 85, 91, 95, 98, 99, 109, 114, 118, 122, 130, 141, 143, 150, 154, 156, 160, 164, 181, 190, 192, 197, 210, 213, 215, 221, 226, 232, 236, 239, 243, 264, 275, 276, 279, 285, 290, 292, 297, 320, 333, 360, 365, 366, 368, 370, 371, 374, 376, 383, 385, 387, 389, 392, 393, 396, 400, 404, 408, 412, 426, 431, 433, 434, 436, 437, 439, 448, 454, 457, 460, 463, 464, 466, 471, 477, 478, 481, 498, 499, 500, 509, 513, 531, 533, 542, 546, 548, 549, 550, 551, 554, 560, 563, 567, 568, 576, 580, 581, 588, 601, 607, 629, 632, 657, 658, 660, 669, 674, 676, 685, 686, 696, 703, 709, 712, 714, 718, 757, 765, 768, 770, 774, 776, 777, 780, 782, 797, 799, 802, 803, 808, 814, 817, 826, 832, 837, 840, 843, 844, 851, 852, 854, 855, 856, 857, 863, 864, 872, 874, 875, 876, 886, 888, 890, 892, 896, 902, 913, 914, 939 | x | 8, 9, 21, 31, 32, 33, 41, 43, 44, 64, 65, 67, 72, 85, 91, 95, 98, 99, 109, 114, 115, 122, 130, 141, 143, 150, 154, 156, 160, 164, 181, 190, 192, 197, 210, 213, 215, 221, 226, 232, 236, 238, 239, 243, 264, 275, 276, 279, 285, 289, 290, 292, 297, 320, 333, 360, 365, 366, 368, 370, 371, 374, 376, 383, 387, 389, 393, 396, 400, 404, 408, 412, 426, 431, 433, 434, 436, 437, 439, 448, 454, 457, 463, 464, 471, 477, 478, 481, 495, 498, 499, 500, 509, 513, 517, 531, 533, 542, 546, 548, 549, 550, 551, 554, 560, 563, 567, 568, 576, 580, 581, 588, 601, 607, 629, 630, 632, 657, 658, 660, 669, 676, 680, 685, 686, 696, 703, 709, 712, 714, 718, 719, 730, 735, 738,757, 765, 768, 770, 774, 776, 777, 780, 782, 797, 799, 802, 803, 808, 814, 817, 826, 832, 840, 843, 844, 851, 852, 854, 855, 856, 857, 863, 864, 872, 874, 875, 876, 886, 888, 890, 892, 896, 902, 913, 914 | 163 | **Purifying selection** |

| Gene | No. aa residues | No. of species | ω (M0) | lmL M7 | lmL M8 | (-2lnaL) | Sgnificance | Paml >95% | SLAC | FEL | MEME | FUBAR | Total no. of site | |
|---|---|---|---|---|---|---|---|---|---|---|---|---|---|---|
| **TLR27** | **2412** | 19 | 0.47030 | -14064,137823 | -14046,750334 | | | 3, 5, 13, 16, 17, 19, 22, 25, 26, 40, 42, 44, 46, 49, 51, 56, 75, 82, 85, 98, 108, 112, 154, 155, 166, 168, 192, 193, 201, 217, 221, 222, 234, 238, 239, 242, 252, 25257, 269, 272, 275, 277, 282, 284, 287, 293, 295, 297, 298, 305, 309, 310, 315, 321, 326, 329, 333, 339, 341342, 351, 355, 358, 366, 370, 372, 378, 380, 381, 393, 395, 397, 400, 401, 407, 410, 414, 423425, 426, 427, 428, 429, 436, 445, 448, 456, 476, 481, 487489, 504, 513, 518, 526, 544, 552, 554, 563, 565, 568599, 600, 607, 622, 631, 633, 634, 635, 638, 654, 668, 672, 759, 773790, 791, 793, 794, 795, 796, 799, 800802, | 802 | 13, 257, 287, 342, 351, 489, 631, 802 | 19, 40, 46, 132, 257, 287, 342, 351, 425, 459, 489, 529, 599, 631, 659, 768, 790, 802 | x | 14 | **Diversifying selection** |
| | | | | x | x | x | x | x | 14, 53, 54, 69, 79, 118, 124, 127, 145, 153, 158, 161, 176, 243, 261, 274, 318, 346, 363, 388, 394, 403, 444, 471, 473, 474, 482, 488, 532, 533, 559, 570, 610, 613, 617, 623, 632, 645, 652, 661, 669, 676, 679, 693, 701, 702, 704, 712, 731, 746, 747, 750, 758, 761, 762, 775, 778 | 14, 33, 37, 43, 53, 54, 69, 72, 76, 79, 89, 91, 118, 124, 125, 126, 127, 143, 145, 149, 153, 158, 161, 162, 176, 178, 182, 228, 230, 243, 246, 261, 273, 274, 302, 318, 320, 323, 337, 340, 344, 346, 347, 357, 363, 374, 379, 388, 394, 402, 403, 442, 444, 453, 455, 468, 470, 471, 473, 474, 479, 482, 488, 492, 495, 501, 532, 533, 534, 542, 549, 559, 564, 570, 571, 582, 583, 584, 588, 605, 610, 611, 613, 617, 621, 623, 625, 632, 645, 647, 650, 652, 661, 664, 667, 669, 674, 676, 679, 690, 693, 694, 695, 701, 702, 704, 712, 721, 727, 731, 735, 738, 740, 746, 747, 749, 750, 751, 758, 759, 761, 762, 775, 778, 801 | x | 14, 37, 43, 53, 54, 69, 72, 76, 79, 89, 118, 124, 126, 127, 143, 145, 149, 150, 153, 158, 161, 176, 178, 182, 213, 223, 243, 274, 318, 320, 323, 337, 344, 346, 347, 357, 363, 379, 388, 394, 403, 417, 442, 444, 453, 455, 460, 470, 471, 473, 474, 482, 488, 492, 495, 501, 532, 533, 534, 549, 559, 570, 571, 582, 583, 584, 588, 610, 611, 613, 617, 619, 623, 632, 645, 647, 652, 661, 664, 669, 674, 676, 679, 690, 693, 701, 702, 704, 712, 721, 731, 735, 738, 740, 746, 747, 750, 751, 758, 761, 762, 775, 778, 792, 801 | 99 | **Purifying selection** |

| Gene | No. aa residues | No. of species | ω (M0) | lmL M7 | lmL M8 | (-2lnaL) | Sgnificance | | Paml >95% | SLAC | FEL | MEME | FUBAR | Total no. of site | |
|---|---|---|---|---|---|---|---|---|---|---|---|---|---|---|---|
| **TLR29** | **2895** | 18 | 0.31571 | -23198,574795 | -23193,811633 | | | | 551, 718 | x | 52, 76, 131, 149, 216, 619, 948 | 51, 71, 73, 76, 83, 131, 149, 152, 178, 216, 257, 263, 288, 303, 317, 351, 419, 468, 525, 551, 613, 619, 799, 910 | x | 6 | **Diversifying selection** |
| | | | | x | x | x | x | | x | 21, 61, 67, 77, 84, 91, 92, 96, 106, 115, 116, 117, 124, 130, 133, 135, 141, 168, 174, 186, 187, 189, 206, 209, 212, 215, 217, 228, 236, 248, 249, 250, 256, 265, 280, 281, 284, 302, 308, 324, 338, 342, 347, 356, 363, 364, 365, 374, 397, 402, 409, 410, 411, 423, 424, 427, 429, 431, 435, 442, 446, 449, 450, 452, 469, 472, 473, 477, 485, 490, 496, 497, 498, 503, 508, 510, 511, 518, 542, 546, 548, 550, 559, 569, 570, 587, 590, 593, 603, 608, 611, 614, 625, 630, 631, 637, 654, 669, 670, 672, 675, 684, 691, 706, 707, 717, 719, 724, 729, 737, 741, 745, 747, 748, 752, 755, 758, 781, 782, 788, 790, 792, 795, 805, 806, 813, 814, 816, 819, 820, 831, 832, 833, 838, 840, 843, 847, 850, 855, 857, 858, 864, 866, 872, 873, 874, 875, 876, 878, 881, 884, 887, 888, 890, 891, 892, 897, 902, 911, 912, 914, 921, 923, 925, 929, 934, 937 | 10, 13, 17, 21, 34, 45, 61, 67, 77, 84, 85, 88, 91, 92, 96, 105, 106, 110, 112, 115, 116, 117, 120, 122, 124, 130, 133, 135, 136, 137, 139, 141, 146, 157, 160, 164, 166, 168, 174, 178, 179, 186, 187, 189, 195, 203, 206, 209, 212, 214, 215, 217, 220, 224, 225, 228, 229, 232, 236, 243, 244, 248, 249, 250, 256, 264, 265, 277, 280, 281, 284, 287, 295, 302, 308, 324, 338, 340, 342, 344, 347, 354, 356, 362, 363, 364, 365, 367, 373, 374, 375, 378, 390, 397, 402, 404, 405, 406, 409, 410, 411, 418, 421, 422, 423, 424, 427, 429, 431, 433, 435, 439, 442, 446, 448, 449, 450, 451, 452, 456, 459, 466, 469, 472, 473, 474, 476, 477, 479, 483, 485, 490, 493, 495, 496, 497, 498, 499, 503, 508, 510, 511, 512, 518, 520, 521, 535, 538, 541, 542, 546, 548, 550, 557, 559, 563, 569, 570, 571, 573, 582, 584, 585, 587, 589, 590, 593, 596, 599, 602, 603, 608, 611, 614, 620, 625, 626, 630, 631, 633, 636, 637, 638, 640, 643, 652, 653, 654, 663, 669, 670, 672, 675, 676, 684, 687, 690, 691, 700, 703, 704, 705, 706, 707, 710, 717, 719, 721, 724, 729, 737, 738, 741, 745, 746, 747, 748, 751, 752, 755, 756, 758, 760, 768, 774, 775, 781, 782, 783, 784, 785, 787, 788, 790, 791, 792, 795, 805, 807, 809, 813, 814, 815, 816, 819, 820, 823, 825, 827, 829, 831, 832, 833, 837, 838, 840, 843, 844, 845, 847, 848, 849, 850, 853, 854, 855, 856, 857, 858, 860, 862, 864, 865, 866, 872, 873, 874, 875, 876, 878, 881, 884, 887, 888, 889, 890, 891, 892, 897, 899, 902, 906, 911, 912, 914, 916, 917, 921, 923, 925, 927, 929, 932, 934, 936, 937 | x | 1, 6, 17, 21, 30, 45, 61, 67, 72, 74, 77, 82, 84, 85, 88, 89, 91, 92, 96, 105, 106, 107, 112, 113, 115, 116, 117, 120, 122, 124, 125, 130, 133, 135, 136, 137, 139, 141, 146, 156, 157, 159, 160, 164, 166, 168, 174, 186, 187, 189, 203, 206, 207, 209, 212, 215, 217, 220, 224, 225, 228, 232, 236, 243, 244, 248, 249, 250, 256, 259, 264, 265, 275, 280, 281, 284, 287, 302, 308, 319, 324, 331, 338, 340, 342, 344, 347, 356, 362, 363, 364, 365, 367, 369, 370, 373, 374, 375, 378, 390, 397, 402, 404, 405, 406, 407, 409, 410, 411, 418, 421, 422, 423, 424, 425, 426, 427, 429, 430, 431, 435, 438, 439, 442, 445, 446, 448, 449, 450, 451, 452, 454, 456, 459, 466, 469, 470, 472, 473, 474, 475, 476, 477, 484, 485, 487, 490, 493, 494, 495, 496, 497, 498, 499, 500, 503, 508, 510, 511, 514, 518, 520, 521, 522, 523, 531, 532, 535, 538, 541, 542, 546, 547, 548, 550, 559, 560, 563, 568, 569, 570, 571, 573, 574, 587, 589, 590, 593, 596, 598, 599, 601, 602, 603, 608, 610, 611, 614, 617, 620, 625, 630, 631, 633, 636, 637, 638, 640, 642, 643, 652, 653, 654, 655, 669, 670, 672, 675, 684, 687, 691, 700, 703, 704, 705, 706, 707, 709, 710, 713, 717, 719, 721, 724, 725, 727, 729, 737, 738, 741, 743, 745, 746, 747, 748, 751, 752, 755, 756, 758, 759, 760, 768, 772, 774, 781, 782, 783, 784, 785, 787, 788, 789, 791, 792, 795, 805, 806, 807, 812, 813, 814, 815, 816, 819, 820, 825, 827, 829, 831, 832, 833, 835, 838, 840, 841, 842, 843, 844, 847, 848, 849, 850, 853, 854, 855, 857, 858, 864, 865, 866, 872, 873, 874, 875, 876, 878, 881, 884, 885, 887, 888, 889, 890, 891, 892, 897, 899, 900, 901, 902, 904, 906, 911, 912, 914, 916, 917, 918, 921, 923, 924, 925, 926, 929, 931, 932, 934, 936, 937, 945 | 274 | **Purifying selection** |

**Supplementary Table 3.** Phylogenetic tests of selection.
